# Enhanced and Unified Anatomical Labeling for a Common Mouse Brain Atlas

**DOI:** 10.1101/636175

**Authors:** Uree Chon, Daniel J. Vanselow, Keith C. Cheng, Yongsoo Kim

## Abstract

Anatomical atlases in standard coordinates are necessary for the interpretation and integration of research findings in a common spatial context. However, the two most-used mouse brain atlases, the Franklin and Paxinos (FP) and the common coordinate framework (CCF) from the Allen Institute for Brain Science, have accumulated inconsistencies in anatomical delineations and nomenclature, creating confusion among neuroscientists. To overcome these issues, we adopted the FP labels into the CCF to merge two labels in the single atlas framework. We used cell type specific transgenic mice and an MRI atlas to adjust and further segment our labels. Moreover, new segmentations were added to the dorsal striatum using cortico-striatal connectivity data. Lastly, we have digitized our anatomical labels based on the Allen ontology, created a web-interface for visualization, and provided tools for comprehensive comparisons between the Allen and FP labels. Our open-source labels signify a key step towards a unified mouse brain atlas.

## Introduction

Anatomical delineation of the brain is critical for elucidation of the anatomical and functional organization of the brain across species^1–5^. Whole brain anatomical atlases provide a spatial framework for examining, interpreting, and comparing experimental data from different studies. For mouse, the most widely used animal model to understand the mammalian brain, the two most commonly used brain atlases are the Franklin-Paxinos atlas^6^(in short, called “FP” hereafter) and the Allen reference atlas^7,8^ (in short, called “Allen” hereafter). Both atlases are largely based on manual delineation by expert neuroanatomists using cytoarchitectonic features based on a variety of staining including Nissl and acetylcholine esterase antibody staining in 2D histological sections.

More recently, the Allen Institute for Brain Sciences released a 3D reference brain with 10 μm isotropic voxel resolution, called the common coordinate framework (CCF)^9^. This new reference brain marks a significant departure from classical neuroanatomy based on 2D sections and provides an excellent platform for the registration of 3D mouse brain imaging datasets collected from emerging high resolution whole brain imaging modalities such as serial two-photon tomography and light sheet microscopy^5,10–12^. More importantly, the CCF facilitates the integration and sharing scientific data from different studies in a common spatial context^13^. The accompanying anatomical labels have smooth delineation across all 3D planes, which enable easy views of 3D perspective of brain regions.

Unfortunately, significant discrepancies exist between the anatomical labels on the Allen CCF and the FP labels. For example, these two atlases often have disagreed anatomical borders and 3D coordinates as well as different nomenclatures for same structures^14,15^.

To make it worse, the latest labels in the CCF released in 2017 also introduced significant changes from its original Allen labels that were based on 2D Nissl stained sections. This has created confusion and mis-interpretation of experimental results^16^. These issues motivated us to create unified and highly segmented anatomical labels in the adult mouse brain based on the CCF. We decided to use the FP labels for our initial anatomical labels because it represents the highest degree of the segmentation in the adult mouse brain, and because a huge body of prior research is based on the FP labels. Here, we adopted the FP labels into the CCF by rigorous alignment using an MRI based atlas and cell type specific transgenic mice marking for distinct anatomical areas^17,18^. We also further segmented labels where cell types could be distinguished within single anatomically defined regions. The resulting labels create a unique opportunity for comprehensive comparisons between the two most commonly used anatomical labels in a common CCF space. Furthermore, we used topographically distinct cortico-striatal projection patterns to add segmentations to the dorsal striatum, which is unsegmented in existing atlases.

Lastly, we have digitized the anatomical labels based on the Allen ontology to facilitate integration of our labels as a neuroinformatics tool^9^. Digitized labels combined with image registration can serve as a powerful tool to automatically quantify signal of interest across whole brain regions in a reference brain^9,10,19,20^. To facilitate its usage, we have made all our newly established digital map data freely available for viewing and downloading from our web-based atlas implementation at http://kimlab.io/brain-map/atlas/.

## Result

### Importing Franklin-Paxinos anatomical labels into the Allen common coordinate framework

We used the FP labels drawn in 2D histological sections for our initial template segmentation. We first imported vector drawing map of FP labels to the Allen CCF (Figure 1A-B). Automated image registration of 2D Nissl sections from the FP atlas to the CCF has been challenging due to different background contents between the two atlases and non-uniform tissue distortion between histological sections in the FP labels. Thus, we used manual adjustment to initially align the FP labels on the CCF coronal sections with 100 μm *z* spacing based on the autofluorescence signals of distinct anatomical features (Figure 1B – C, yellow arrows as examples). Autoflourescent background in the CCF provides rich anatomical information in both cortical and subcortical regions. For example, distinct contrast in the barrel field provides strong evidence to delineate layer 4 for the somatosensory barrel cortex (Figure 1B-C, red arrows).

**Figure 1.**
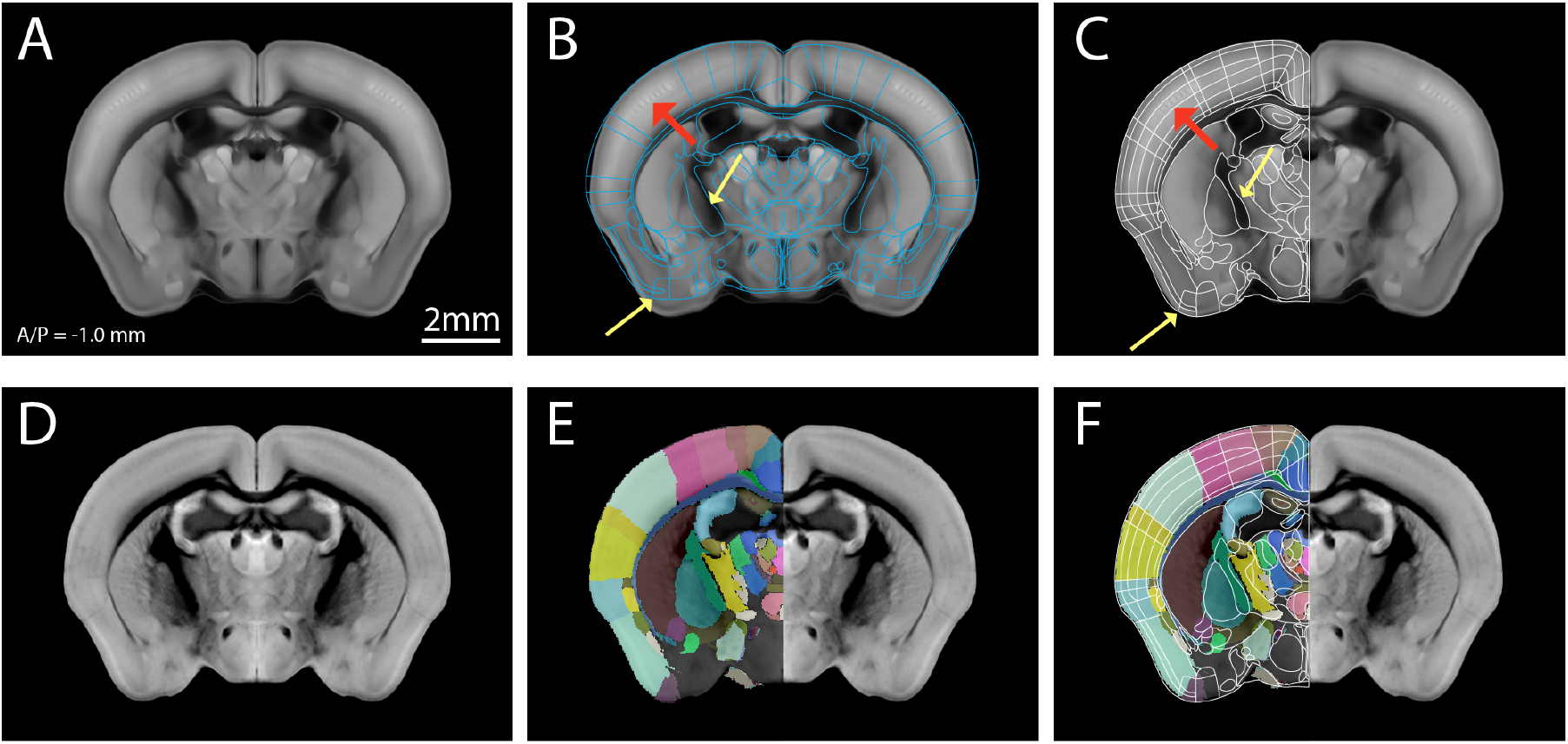
Initial import and alignment of the Franklin-Paxinos labels onto the Allen Common Coordinate Framework. (A) The Allen Common Coordinate Framework (CCF) that serves as base anatomical platform. A/P represent Bregma anterior/posterior coordinate. (B) Initial import of the Franklin-Paxinos (FP) vector labels in the CCF. (C) Manual alignment based on anatomical features in the CCF. Yellow arrows highlight distinct anatomical boundaries based on edges and white matter track. Red arrows indicate layer 4 in the somatosensory barrel cortex. (D) MRI images registered to the same CCF plane in (A). (E) Original FP based labels drawn in the MRI atlas registered to the CCF. Lack of labels in hypothalamic and amygdala regions are due to missing labels in the original MRI labels. (F) Further adjustment of anatomical delineation (white lines) based on the MRI labels.

To further assist 2D label alignment in the context of contiguous 3D planes, we used a high resolution magnetic resonance imaging (MRI) atlas with the FP labels in most brain regions^17,21,22^. We first registered the MRI reference brain to the CCF and transformed the MRI labels to fit in the CCF (Figure 1D-E). Although the MRI labels are not as detailed as Nissl based FP labels, it provided an independent way to align and to further adjust our initial alignment in 3D space (Figure 1F). The MRI labels were particularly useful to align segmentations in the isocortex (also called “neocortex”) (Figure 1F).

### Fine label alignment and further segmentations using cell type-specific transgenic mice

Previously, histological staining with specific markers (e.g., acetylcholine esterase, or parvalbumin) on 2D sections has been used to guide detailed delineation in anatomical regions^6^. We utilized a similar approach using 14 different transgenic mouse lines that mark specific neuronal subtypes^10,18,23^ (called “marker brain” hereafter) to highlight anatomical boundaries otherwise often not visible in the CCF tissue autoflourescent background. Marker brains imaged by STPT were registered to the CCF, and their signals were overlaid in the CCF to highlight cell type based anatomical features (Figure 2 and S1, Table S1). For example, Choline acetyltransferase (Chat)-Cre mice crossed with Cre dependent reporter mouse expressing nuclear tdTomato (Ai75) were used to delineate brain regions enriched with cholinergic neurons such as the basal forebrain and the hindbrain areas (Figure 2A)^24^. Parvalbumin (PV)-Cre crossed with Cre dependent reporter mouse expressing nuclear GFP (H2B-GFP) has been very useful to delineate structures in the thalamus, midbrain, and hindbrain (Figure 2B)^6,25^. Somatostatin (SST)-Cre crossed with the H2B-GFP reporter mouse has been useful for amygdala, hypothalamus, olfactory regions, and subcortical regions, such as the bed nucleus of the stria terminalis (BST) (Figure 2C)^26^. Oxytocin receptor (OTR)-Cre crossed with Cre dependent reporter mouse expressing tdTomato (Ai14) highlighted selected brain regions including dorsal endopiriform nucleus (DEn), CA2 in the hippocampus, amygdala and entorhinal regions (Figure 2D)^27^. Lastly, we used cortical layer specific Cre mice crossed with Ai75 to validate our cortical layer (L) delineation. We used Ctgf-Cre for L6b, Ntsr1-Cre for L6, Rbp4-Cre for L5, and Cux2-Cre for L2/3 (Figure 2E and S1)^28^. Additional marker brains were utilized to delineate several more brain regions. For example, Ctgf-Cre was further used for delineations of DEn and structures of thalamus, amygdala, and isocortical areas (Figure S1). The full list of marker brains and their expression in anatomical regions is summarized in Table S1.

**Figure 2.**
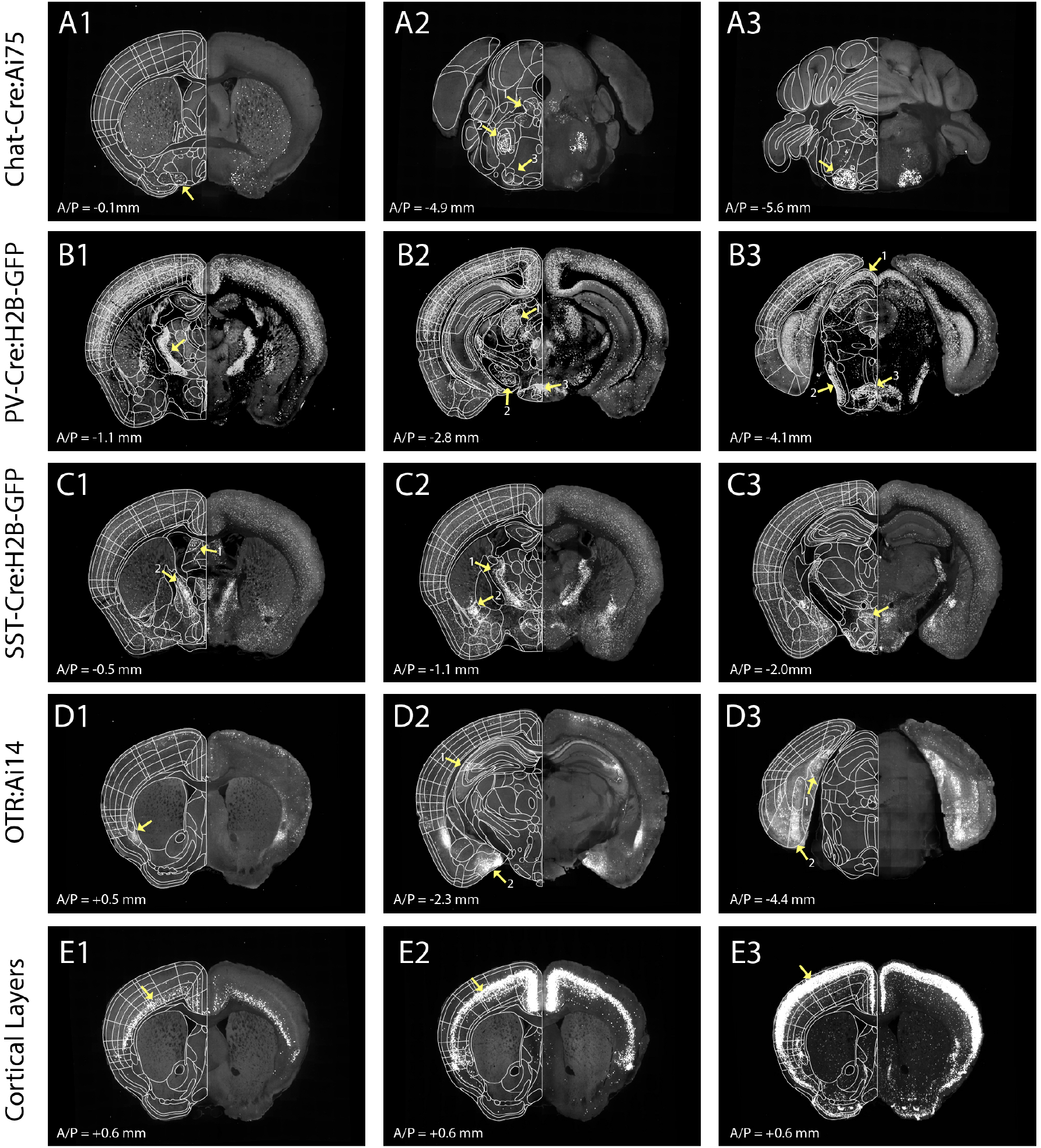
Marker brains to assist further alignment of anatomical labels. (A-E) Examples of different marker brains registered to the CCF that helped to align subregions as highlighted with yellow arrows. A/P represent Bregma anterior/posterior coordinates. (A) Chat-Cre:Ai75 brain to delineate (A1) the basal forebrain structures including the nucleus of the horizontal limb of the diagonal band (arrow). It was also used to delineate (A2) midbrain areas, including the laterodorsal tegmental nucleus (arrow1), themotor trigeminal nuclei (arrow 2), the lateral superior olive (arrow 3), and (A3) the facial nucleus (arrow). (B) PV-Cre:H2B-GFP brain to delineate (B1) the reticular nucleus (arrow), (B2) th anterior pretectal nucleus (arrow 1), the substantia nigra, reticular part (arrow 2), and the retromamillary nucleus (arrow 3) as well as (B3) the superficial gray layer superior colliculus (arrow 1), the ventral nucleus of the lateral lemniscus (arrow 2), and the reticulotegmental nucleus of the pons, pericentral part (arrow 3). (C) SST-Cre:H2B-GFP brain to delineate (C1) the cerebral nuclei, such as the lateral septal nucleus, dorsal part (arrow 1) and the bed nuclei of the stria terminalis medial division posteromedial part (arrow 2), (C2) the reticular nucleus (arrow 1) and the central amygdaloid nuclei (arrow 2), and (C3) hypothalamic structures, such as the dorsomedial hypothalamic nuclei dorsal and ventral parts (arrow). (D) OTR:Ai14 brain to delineate (D1) the dorsal endopiriform nucleus (arrow), (D2) CA2 (arrow 1), the posteromedial cortical amygdala (arrow 2), and (D3) the caudomedial entorhinal cortex (arrow1) as well as the postsubiculum (arrow 2). (E) Cortical layers defined by (E1) Ntsr-Cre:Ai75 for layer 6, (E2) Rbp4-Cre:Ai75 for layer 5, and (3) Cux2-Cre:Ai75 for layer 2/3.

While utilizing marker brains, distinct cell populations were observed within specific substructures. We used this information as a way to further segment structures in the thalamus, hypothalamus, and hindbrain. For example, using PV-Cre and Ctgf-Cre marker brains, ventral posteromedial nucleus of the thalamus (VPM) was further segmented into dorsal and ventral parts (VPMd and VPMv, respectively) (Figure 3A). We observed densely packed cell population in VPMd in both lines, contrasting the loosely scattered cells in VPMv (yellow arrows in Figure 3A). Using OTR-Cre and Ctgf-Cre marker brains, posterior hypothalamic nucleus (PH) was segmented into nuclear dorsal and ventral parts (PHnd and PHnv, respectively) with higher expression in PHnd (Figure 3B). Lastly, medial vestibular nucleus, parvicellular part (MVp) was further divided into dorsal and ventral parts (MVpd and MVpv, respectively) based on density difference from SST-Cre and PV-Cre marker brains (Figure 3C). We added 10 new subdivisions (Table S2).

**Figure 3:**
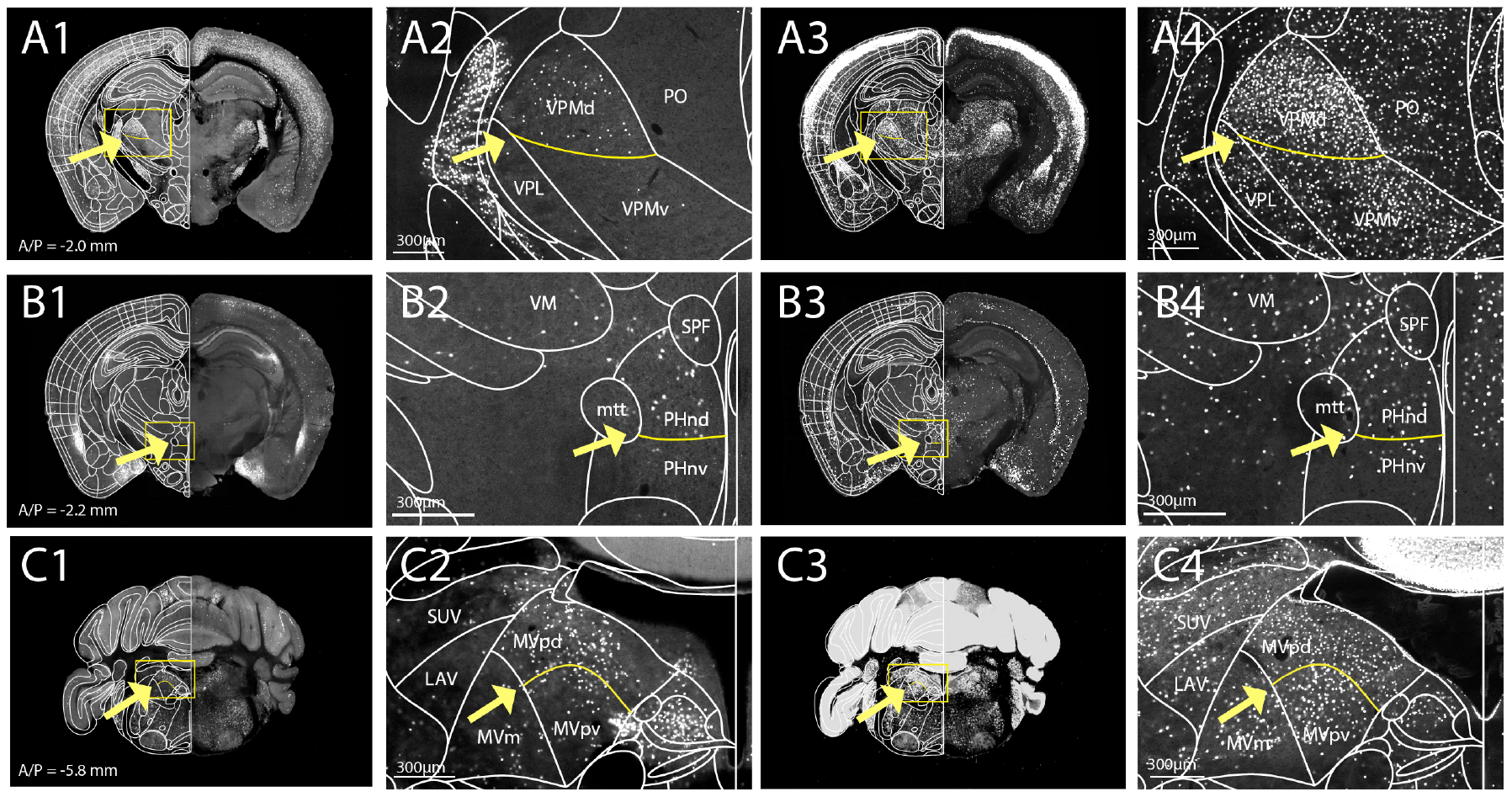
Additional segmentations based on distinct expression from marker brains. (A-C) Examples of marker brains to further segment structures. New segmentations are marked by yellow lines. (A) PV-Cre:H2B-GFP (A1-2) and Ctgf-Cre:Ai75 (A3-4) marker brains were utilized to further segment ventral posteromedial nucleus of the thalamus (VPM) to dorsal and ventral parts (VPMd and VPMv, respectively). (B) OTR-Cre:Ai14 (B1-2) and Ctgf-Cre:Ai75 (B3-4) used to segment dorsal and ventral parts (PHnd and PHnv, respectively) of the posterior hypothalamic nucleus (PHn). (C) SST-Cre:H2B-GFP (D1-2) and PV-Cre:H2B-GFP (D3-4) used to segment the medial vestibular nucleus, parvicellular part (MVp) to dorsal and ventral parts (MVpd and MVpv, respectively). See Table S2 for full names of acronyms.

### Long-range projection based anatomical segmentation

Previously, anatomical segmentations were largely based on cytoarchitectonic features^6 7^. Although highly useful, this approach cannot be applied to the dorsal striatum without such features. Thus, dorsal striatum remains unsegmented in both FP and Allen atlases despite its prominent size and heterogeneous functions in the brain. Recent studies has shown that different parts of the dorsal striatum receive topographically distinct cortical inputs^29–31^. We decided to use a similar approach to segment the dorsal striatum based on distinct cortico-striatal projections. We downloaded 129 datasets with anterograde tracing using C57bl/6 mice covering the entire isocortical areas from the Allen connectivity dataset^20^ and registered all of these brains to the CCF (Figure 4A-B). Then, we averaged the projection datasets from the 10 different cortical regions for each anatomically distinct dorsal striatum projection pattern (Figure 4B). We superimposed the projection dataset on the CCF and delineated different striatal areas based on cortico-striatal projection data (Figure 4B). We observed different striatal regions with either distinct input from one cortical group or convergent inputs from multiple regions, which is consistent with previous studies^29,30^(Figure 4C-D). We added new delineations to the existing labels (Figure 4D, Table S2).

**Figure 4:**
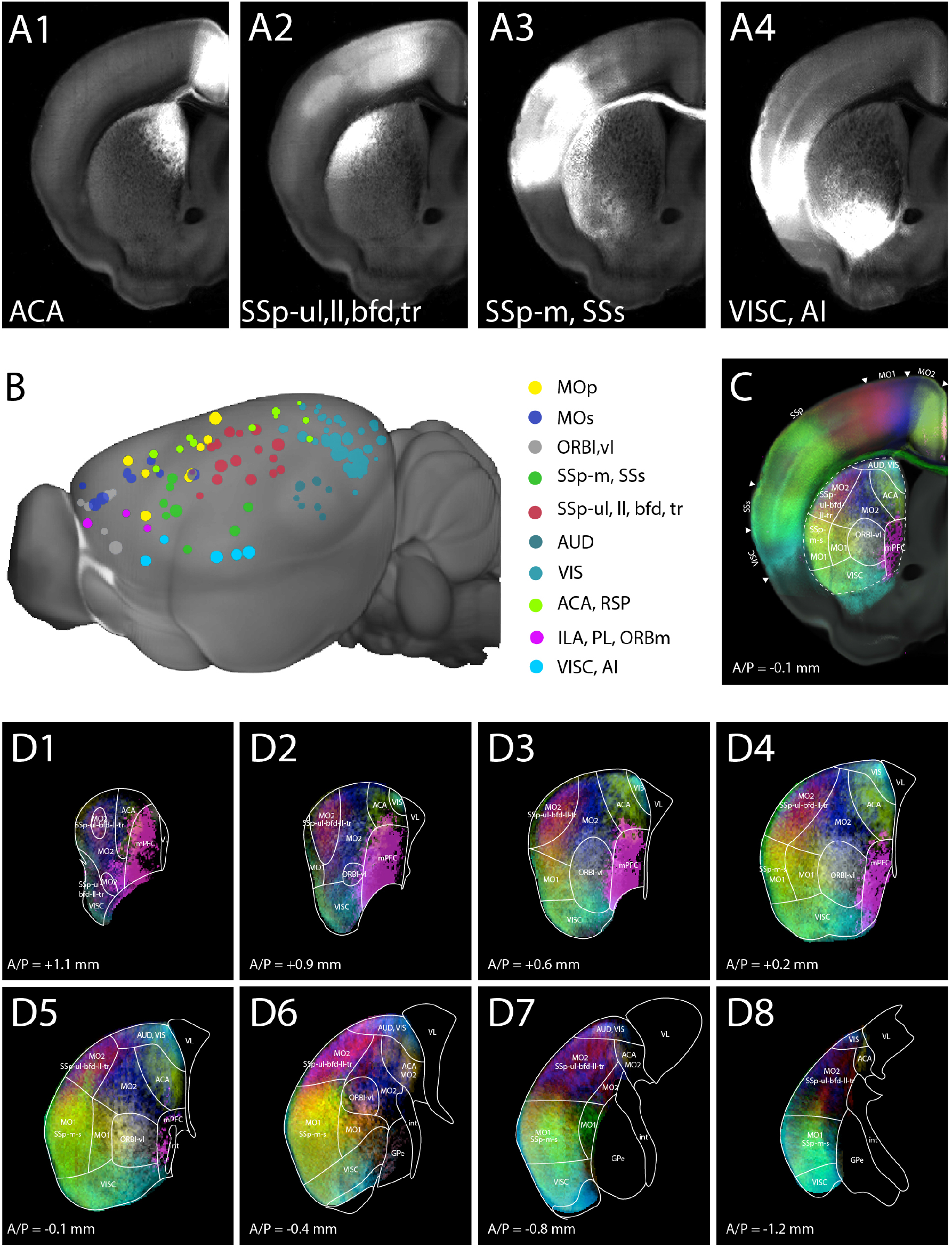
Cortico-striatal projection based striatum segmentations. (A) Anterograde tracing datasets from different cortical domains registered into the CCF. A1 for the anterior cingulate cortex (ACA), A2 for the primary somatosensory cortex (SSp), upper limb (ul), lower limb (ll), barrel field (bfd), and trunk (tr) area. A3 for the SSp, mouth (m) and secondary (s). A4 for the visceral (VISC) and the agranular insular cortex (AI). (B) 129 datasets clustered into 10 groups based on cortical input regions. Datasets in the same cluster have the same color. (C) Example of striatal segmentation based on cortico-striatal projection patterns. (D) Representative images of new dorsal striatum segmentations throughout several Bregma A/P planes. Full name of acronym can be found in Table S2.

### Digitization and hierarchical organization of anatomical labels

Digital atlases with distinct label values for each anatomical region have been very useful neuroinformatics tools to automatically quantify target signals in different anatomical regions when combined with image registration^10,19^. Thus, we assigned a unique ID in each label (Figure 5A-C). We adopted and arranged numerical IDs for each structure in a hierarchical manner based on the Allen ontology (Figure 5E)^8^. In the digitization process, we first found comparable brain regions between the FP and the Allen labels. To accommodate the higher degree of segmentation in the FP labels, 471 more structure IDs were created (Table S2). For example, PAG consists of several subdivisions that plays various functions including expression of fear behavior^32^. PAG, which is considered as a single structure in the Allen labels, is further segmented into dorsomedial, lateral, dorsolateral, ventrolateral, pleoglial, and p1 divisions (DMPAG, LPAG, DLPAG, VLPAG, PlPAG, and p1PAG, respectively) in FP labels. The boundaries of the subdivisions were delineated by observing cell density differences between each division with SST-Cre expression (Figure 5D1). Each subdivided region was given new unique numerical IDs and assigned within its parent structures (Figure 5D2, E).

**Figure 5:**
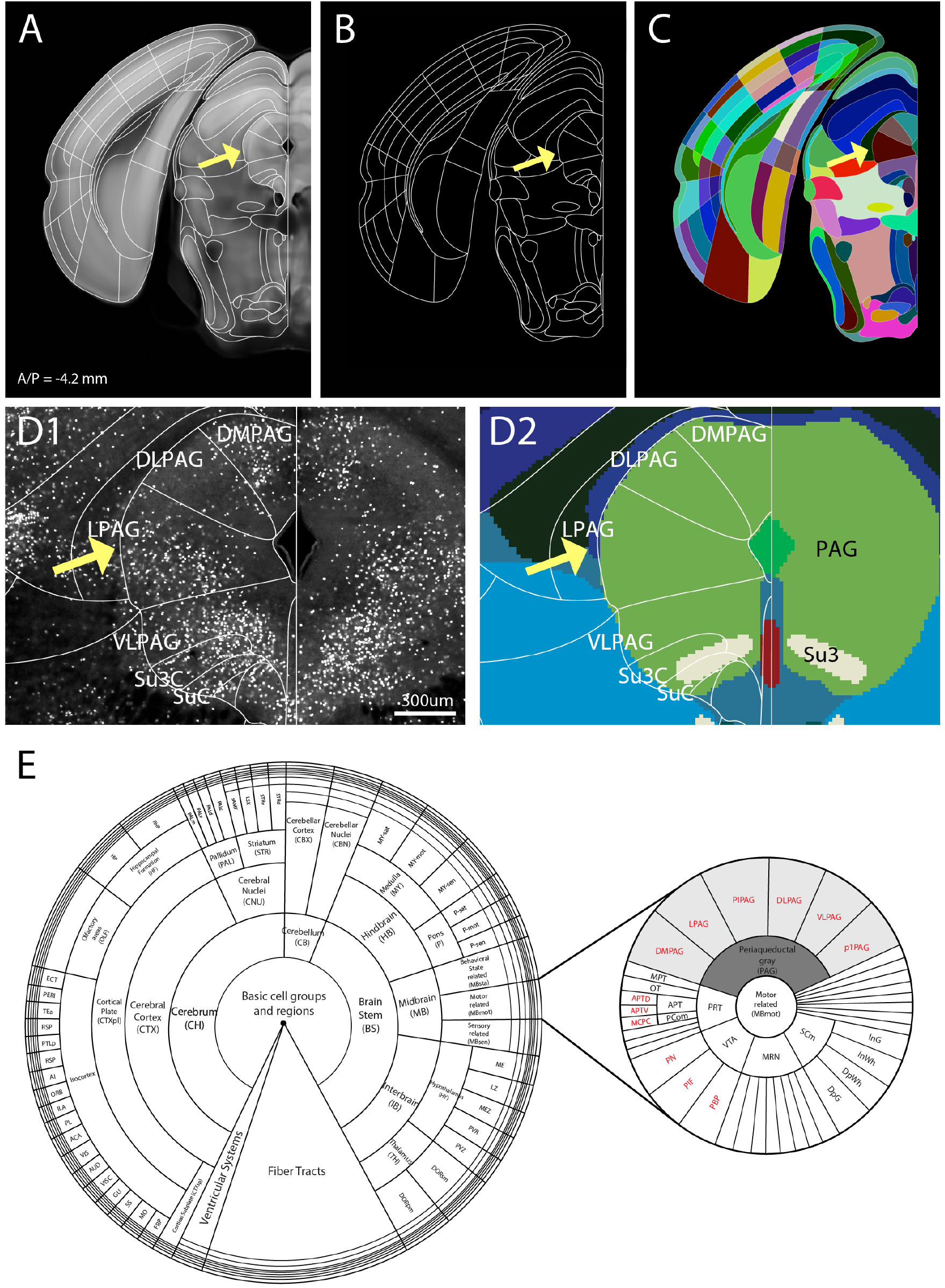
Digitization of anatomical structures. (A) Example of our highly segmented label on the CCF. Yellow arrows highlight the lateral subdivision of periaqueductal gray (PAG). (B) Exported delineation lines. (C) Digitization of labels with unique numerical ID for each anatomical structure. Different color of each structure pertains to different number. (D) SST-Cre:H2B-GFP showed distinct subregions in PAG with different cell density level. (D1) Our labels (white font) divide PAG into 6 different subregions, as can be seen with the specific enrichment of SST neurons in DLPAG and LPAG (yellow arrow). (D2) In contrast, the Allen labels (color labels in the background) showed only 2 segmentations within the PAG (black font). (E) Hierarchical organization of anatomical labels based on the Allen ontology. Numerical IDs of individual structures assigned within parent structures for region-level and individual structure-level data analysis. For example, the PAG (shaded dark gray) is the parent structure of 6 subdivided structures (shaded light gray). Red font labels refer to structures further divided by the FP labels that are not present in the Allen labels.

Since the nomenclature and abbreviations in same structures are often different between the FP and the Allen labels, we systematically compared between the two labels. For example, cingulate cortex, area 24b (A24b) in the FP labels matches to the anterior cingulate area, dorsal part (ACAd) in the Allen labels. We included the complete list of comparisons between the two labels, unique brain region IDs, and hierarchical arrangement in Table S2. This information can be utilized to compare the nomenclature within any brain regions between the two atlases.

### Comparison between Allen and FP based anatomical labels

Because our anatomical labels adopted from the FP labels were aligned in the Allen CCF, we can compare and contrast difference between two most commonly used anatomical labels in the same space (Figure 6). We also included the original Allen labels drawn in Nissl stained sections as additional comparison (last column of Figure 6). Our labels have overall finer segmentations than the Allen labels. For example, the zona incerta (ZI) is a part of subthalamic nucleus that plays an important role in several behaviors such as pain processing and defensive behavior^33,34^. We previously found that parvalbumin (PV) neurons are heavily enriched in ventral ZI^10^. The FP labels segmented PV enriched ventral ZI separately from dorsal ZI while both the original and the new Allen labels have only one segmentation for ZI (Figure 6A). Moreover, Allen and FP labels often use different boundaries even in similar brain regions. For example, substantia innominata (SI) in the Allen labels is a part of the basal forebrain structure that is important in attention and learning^35,36^. In the FP label, the matching region is composed of ventral pallidum (VP), substantia innominata basal (SIB), and extended amygdala (EA). In our marker brains, VP and EA are marked by cholinergic and somatostatin neurons, respectively (Figure 6B)^24^. Moreover, large portion of EA was included as a part of lateral preoptic area (LPO) in the new Allen labels (but not in the original labels), which does not match with our border between hypothalamus and basal forebrain (yellow arrows in Figure 6B). Discrepancies between anatomical borders extend to many different areas including cortical areas. For example, we noticed that boundary between motor and somatosensory cortex in the latest Allen labels has been dramatically shifted from its original label (yellow arrows in Figure 6C). Our labels match better to the original Allen labels than to the latest version, consistent with the existence of layer 4 in the somatosensory area, but not in the motor area, and with patterns of cortical layer specific marker brains (Figure 6C). Moreover, the latest Allen labels simplified segmentation in some key regions that are functionally subdivided. For example, the bed nucleus of the stria terminalis (BST) in the original Allan atlas was divided into different subregions, but is no longer subdivided in the new atlas (Figure 6D4). BST subdivisions play important roles in distinctive behaviors (e.g., anxiety and social behavior) and have unique anatomical connections^37–41^. Our labels are highly segmented in the BST (Figure 6D).

**Figure 6:**
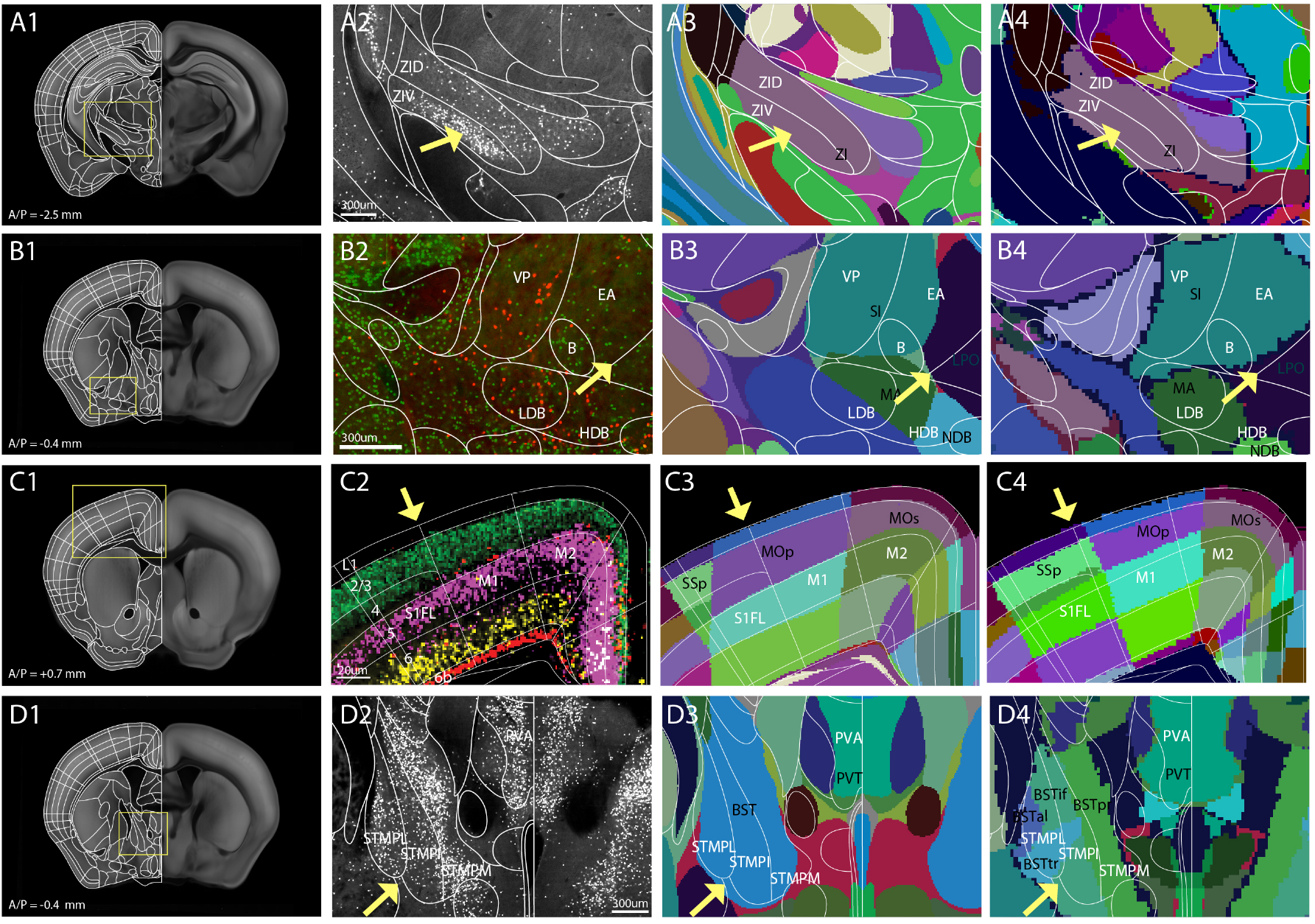
Comparison between the Allen and our new labels. The first column: Our highly segmented labels on the Allen CCF, The second column; our labels (white lines) with marker brain background, The third column: comparison between our labels and the latest Allen labels (colored background), The fourth column: comparisons between our labels and the original Allen labels (colored background). (A2 – D4) Anatomical names in black and white are from the Allen and our labels, respectively. (A2-4) PV-Cre:H2B-GFP (A2) to identify subregions in zona incerta (ZI). Low in dorsal and high in ventral parts (ZID and ZIV, respectively) in our labels while the original and the new Allen labels have a single combined structure for ZI. (B2-4) (B2) Virtual overlay of Chat-Cre:Ai75 (red) and SST-Cre:H2B-GFP (green) to compare basal forebrain regions. (B3-4) Our labels further segregate the single structure defined as the substantia innominate (SI, Allen) into the ventral pallidum (VP) and the extended amygdala (EA). Yellow arrow highlights border between basal forebrain and hypothalamus. (C2-4) Disagreed borders between the somatosensory and the motor cortices. Yellow arrow highlights border between the somatosensory and motor cortices. (C2) Virtual overlay of pseudo colored Cux2:Ai75 (L2/3, green), Rbp4:Ai75 (L5, magenta), Ntsr1:Ai75 (L6, yellow), and Ctgf:Ai75 (L6b, red). Note the lack of Cux2:Ai75 and Rbp4:Ai75 signal in the layer 4 of the somatosensory cortex. (D2-4) The BST is divided into several subregions in our labels compared to a single structure of the BST in the new Allen labels, despite the original version with finer delineations for this structure. See table S2 for the abbreviation.

### Web-based atlas visualization and sharing

The web-visualization platform for digital atlases enables easy identification of anatomical labels across different sections and comparison across different atlases^3,13^. Thus, we created a website (http://kimlab.io/brain-map/atlas/) to visualize and share our anatomical labels. The web visualization includes easy identification of anatomical labels in the background CCF. All labels and associated files are freely available for downloading (Supplementary File 1 and 2). This open source data sharing will facilitate to further refinement of anatomical labels and to integrate data interpretation within this single anatomical platform.

## Discussion

Here, we present highly segmented open source anatomical labels on the Allen CCF, which are easily accessible via our website. Our labels are largely based on FP labels with new cortico-striatal projection based segmentations in dorsal striatum and further segmentations based on fluorescent transgenic markers.

A reference atlas serves a critical role in understanding spatial context of the brain^8,10,42,43^. However, independently generated atlases with different nomenclature and boundaries can make it difficult to integrate data from different studies^13^. Significant effort has been made to standardize a rodent brain atlas as a key neuroinformatics tool to facilitate data exchange and to enhance reproducibility between different studies^3,13,44^ For example, the International Neuroinformatics Coordination Facility established digital atlas infrastructure for a common spatial framework such as the scalable brain atlas under FAIR (Findable Accessible Interoperable Reproducible) principles^13,44^. Recently, the Allen CCF generated from iterative averaging of over 1000 different mouse brain samples provides the highest resolution 3D digital atlas platform^9^. There has been significant encouragement by funding agencies (e.g., BRAIN initiative) to use the CCF as a common anatomical framework for functional and anatomical studies to facilitate seamless exchange between results from different studies^45^. To further support this trend, new computational tools are being developed to integrate individual datasets (e.g., 3D imaging or even 2D histological sections) in the standard atlas framework^37–39,46^. While the CCF provides an ideal atlas platform with high resolution 3D images, its associated anatomical labels released in 2017 have been controversial due to fewer fine segmentations and significant changes in their anatomical borders from the original version. Moreover, inconsistencies in borders and nomenclatures compared to the widely used FP labels make it difficult to compare findings from studies that use different atlases. Our labels are ideally suited to resolve the issues.

Our strategy was to establish the FP based anatomical labels in the Allen CCF. We used a series of steps to rigorously align the FP labels in the Allen CCF. We further generated finer segmentations based on marker brains that highlight specific anatomical regions otherwise not visible in the background^18^. These strategies enabled us to establish highly detailed FP based labels in the Allen CCF. Our systematic comparison between the two atlases marks an important first step towards a unified anatomical label in a common atlas platform. As neuroscience research becomes increasingly collaborative, it is essential to have consistency in anatomical labels to specify regions of interest. By integrating FP based labels in the CCF, our labels can be used to facilitate the comparison of anatomical interpretations from past and future studies regardless of the atlas used.

We also used an cortico-striatal long-range connectivity to finely segment the dorsal striatum. Projectome-based atlasing provides an alternative way to segment brain regions that do not have distinct cytoarchitectonic features. Since brain-wide projectome data are becoming increasingly available in open source platforms^20,47–49^, similar approaches can be used to segment other brain regions with distinct projection patterns. Moreover, since this anatomical connectivity is related to functional interactions between neural circuitry, connectivity based anatomical segmentation can provide a unique opportunity to integrate functional circuit in the anatomical map.

Our digitized anatomical labels can be easily integrated into data processing pipelines to automatically quantify target signals throughout anatomical regions in the whole brain. We previously built such a pipeline to quantitatively map neural activity based on c-Fos induction, GABAergic cell subtypes, and long-range neural connectivity^10,19,48^. Moreover, mapping pipelines are increasingly available for high-resolution 3D image data and histological sections^11,50,51^. With image registration to the CCF, our digitized labels can serve as an invaluable neuroinformatics tool to examine target signals in the FP based labels as well as the built-in Allen CCF labels.

Moving forward, by integrating two most popular brain segmentations in the same 3D anatomical context, it will help to build unified anatomical labels for the mouse brain in the future^3,13,52^. To facilitate such work, we are making all the data freely available to visualize and download via our website. We envision that similar approaches can be taken to integrate independently generated atlases within animal species including humans.

## Material and Methods

### Animals

All animal work has been approved by the Institutional Animal Care and Use Committee of Penn State University College of Medicine. We used following transgenic mice to fluorescently label specific cell types (marker brains). For Cre drivers, we used OT-Cre (Jax: 024234), Avptm-Cre (Jax: 023530), OTR-Cre (gift from Nishimori lab, Tohoku University). For Cre dependent reporter mice, we use Ai14 (Jax:007908). We crossed cell type specific Cre driver mice with Ai14 to create maker brains. We used both male and female mice at ~2 - 3 months old. All mice were group housed in 12/12 light/dark cycle (6am light on, 6pm off) with access to food and water ad libitum. Other marker brains were downloaded from either publically available BICCN dataset or previously published database^10^. Because we observed highly stereotypical expression in each marker brain, we used one representative brain per each marker line for our anatomical work. The complete list of the maker brain with their source is listed in the Table S1.

### Sample preparation and imaging of cell type specific transgenic mice

Transgenic mice were perfused by using cardiac perfusion with 0.1M phosphate buffer (PB) followed by 4% paraformaldehyde (PFA). Brains were post-fixed with 4% PFA at 4°C overnight and transferred to 0.05M PB until imaging. Detailed protocol for the STPT imaging was described previously^10^. Briefly, a fixed brain was embedded in oxidized 4% agarose and cross linked by 0.05M sodium borate buffer at 4°C overnight. We used Tissuecyte 1000 (Tissuevision) to perform serial two-photon tomography imaging. We used 970nm wavelength laser and acquired a series of images at 1 μm X-Y resolution in every 50 μm *z* sections. We used custom-built algorithms to reconstruct the whole brain. Our imaged brains and downloaded marker brains were registered to the CCF using open source program (Elastix)^53^ as described previously^10^.

### Importing and modifying the FP labels to the Allen CCF

We originally obtained vector drawing of Nissl 2D section from Paxinos and Franklin’s the Mouse Brain in Stereotaxic Coordinates, 3^rd^ edition^6^. We also used the 4^th^ version to incorporate latest updated labels. We used a vector drawing tool (Adobe Illustrator) for our label work. We downloaded the Allen CCF and associated labels from the Allen Institute for Brain Sciences API (http://help.brain-map.org/display/mousebrain/API), and generated coronal slices (10 μm isotropic) using Image-Stacks-Reslices in FIJI (NIH). This produced 1320 Z coronal slices. Then, we selected one coronal slice in every 10 slices from Z95 to Z1315 using Image-Stacks-Tools-Make Substack in FIJI, generating 123 coronal images with 100 μm *z* spacing. We identified matching *z* planes between the FP atlas and the CCF using distinct anatomical landmarks (e.g., fiber track, and ventricles). To aid our label alignment in 3D, we downloaded MRI labels from different brain regions from publically available database (https://imaging.org.au/AMBMC/AMBMC). We combined labels from different brain regions to reconstruct the MRI labels using FIJI (NIH). Then, we registered the MRI atlas with the FP based labels to the CCF using Elastix. The MRI labels were particularly useful to align boundaries in cortical areas. We loaded cell type specific labeling from different transgenic mice and MRI labels as separate layers on the Illustrator, and used the information to further adjust anatomical delineations. To accommodate the FP labels (mostly 120 μm *z* spacing) in 100 μm *z* spacing, we used 5^th^ section of every 6 FP labels twice in the initial alignment and used the MRI atlas and marker brains to further modify the labels across the 3D plane. Once the FP labels were imported in the matching plane of the CCF on Adobe Illustrator, we used linear translation to stretch the FP labels to fit the CCF roughly. Then, we performed finer alignment manually based on specific landmarks of the brain with distinct contrast (e.g., fiber tracts). In selected areas (e.g., hypothalamus), boundaries were removed entirely and re-drawn based on key features of the CCF and distinct cell populations. In caudal areas, we often used 2-3 different FP planes to create hybrid labels to fit the CCF background as well as cell type specific features of the selected plane.

### Cortico-striatal projection based segmentation in dorsal striatum

We downloaded 129 datasets with anterograde virus injection in different cortical areas from C57bl/6 mouse line using Allen connectivity database (http://help.brain-map.org/display/mouseconnectivity/API). All downloaded datasets were registered to our modified CCF with 100 um *z* spacing using Elastix. After the image registration, we removed the autoflourescent background of each sample using binary thresholding (FIJI). We clustered projection dataset into 10 groups based on their cortical injection sites and averaged projection signals in the same group using FIJI. Then, we imported the projection data into Illustrator as separate layers and used them to further segment the dorsal striatum.

### Digitization of anatomical labels

Our labels were first compared to segmented regions of the Allen labels. We used ontologically arranged Allen label numbering system as a template to digitize our labels (Table S2). All labels were imported onto FIJI, and each region was selected using wand tool and assigned specific anatomical identification numbers using the Process-Math-Add function. If our labels matched the Allen labels, we assigned the same Allen anatomical identification numbers. If our labels were not found in the Allen labels (e.g., finer segmentation in our labels), we assigned new unique identification numbers. If there was significantly disagreed border delineation of matching structures with similar nomenclature, we maintained the same ID number for that specific structure.

## Acknowledgement

We thank Rhea Sullivan in helping generating cortico-striatal projectome data, and Pavel Osten and Piotr Majka for critical reading and editing the manuscript. This publication was made possible by a NIH grant (R01MH116176) and Tobacco Cure Funds from the Pennsylvania Department of Health to Y.K. and facilitated by NIH grant 1R24OD18559-01-A2 to K.C. Its contents are solely the responsibility of the authors and do not necessarily represent the views of the funding agency.

## Contributions

Conceptualization, Y.K.; label alignment and digitization, U.C.; Dorsal striatum segmentation, Y.K.; Web visualization, D.V.; Manuscript preparation, Y.K., U.C., K.C.

---

**Figure S1, Related to Figure 2:**
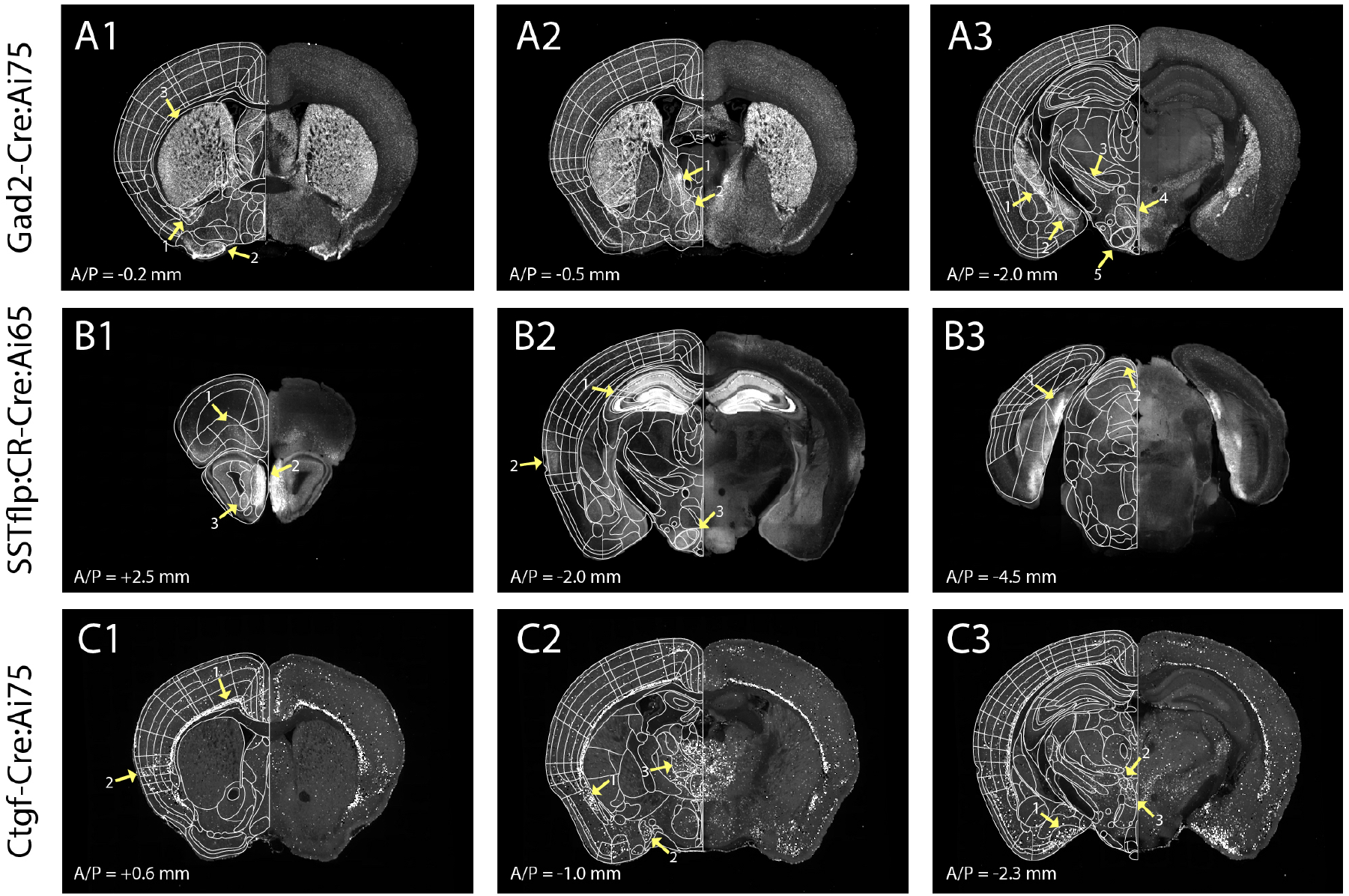
Additional marker brains used for alignment of anatomical borders. (A) Gad2-Cre:Ai75 brain to delineate (A1) the interstitial nucleus of the posterior limb of the anterior commissure (arrow 1), the olfactory tubercle (arrow 2) and the caudate putamen (arrow 3), (A2) the bed nuclei of the stria terminalis medial division posteromedial part (arrow 1), the striohypothalamic nucleus (arrow 2), and (A3) the central amygdaloid nuclei (arrow 1), the medial amygdala posterodorsal division (arrow 2), the zona incerta (arrow 3) and other hypothalamic structures such as the dorsomedial hypothalamic nucleus, ventral part (arrow 4) and the ventromedial hypothalamic nucleus (VMH, arrow 5). (B) SSTflp:CR-Cre:Ai65 transgenic mouse line to delineate following structures: (B1) the orbital cortex area (arrow 1), layers of the olfactory bulb (arrow 2), the anterior olfactory area, ventral part (arrow 3), (B2) the CA2 of hippocampus (arrow 1), the ectorhinal cortex (arrow 2), the ventromedial hypothalamic nucleus (arrow 3), (B3) the postsubiculum (arrow 1), and superficial gray layer of the superior colliculus (arrow 2). (C) Ctgf-Cre:Ai75 transgenic mouse line to delineate following structures: (C1) the granular insular cortex (arrow), (2) the dorsal endopiriform nucleus (arrow 1), the medial amygdalar nucleus, anterodorsal (arrow 2), the thalamic structures such as anteromedial thalamic nucleus (arrow 3), (3) the posteromedial cortical amygdala (arrow 1), the subparafascicular thalamic nucleus (arrow 2), and the posterior hypothalamic nucleus, dorsal part (arrow 3).

## Supplementary Tables and Files

**Table S1, Related to Figure 2 and S1: Transgenic mouse list**

List of all transgenic mouse brains used for fine label alignments and further segmentations. Cell type specific drivers and corresponding reporter lines are outlined, as well as the specific structures that were highlighted from each line. See Table S2 for the abbreviation.

**Table S2, Related to Figure 5: Label identification numbers**

List of the FP based label full names, abbreviation, numerical ID, parent structures for the ontology, and corresponding Allen label nomenclature. Lack of clearly matched areas between our FP based labels and the Allen labels was left blank.

**Supplementary File 1: A series of digitized labels in every 100 μm z spacing**

The file name contains the information about Bregma anterior posterior coordinate of each image

**Supplementary File 2: A series of matched CCF background planes in every 100 μm z spacing**

Z numbers in the file name are matched with label numbers in Supplementary File 1

