## Supplementary figures and images for "Enhanced and Unified Anatomical Labeling for a Common Mouse Brain Atlas"

### 1_AP+4.3.tif

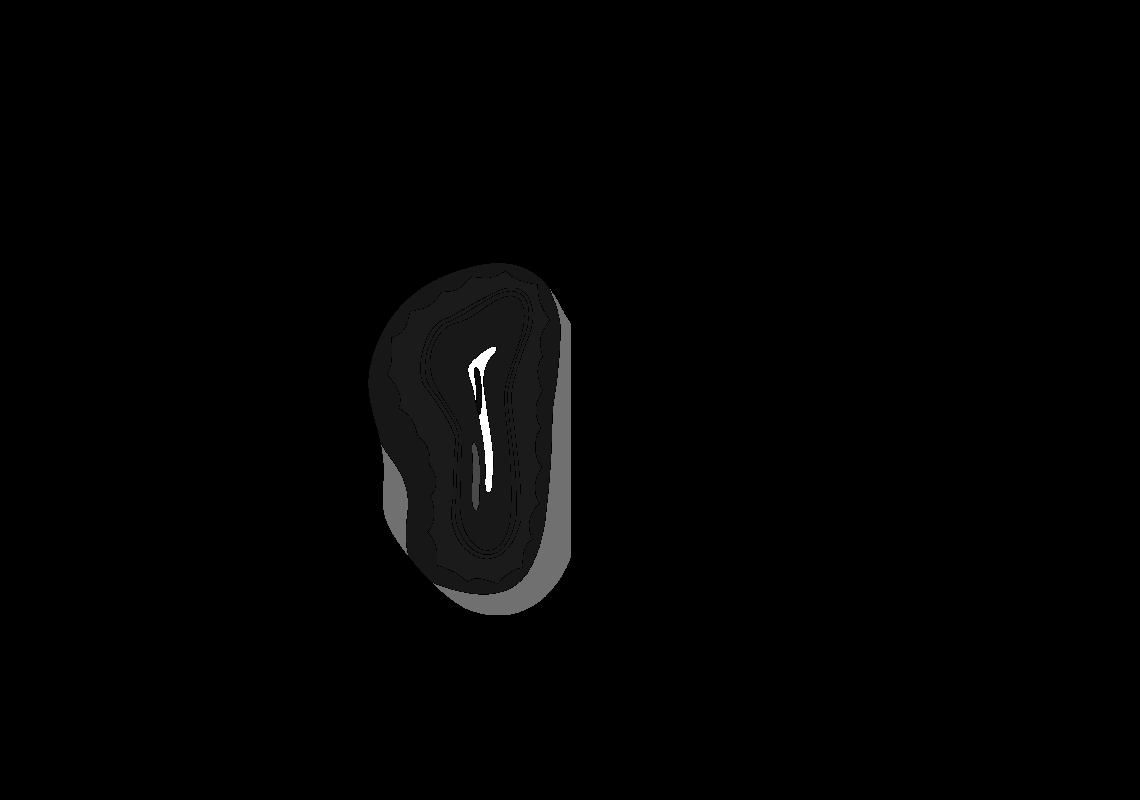

### 4_AP+4.0.tif

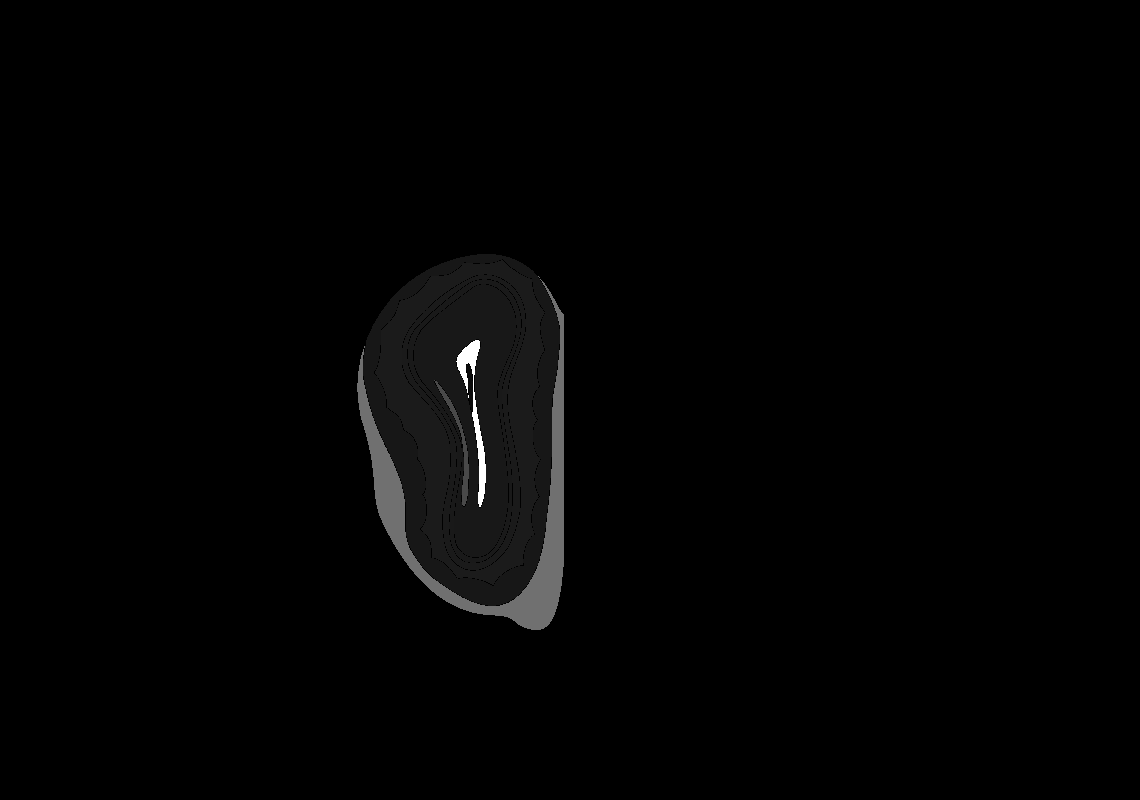

### 5_AP+3.9.tif

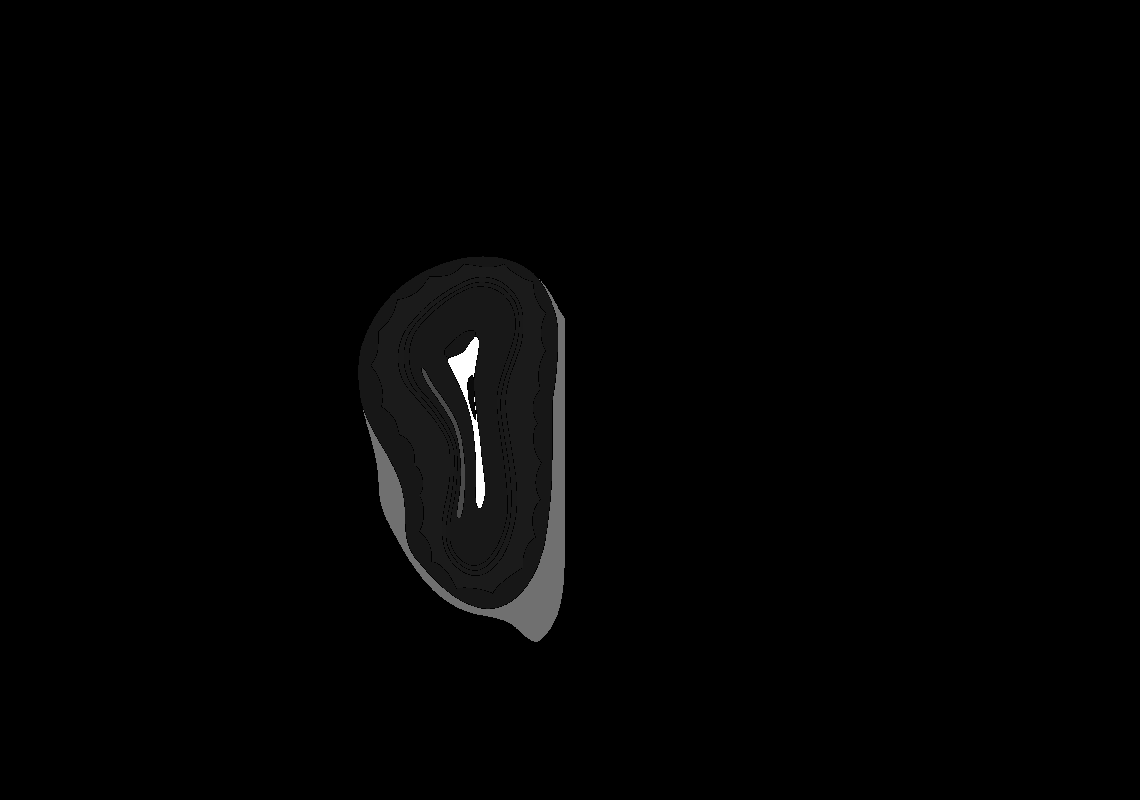

### 10_AP+3.4.tif

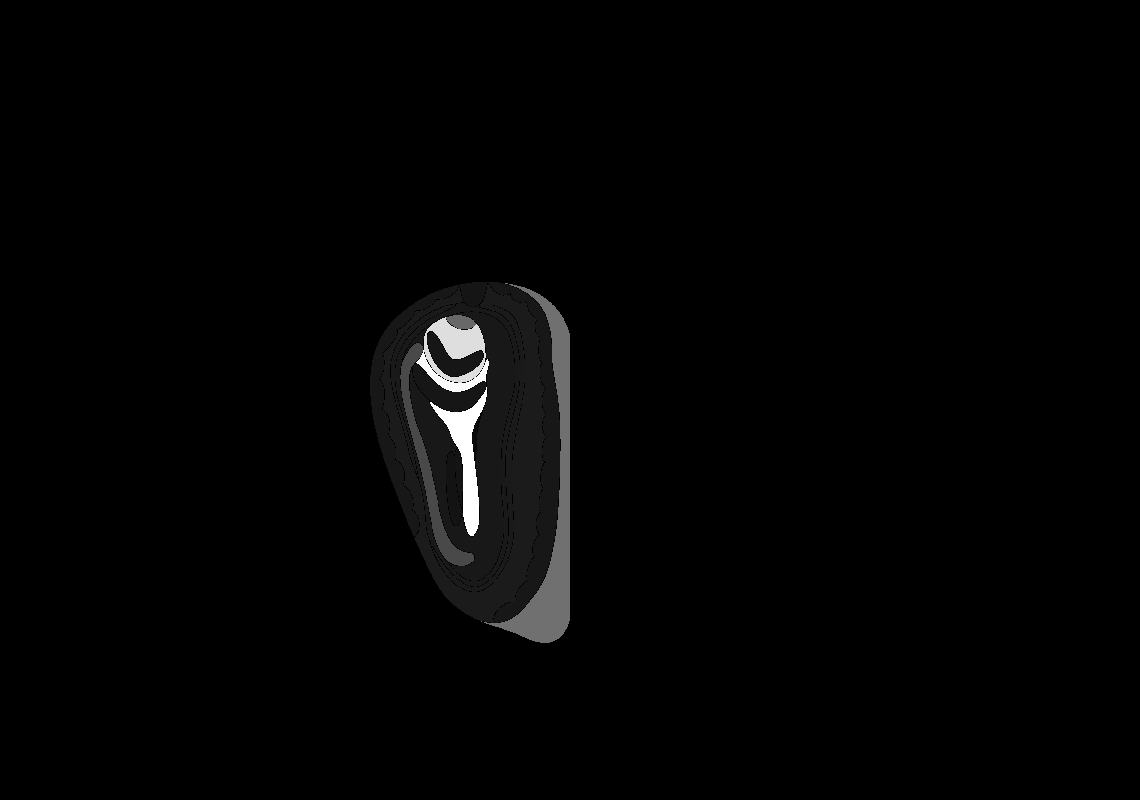

### 17_AP+2.7.tif

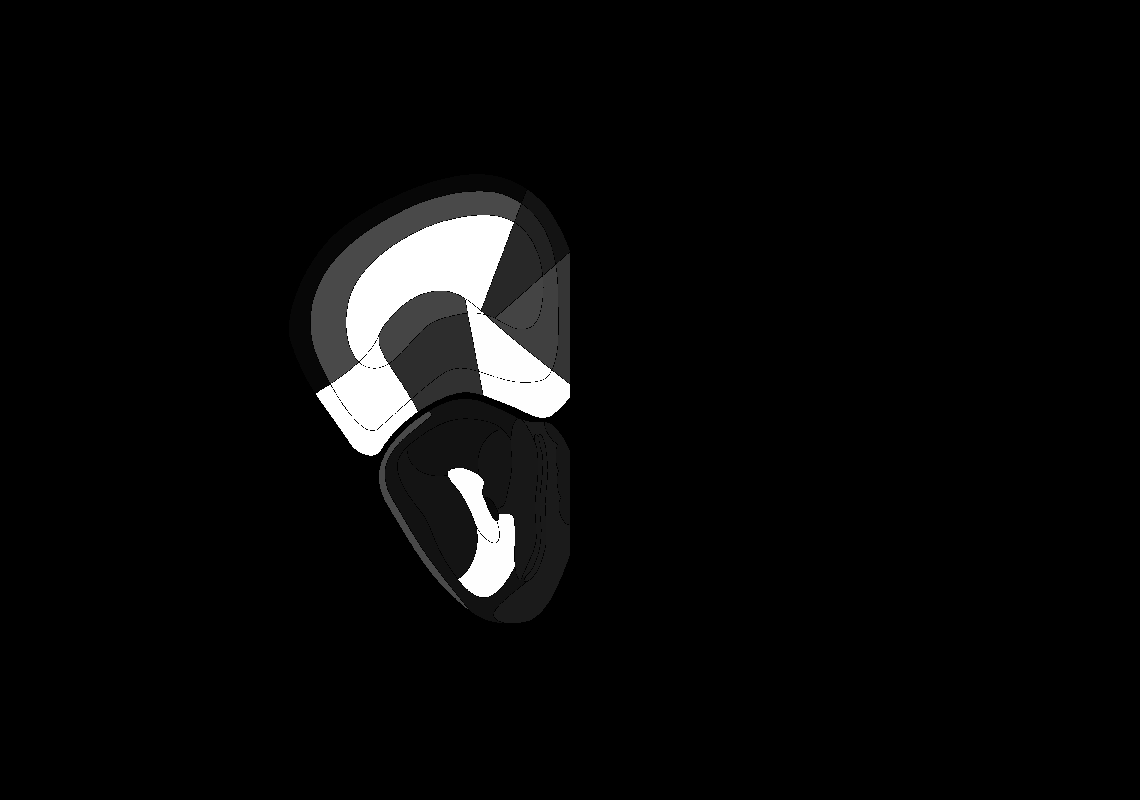

### 20_AP+2.4.tif

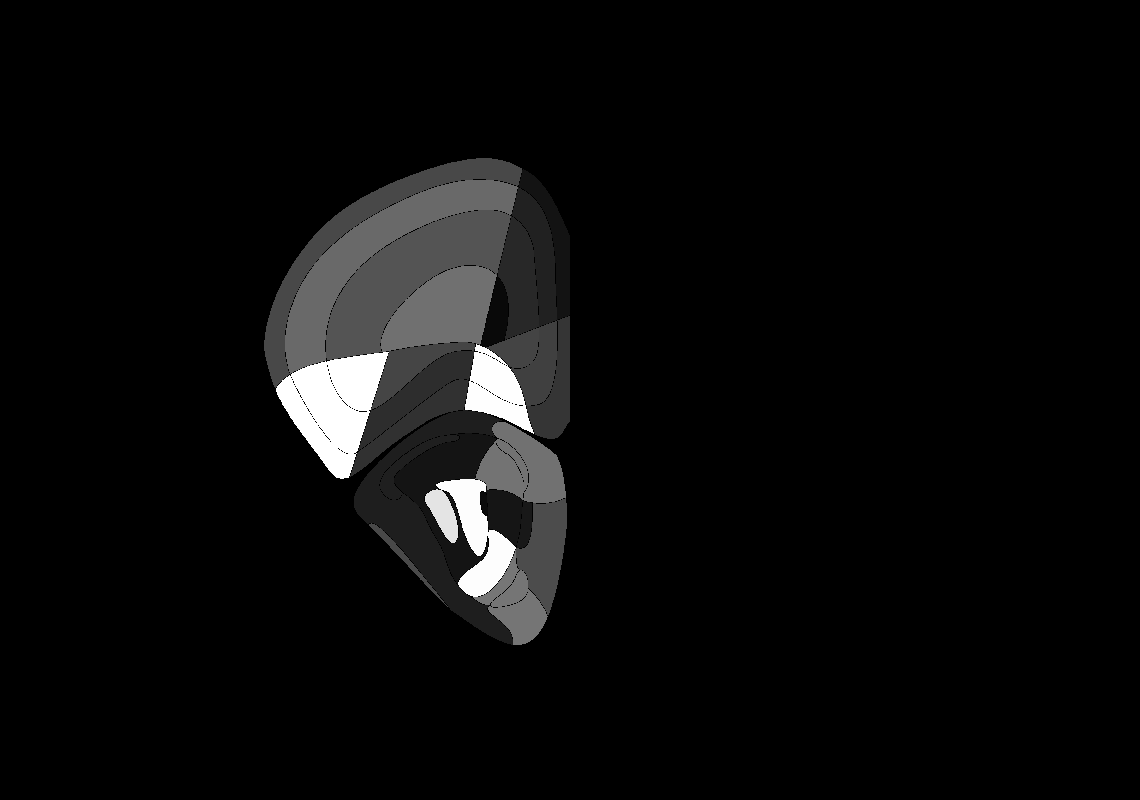

### 42_AP+0.2.tif

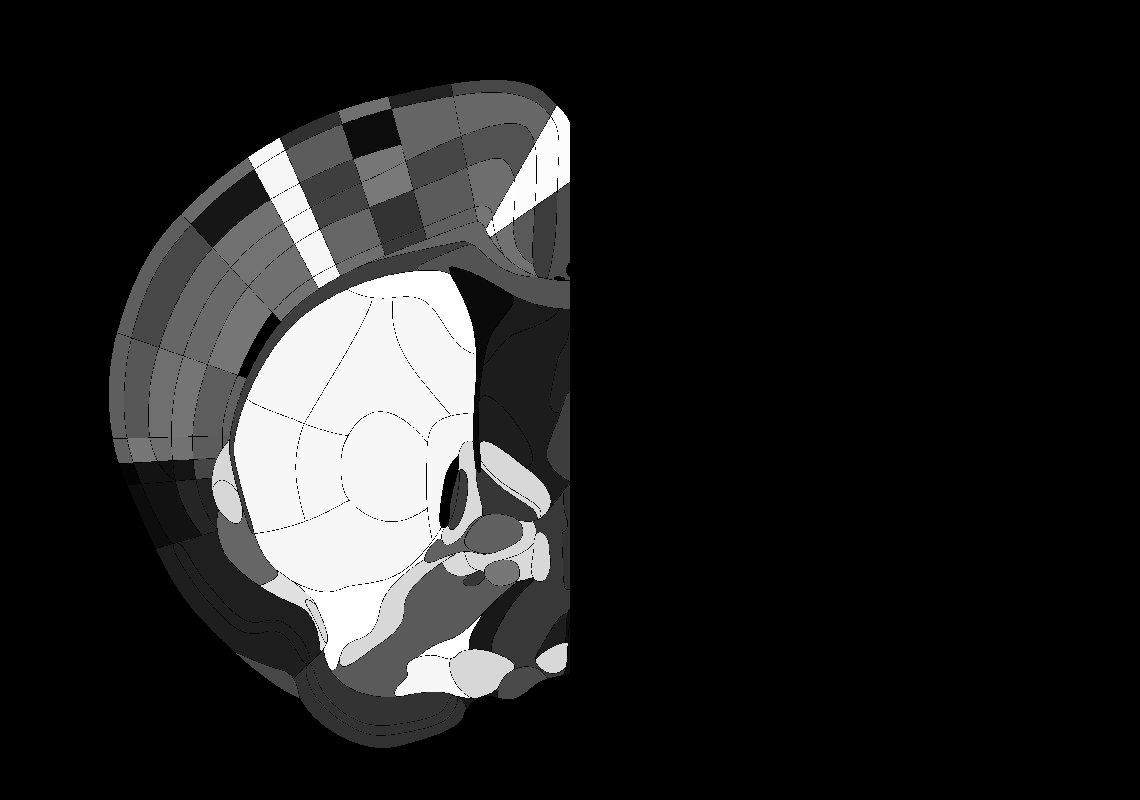

### 45_AP-0.1.tif

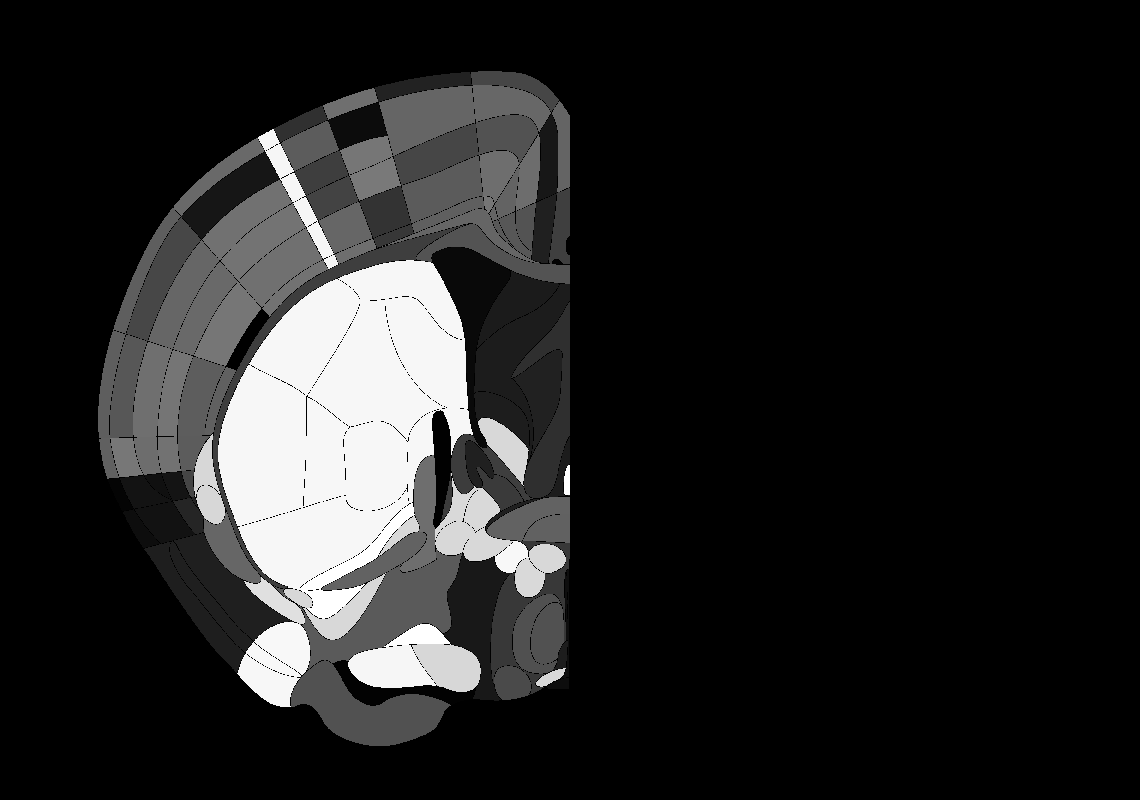

### 51_AP-0.7.tif

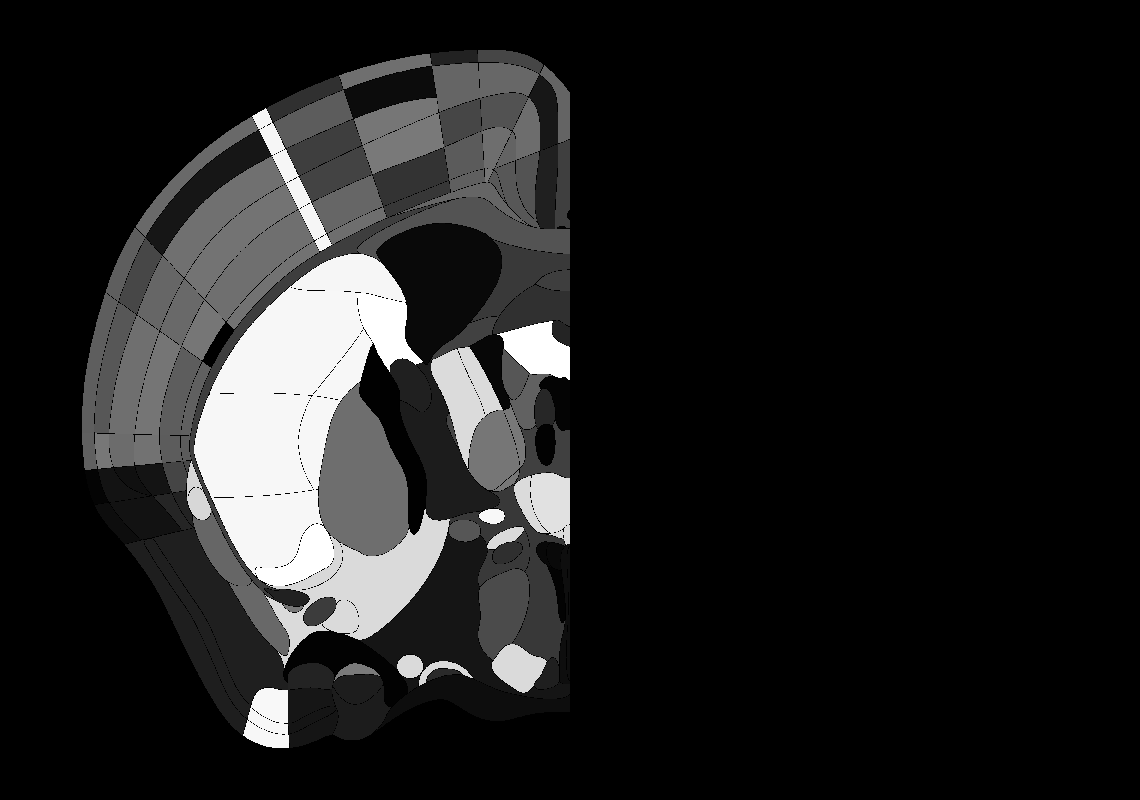

### 52_AP-0.8.tif

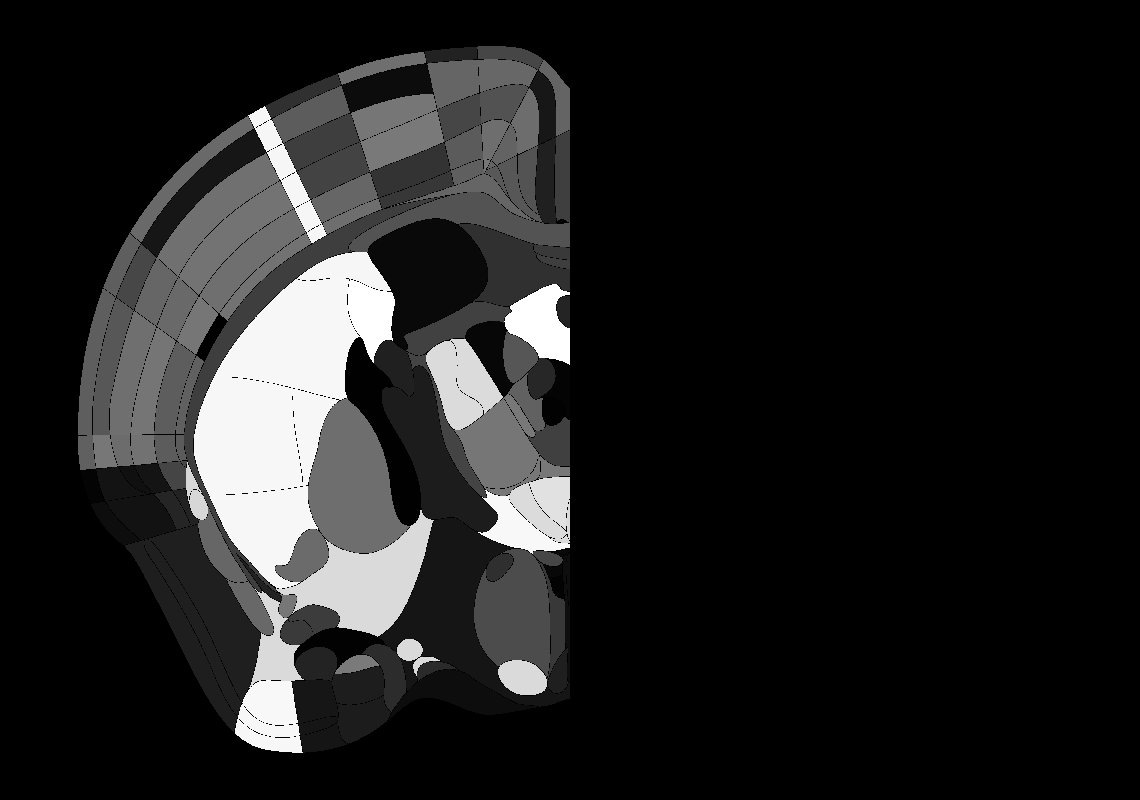

### 56_AP-1.2.tif

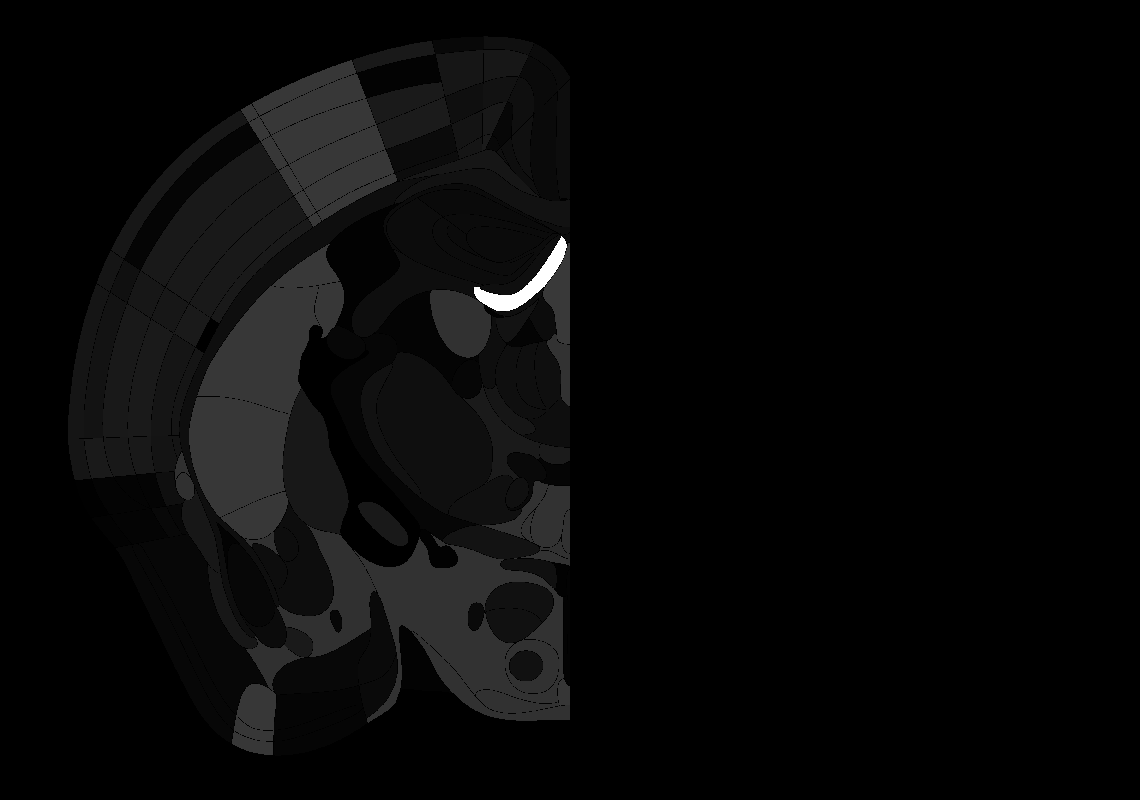

### 61_AP-1.7.tif

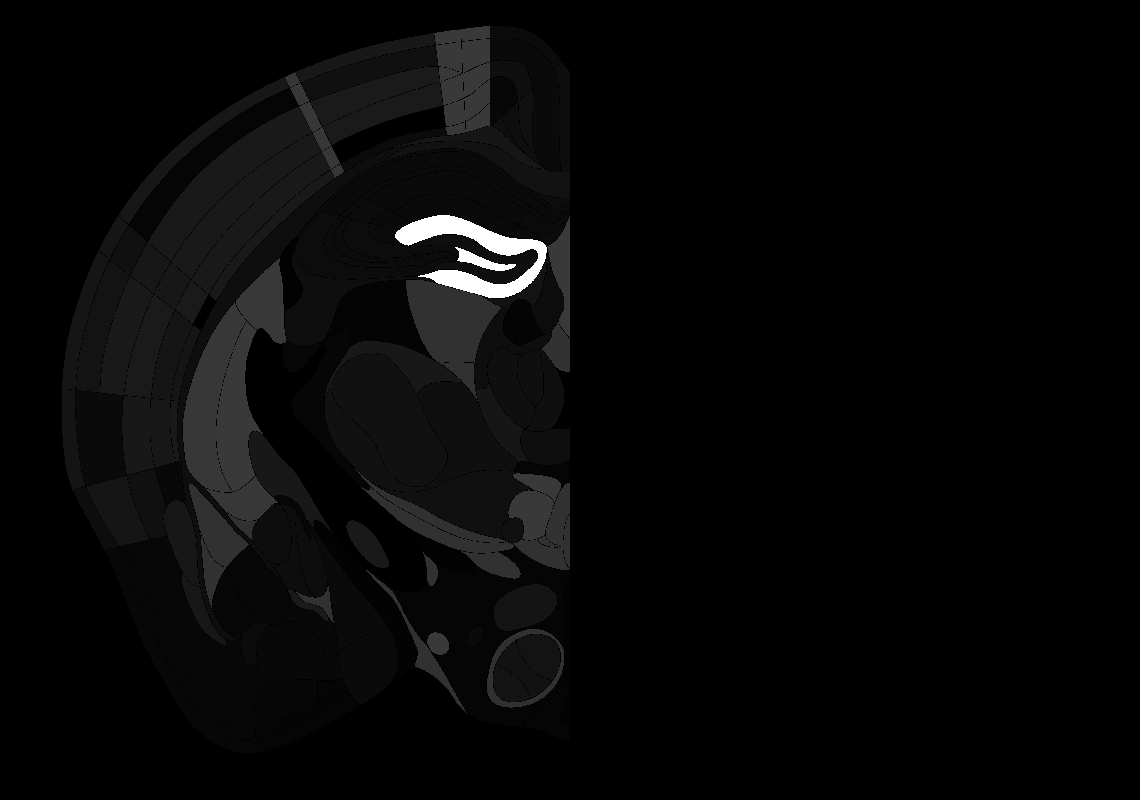

### 75_AP-3.1.tif

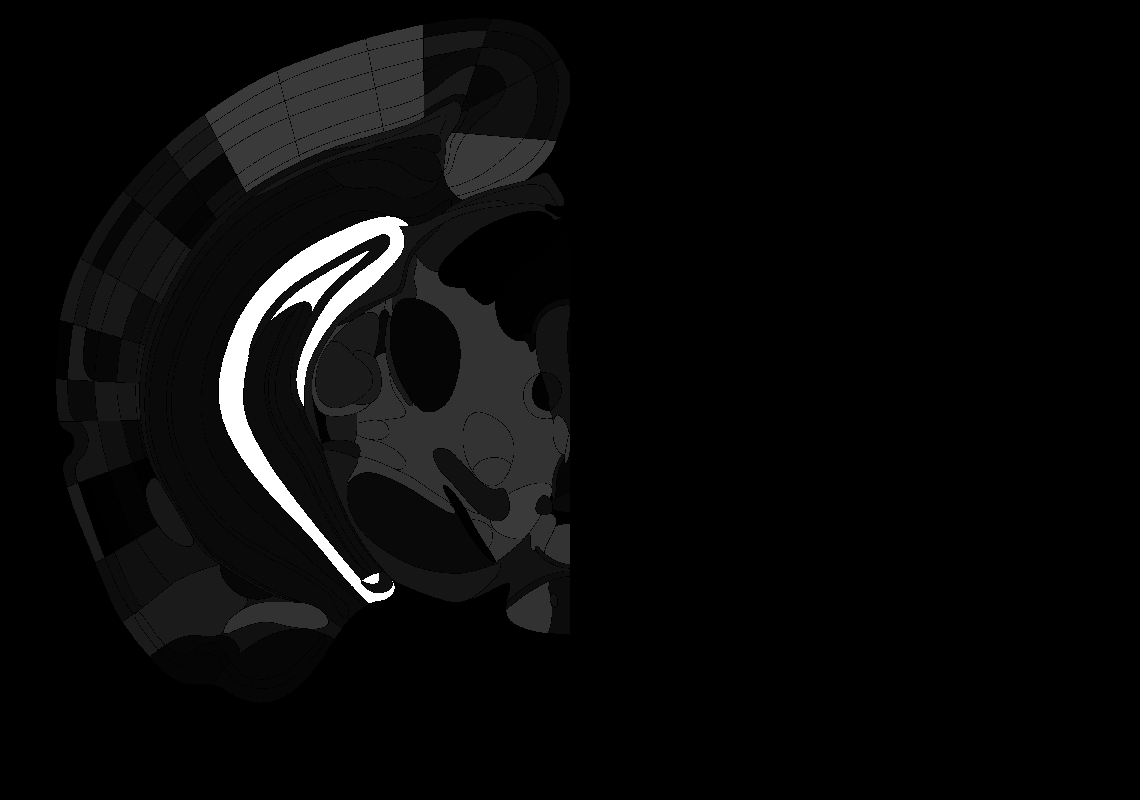

### 78_AP-3.4.tif

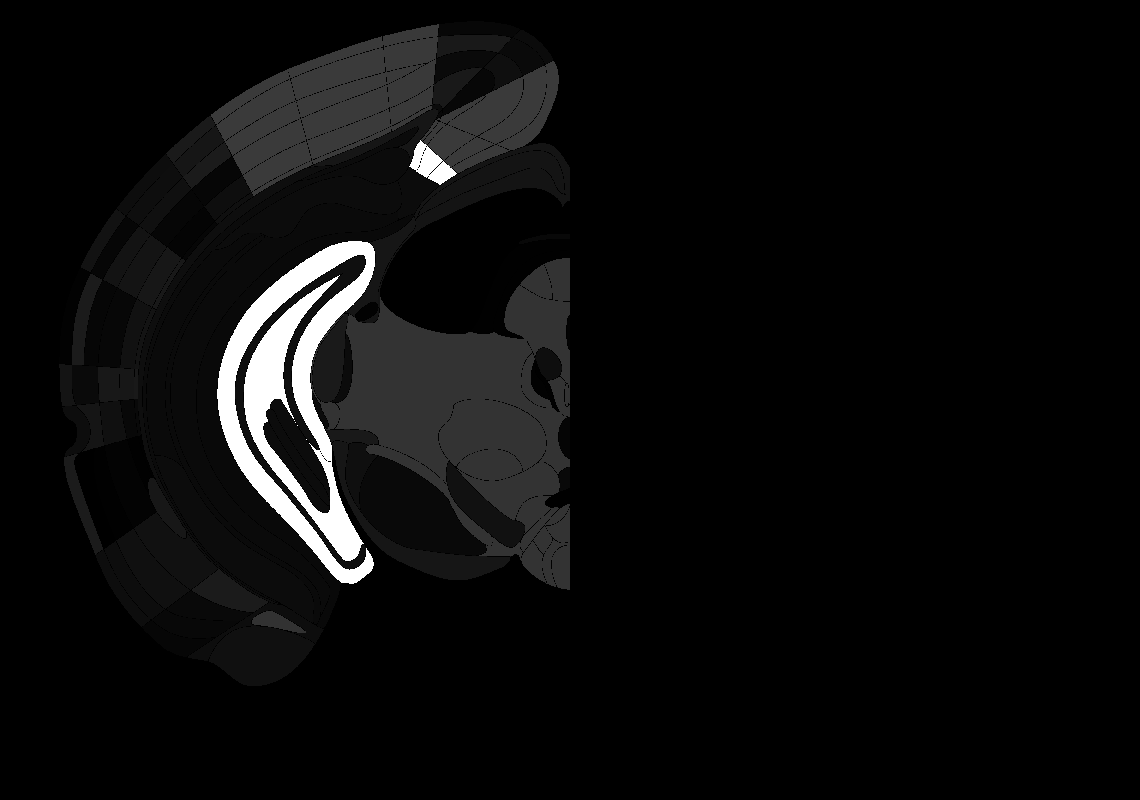

### 85_AP-4.1.tif

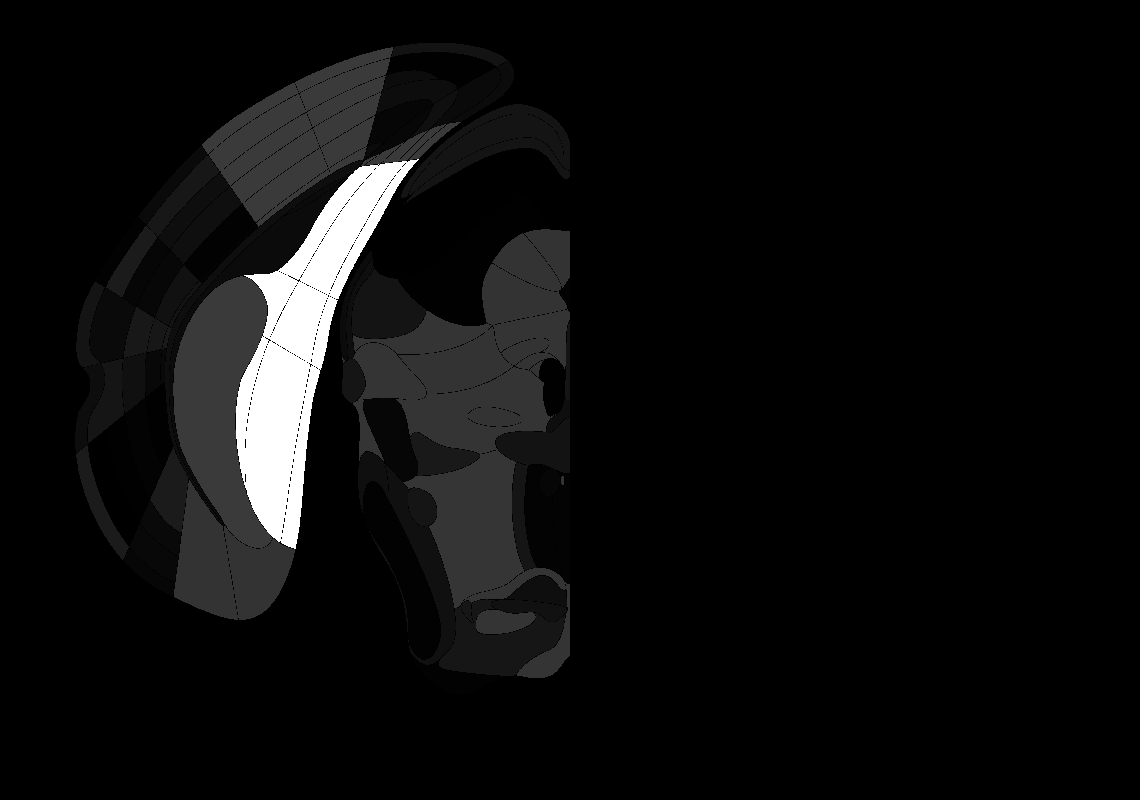

### 88_AP-4.4.tif

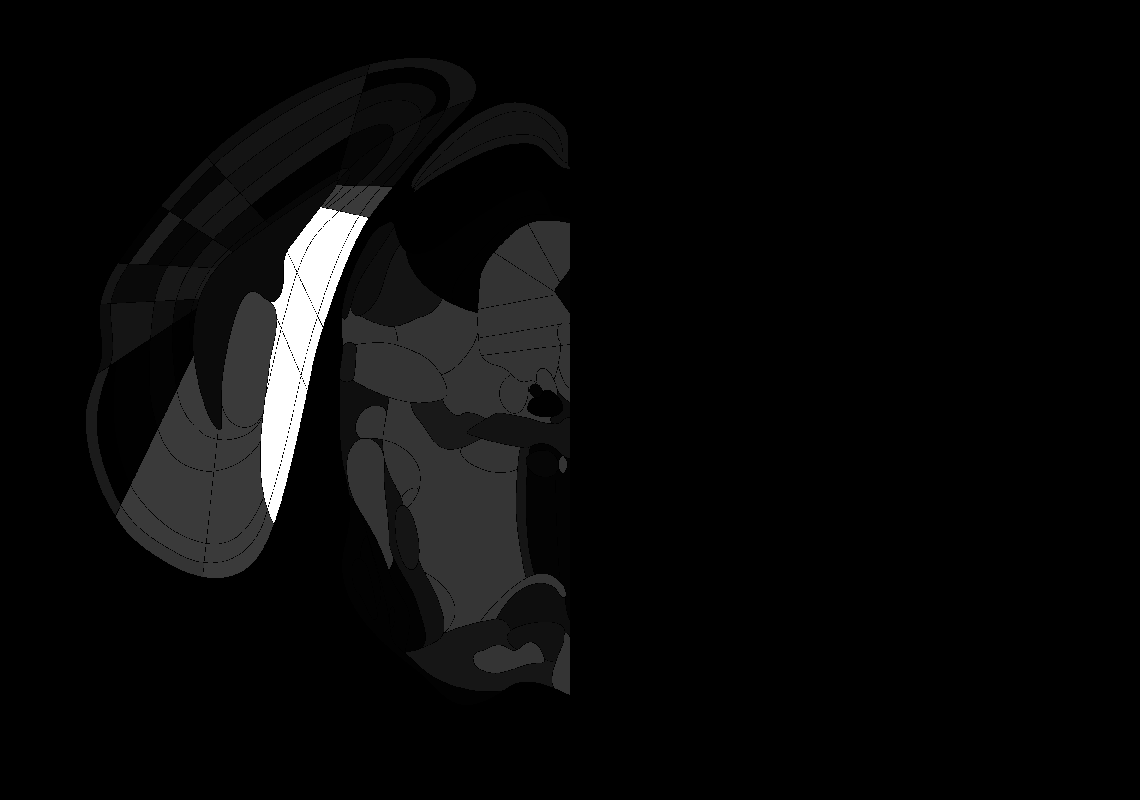

### 91_AP-4.7.tif

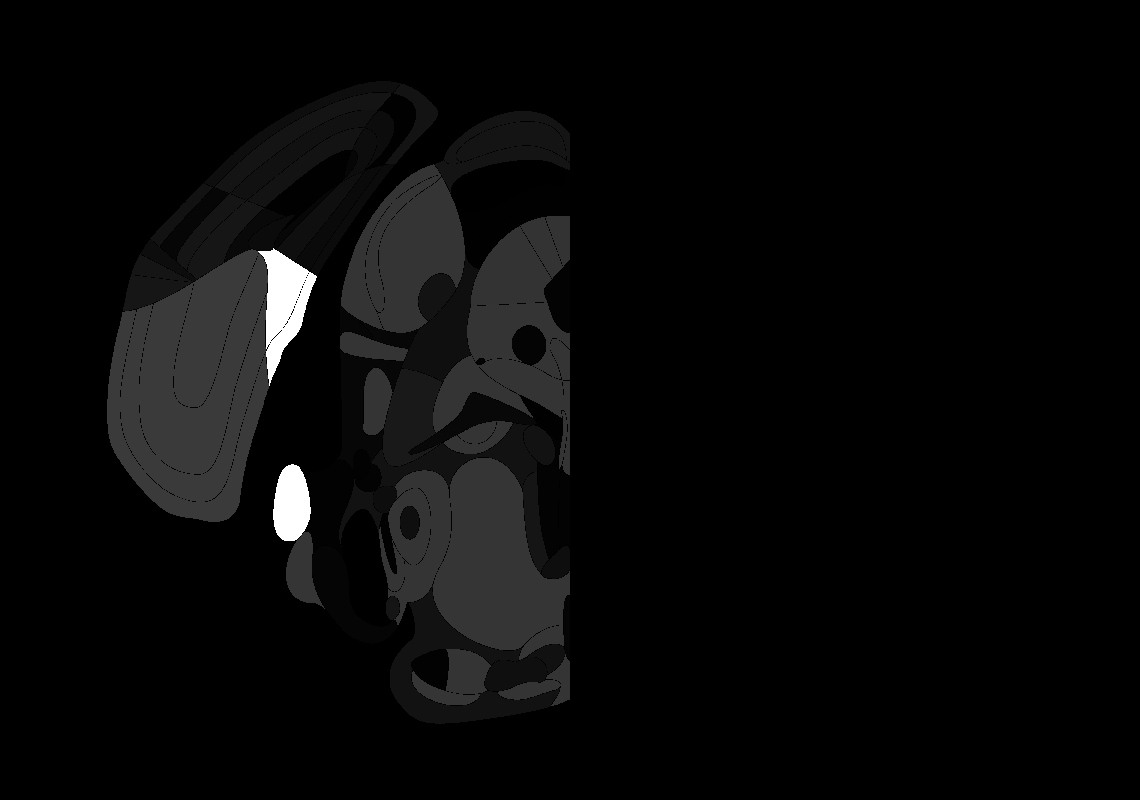

### 92_AP-4.8.tif

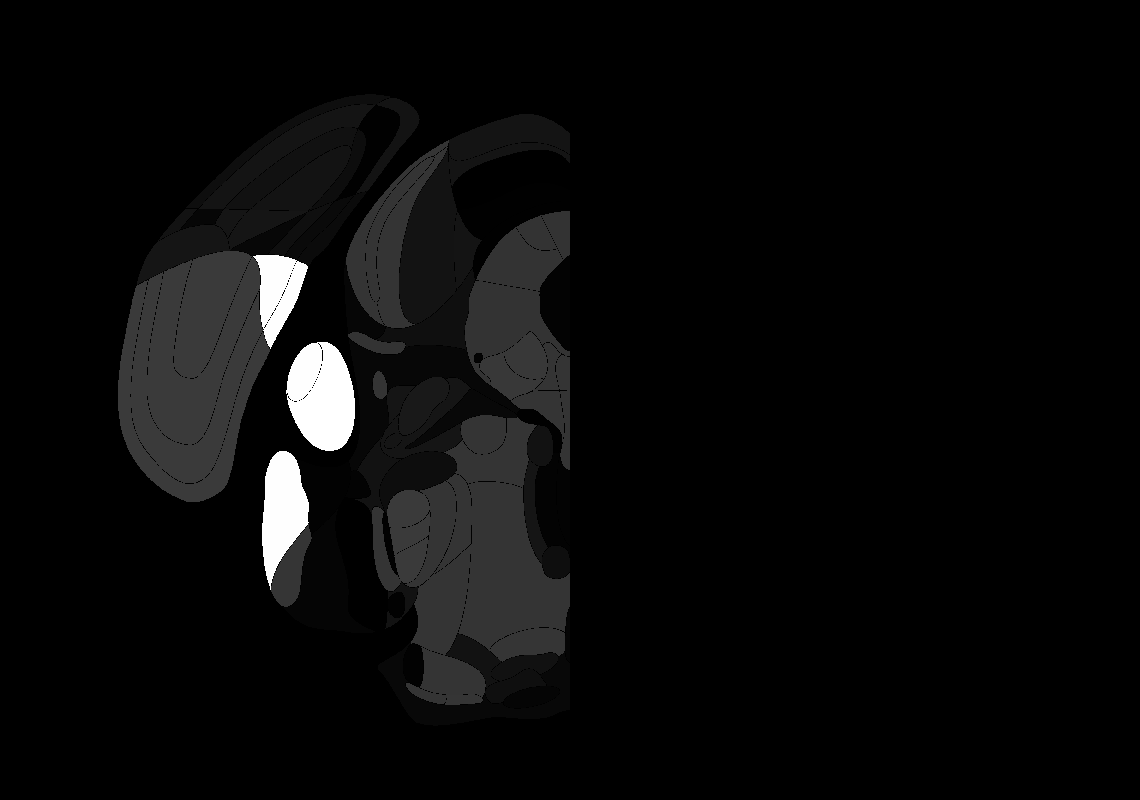

### 96_AP-5.2.tif

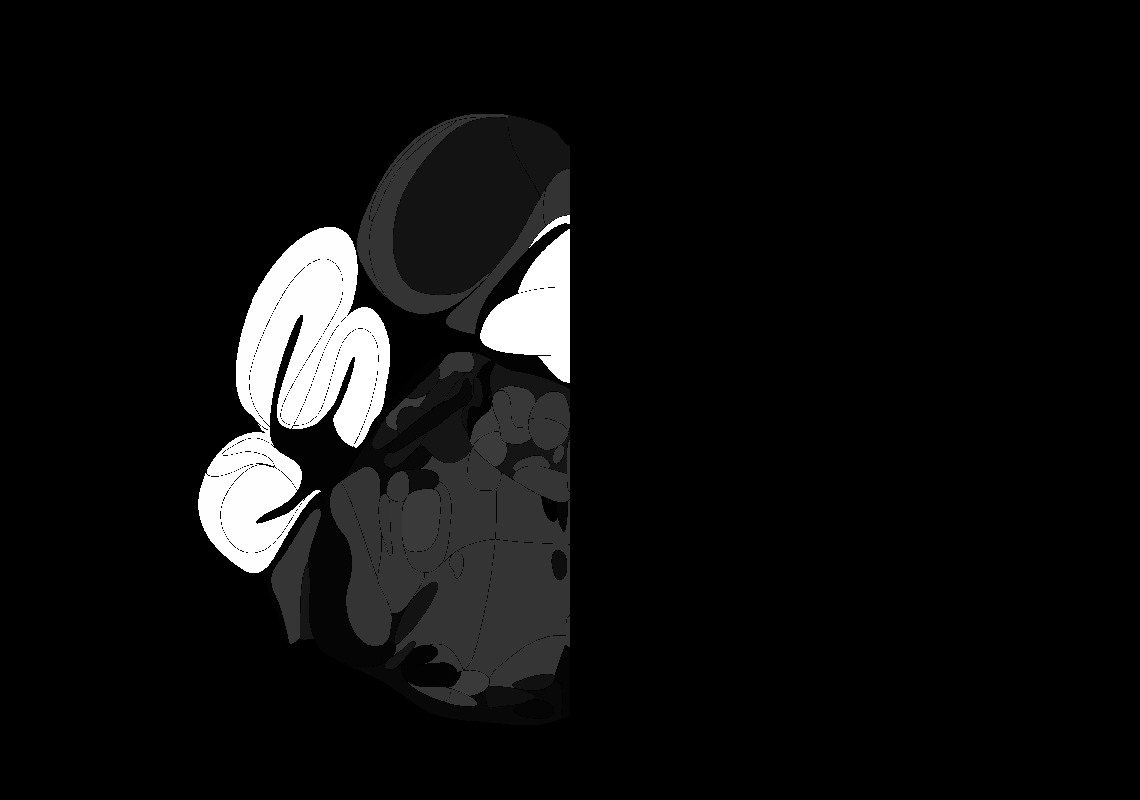

### 108_AP-6.4.tif

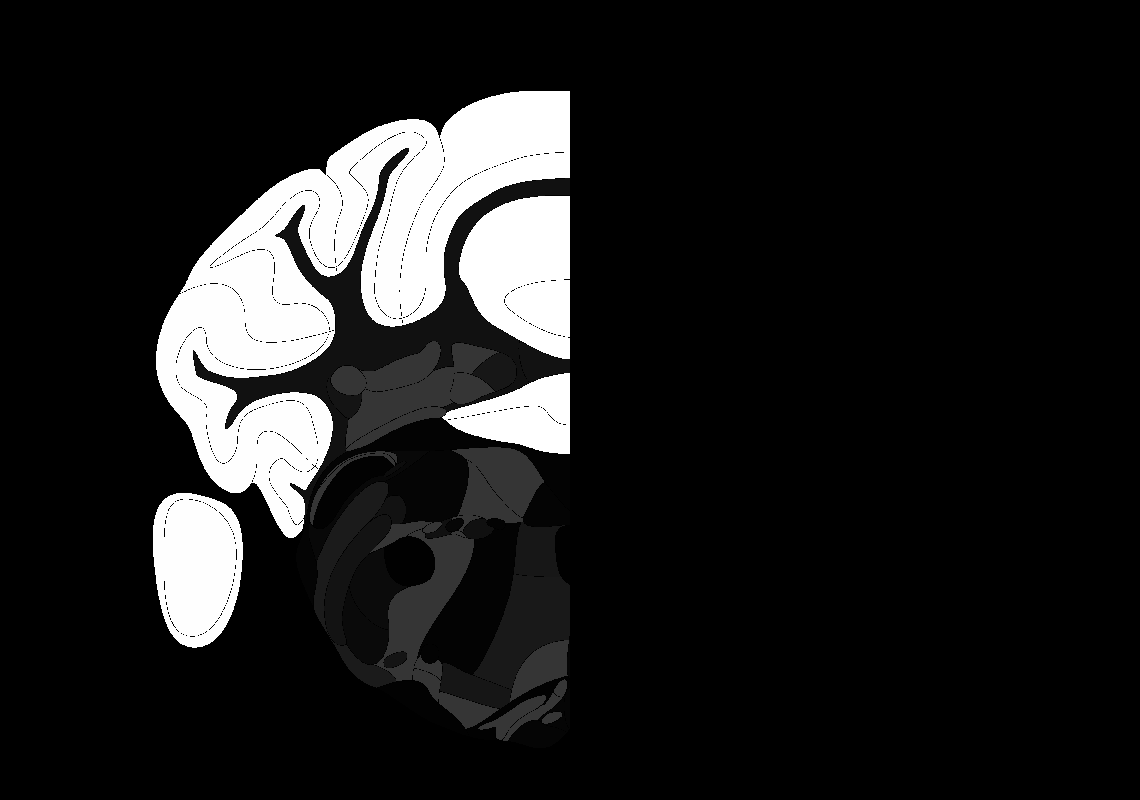

### 116_AP-7.2.tif

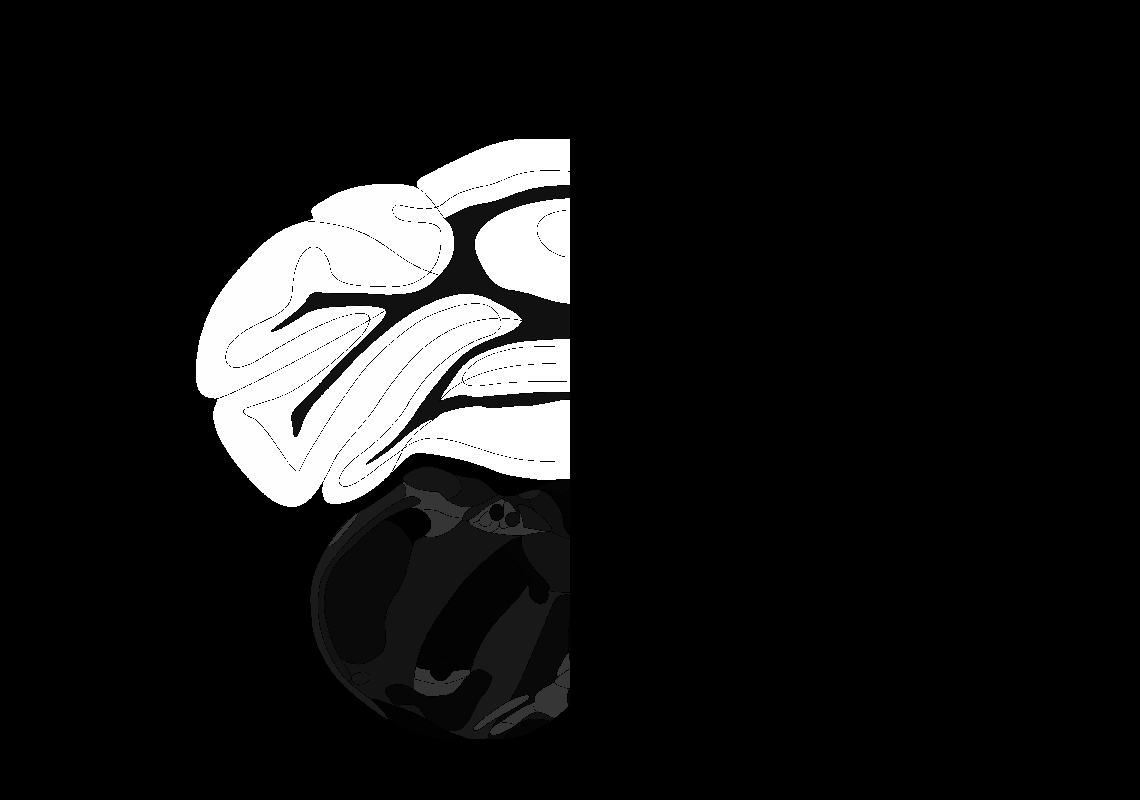

### 121_AP-7.7.tif

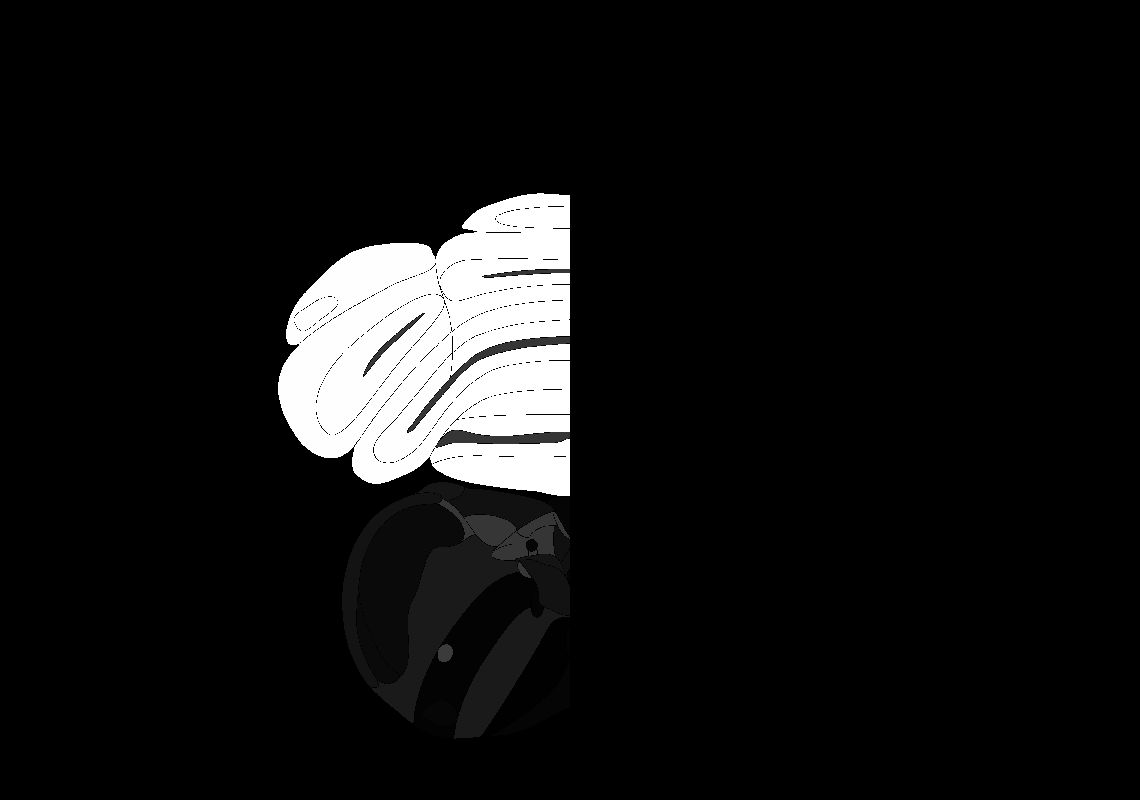

### 122_AP-7.8.tif

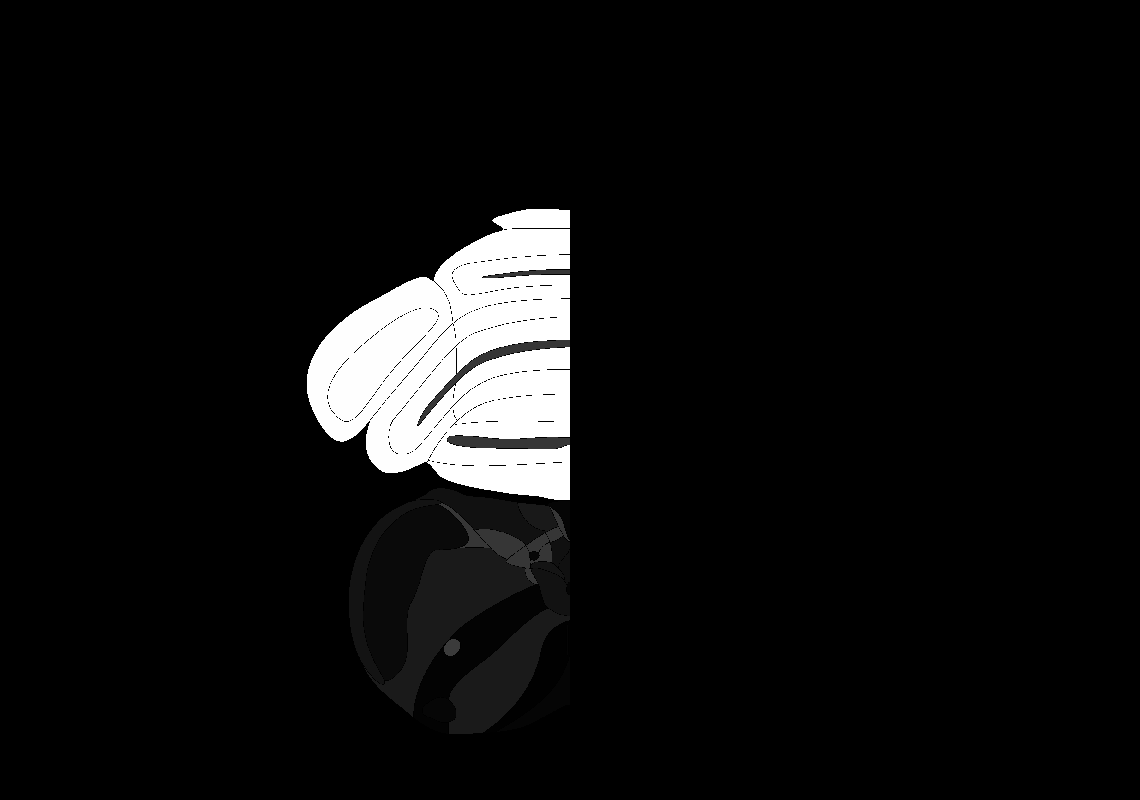

### AllenCCF_Z001.tif

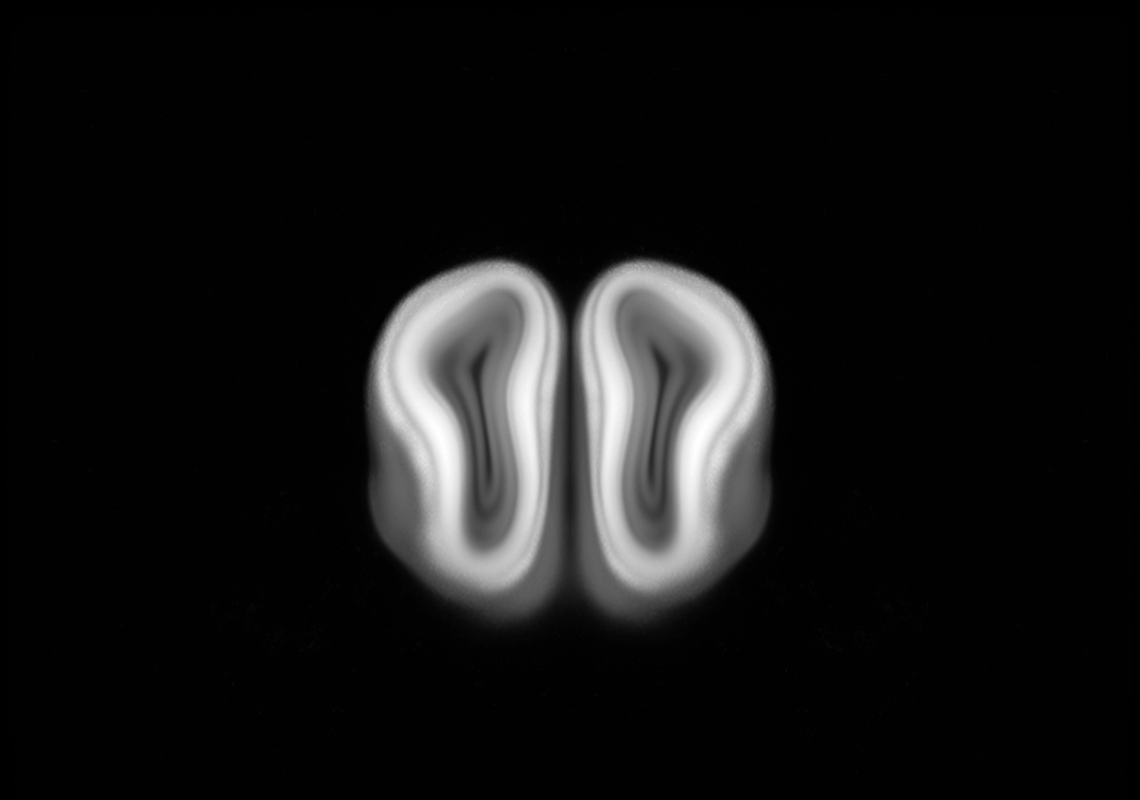

### AllenCCF_Z002.tif

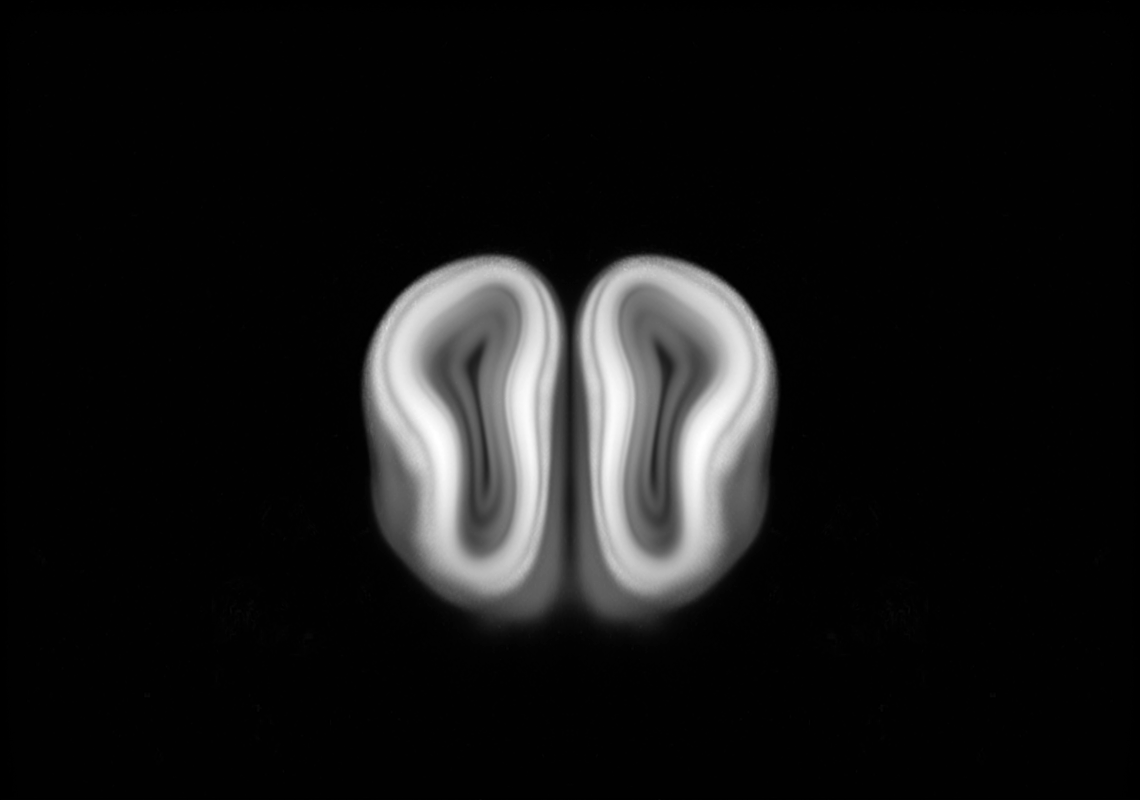

### AllenCCF_Z003.tif

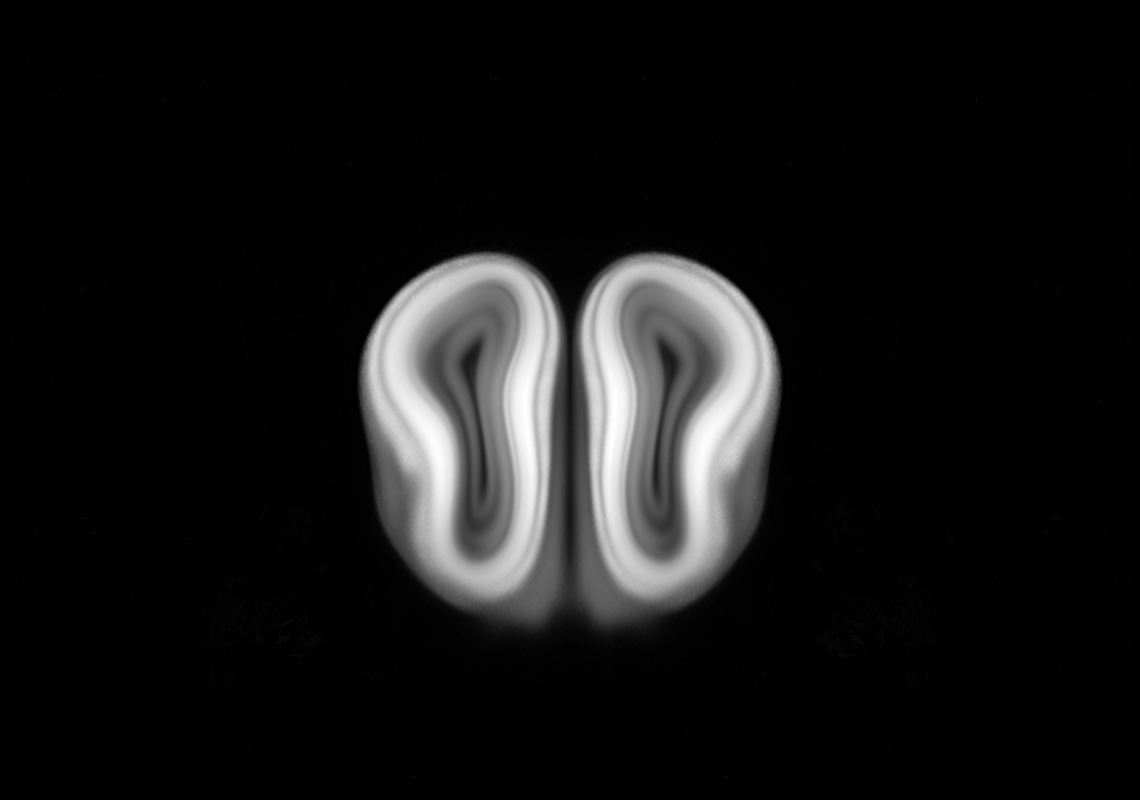

### AllenCCF_Z004.tif

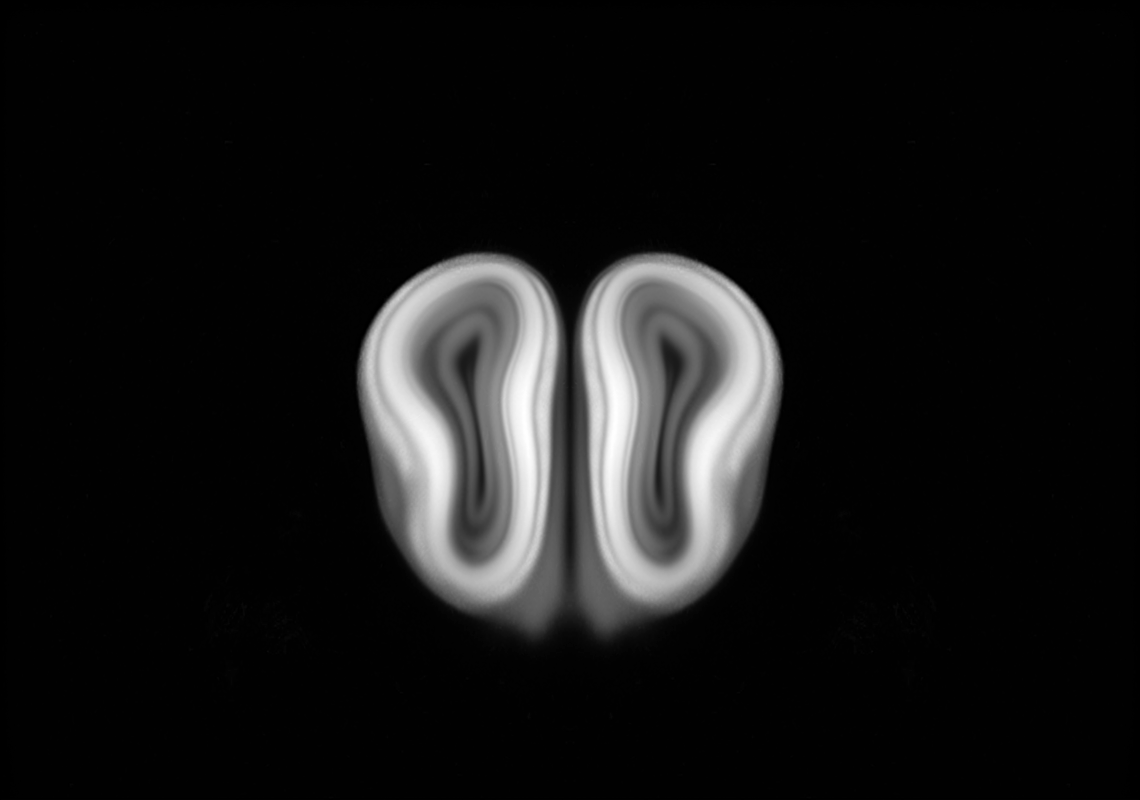

### AllenCCF_Z005.tif

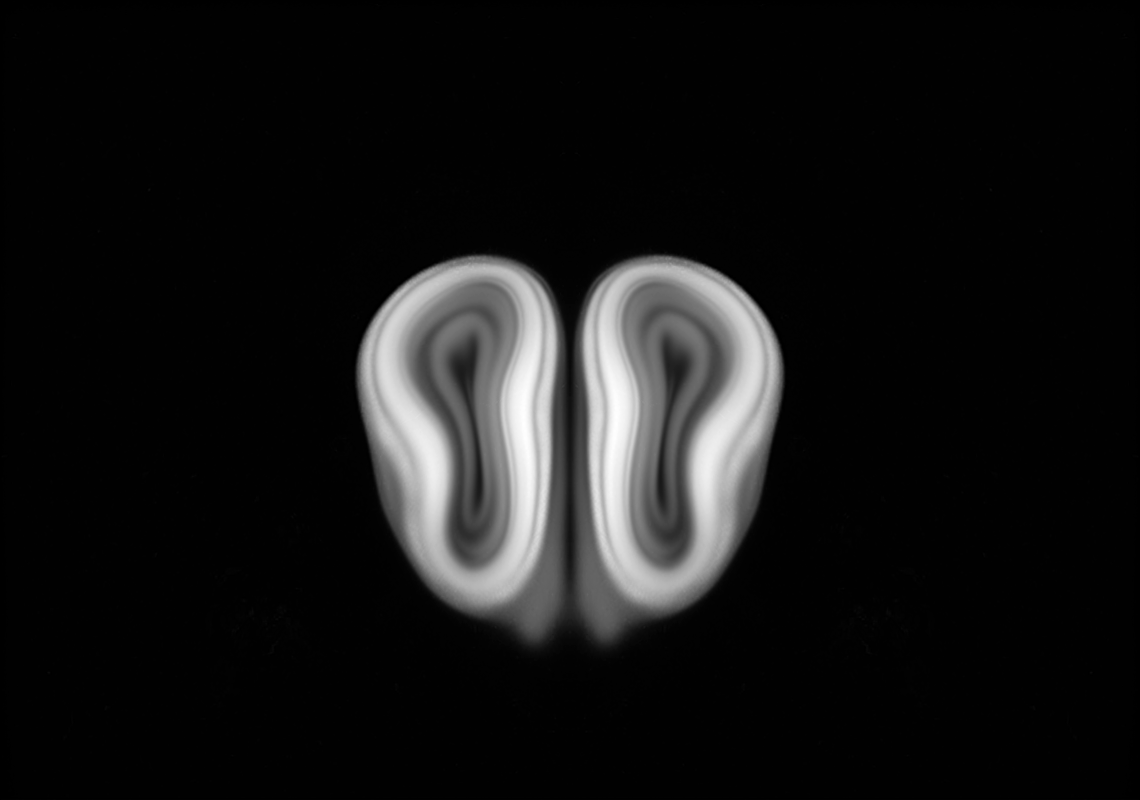

### AllenCCF_Z006.tif

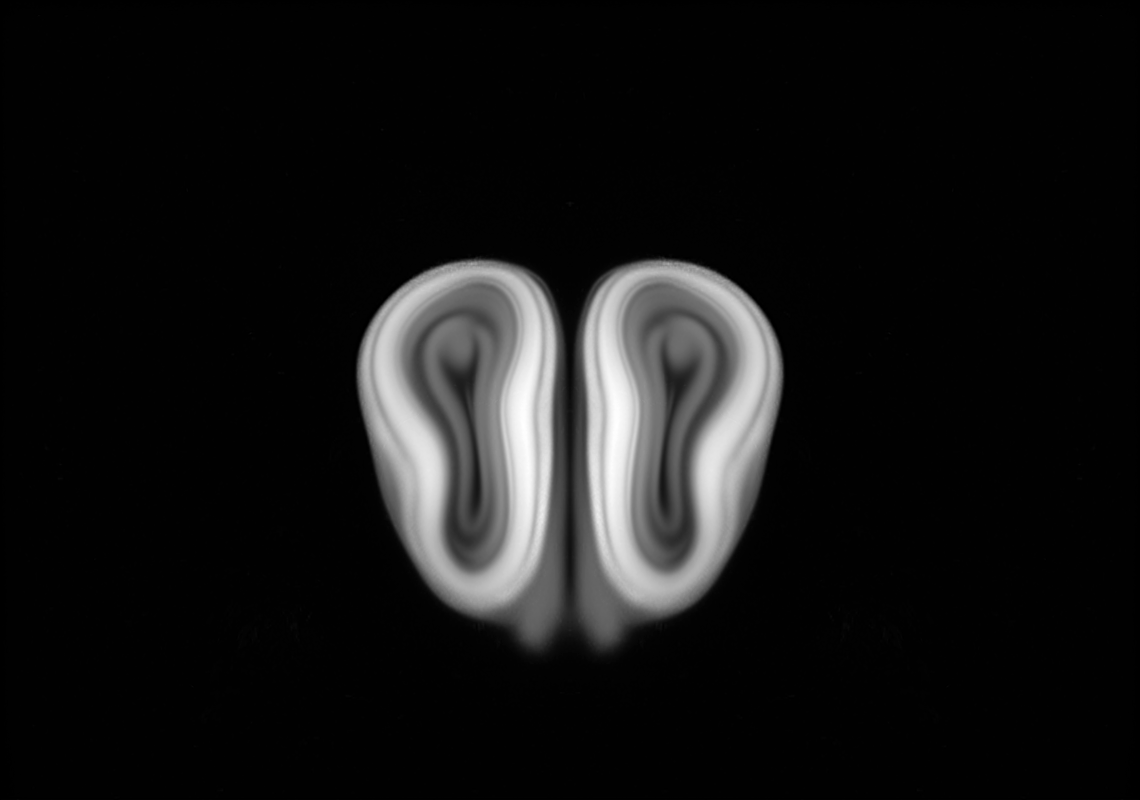

### AllenCCF_Z007.tif

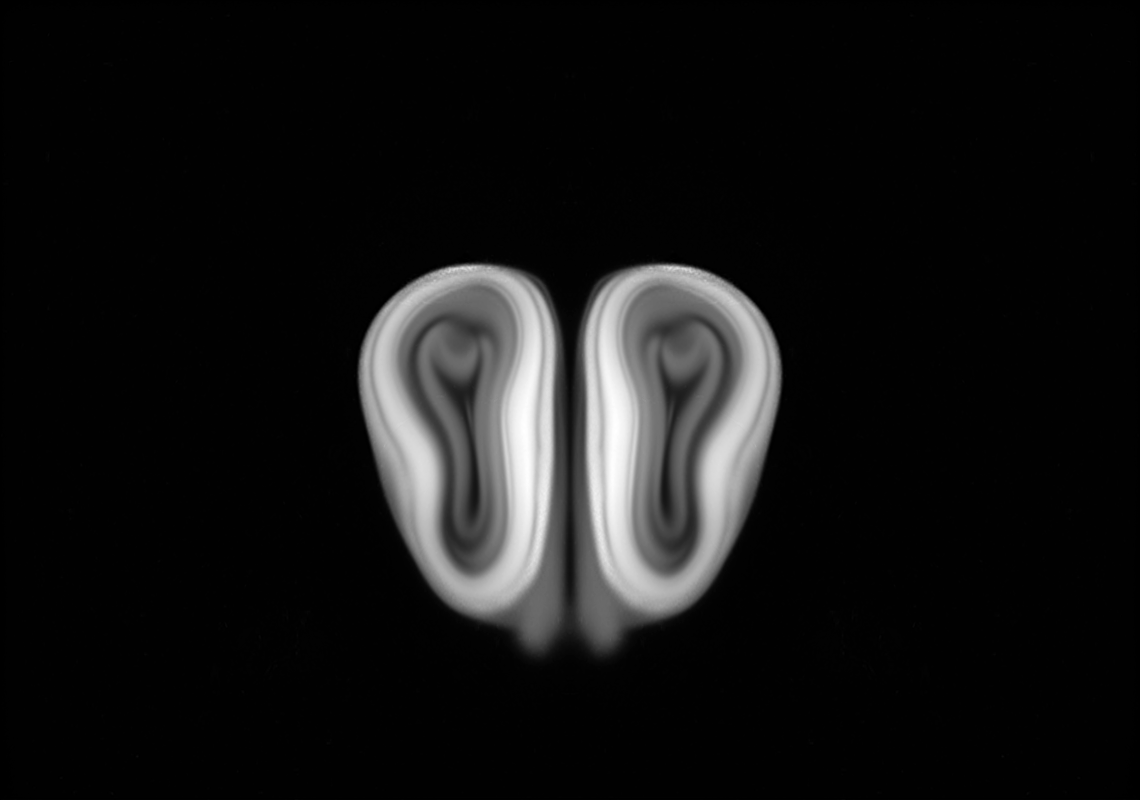

### AllenCCF_Z008.tif

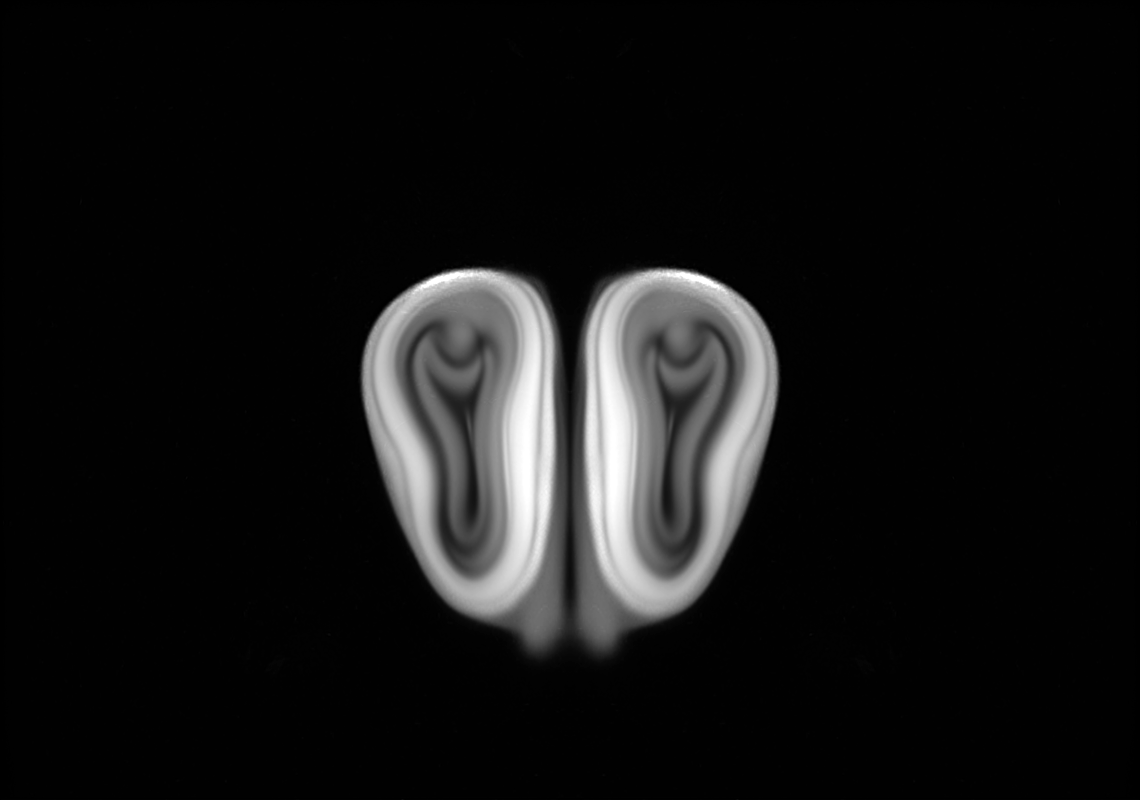

### AllenCCF_Z009.tif

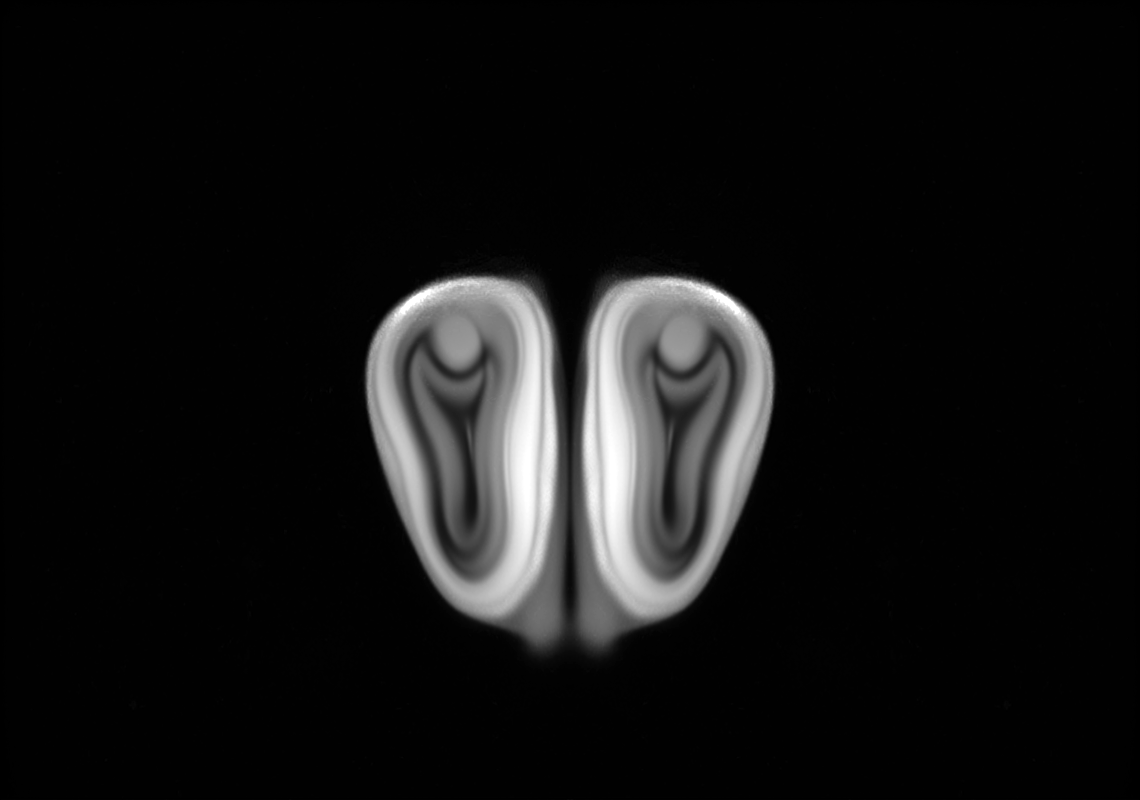

### AllenCCF_Z010.tif

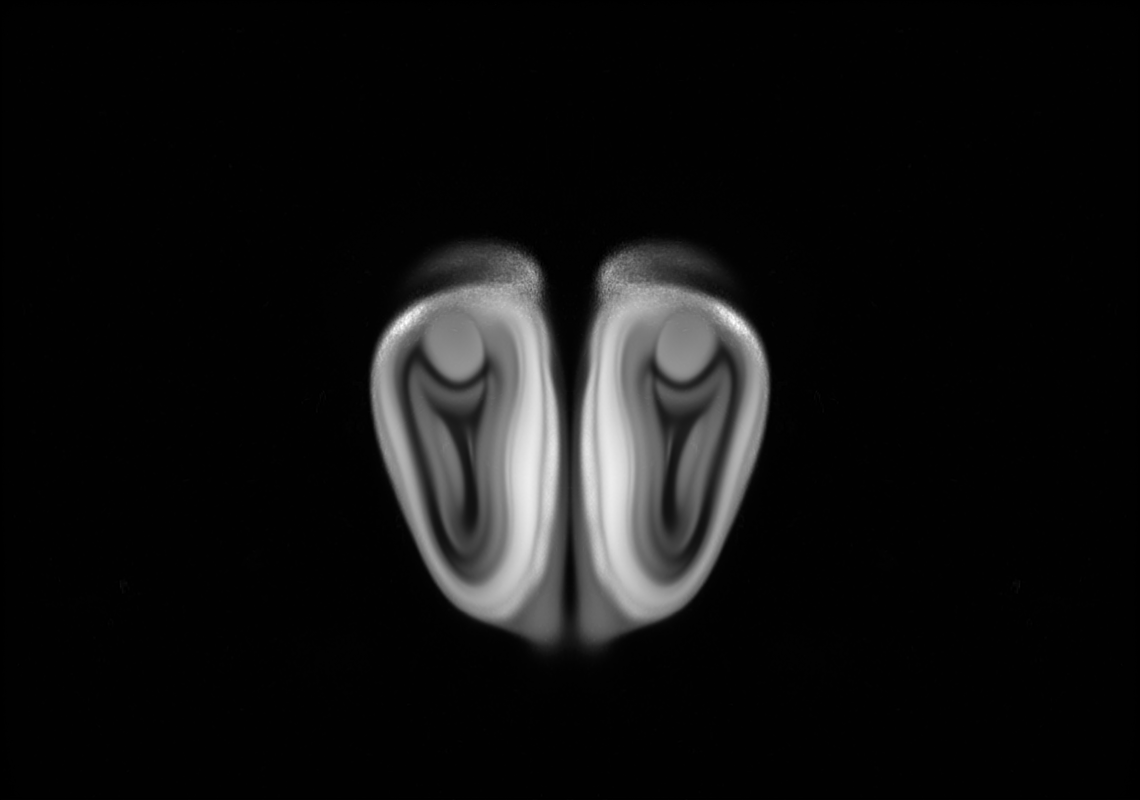

### AllenCCF_Z011.tif

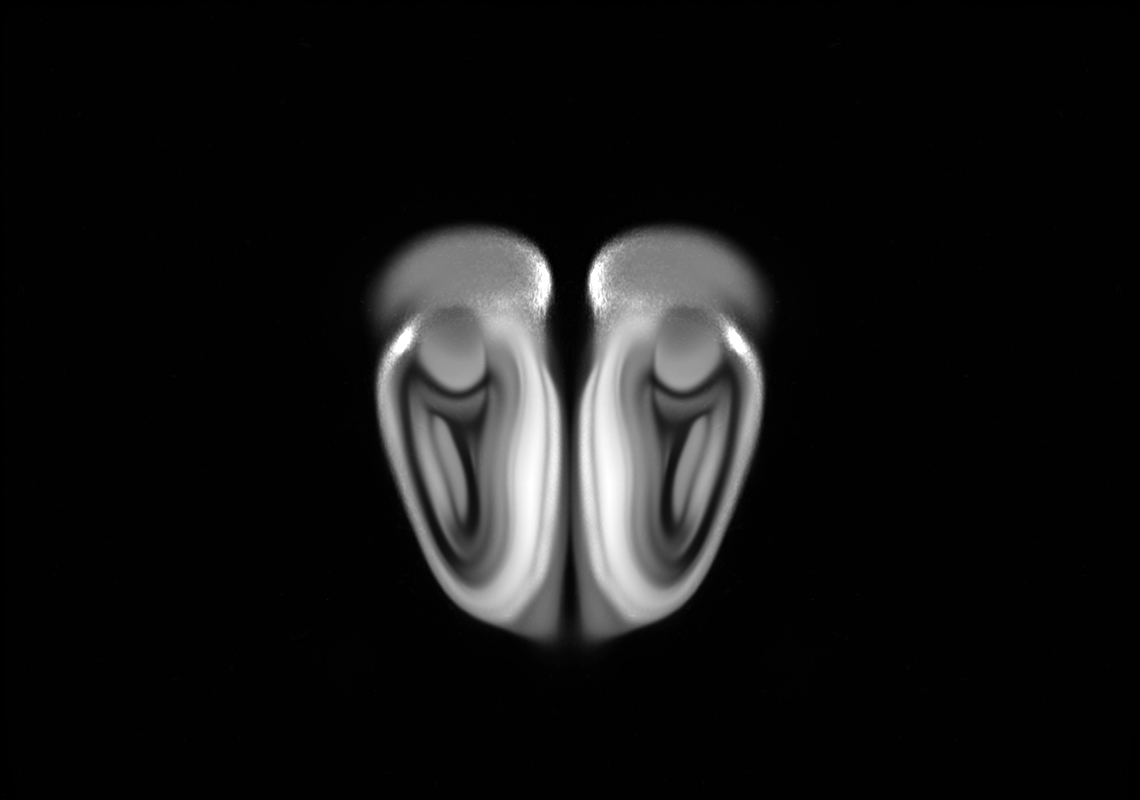

### AllenCCF_Z012.tif

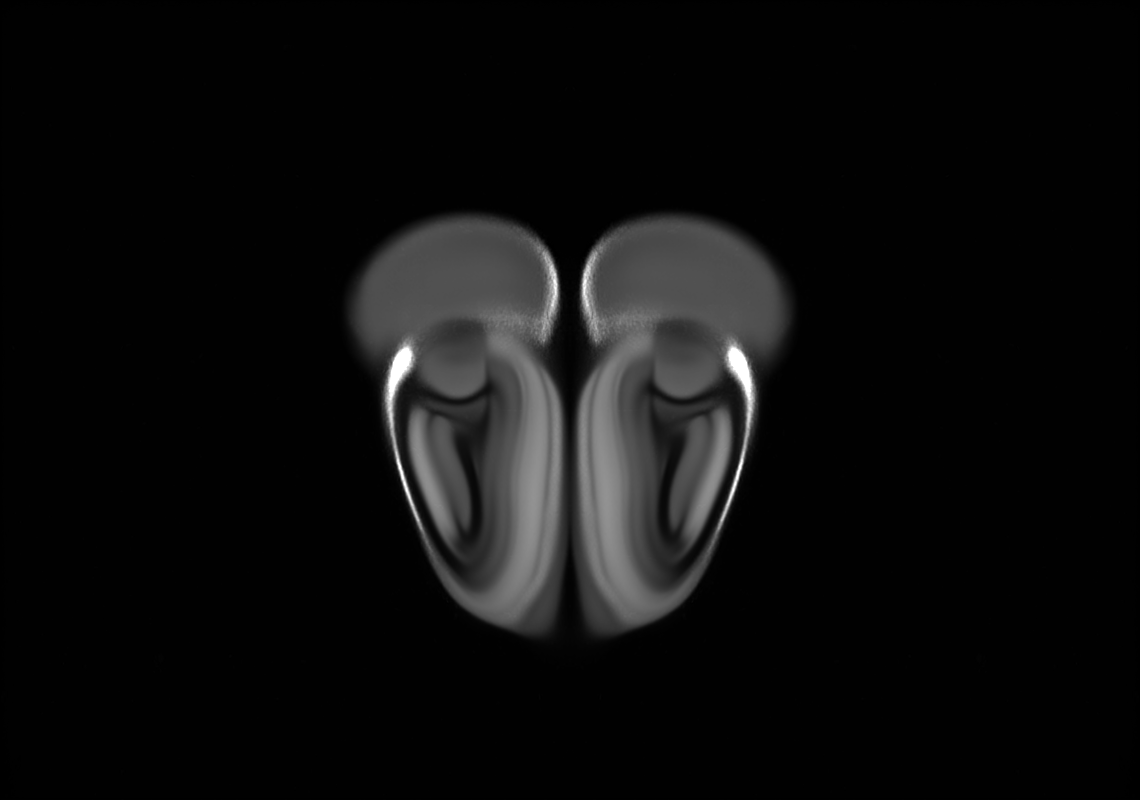

### AllenCCF_Z013.tif

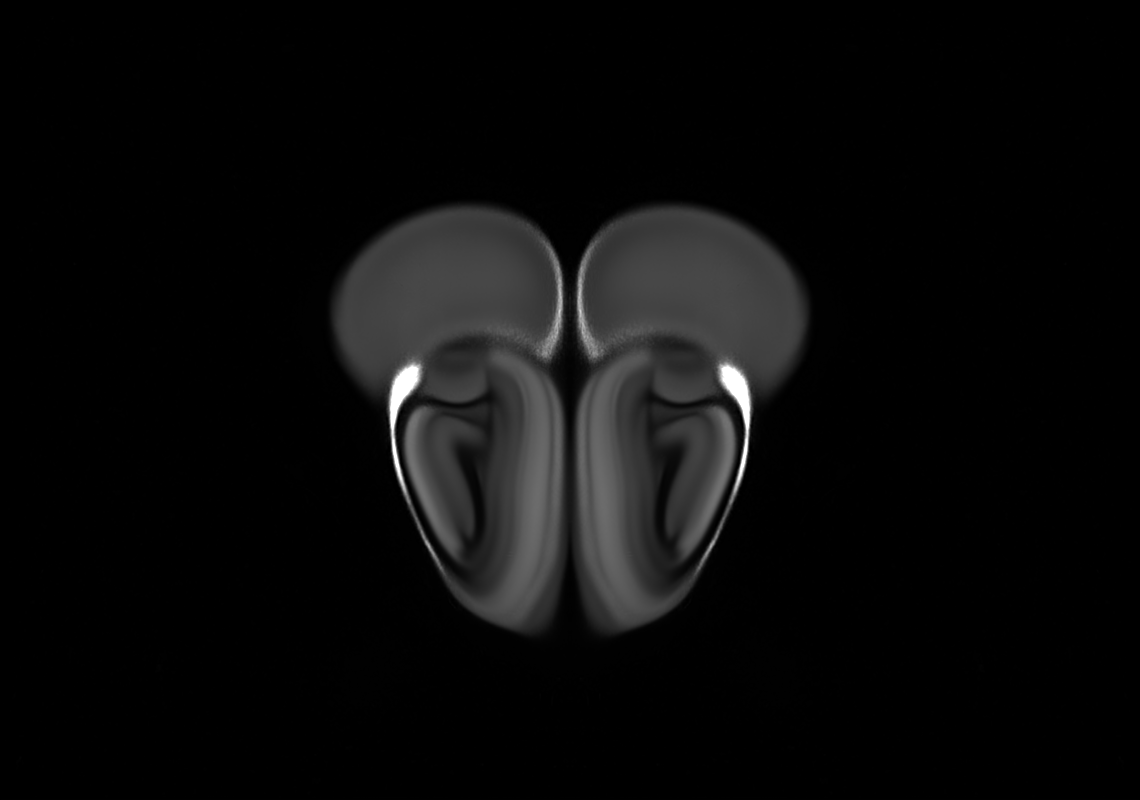

### AllenCCF_Z014.tif

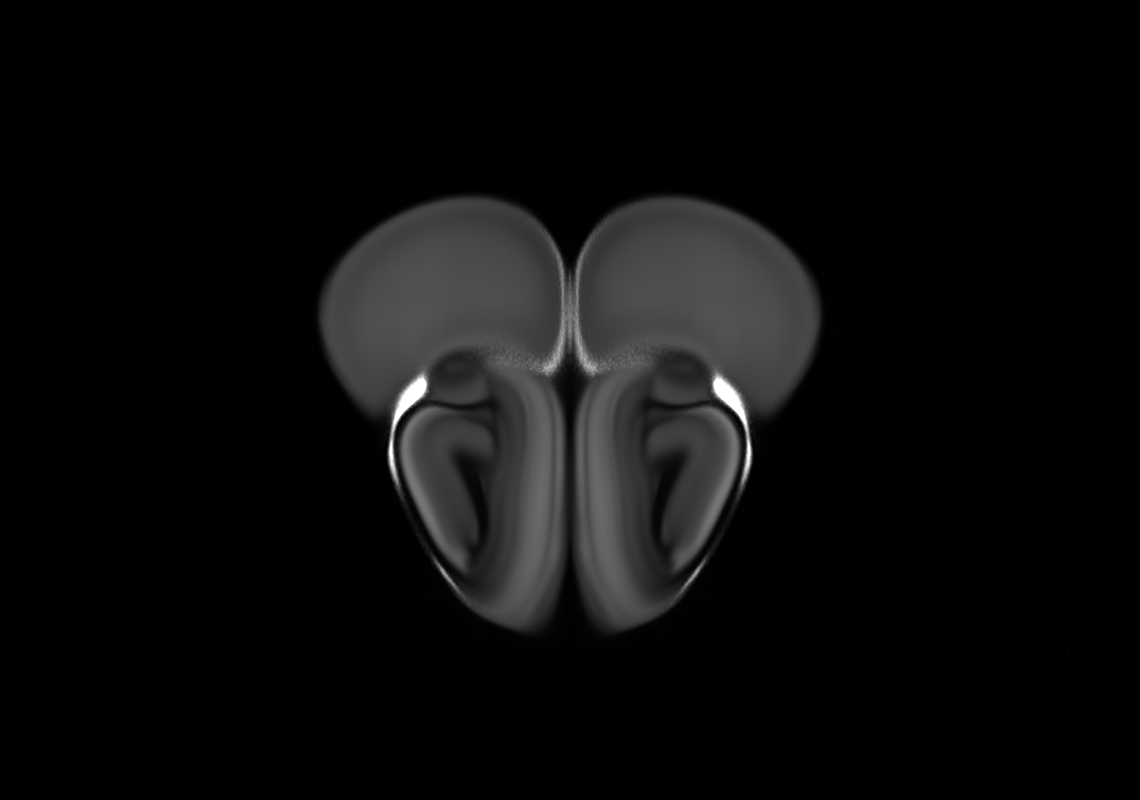

### AllenCCF_Z015.tif

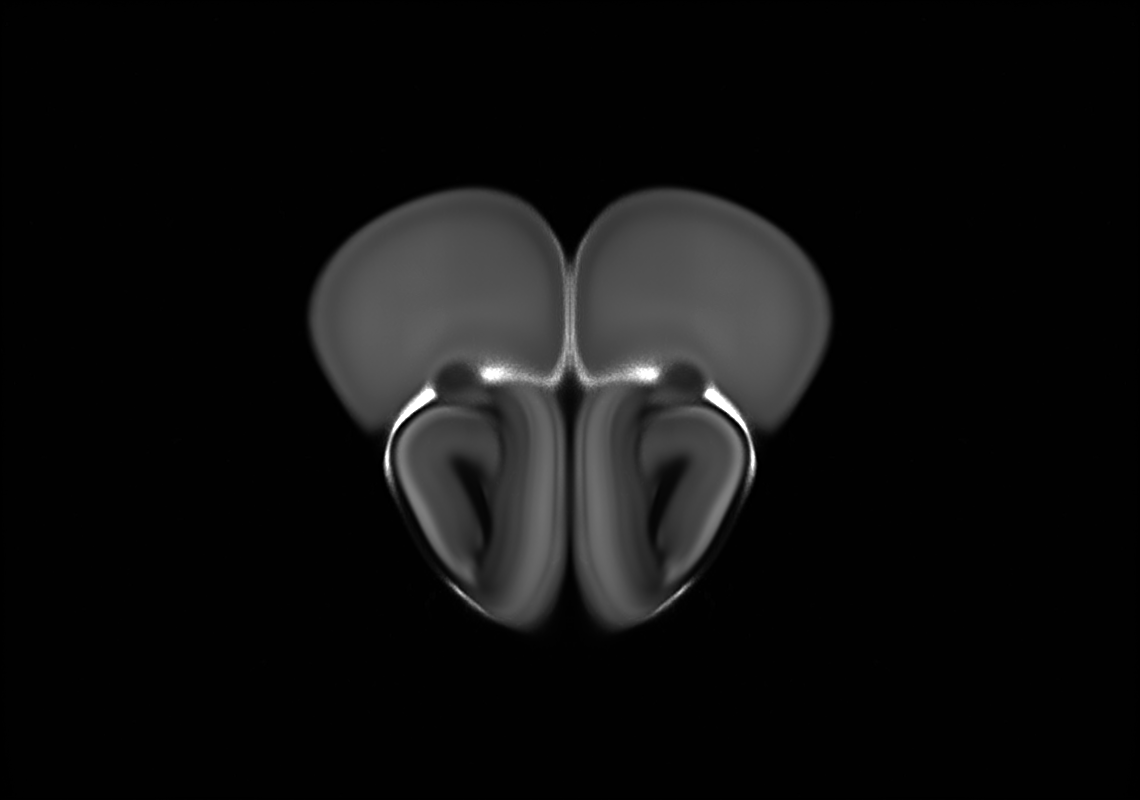

### AllenCCF_Z016.tif

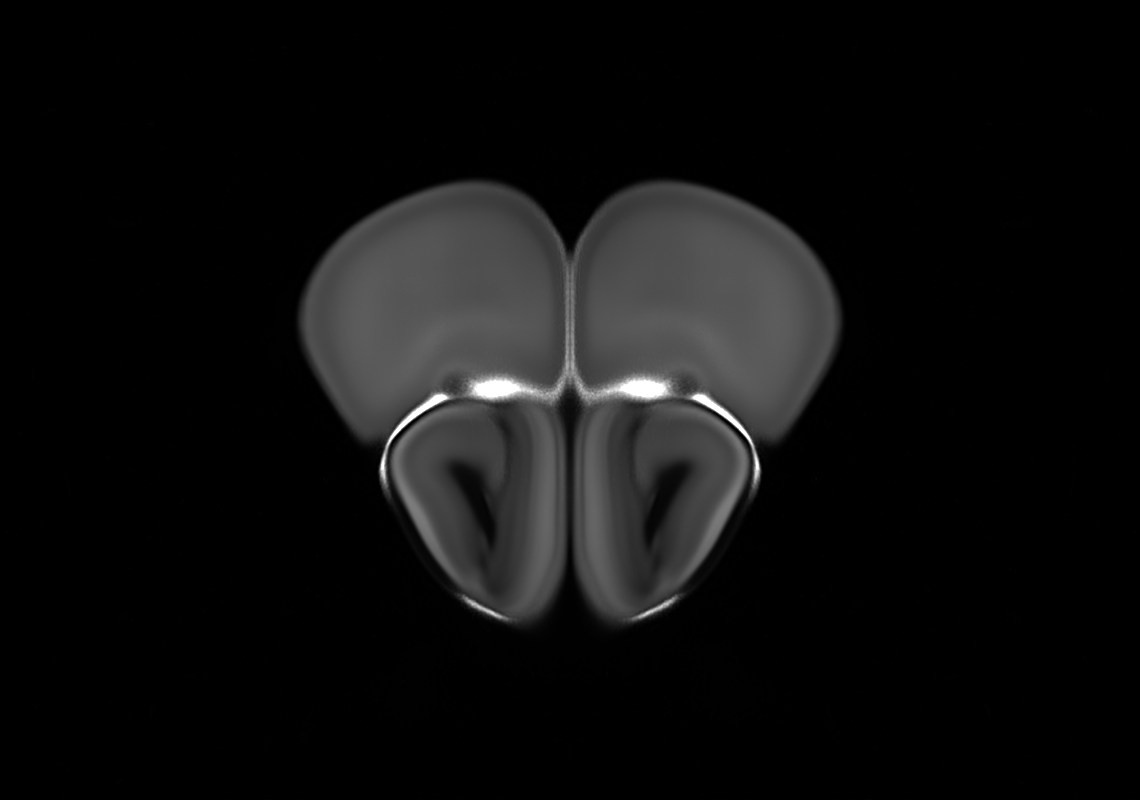

### AllenCCF_Z017.tif

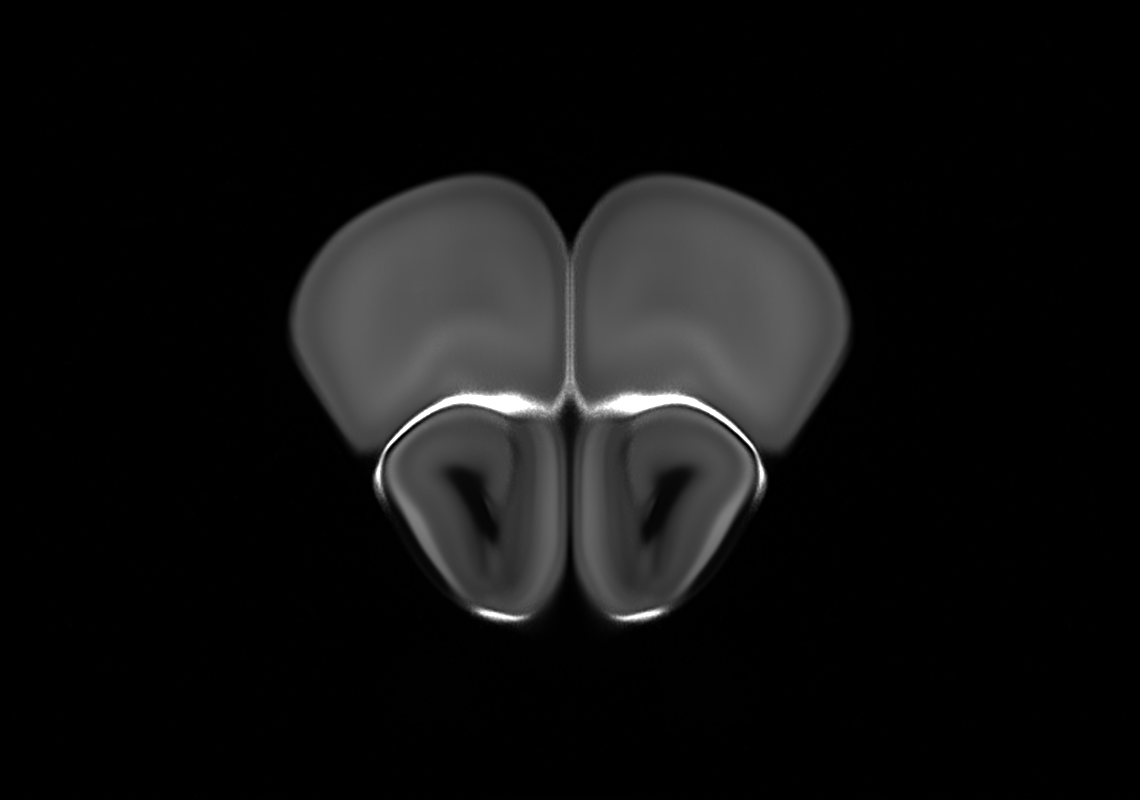

### AllenCCF_Z018.tif

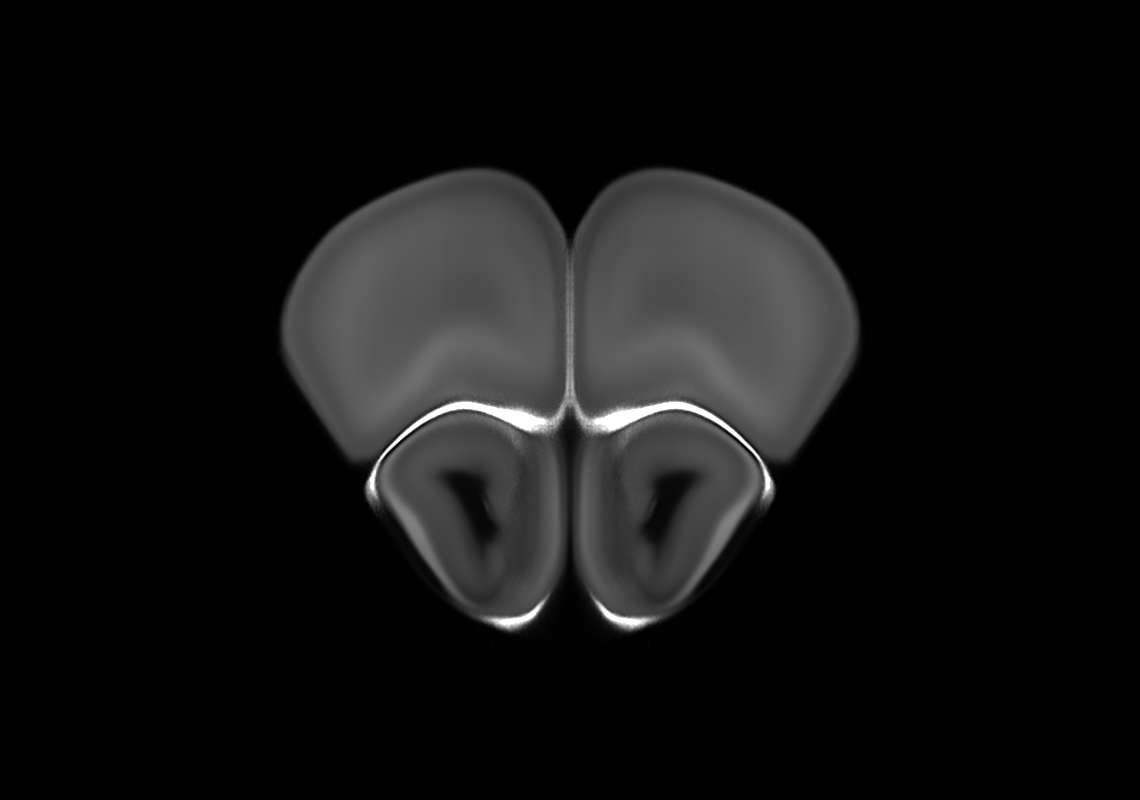

### AllenCCF_Z019.tif

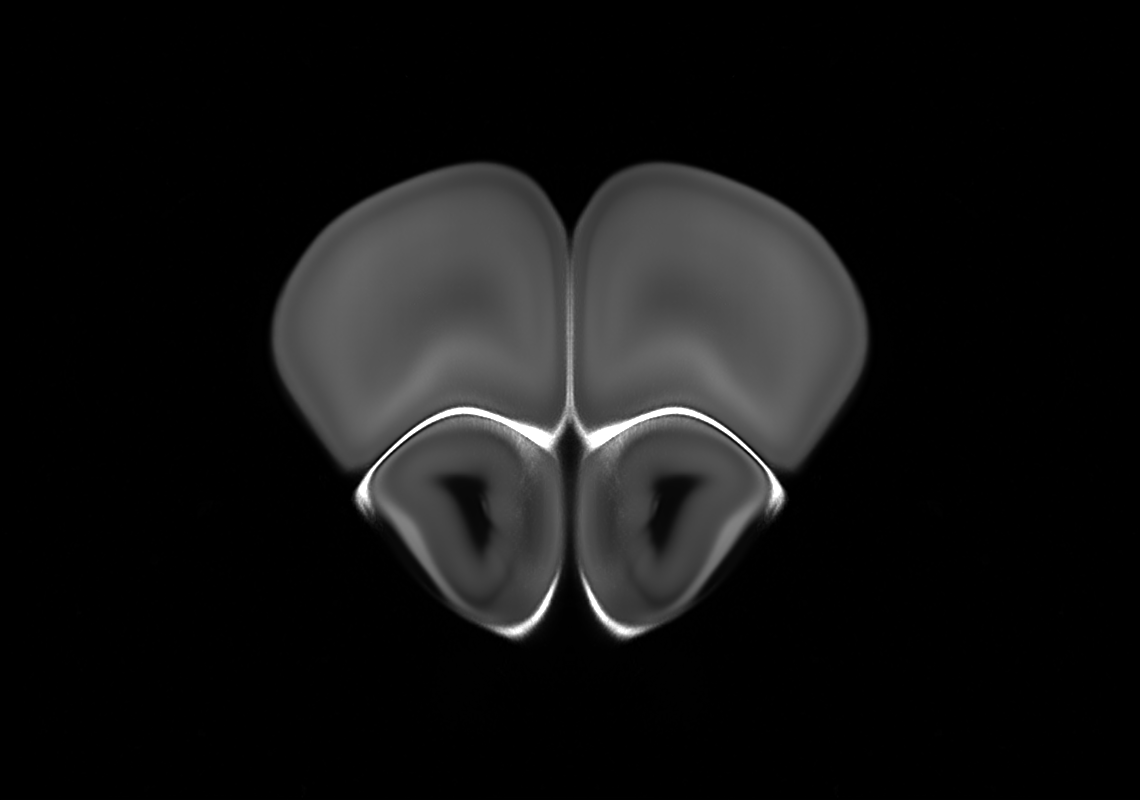

### AllenCCF_Z020.tif

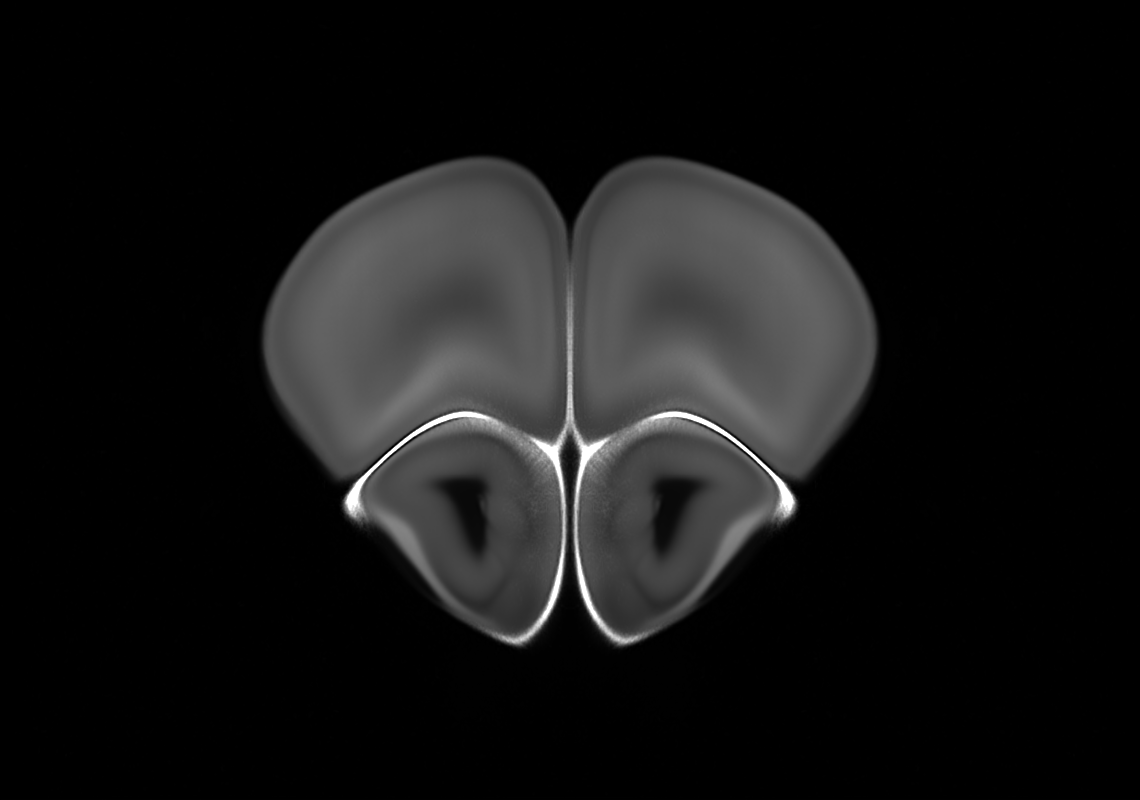

### AllenCCF_Z021.tif

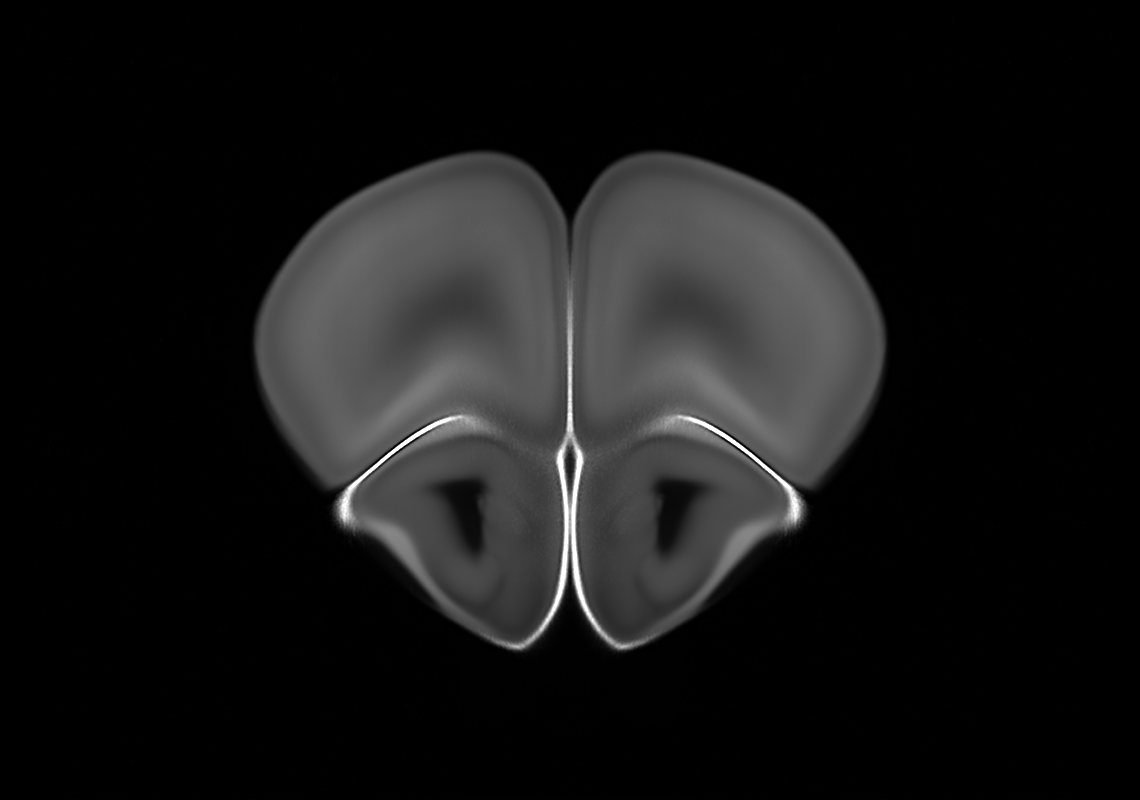

### AllenCCF_Z022.tif

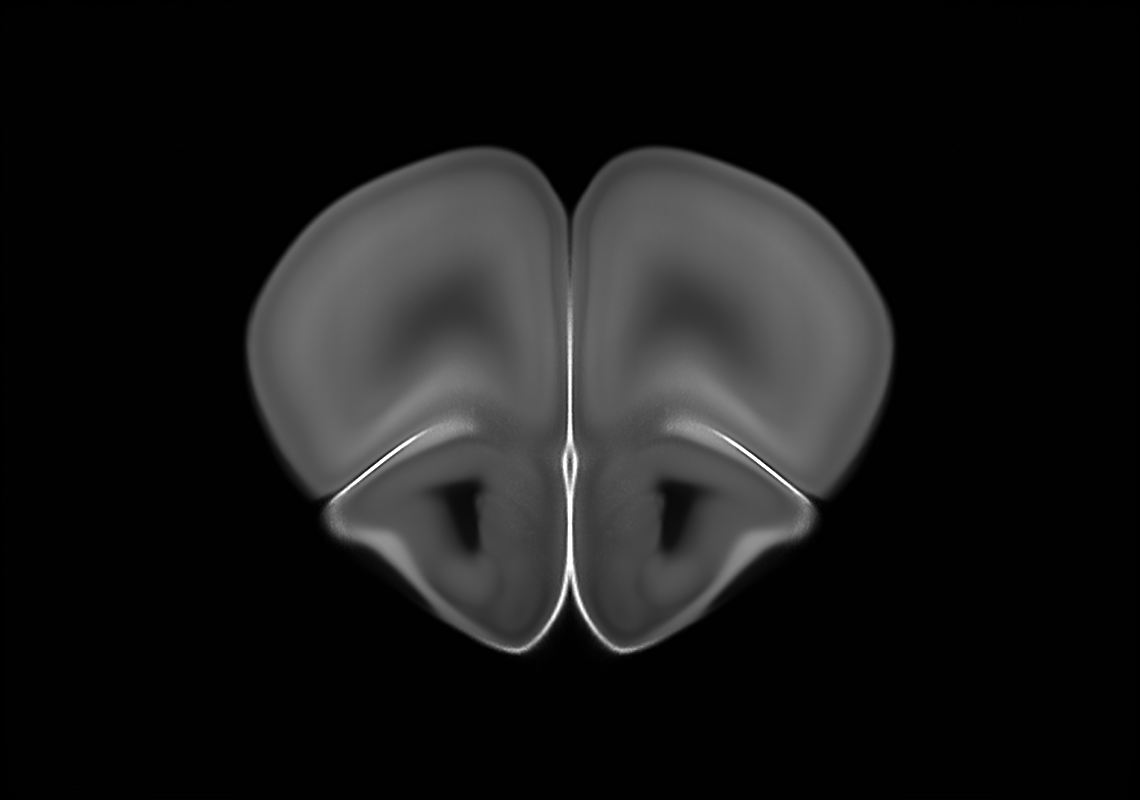

### AllenCCF_Z023.tif

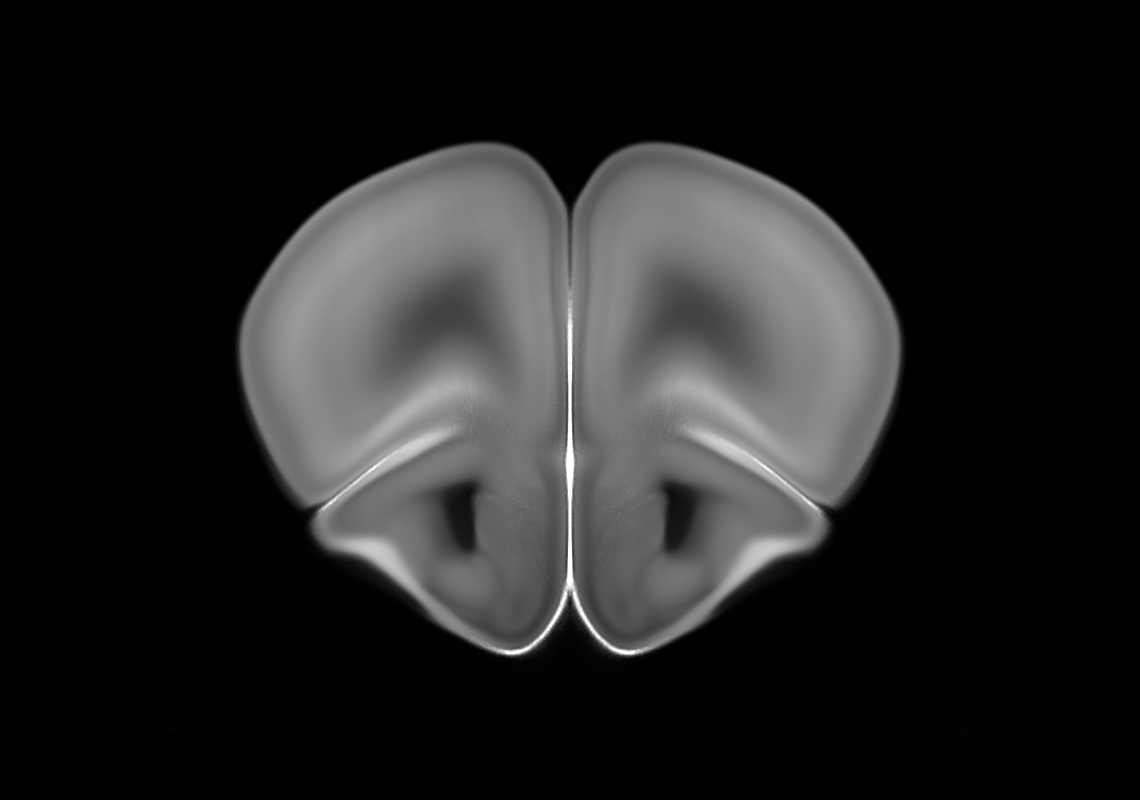

### AllenCCF_Z024.tif

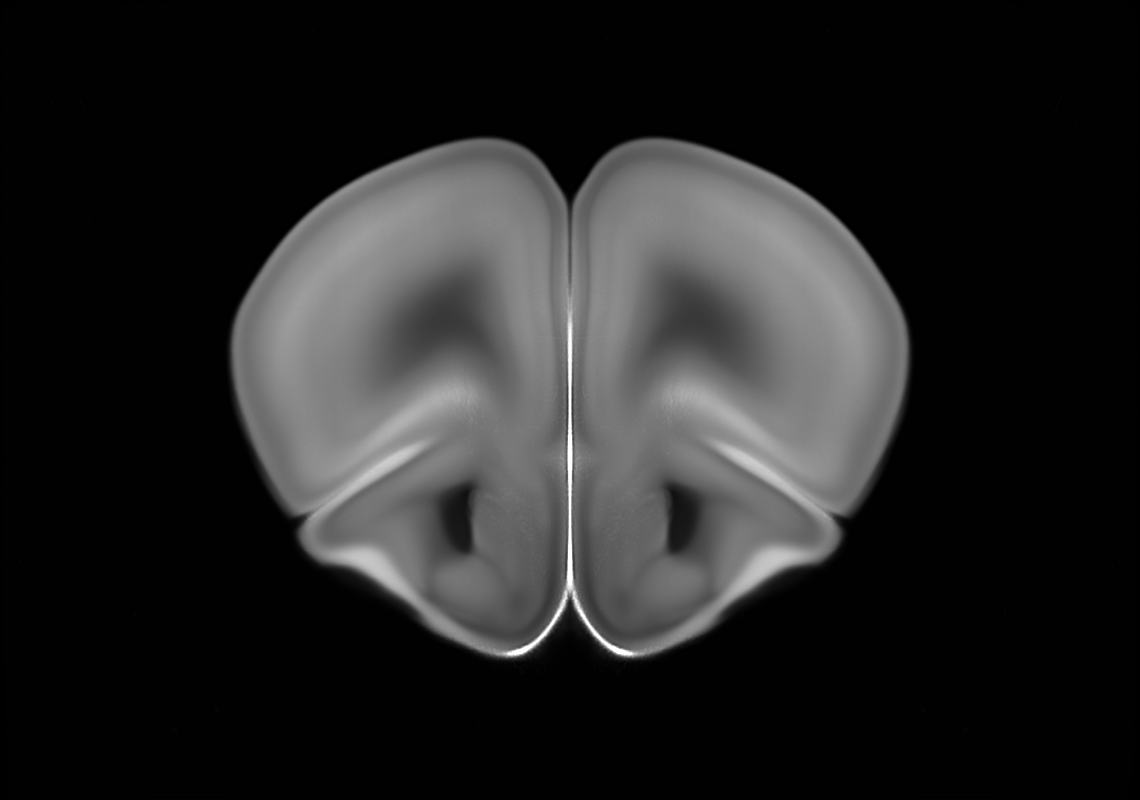

### AllenCCF_Z025.tif

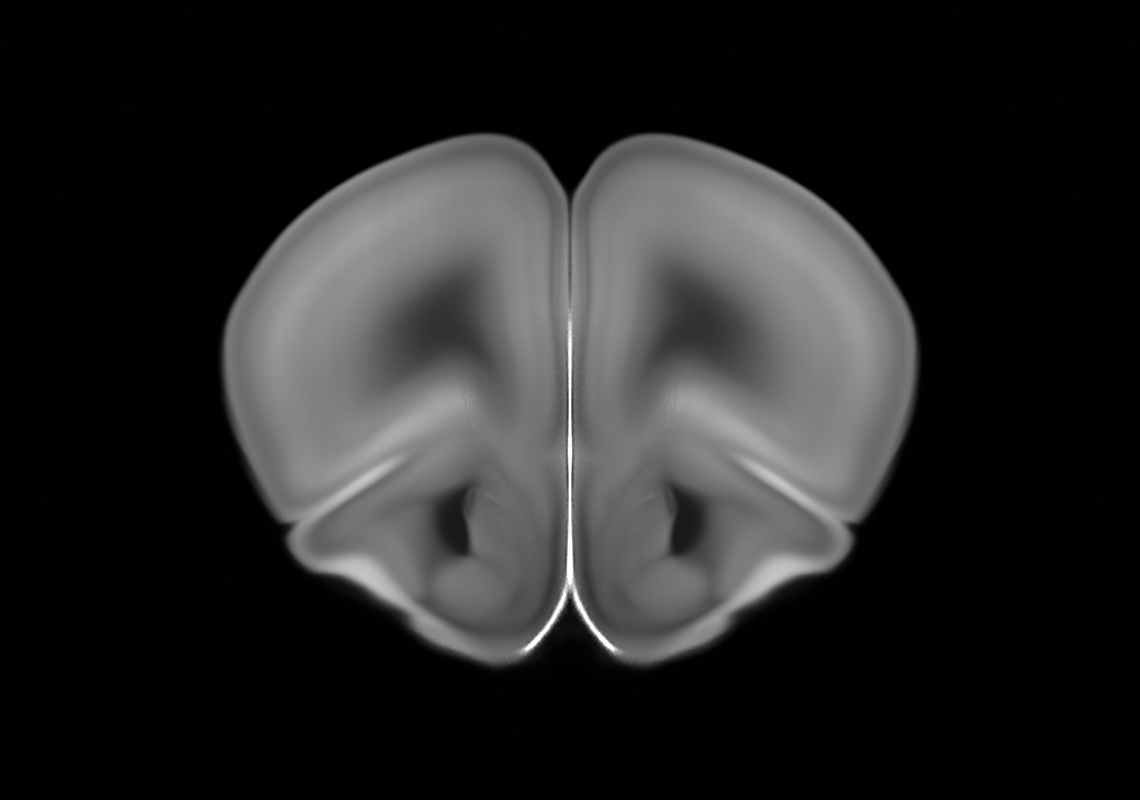

### AllenCCF_Z026.tif

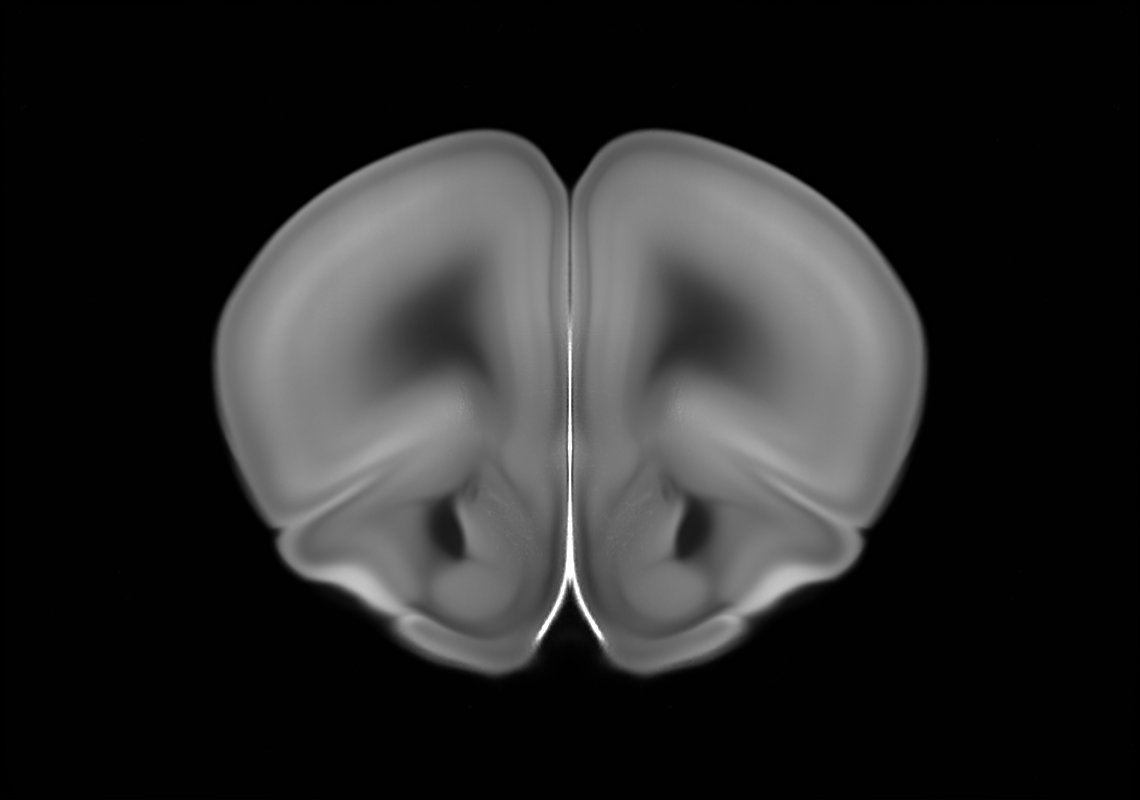

### AllenCCF_Z027.tif

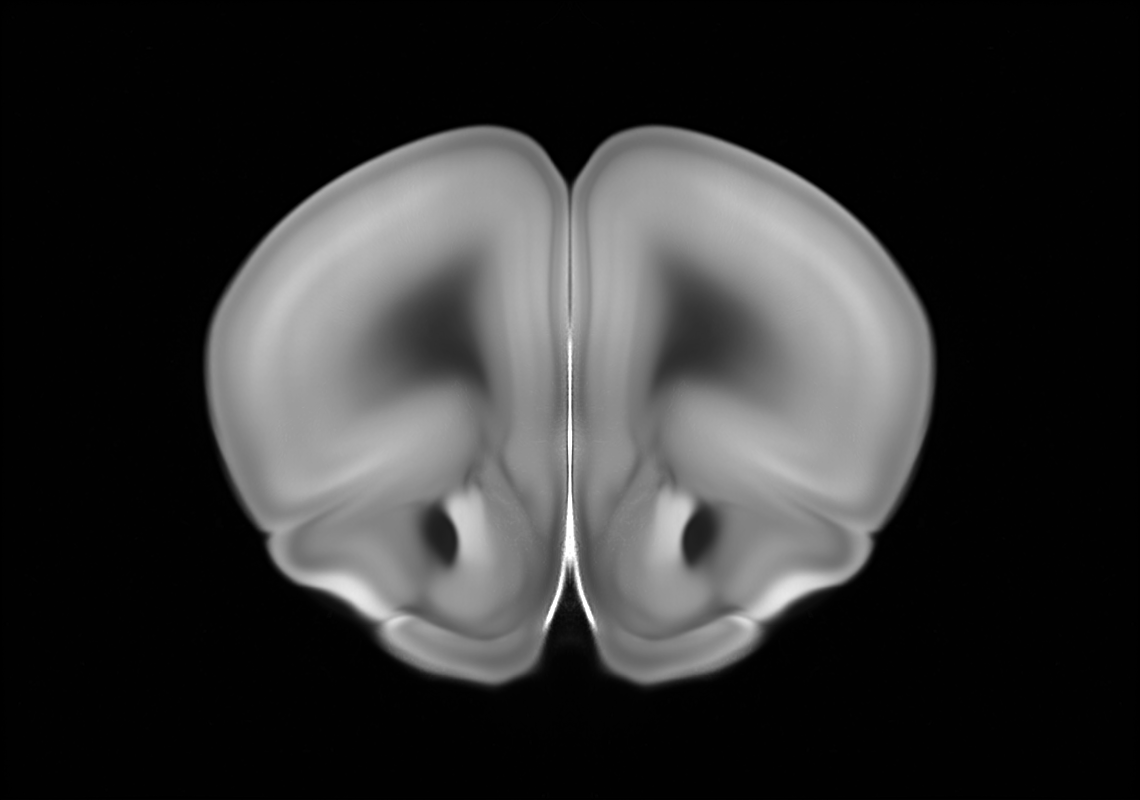

### AllenCCF_Z028.tif

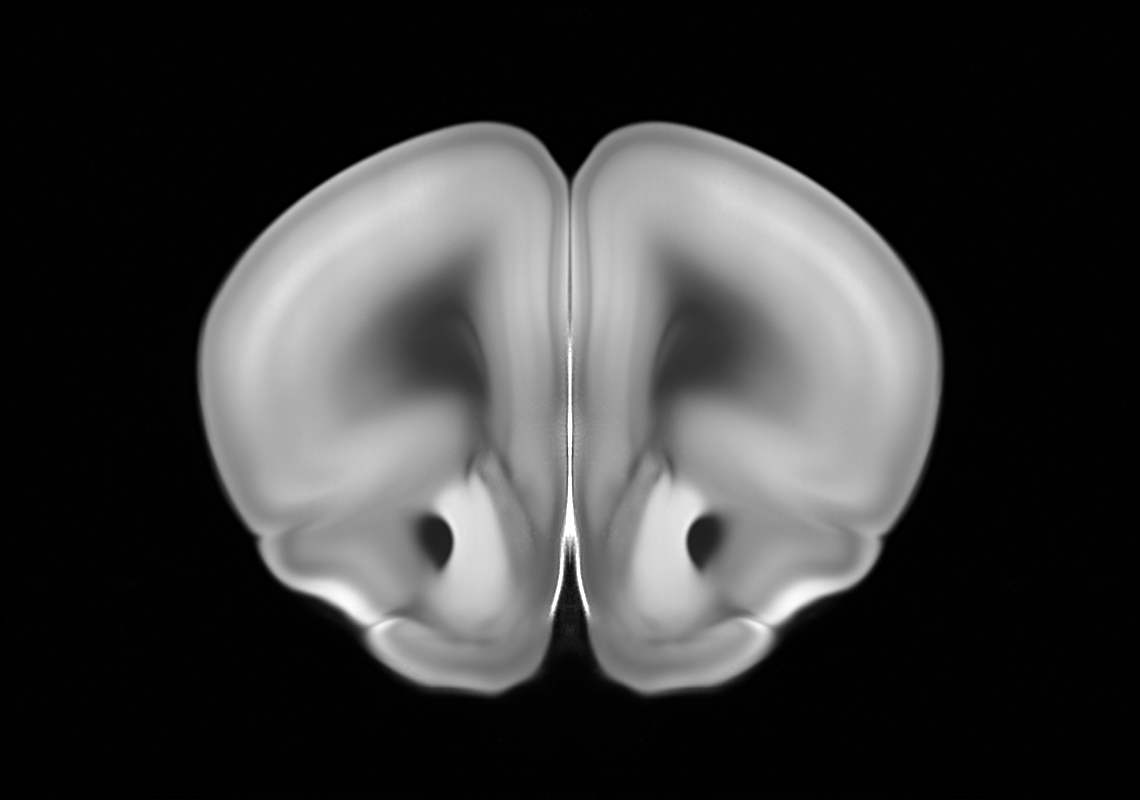

### AllenCCF_Z029.tif

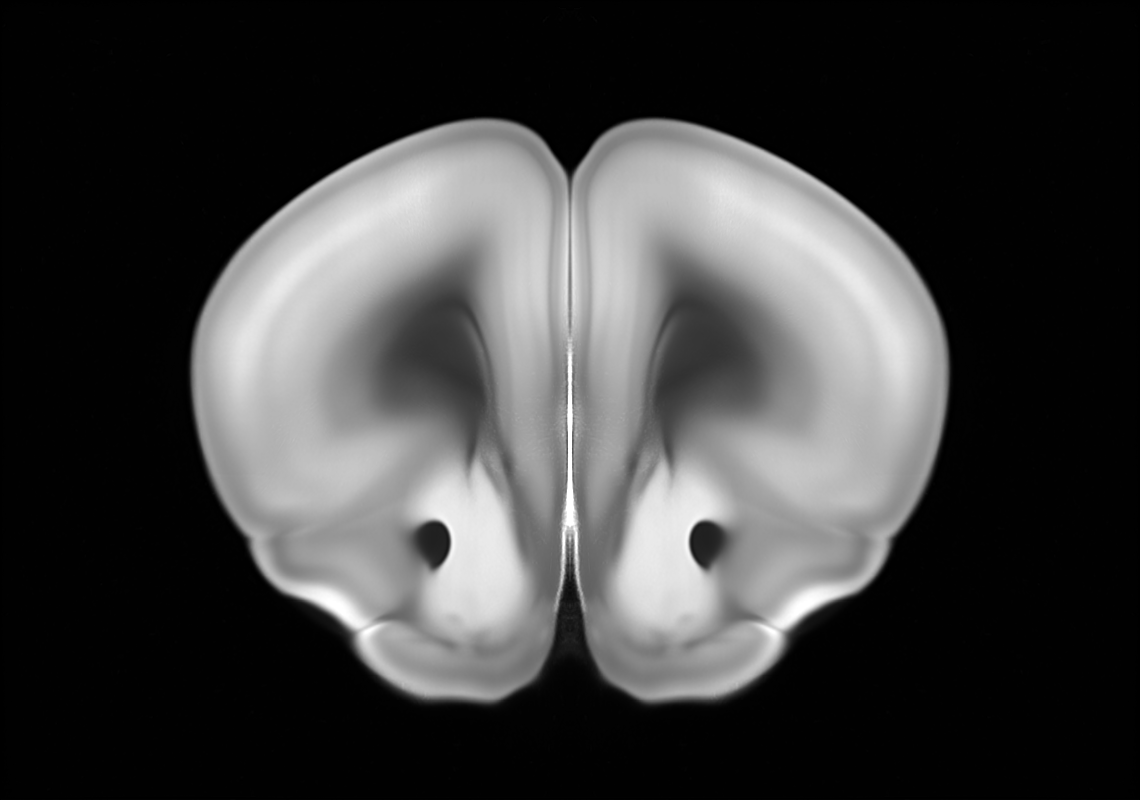

### AllenCCF_Z030.tif

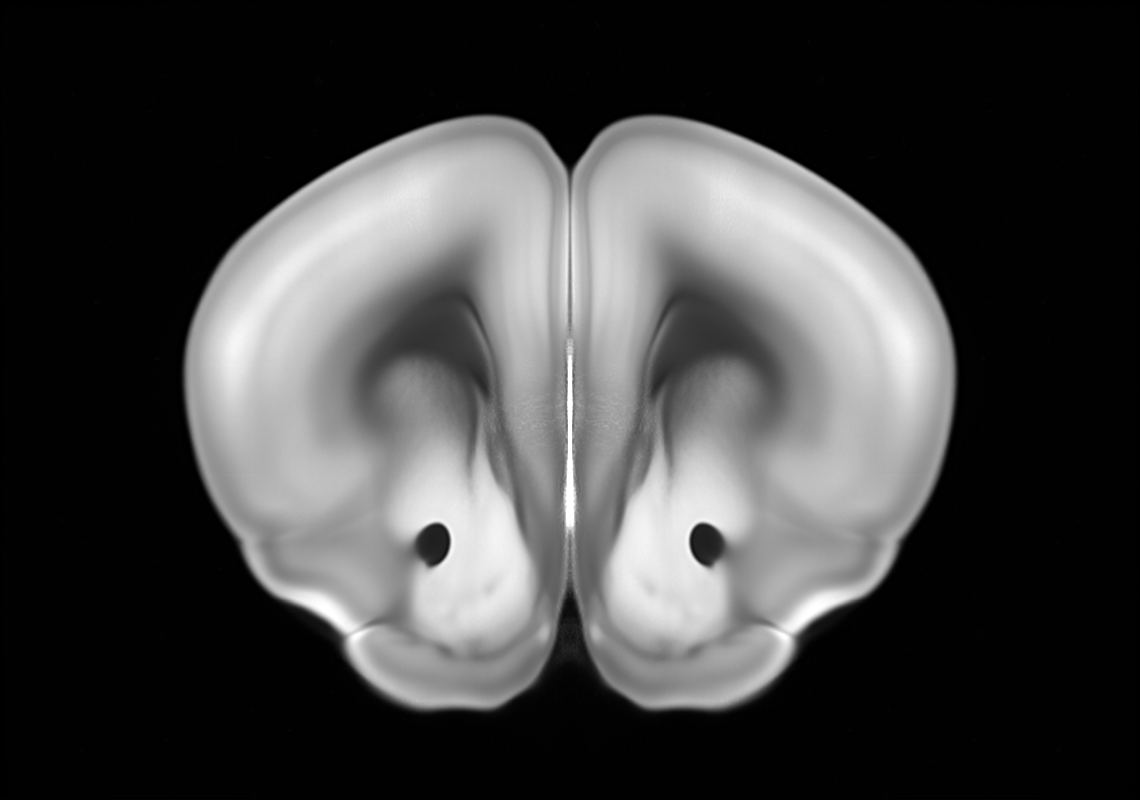

### AllenCCF_Z031.tif

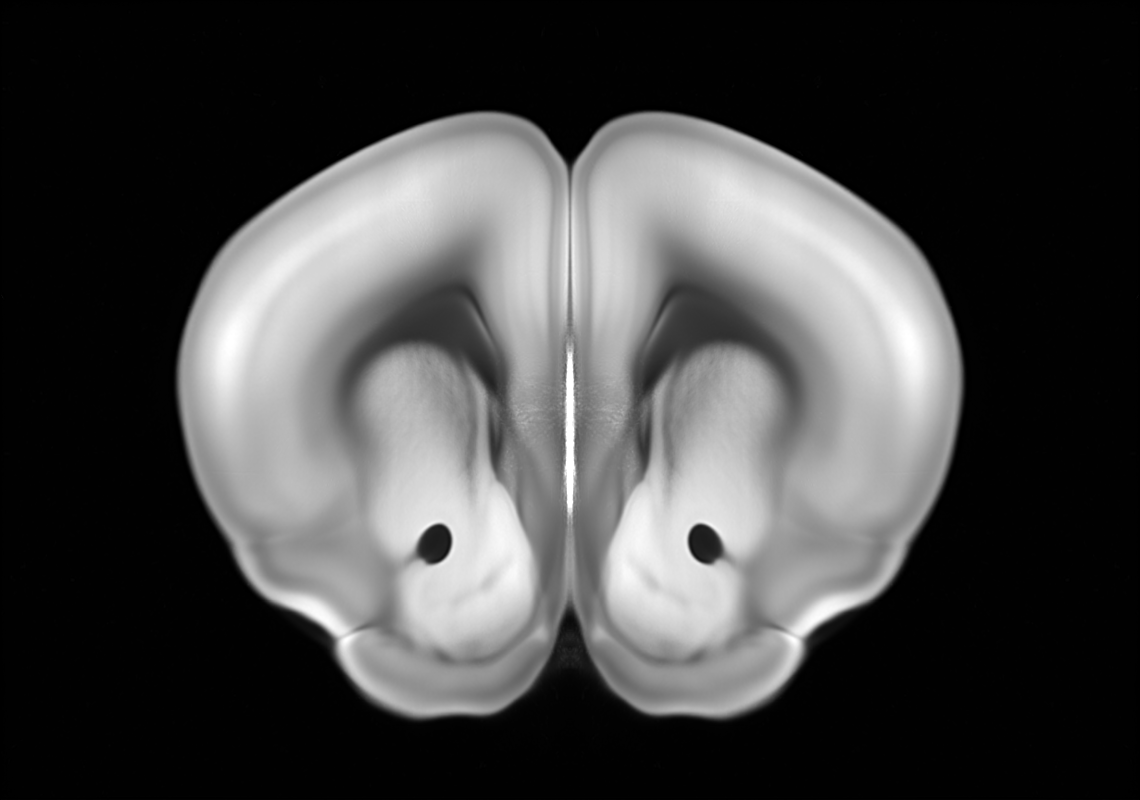

### AllenCCF_Z032.tif

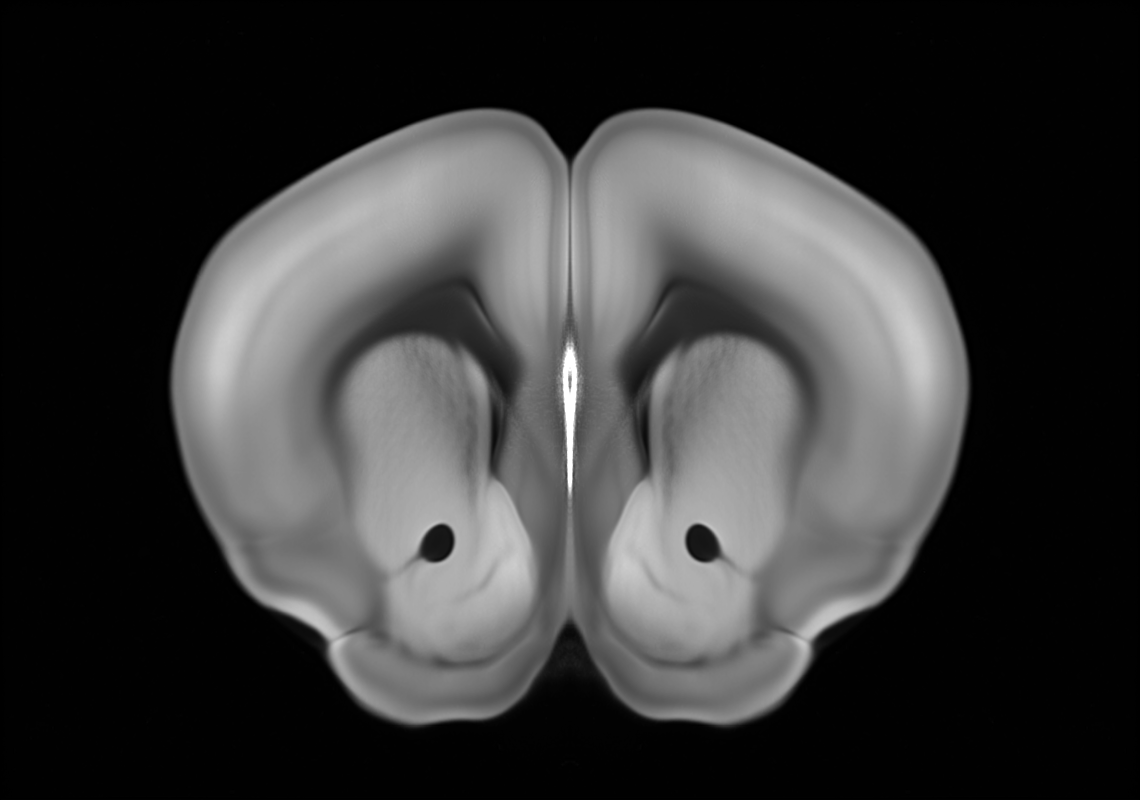

### AllenCCF_Z033.tif

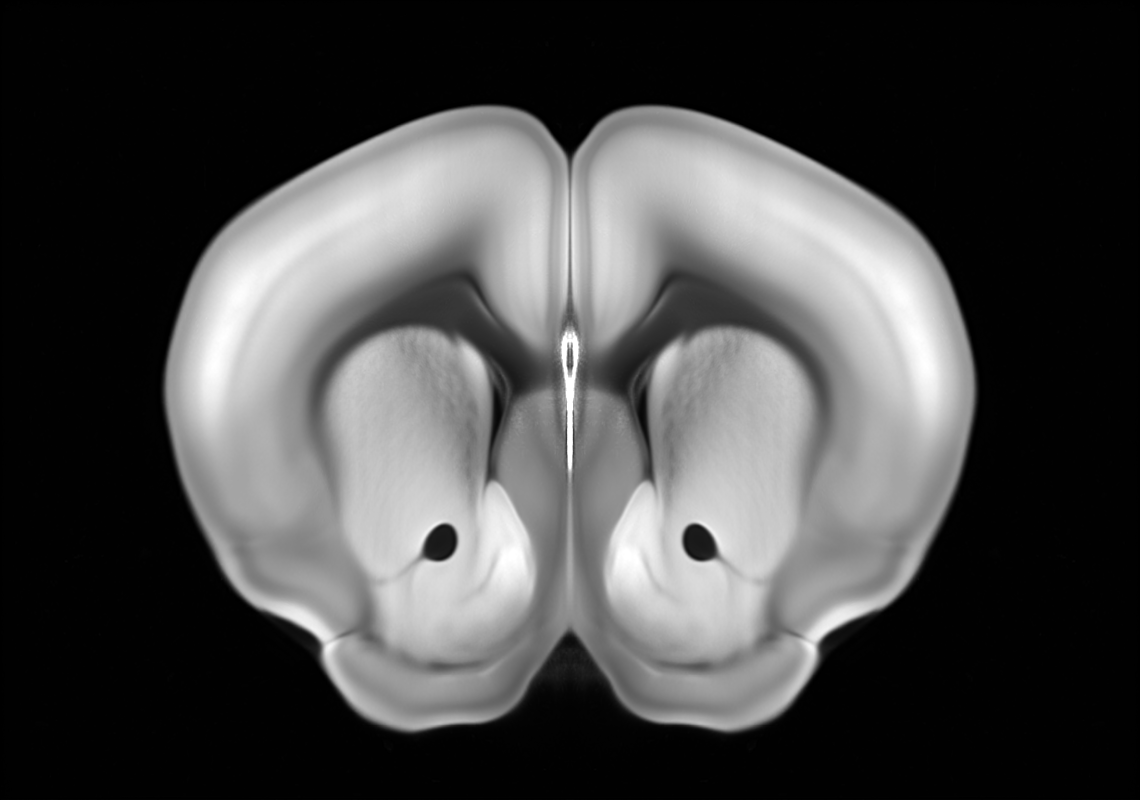

### AllenCCF_Z034.tif

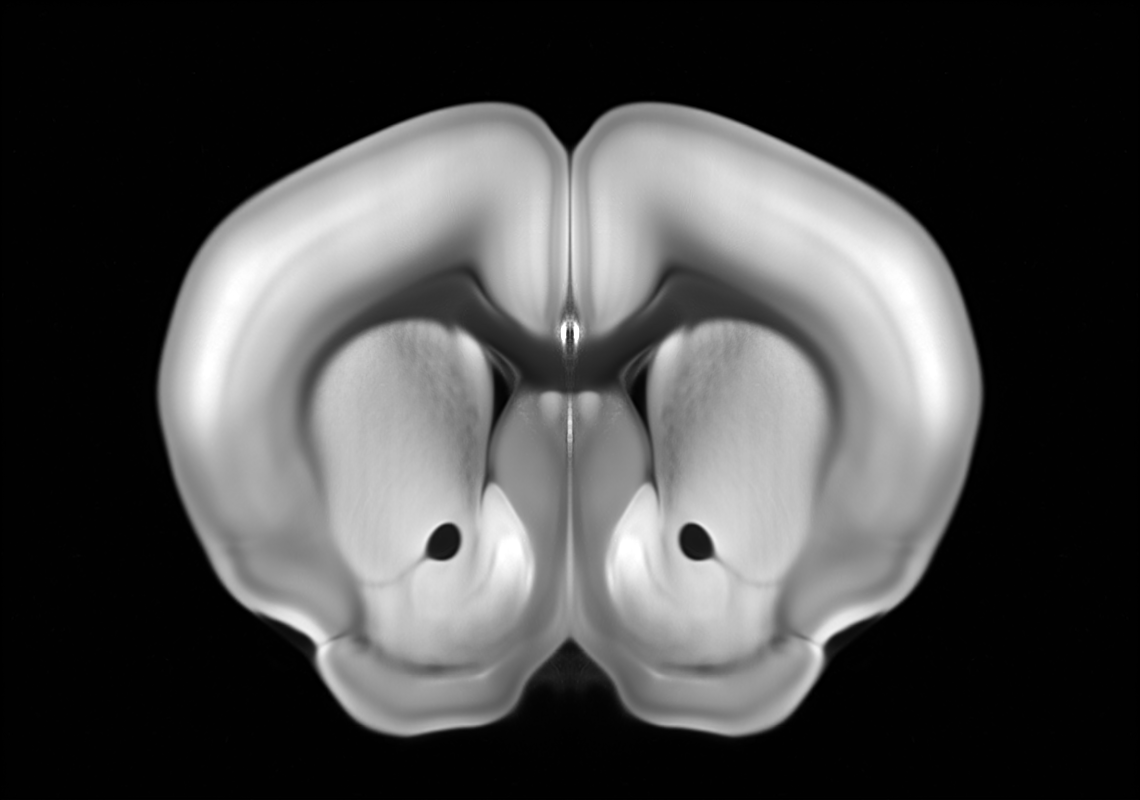

### AllenCCF_Z035.tif

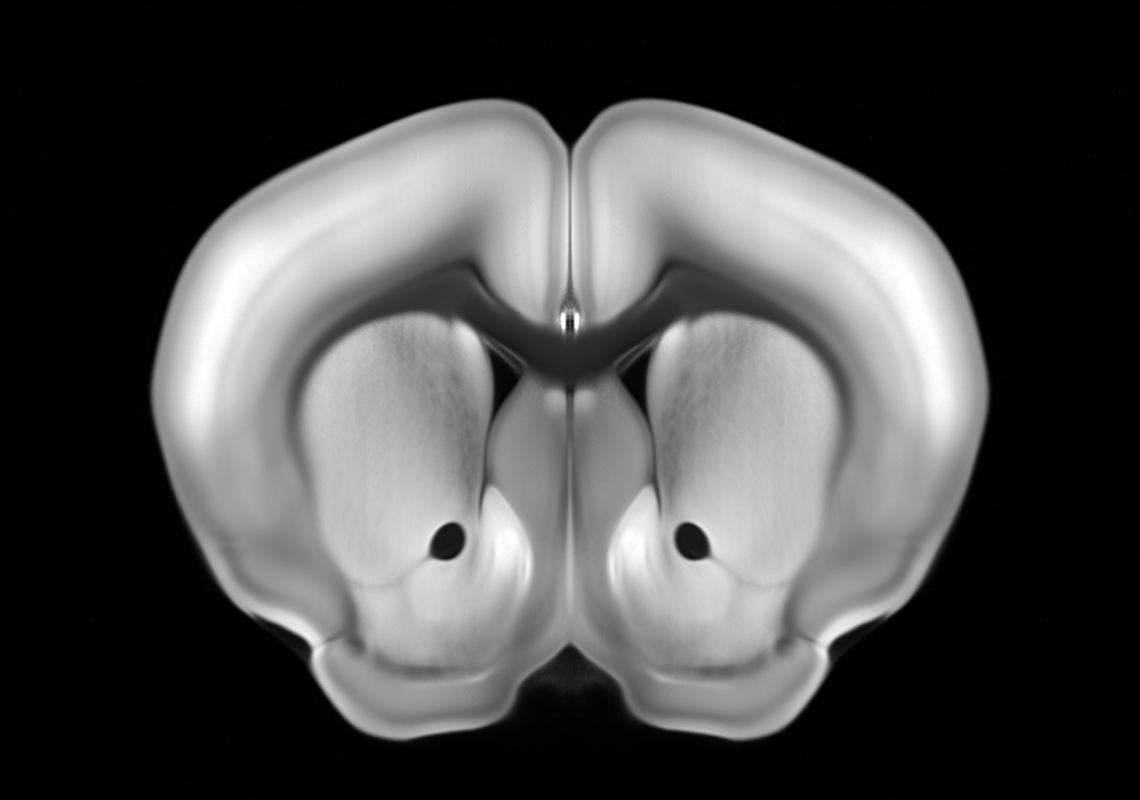

### AllenCCF_Z036.tif

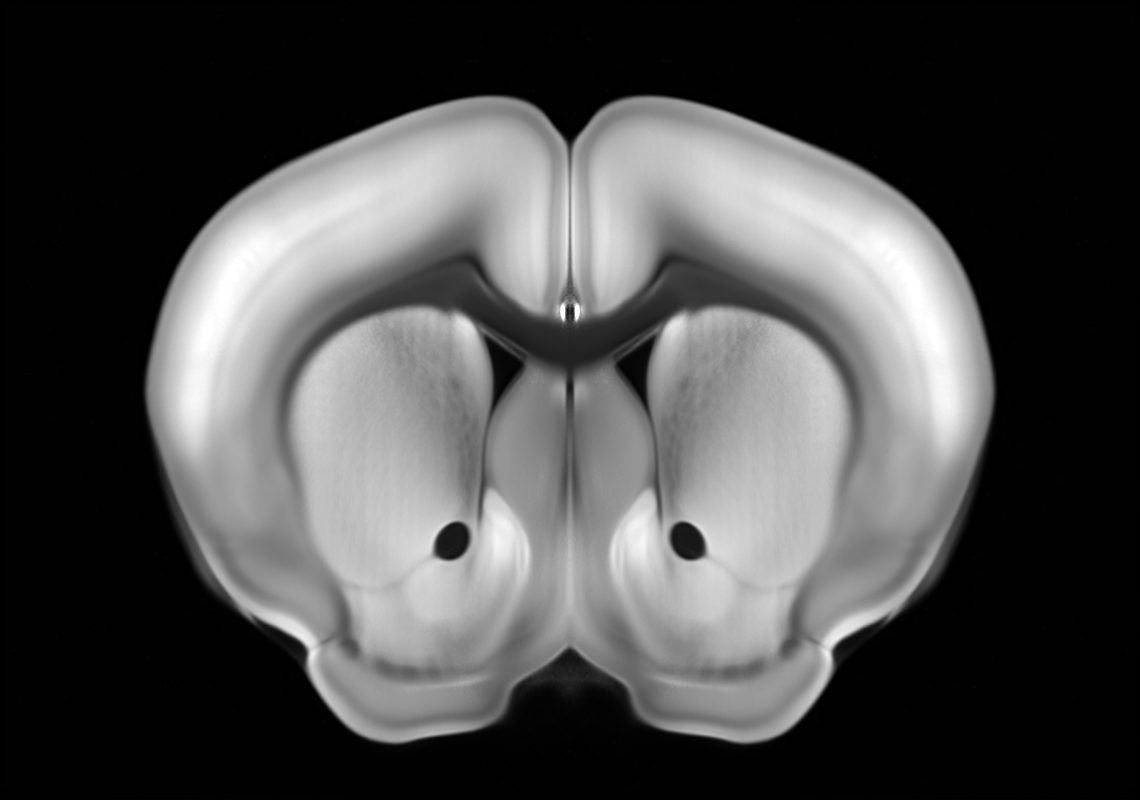

### AllenCCF_Z037.tif

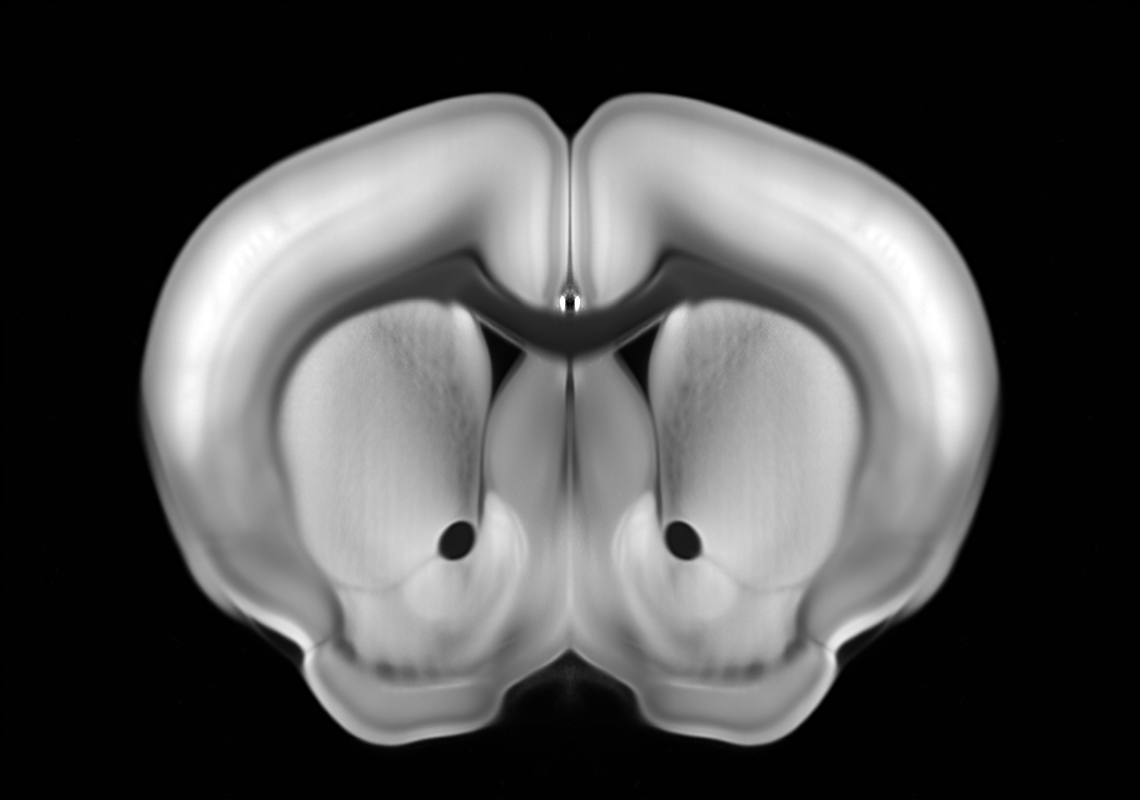

### AllenCCF_Z038.tif

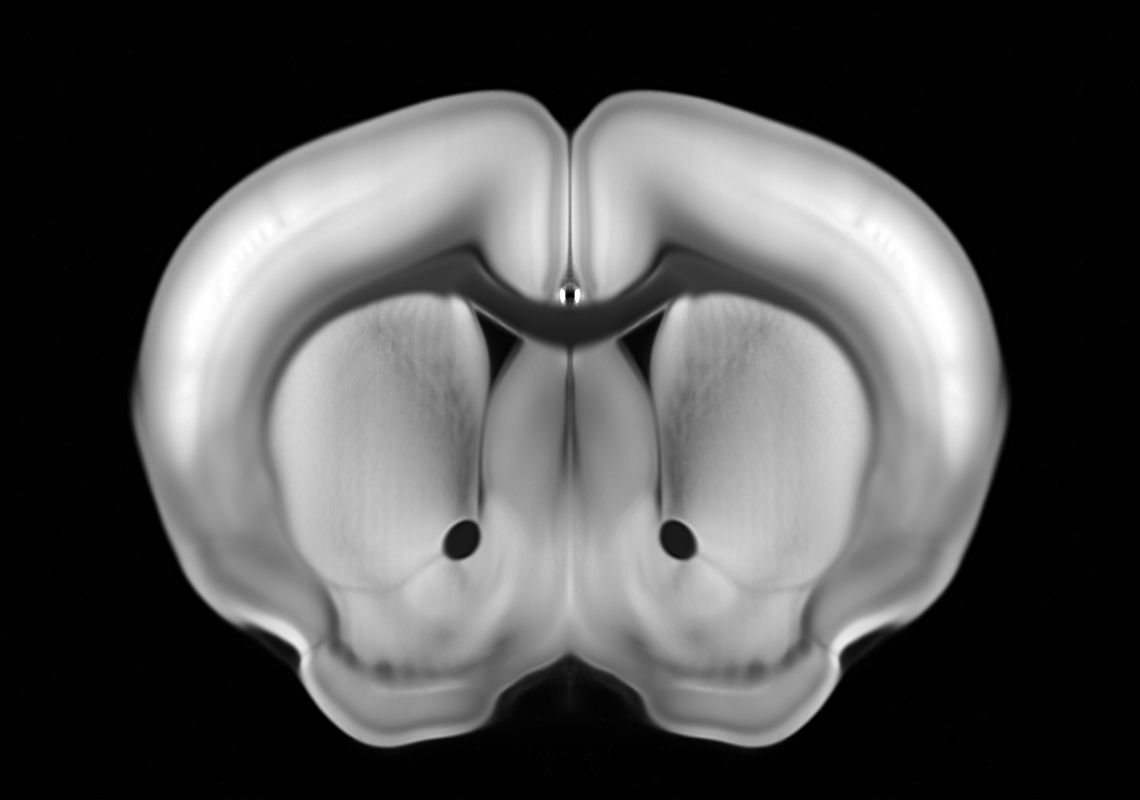

### AllenCCF_Z039.tif

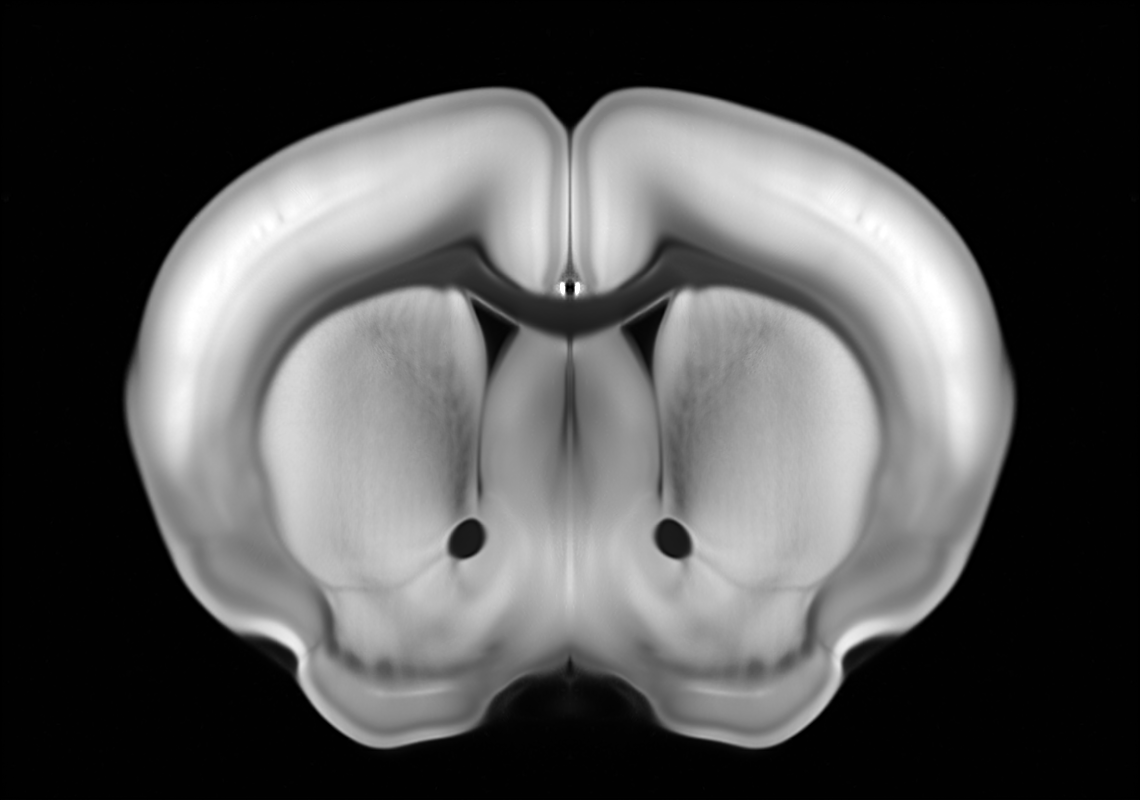

### AllenCCF_Z040.tif

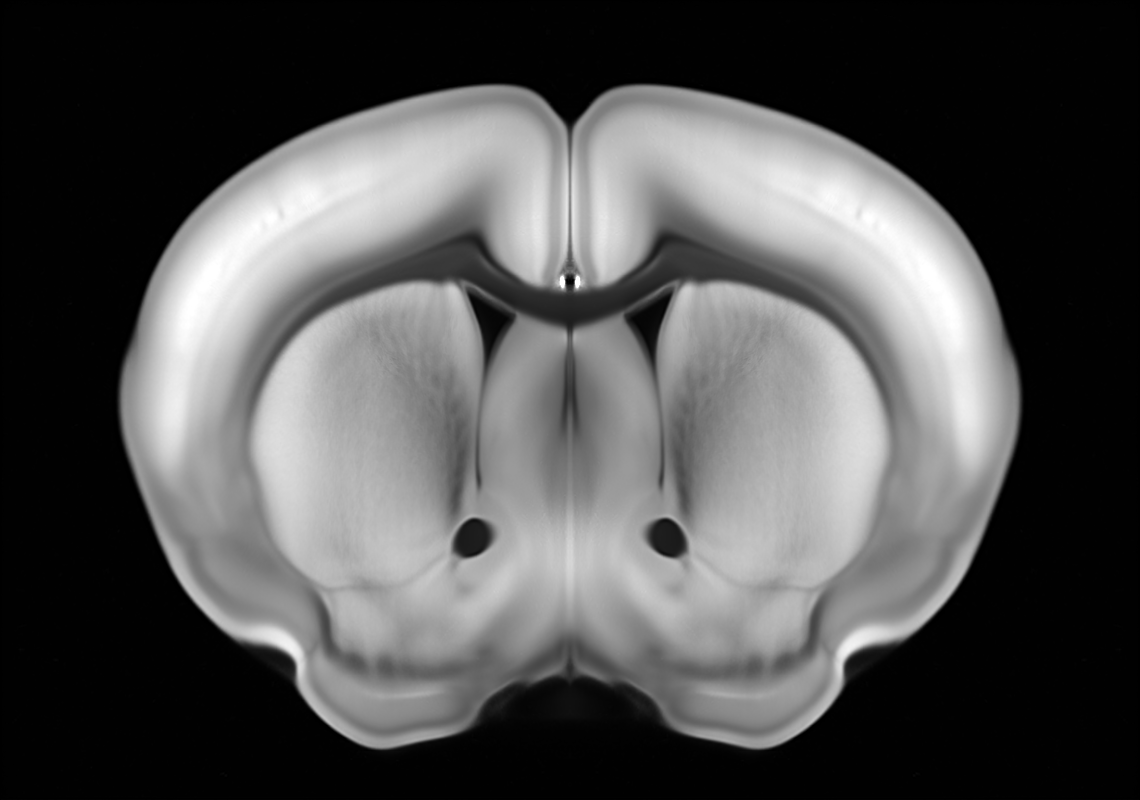

### AllenCCF_Z041.tif

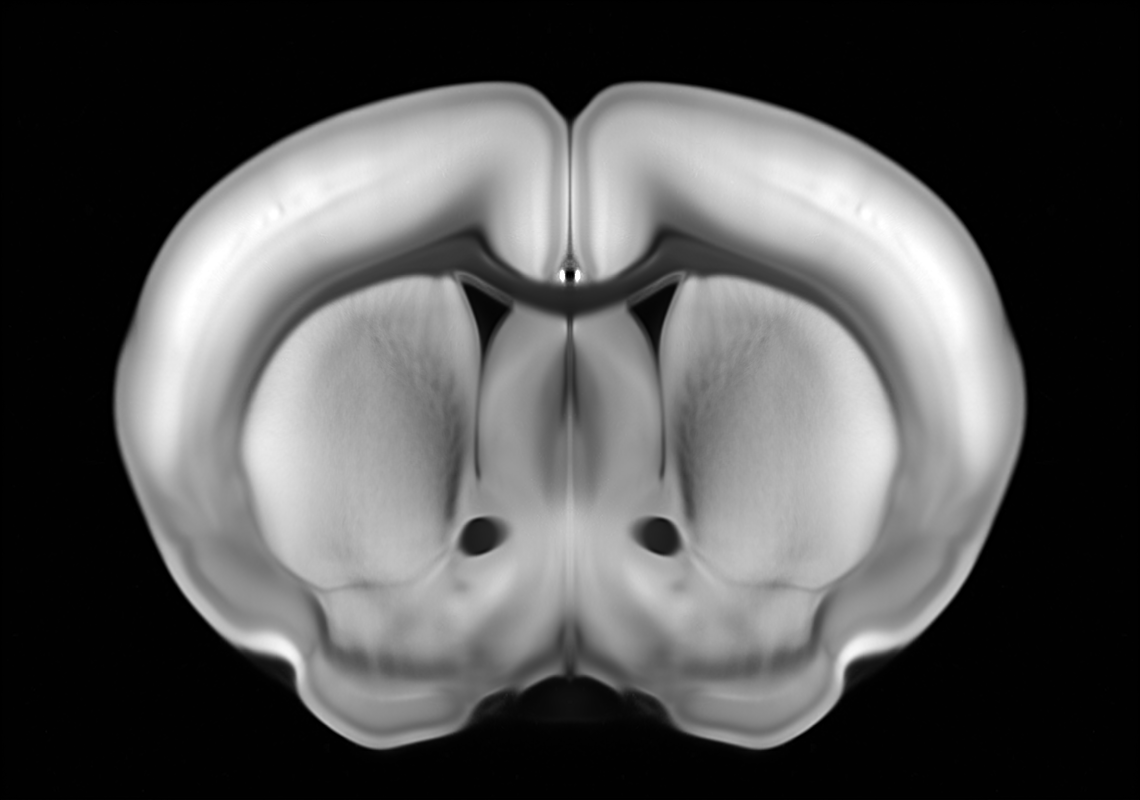

### AllenCCF_Z042.tif

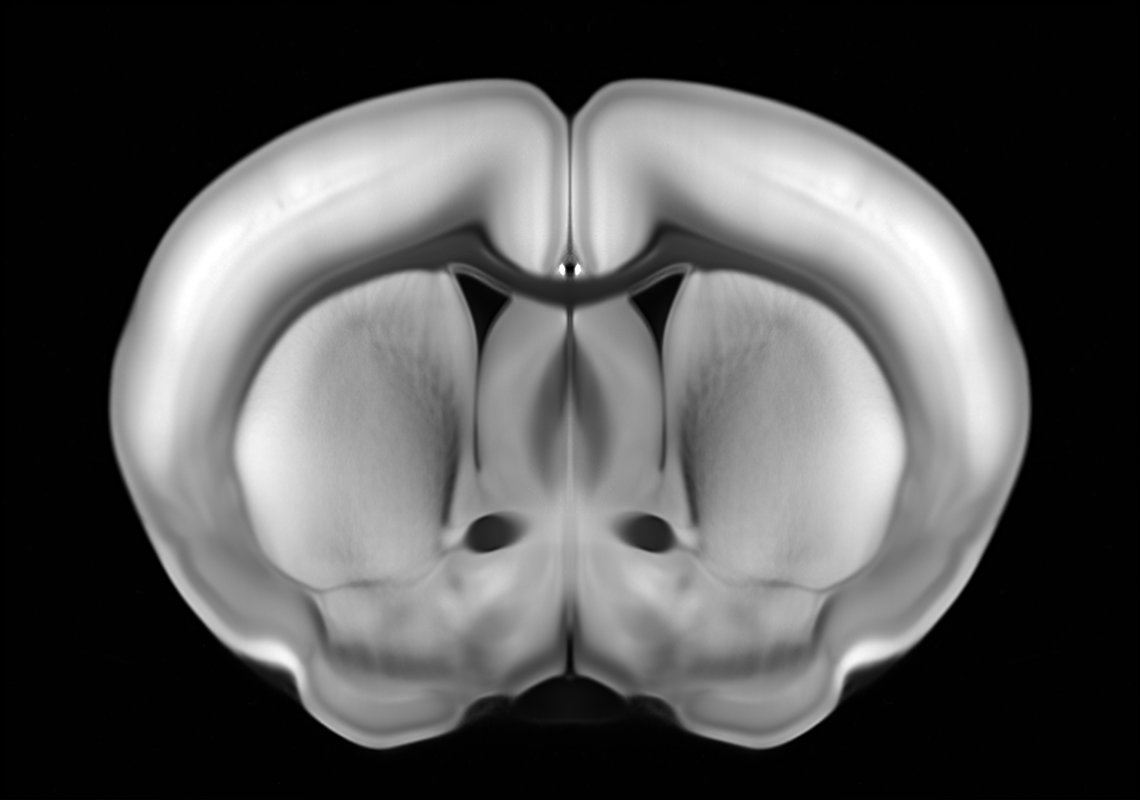

### AllenCCF_Z043.tif

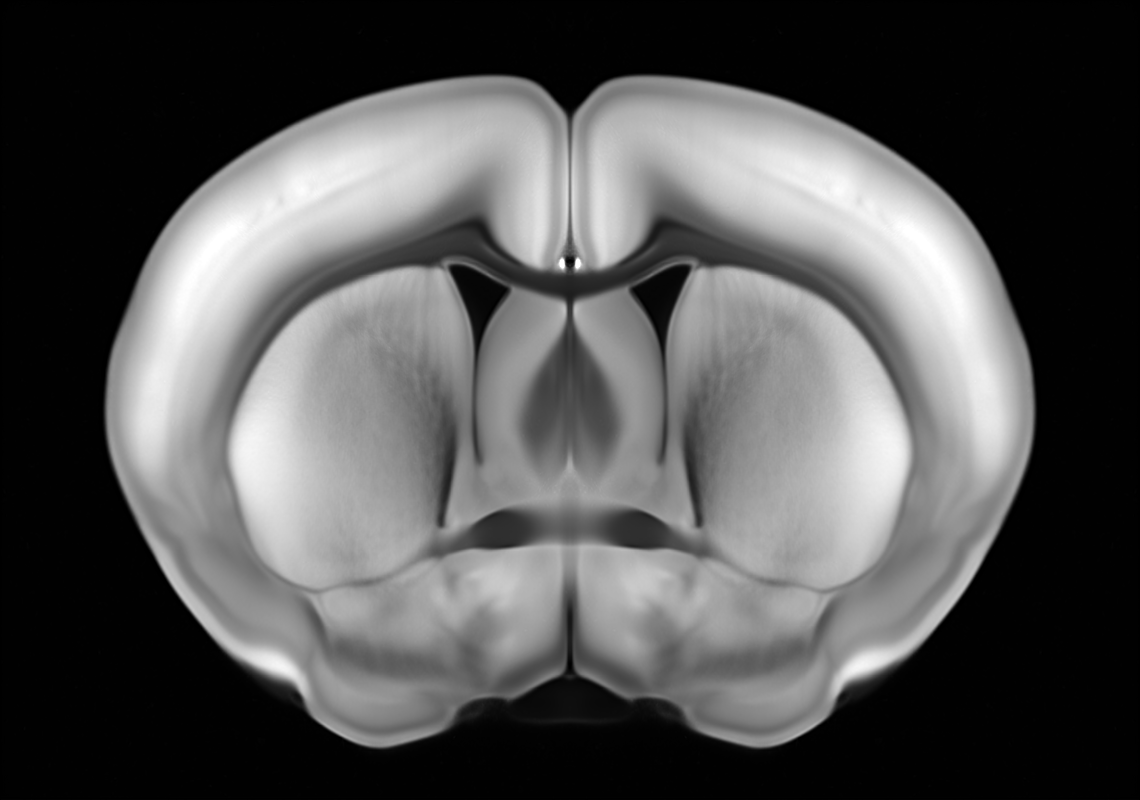

### AllenCCF_Z044.tif

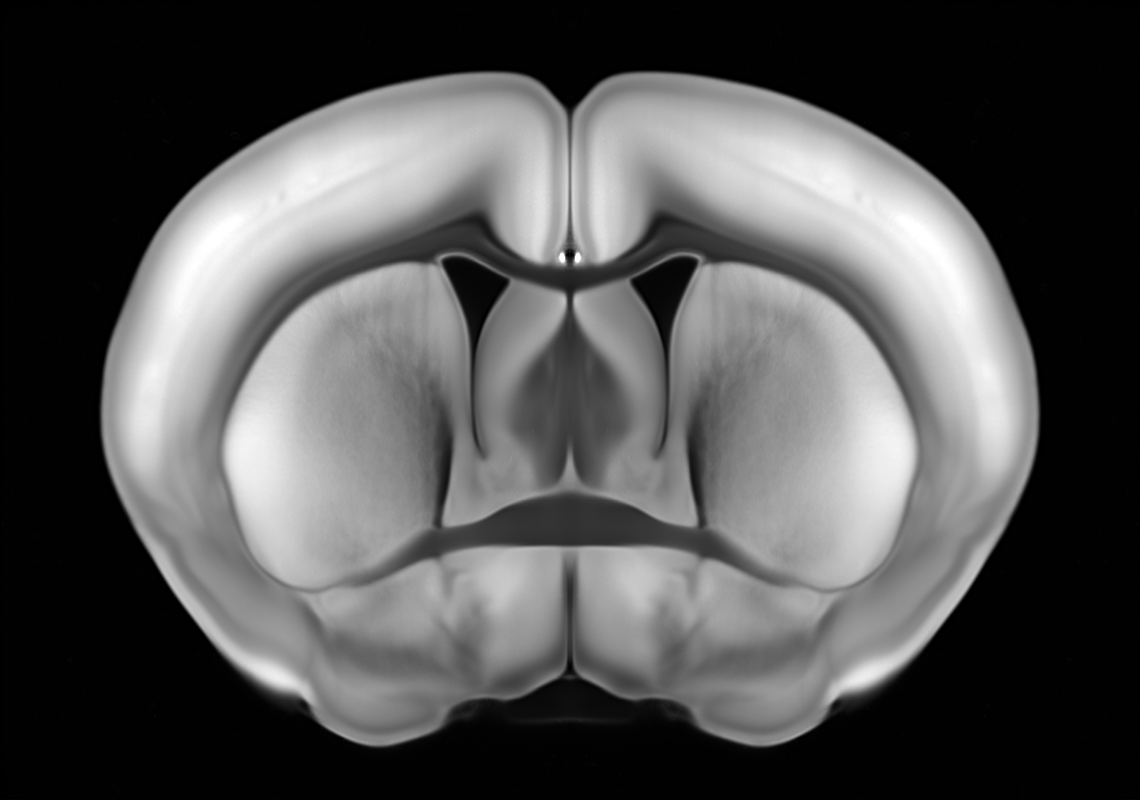

### AllenCCF_Z045.tif

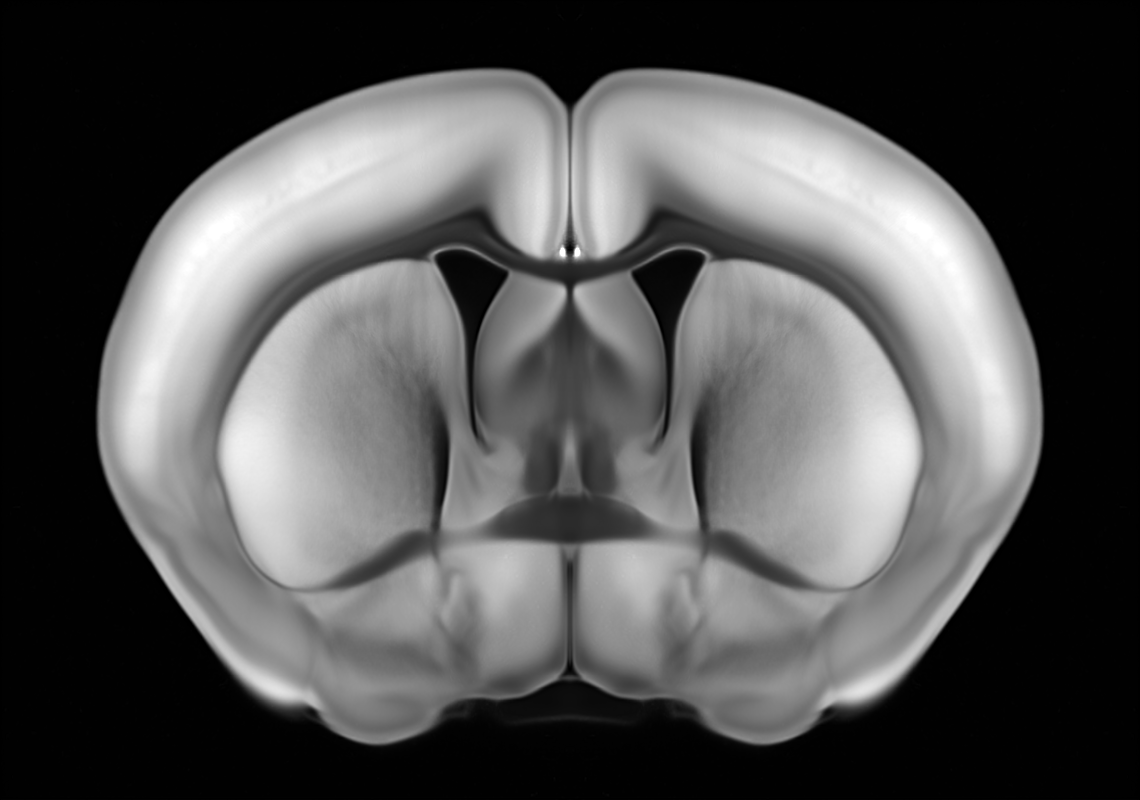

### AllenCCF_Z046.tif

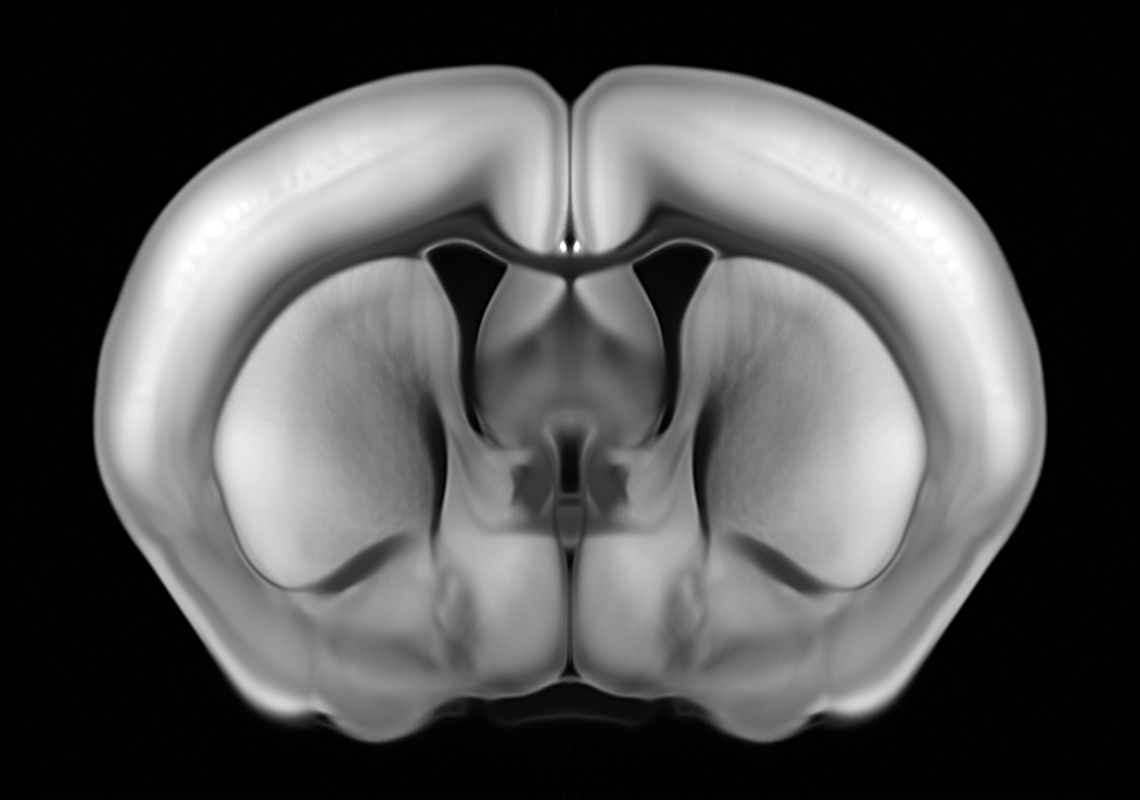

### AllenCCF_Z047.tif

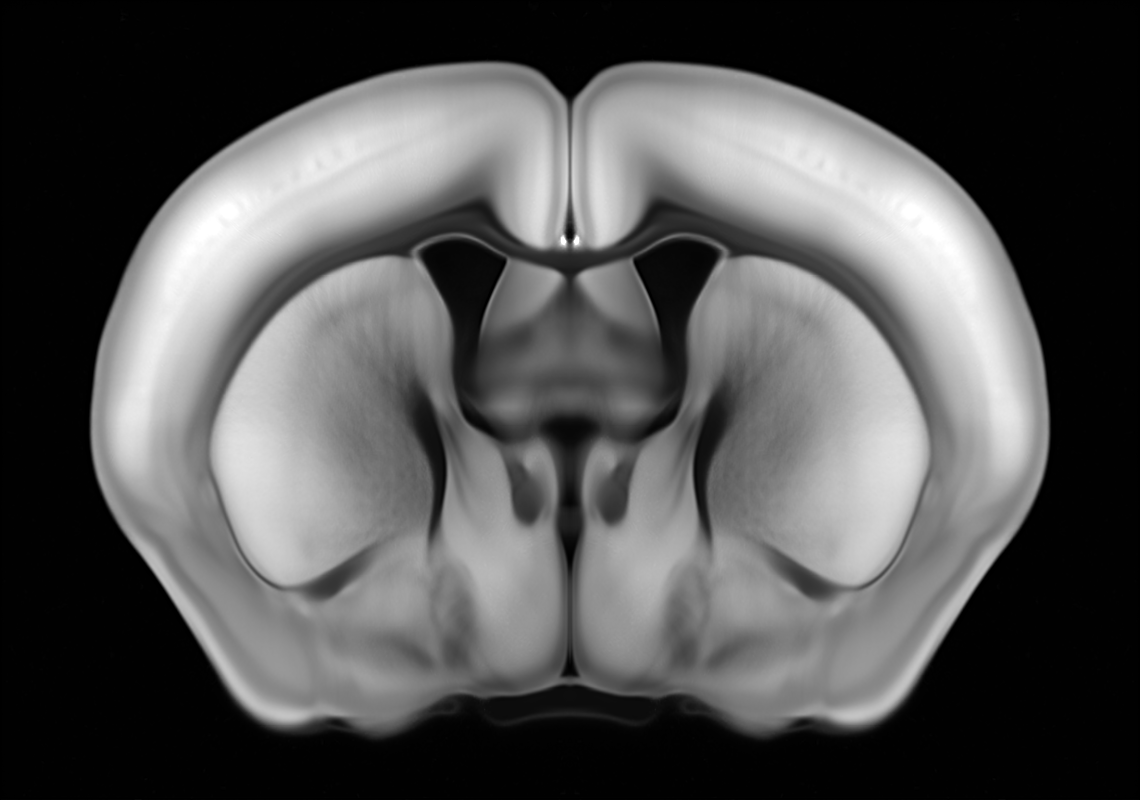

### AllenCCF_Z048.tif

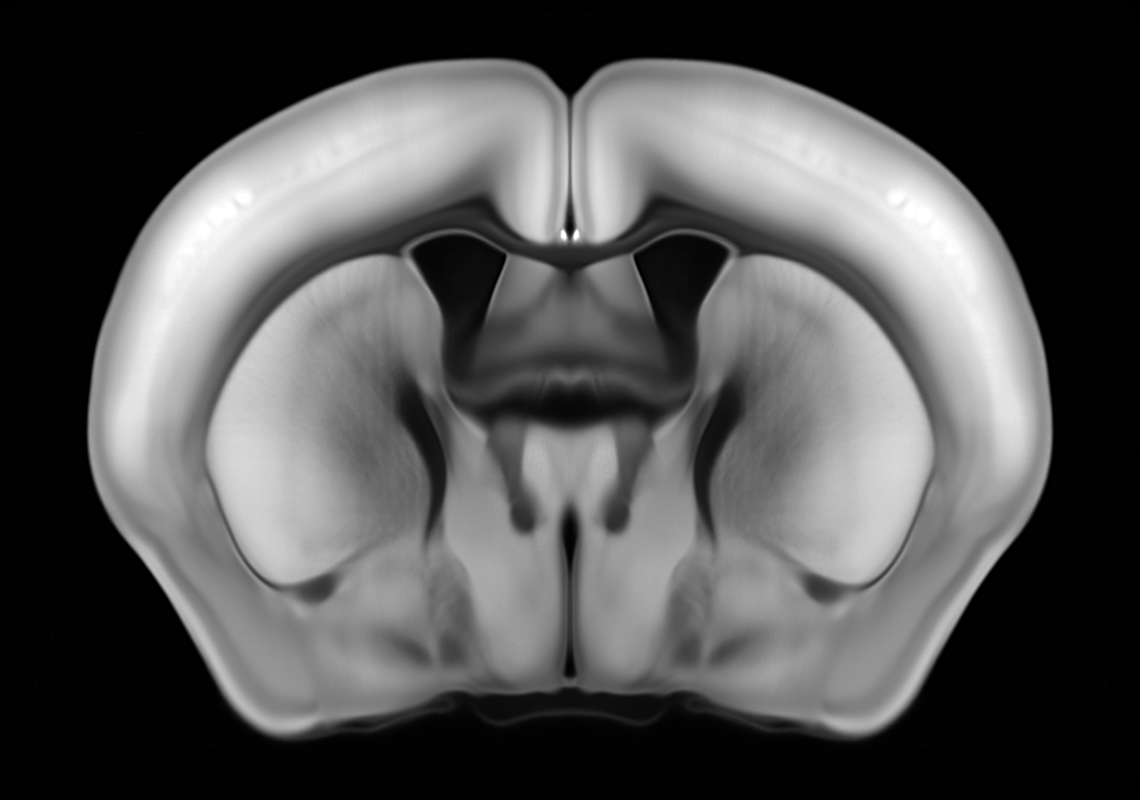

### AllenCCF_Z049.tif

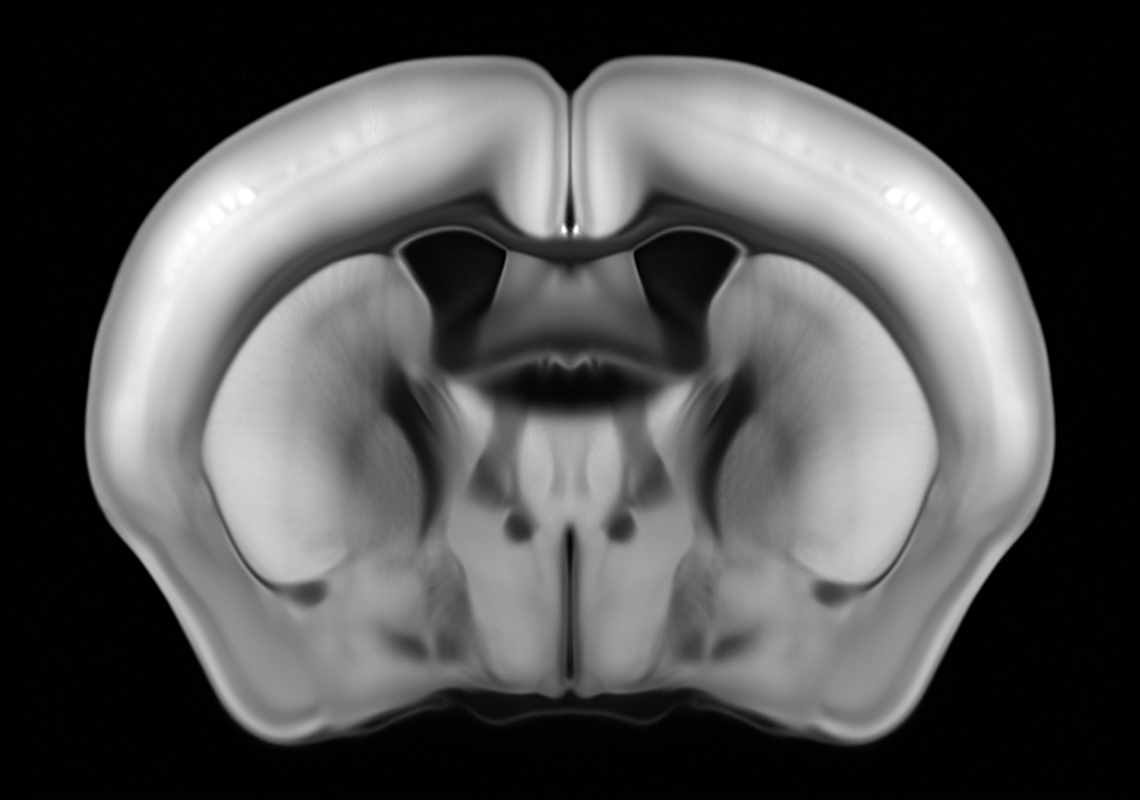

### AllenCCF_Z050.tif

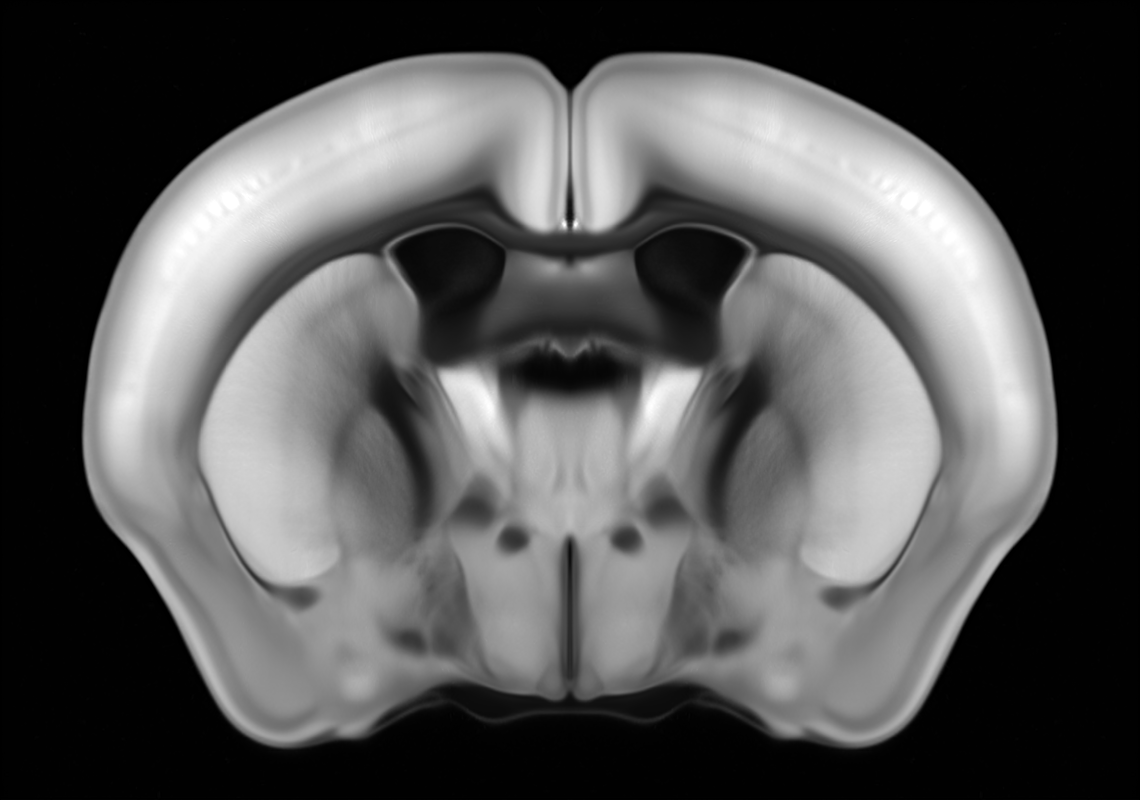

### AllenCCF_Z051.tif

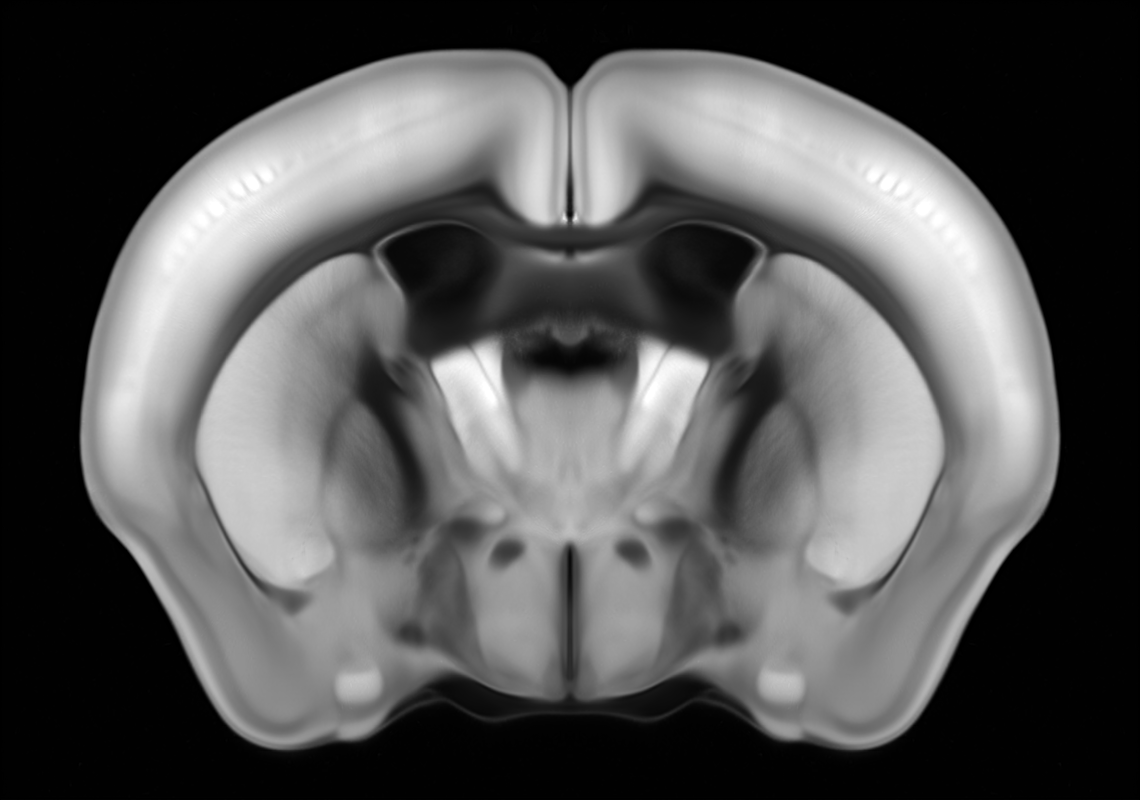

### AllenCCF_Z052.tif

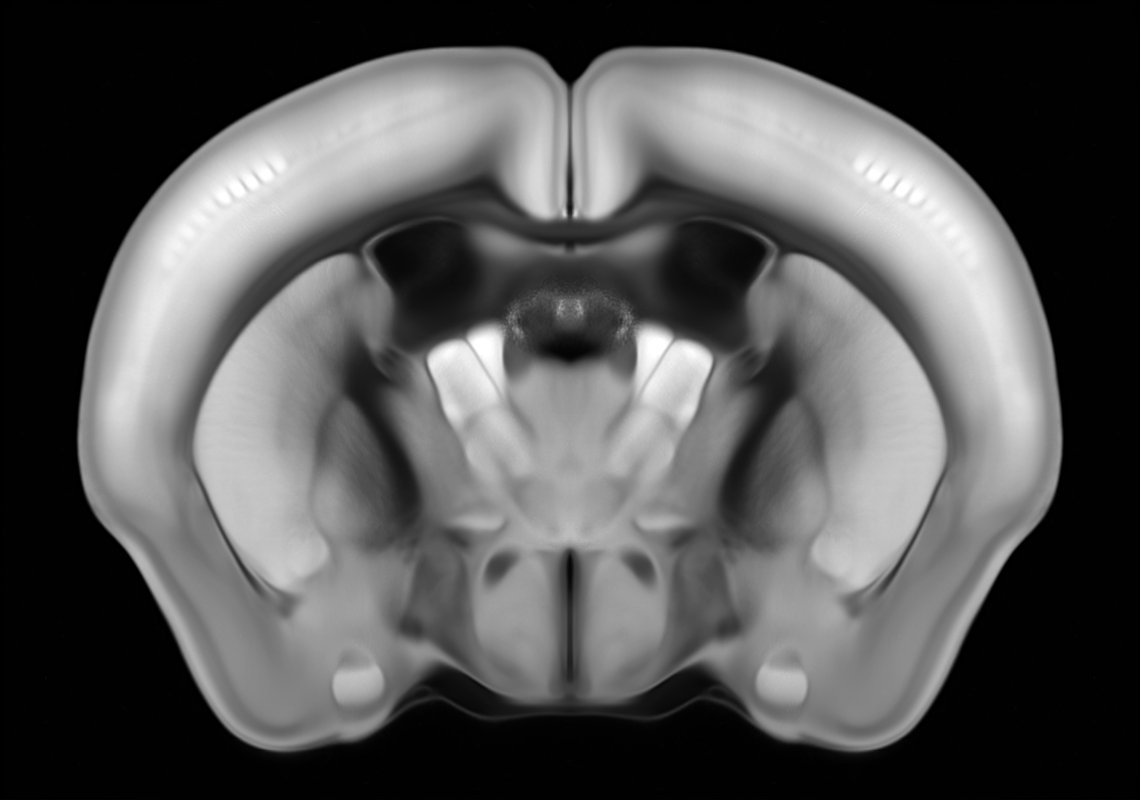

### AllenCCF_Z053.tif

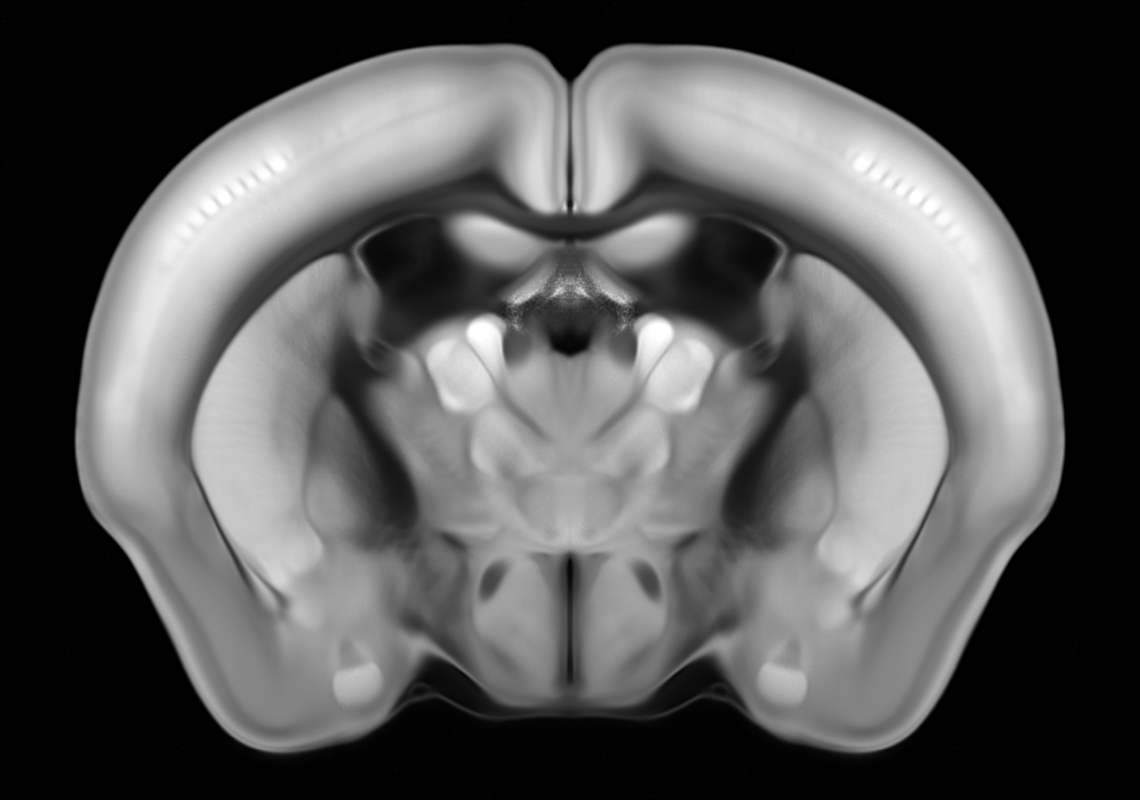

### AllenCCF_Z054.tif

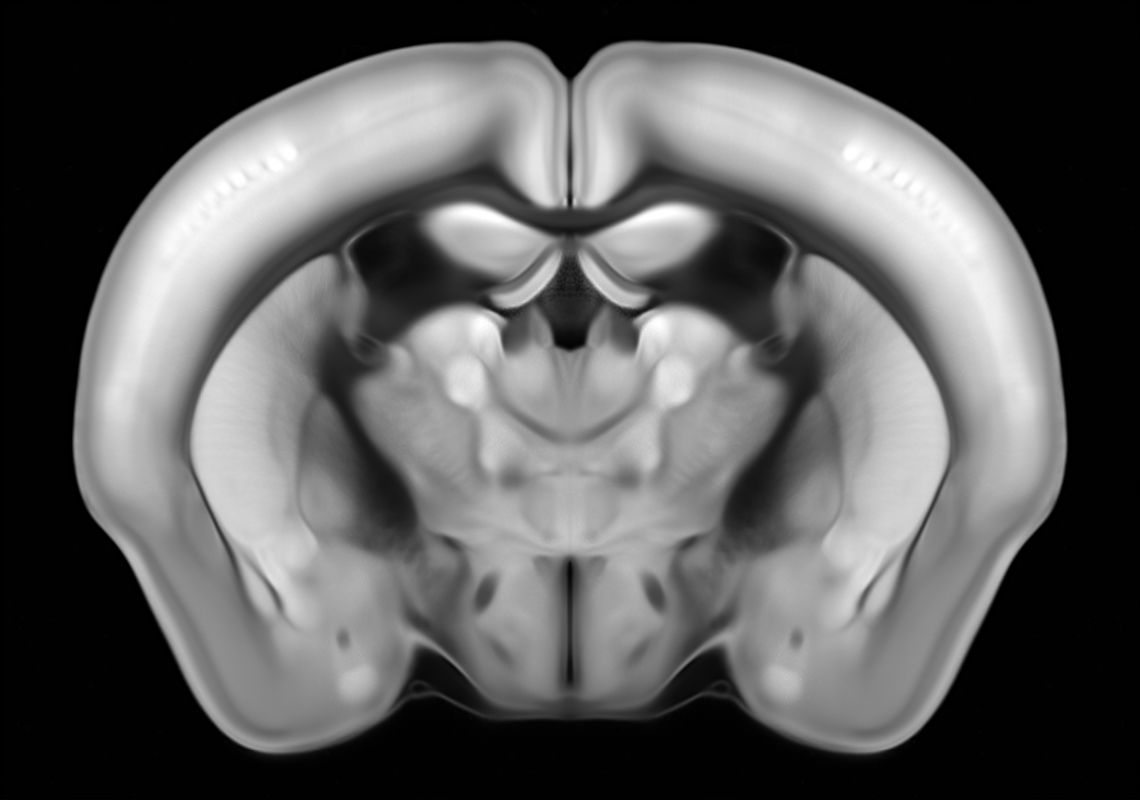

### AllenCCF_Z055.tif

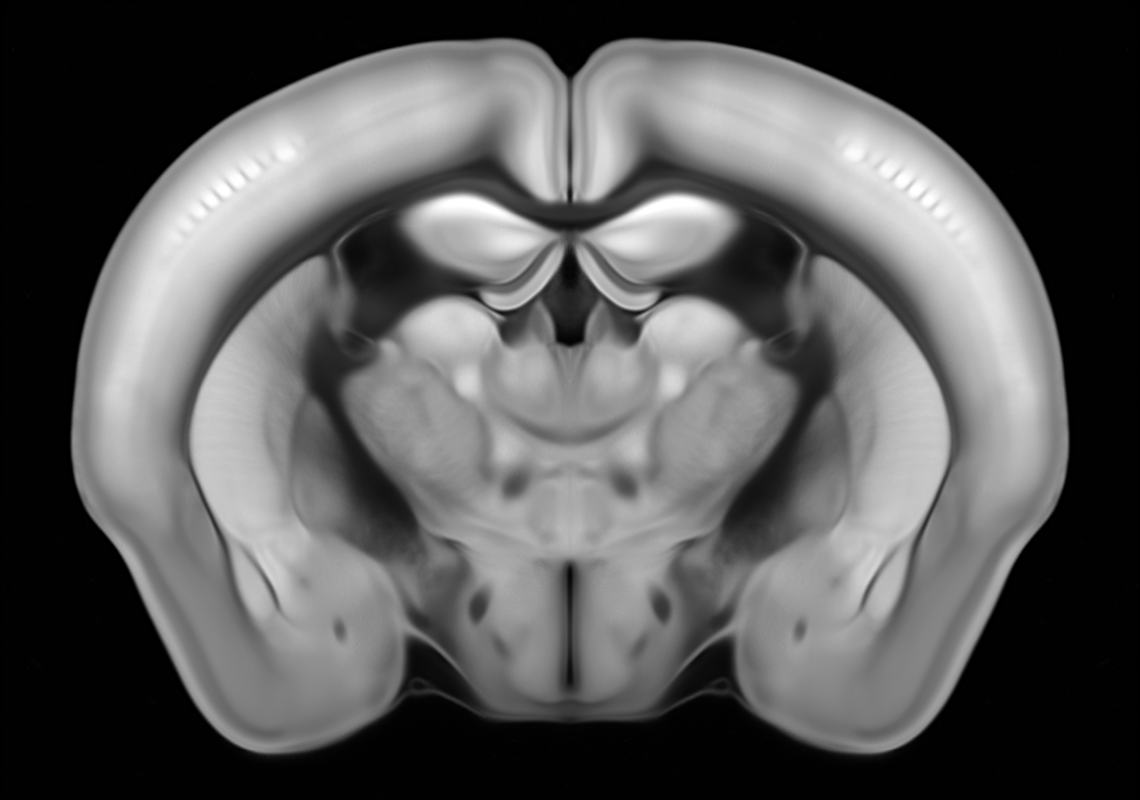

### AllenCCF_Z056.tif

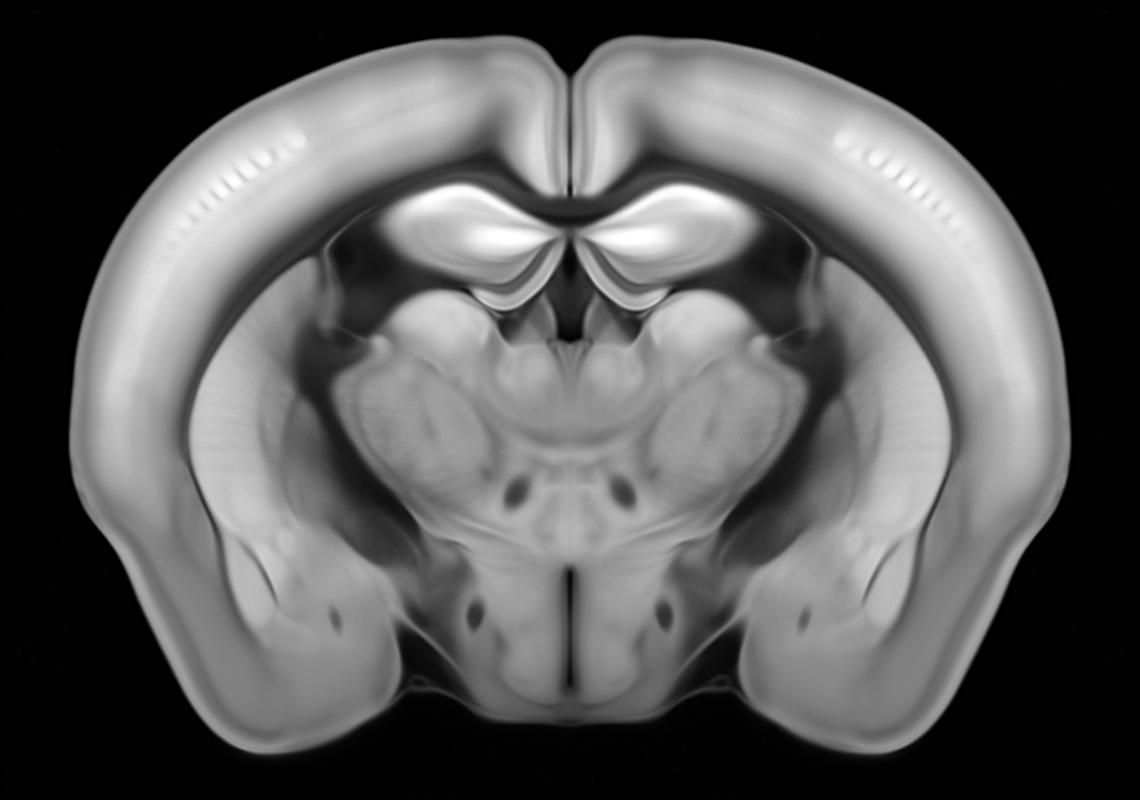

### AllenCCF_Z057.tif

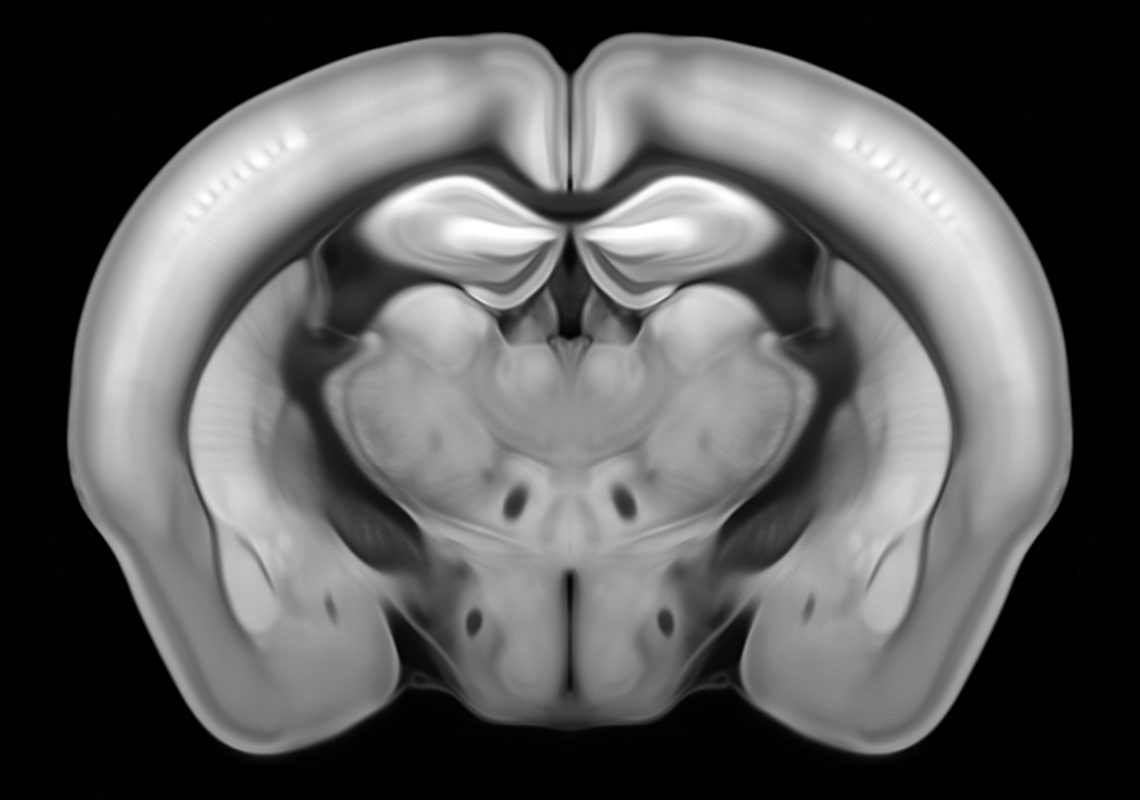

### AllenCCF_Z058.tif

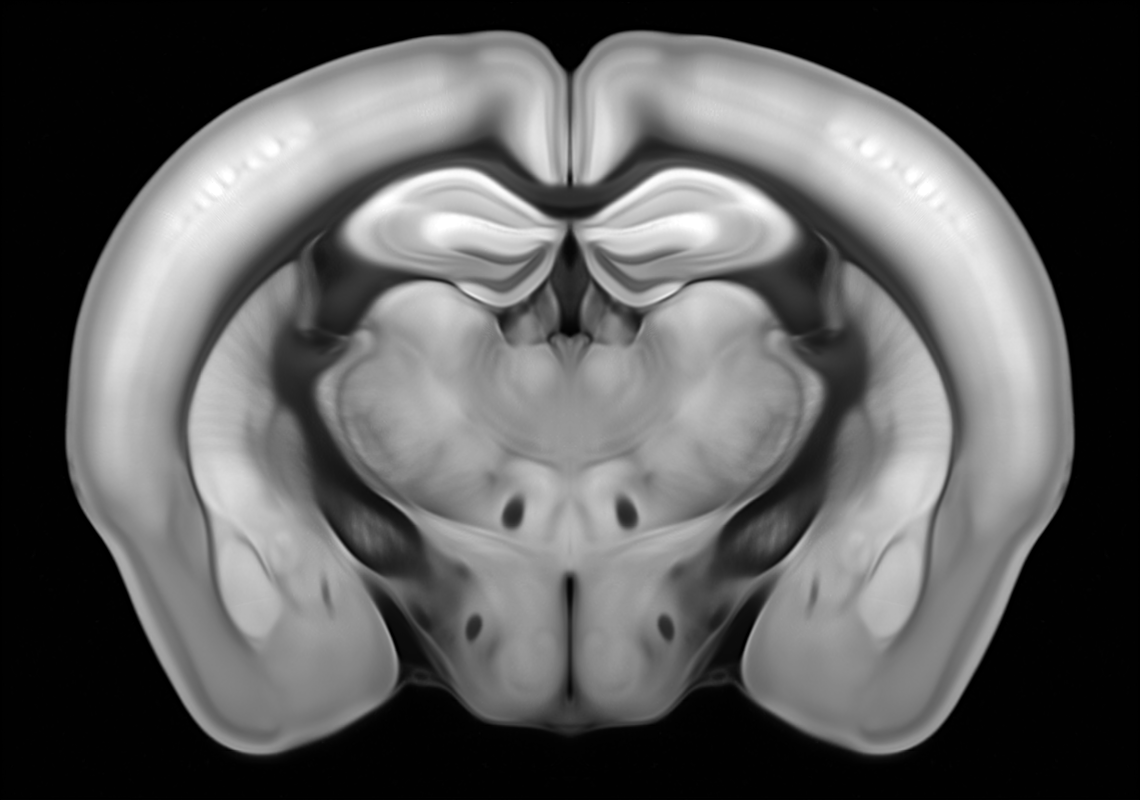

### AllenCCF_Z059.tif

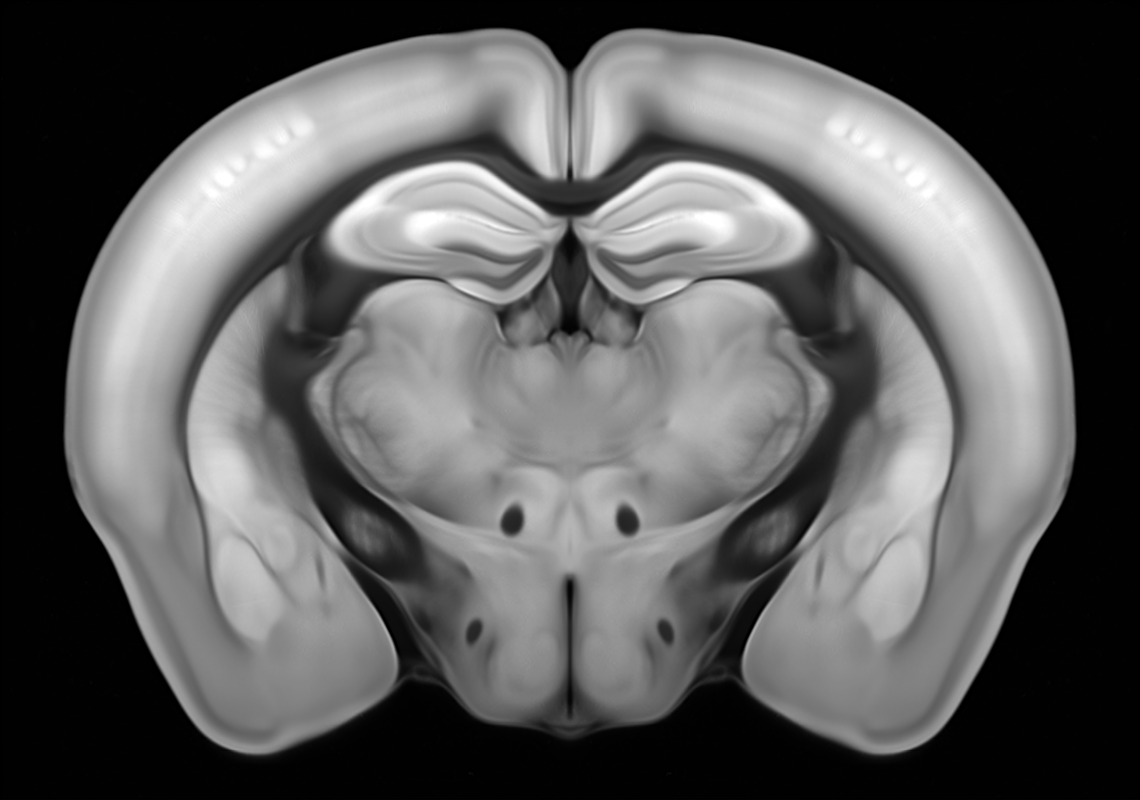

### AllenCCF_Z060.tif

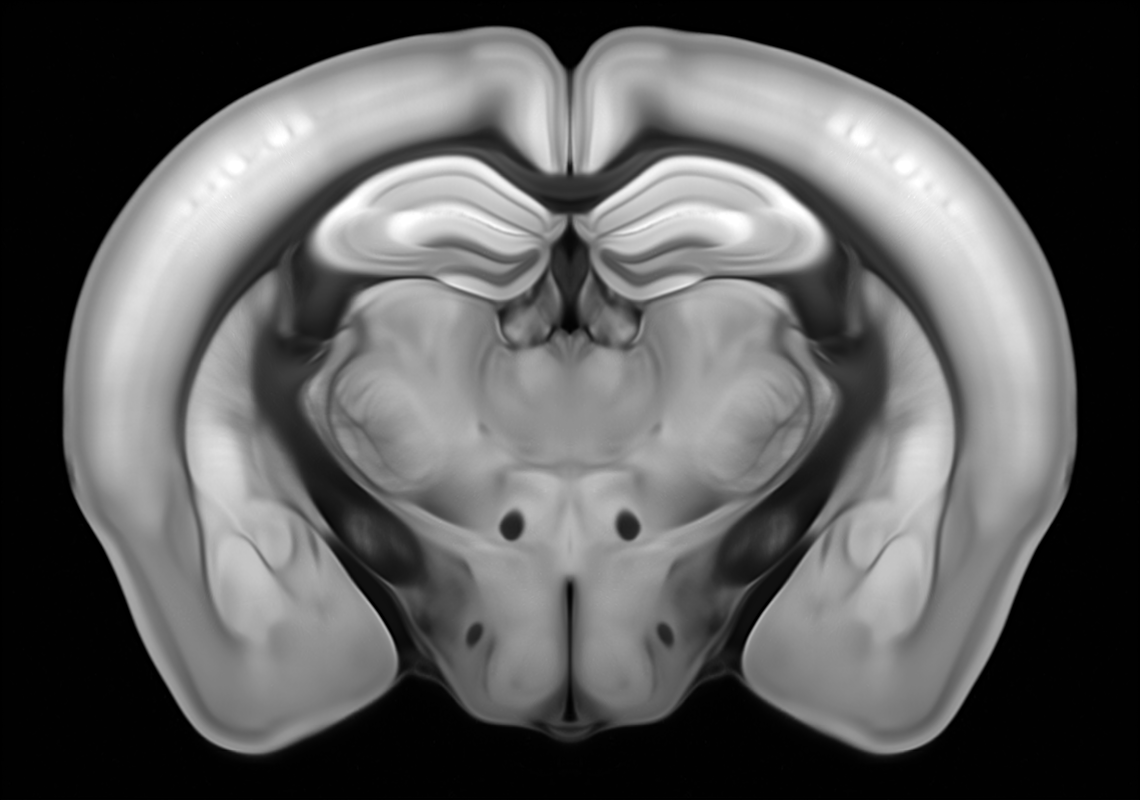

### AllenCCF_Z061.tif

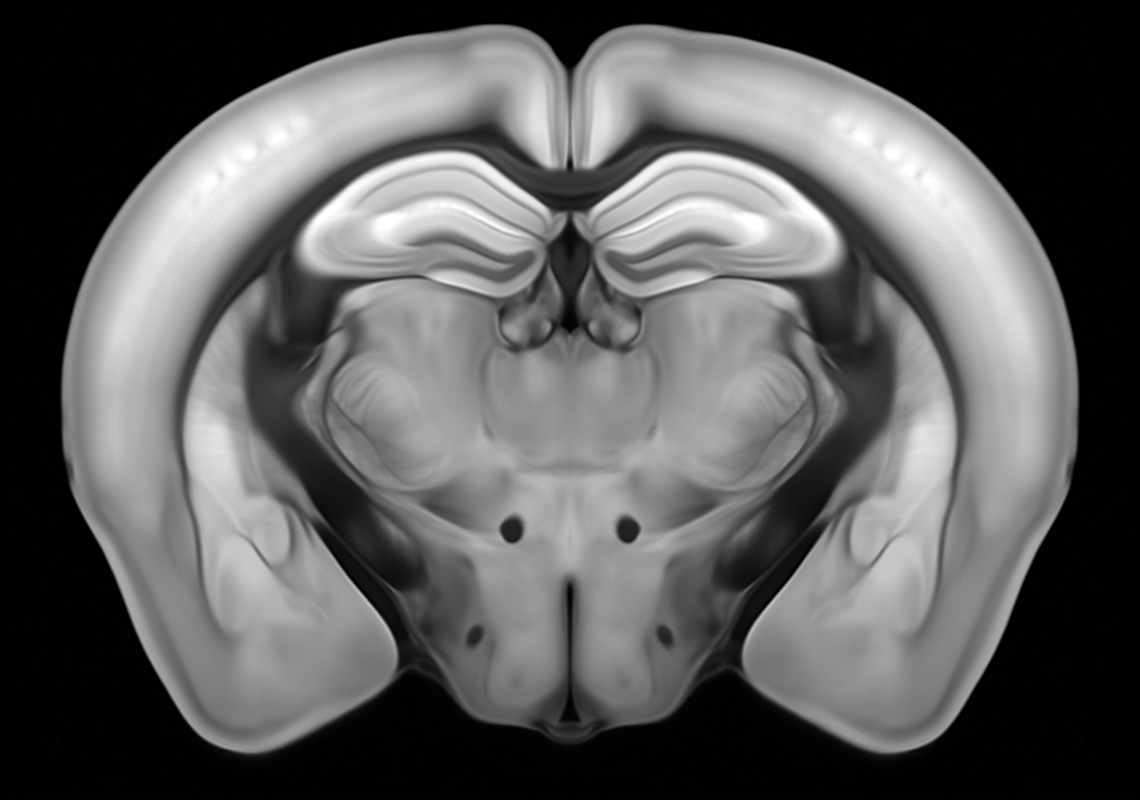

### AllenCCF_Z062.tif

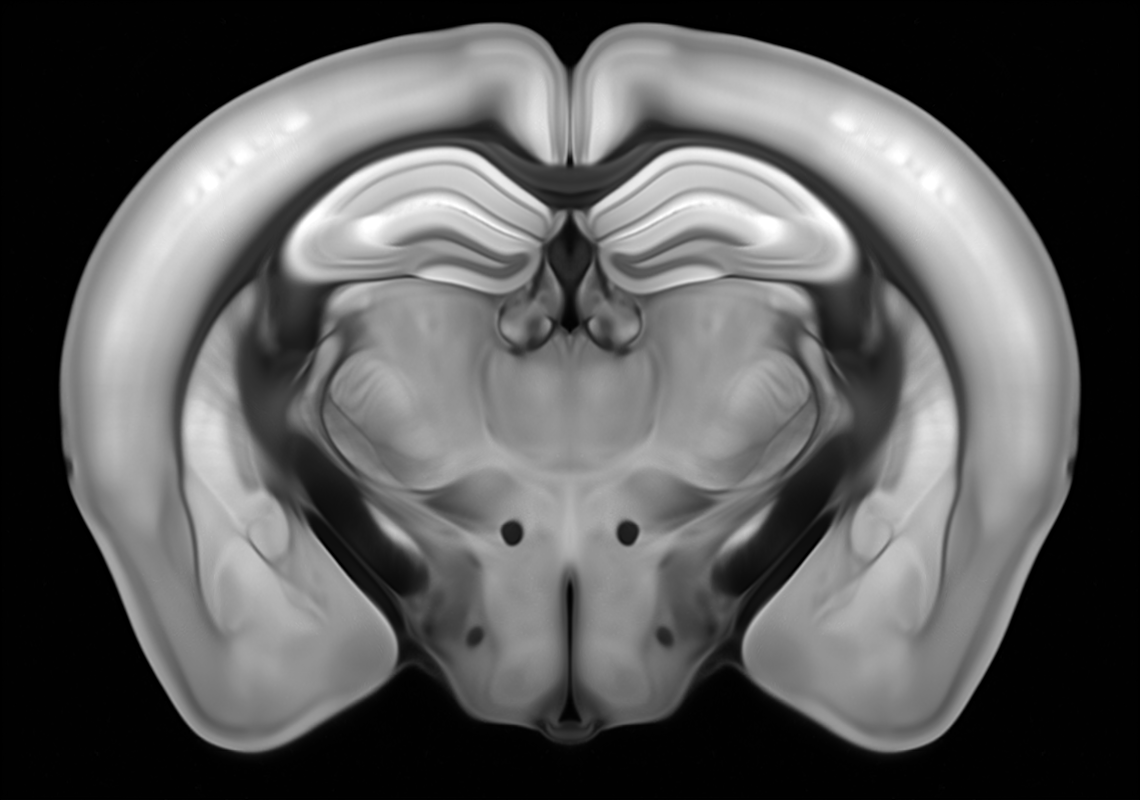

### AllenCCF_Z063.tif

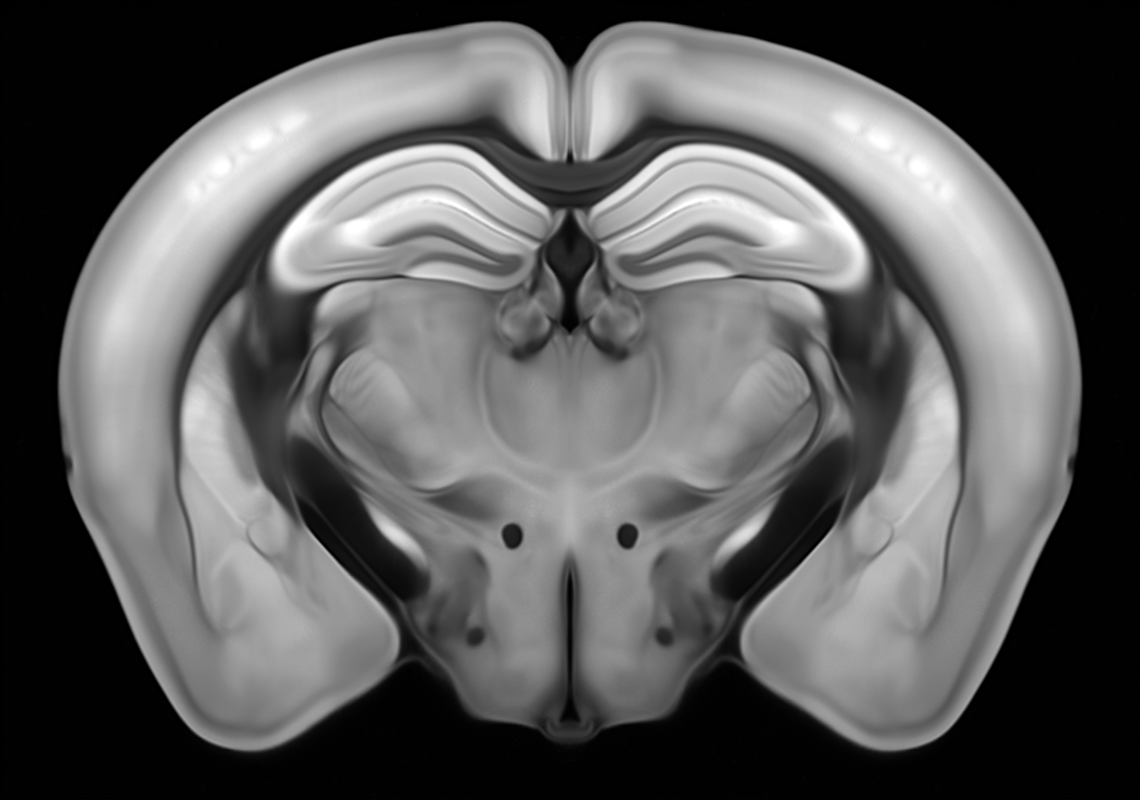

### AllenCCF_Z064.tif

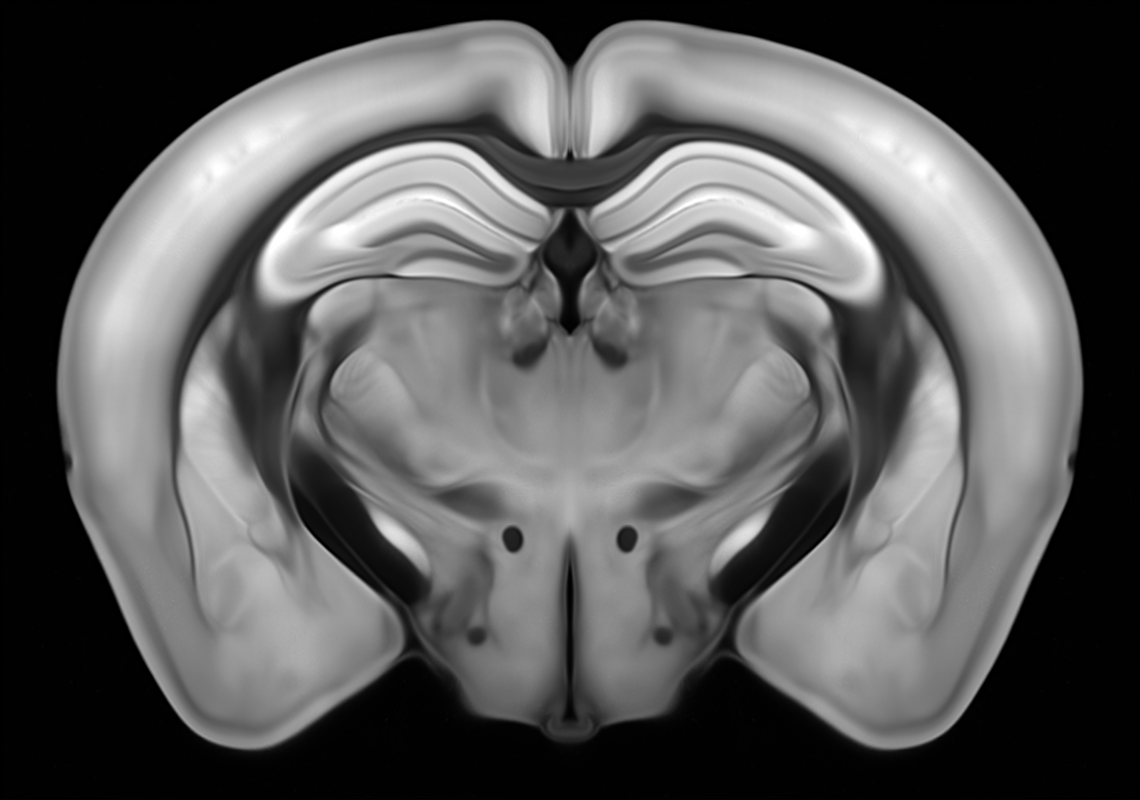

### AllenCCF_Z065.tif

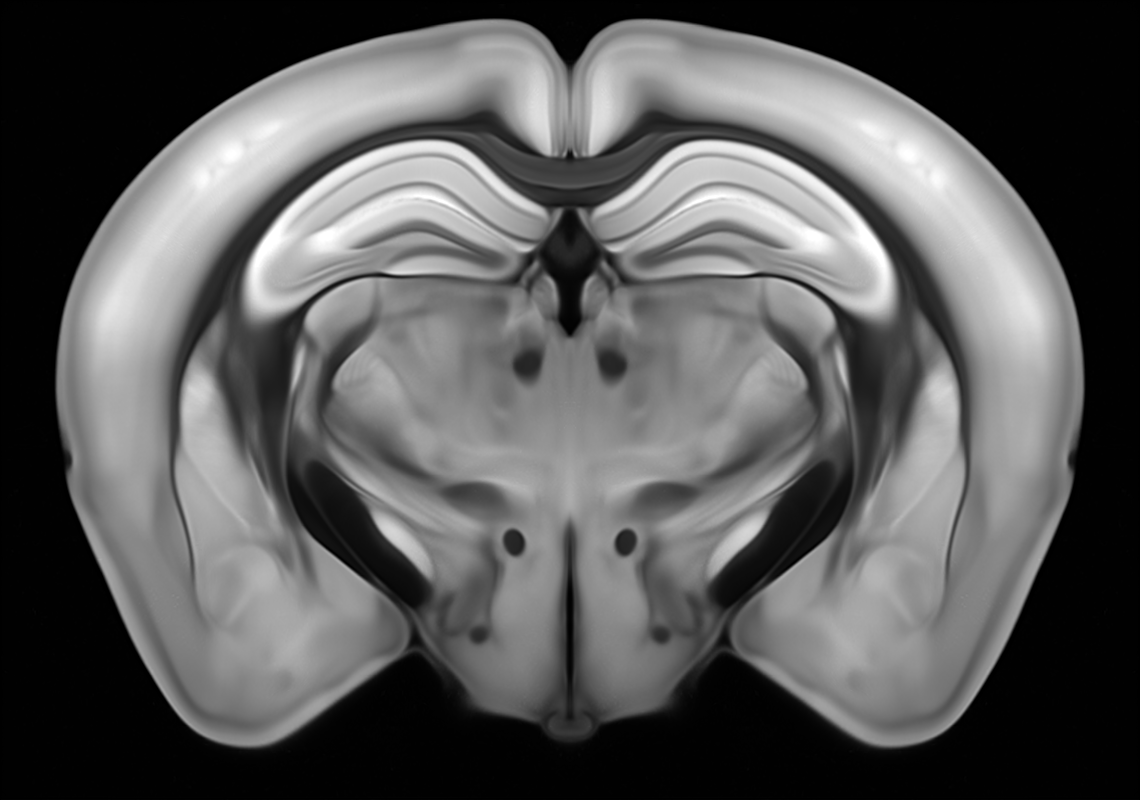

### AllenCCF_Z066.tif

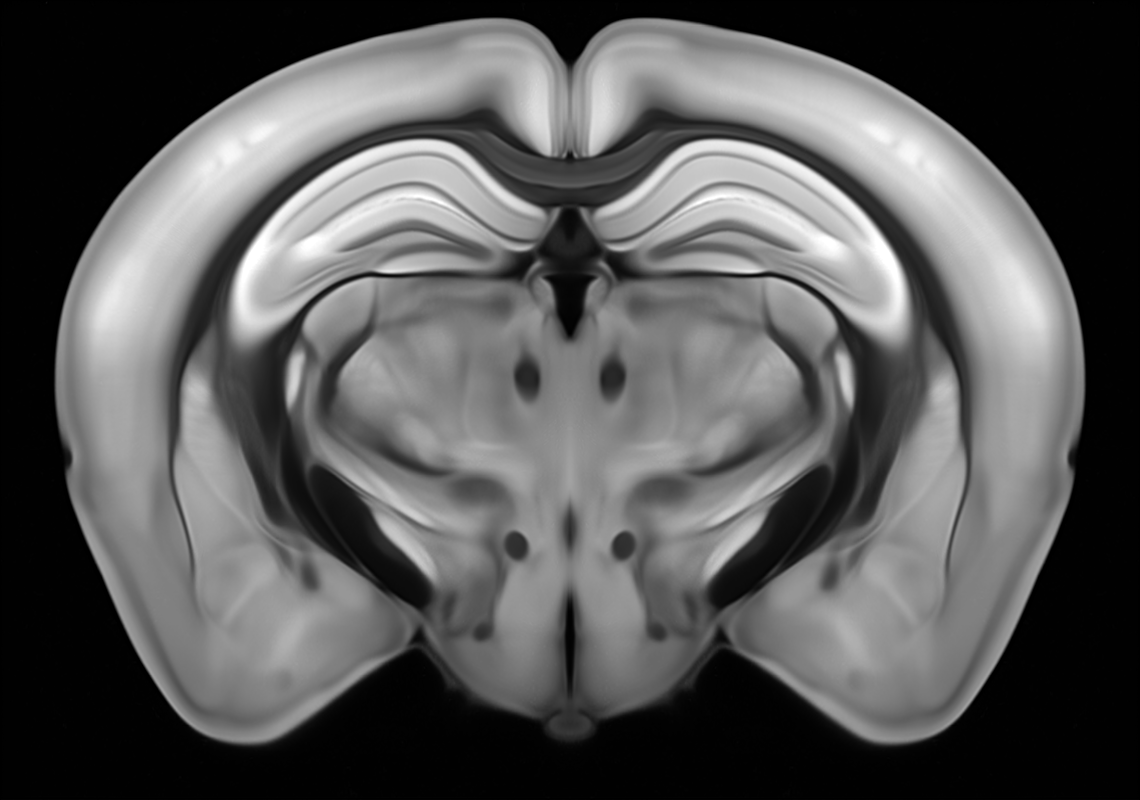

### AllenCCF_Z067.tif

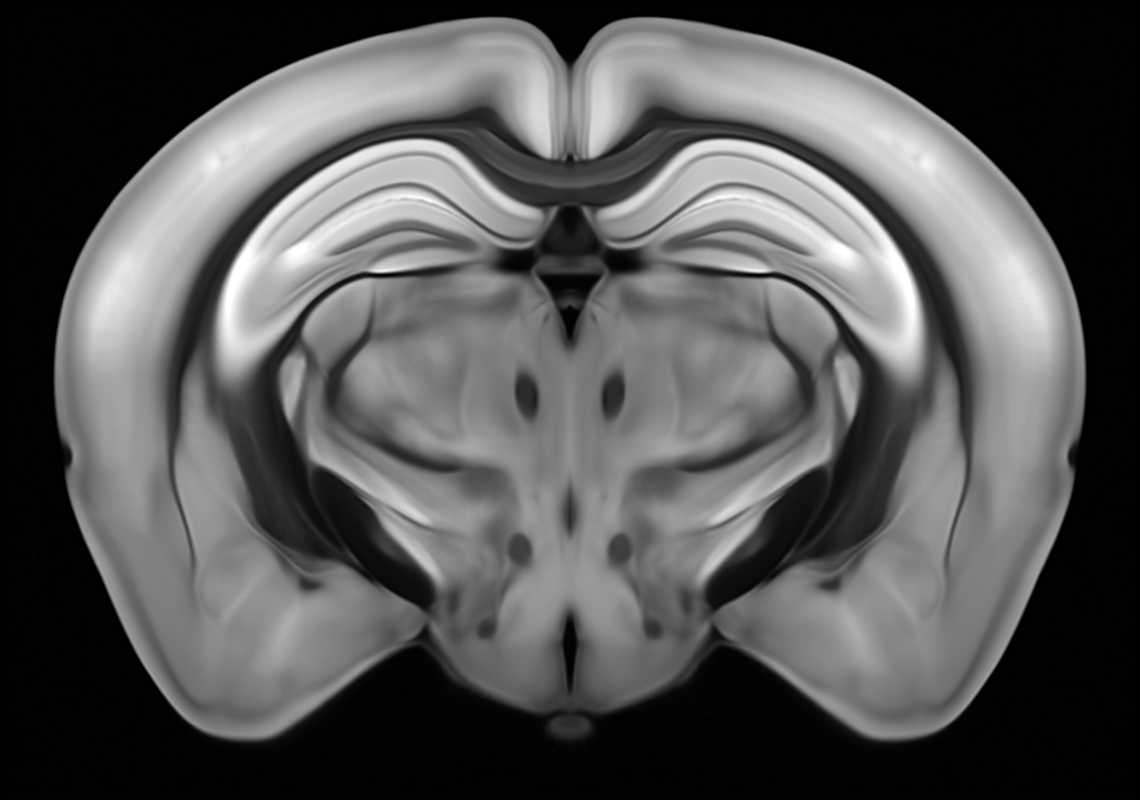

### AllenCCF_Z068.tif

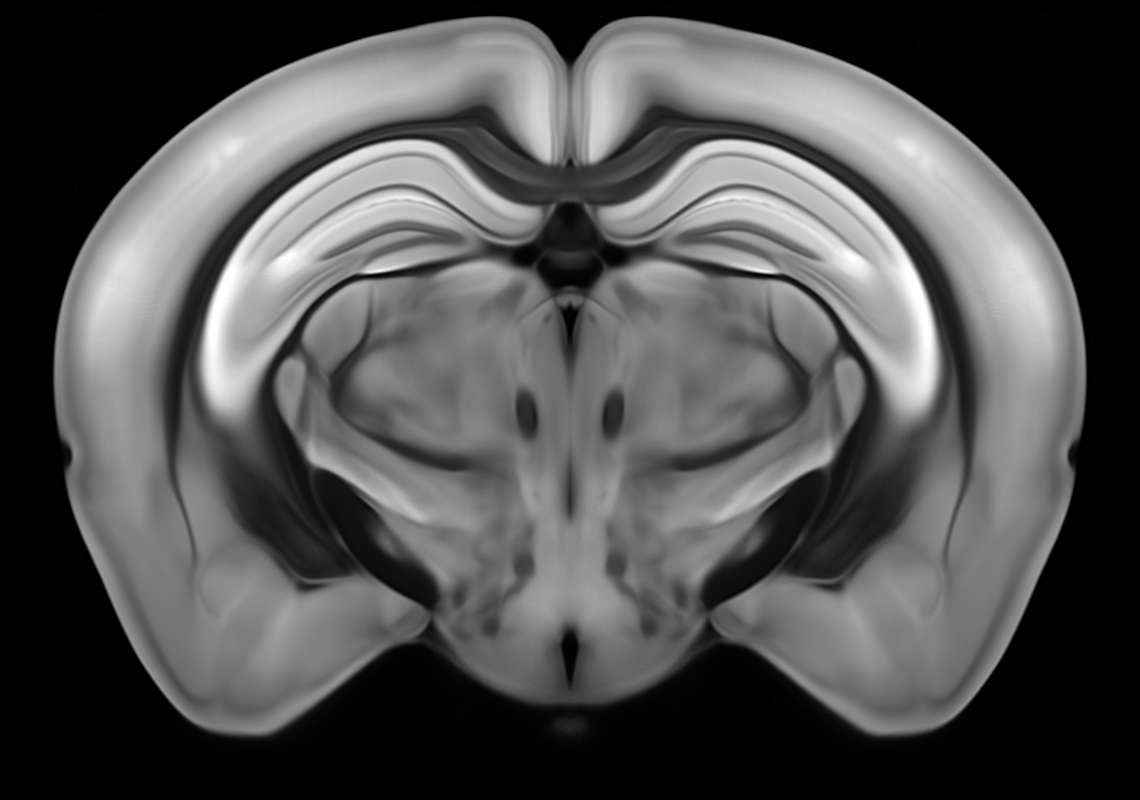

### AllenCCF_Z069.tif

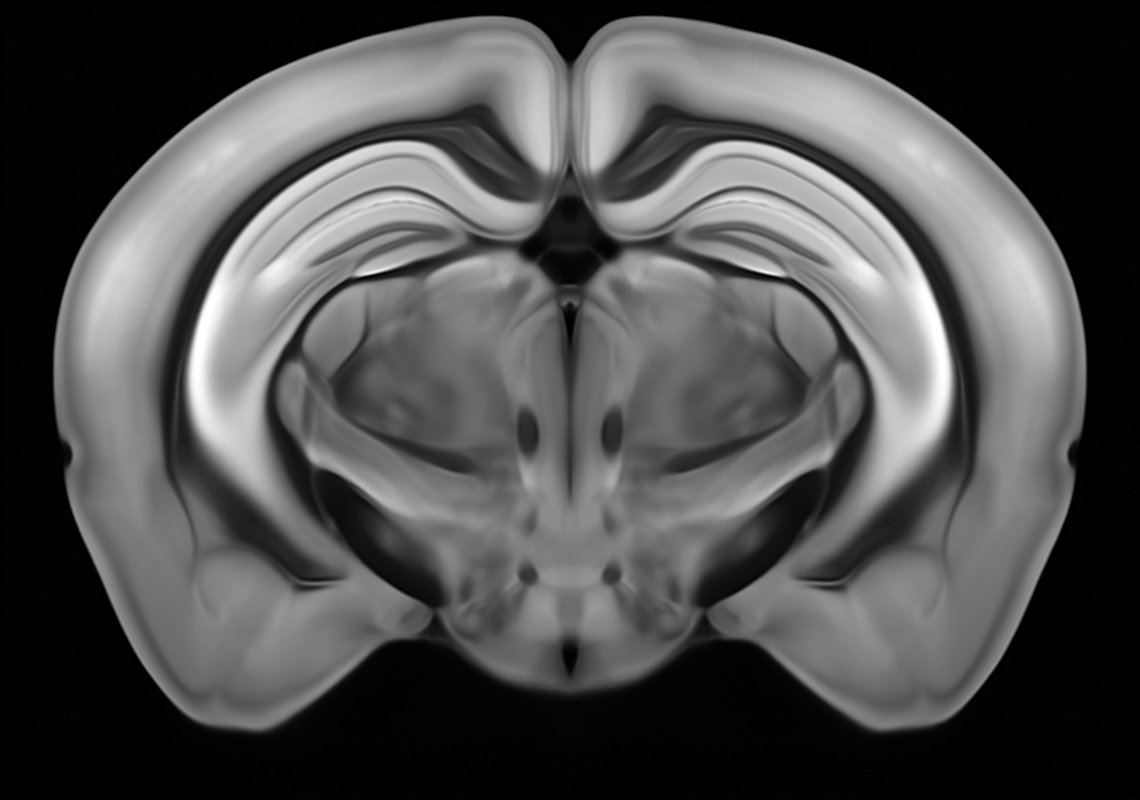

### AllenCCF_Z070.tif

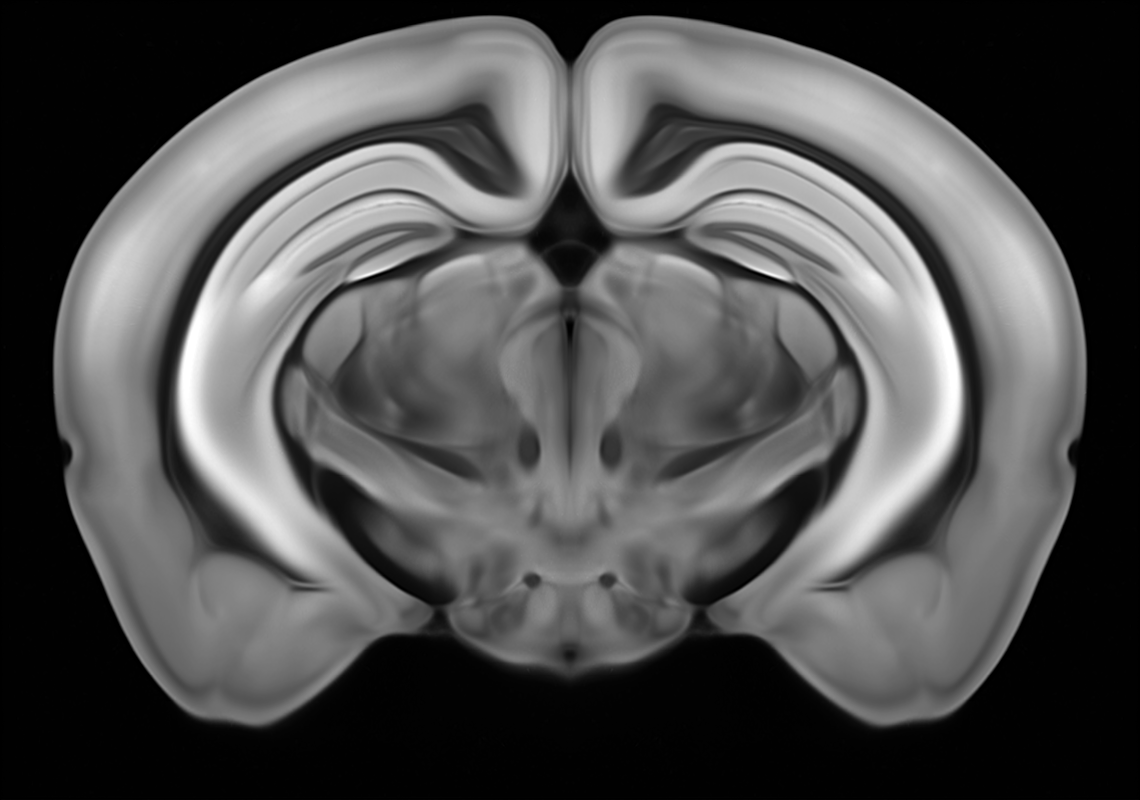

### AllenCCF_Z071.tif

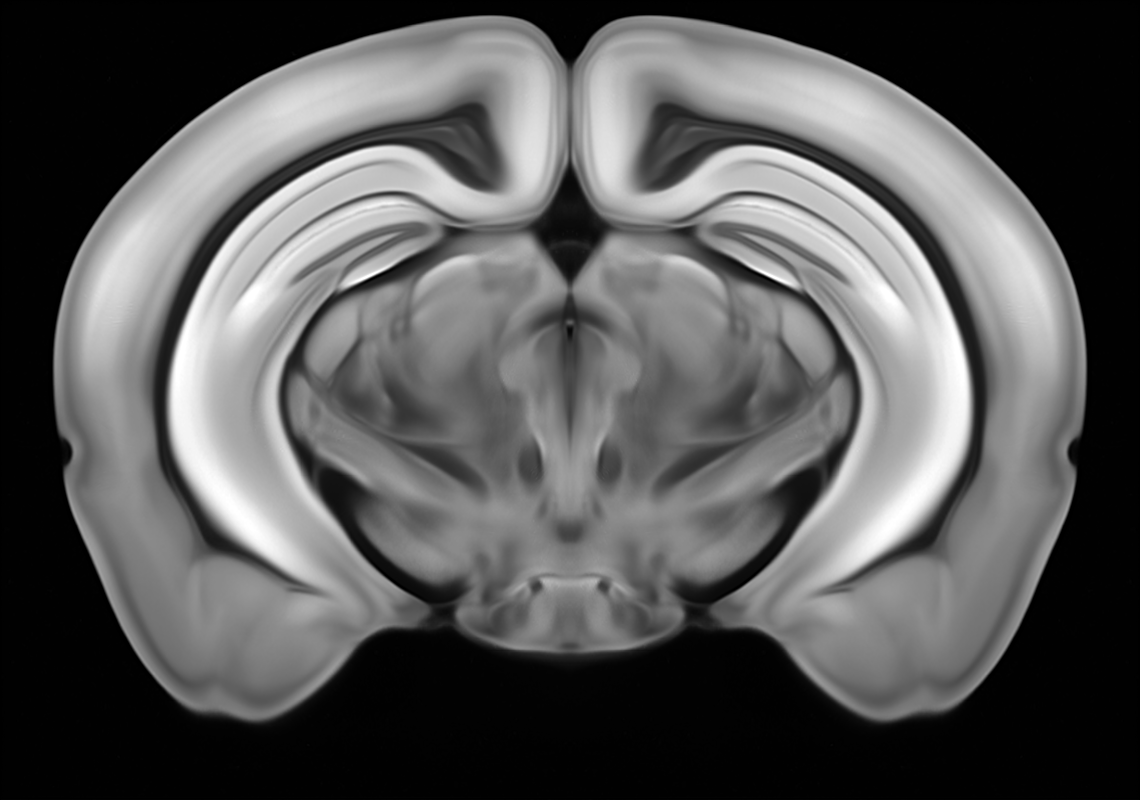

### AllenCCF_Z072.tif

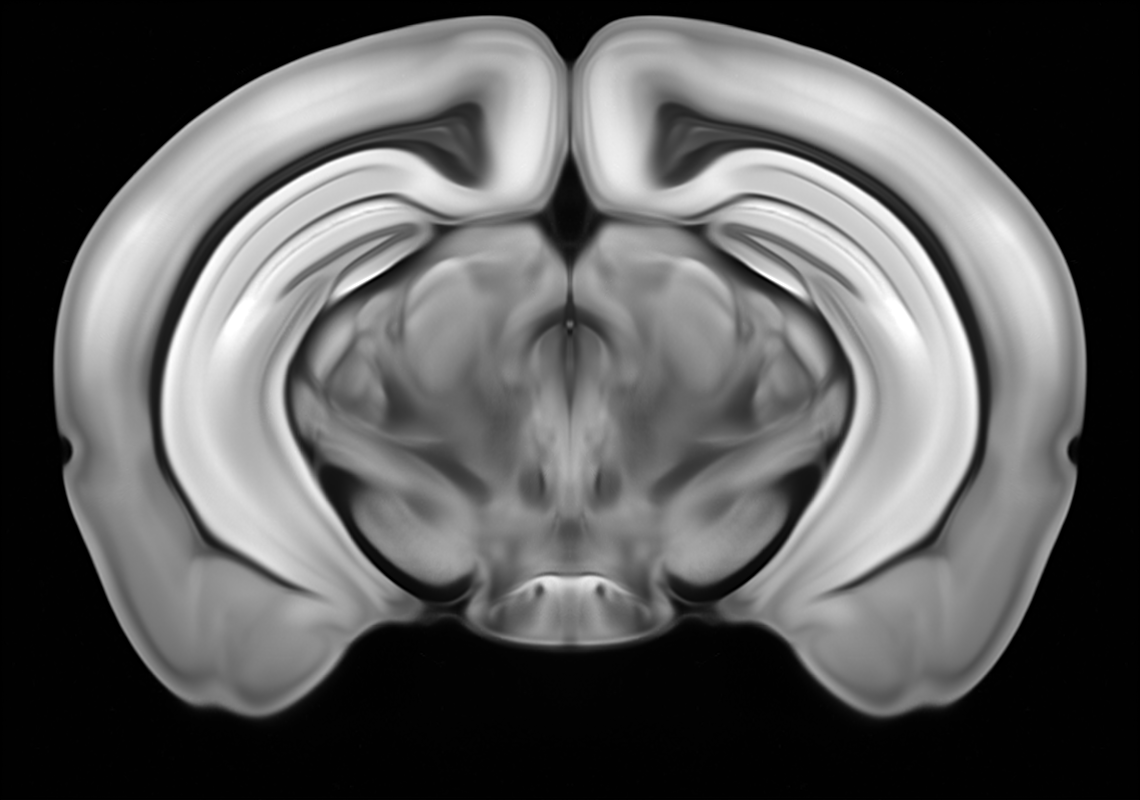

### AllenCCF_Z073.tif

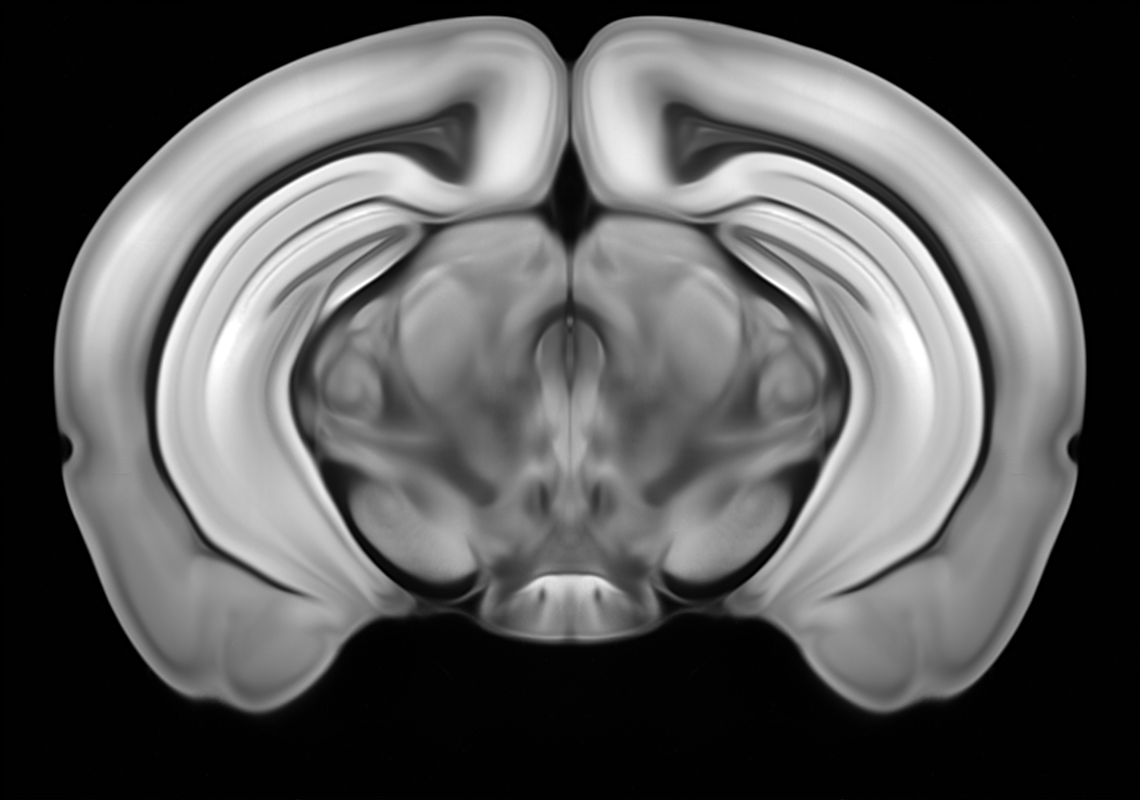

### AllenCCF_Z074.tif

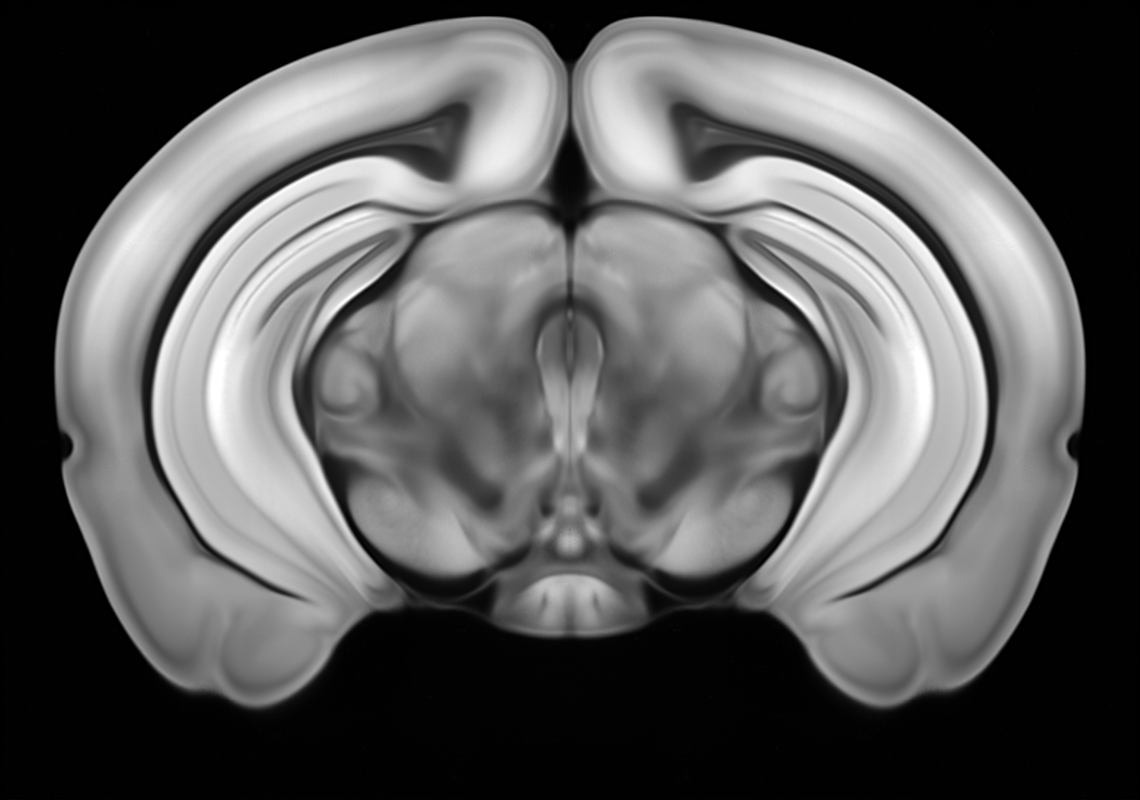

### AllenCCF_Z075.tif

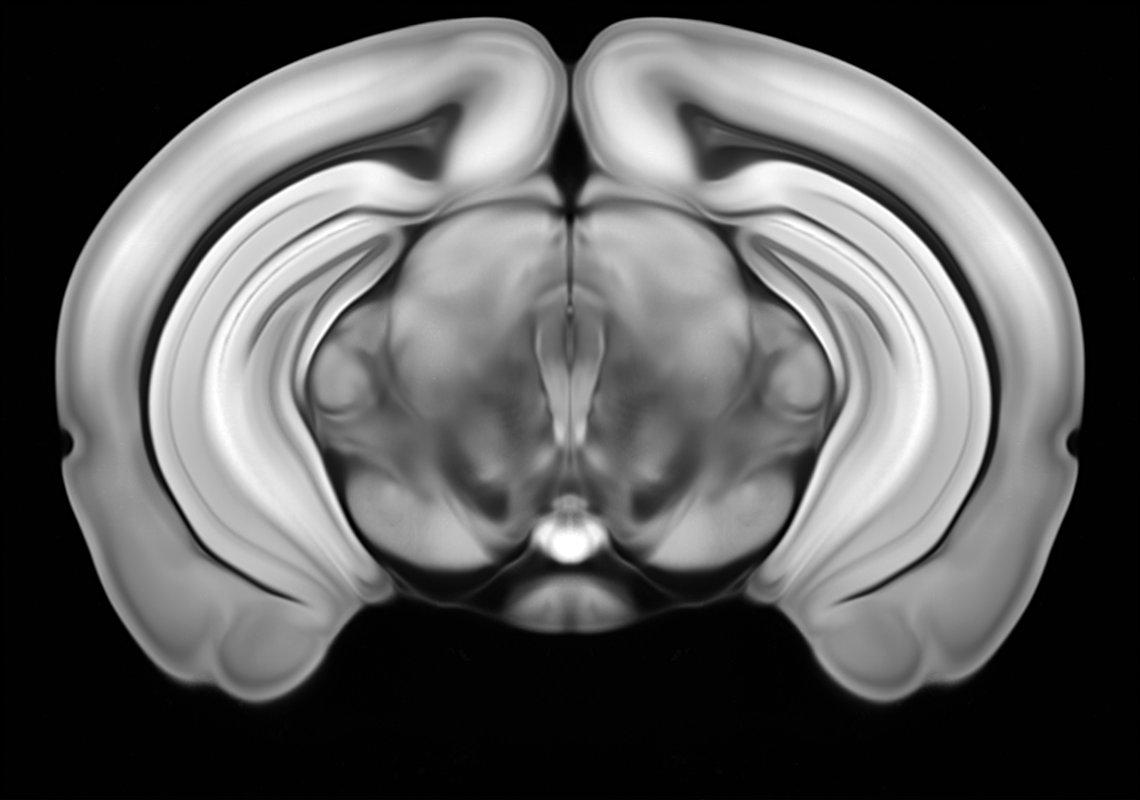

### AllenCCF_Z076.tif

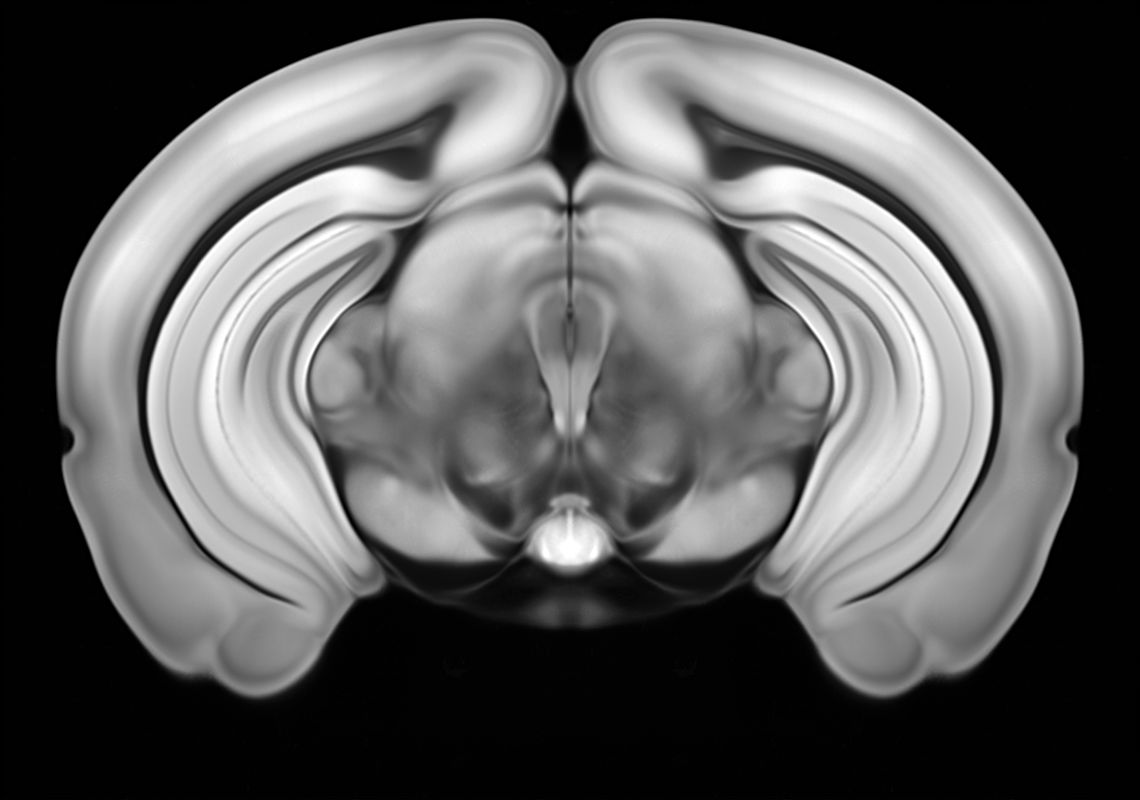

### AllenCCF_Z077.tif

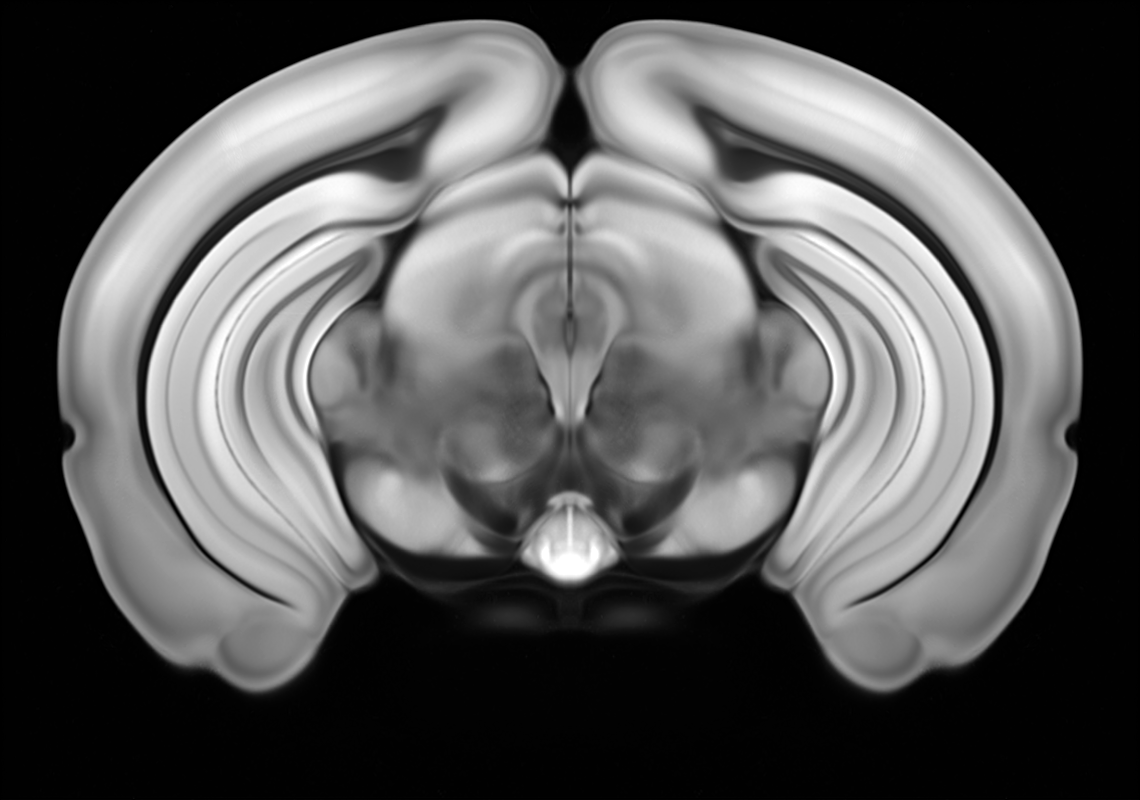
