## Supplementary figures and images for "Enhanced and Unified Anatomical Labeling for a Common Mouse Brain Atlas"

### 2_AP+4.2.tif

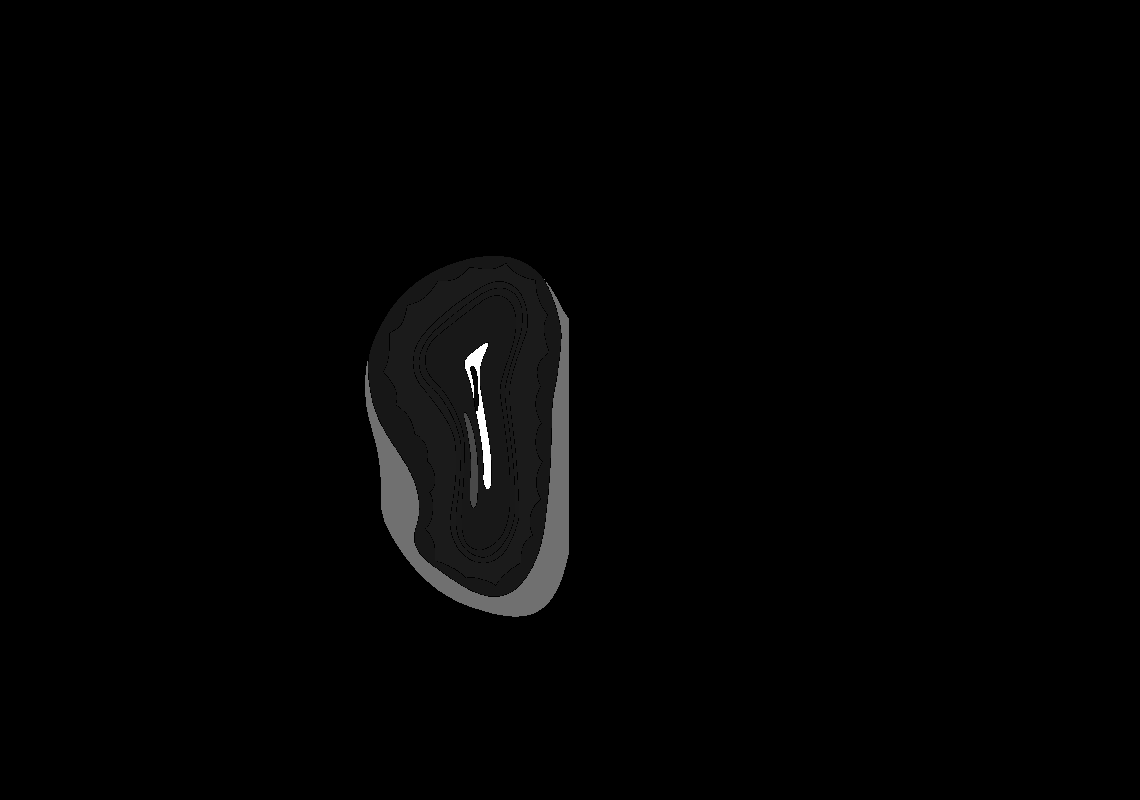

### 3_AP+4.1.tif

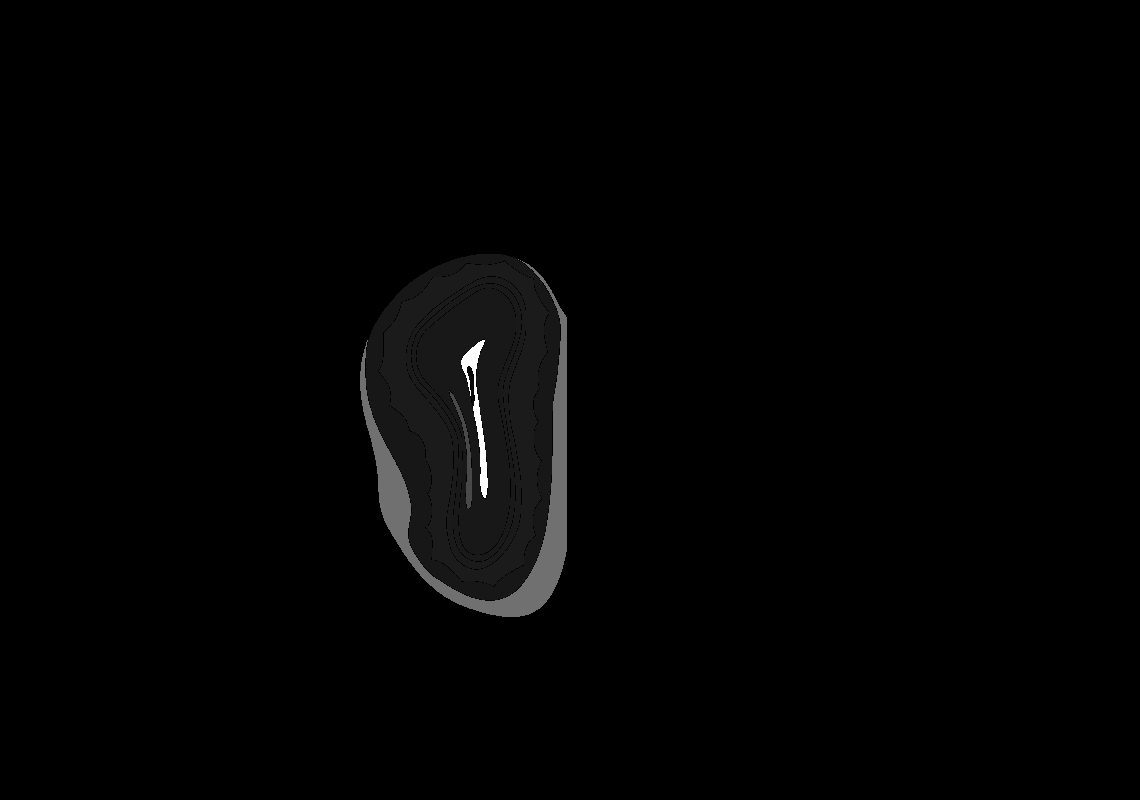

### 6_AP+3.8.tif

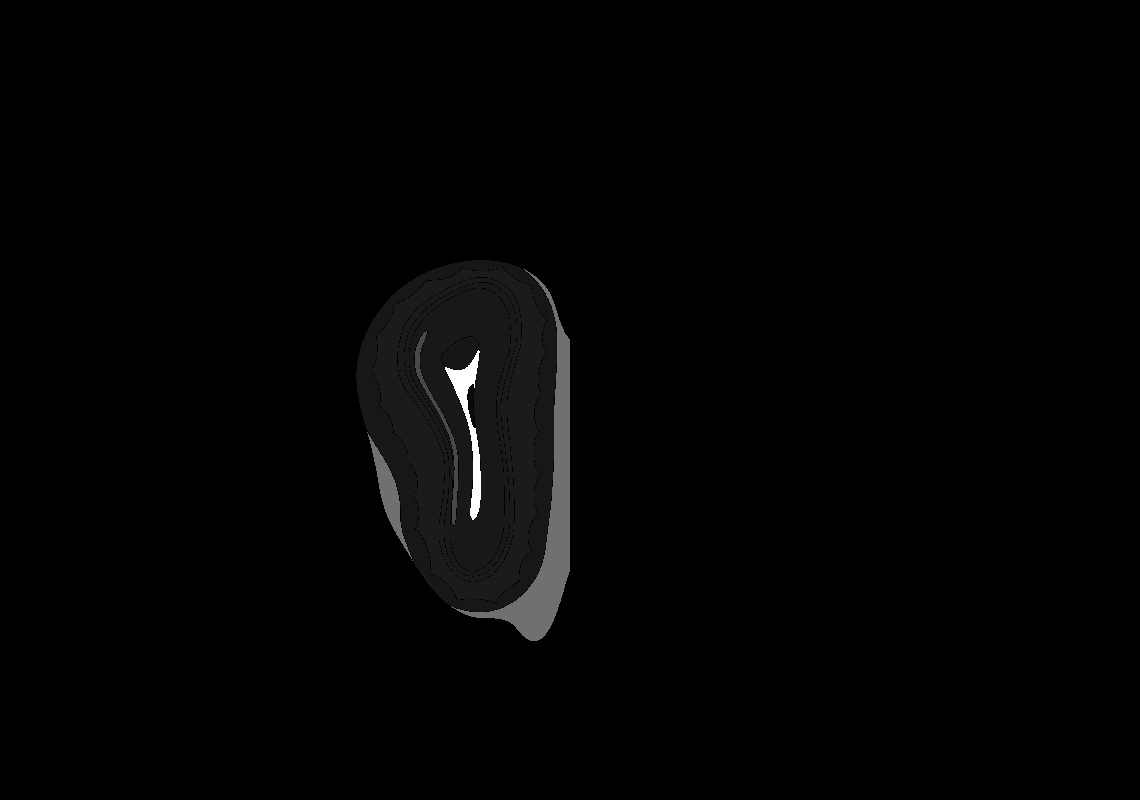

### 7_AP+3.7.tif

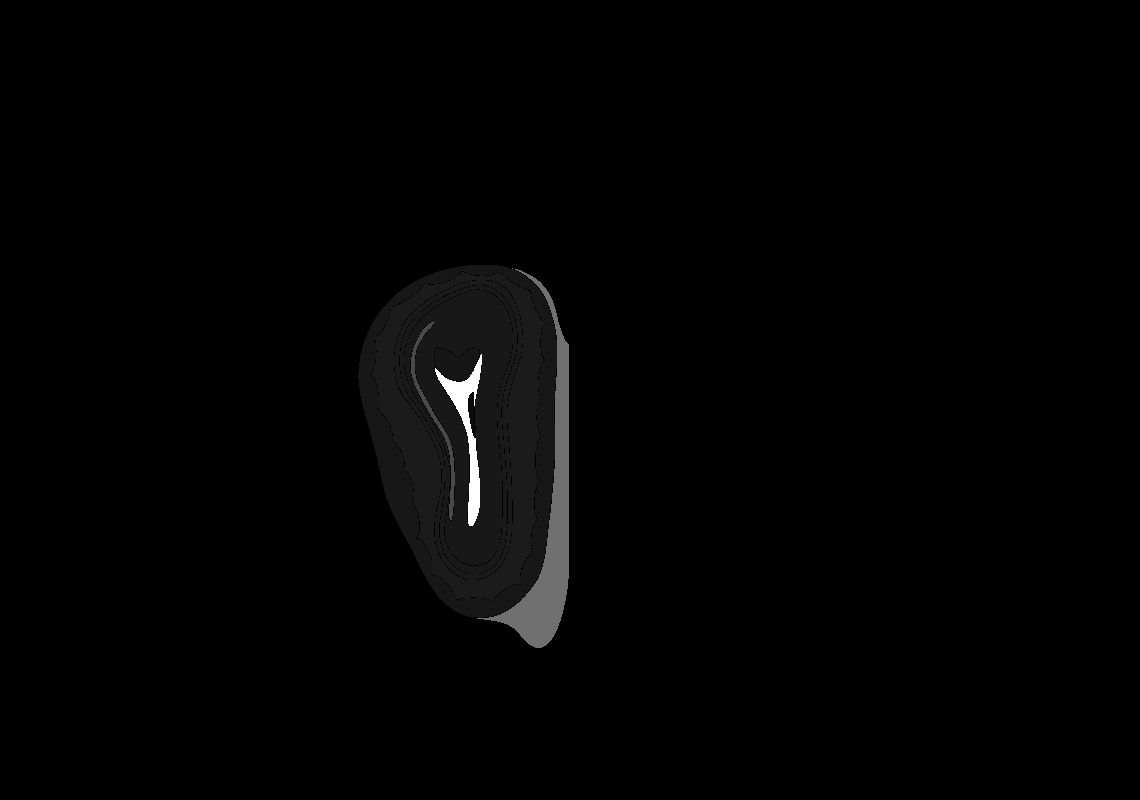

### 8_AP+3.6.tif

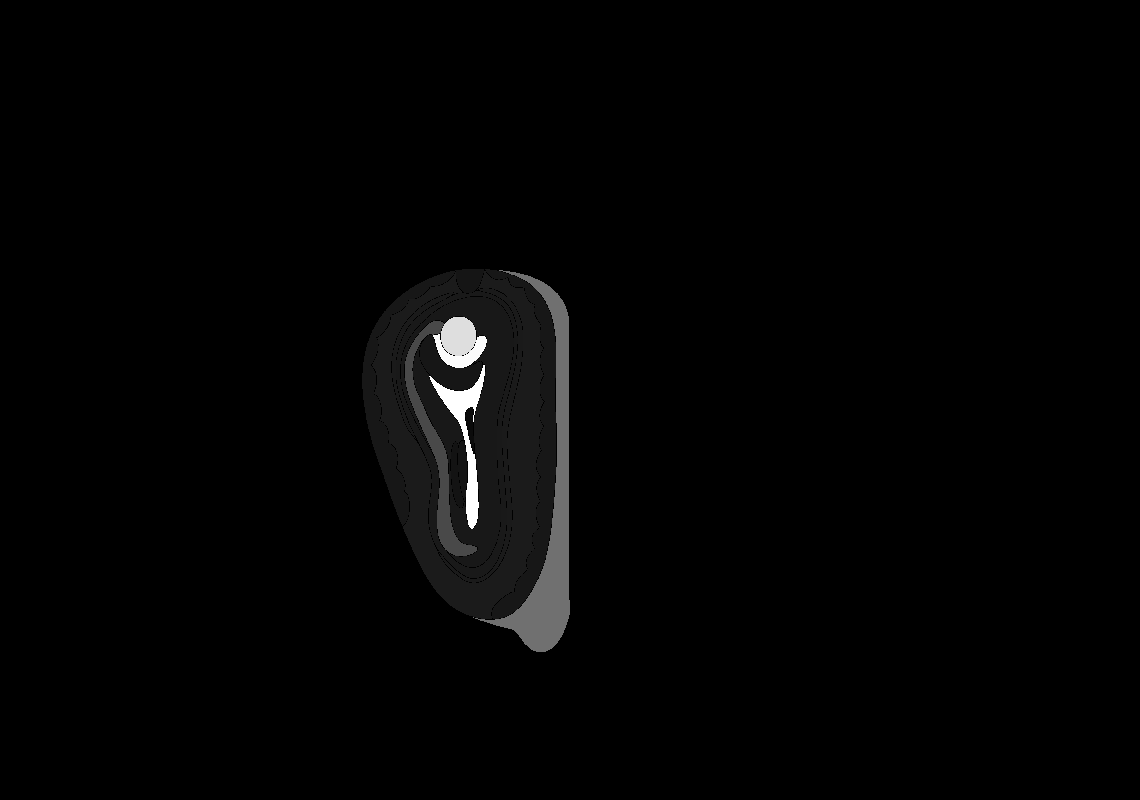

### 9_AP+3.5.tif

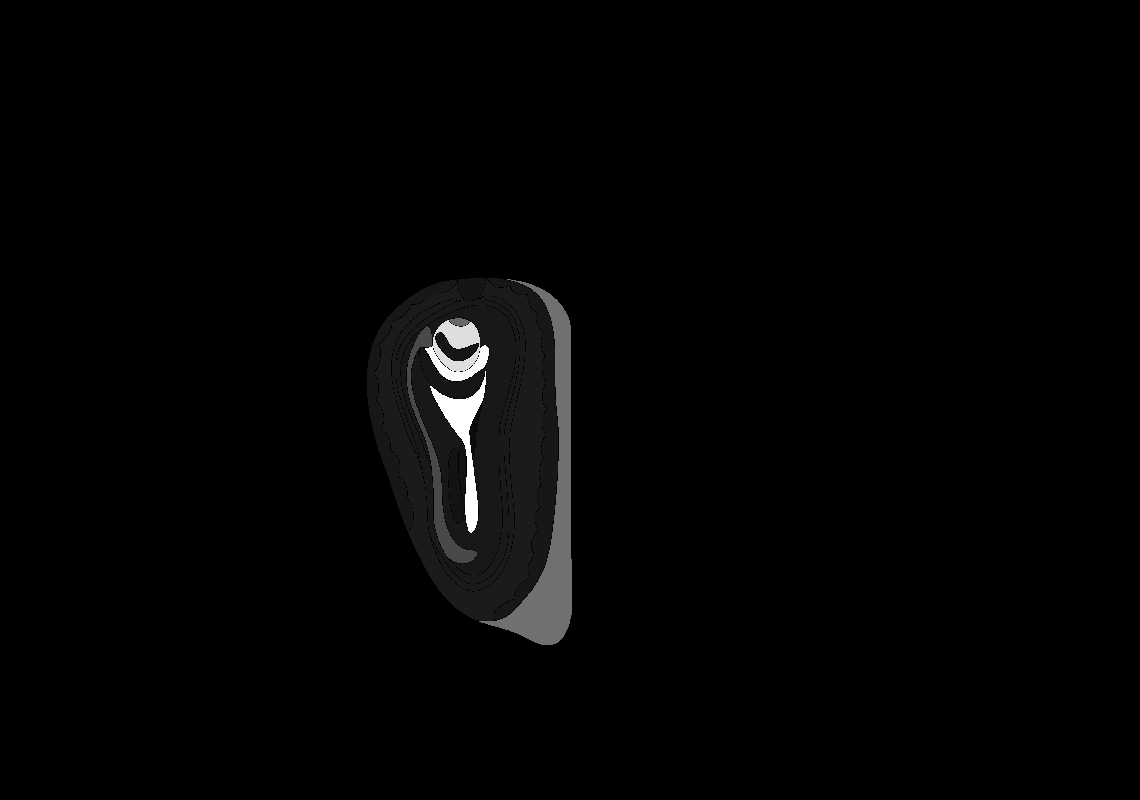

### 11_AP+3.3.tif

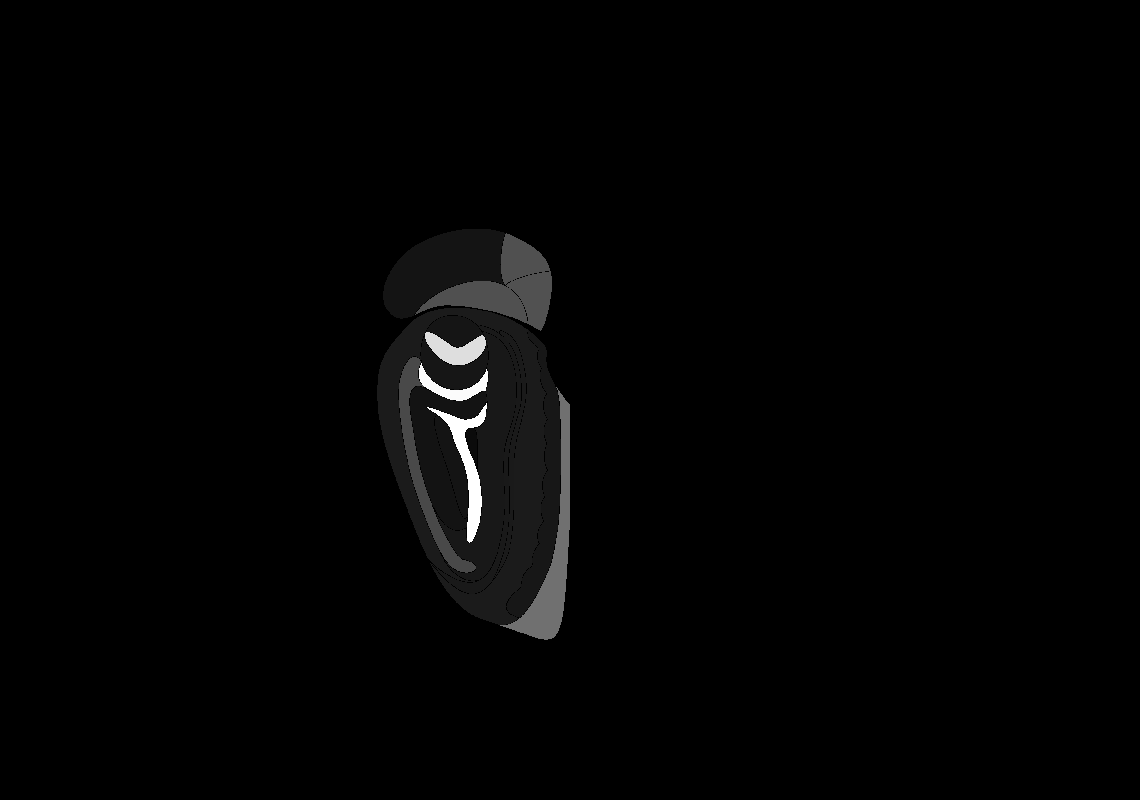

### 12_AP+3.2.tif

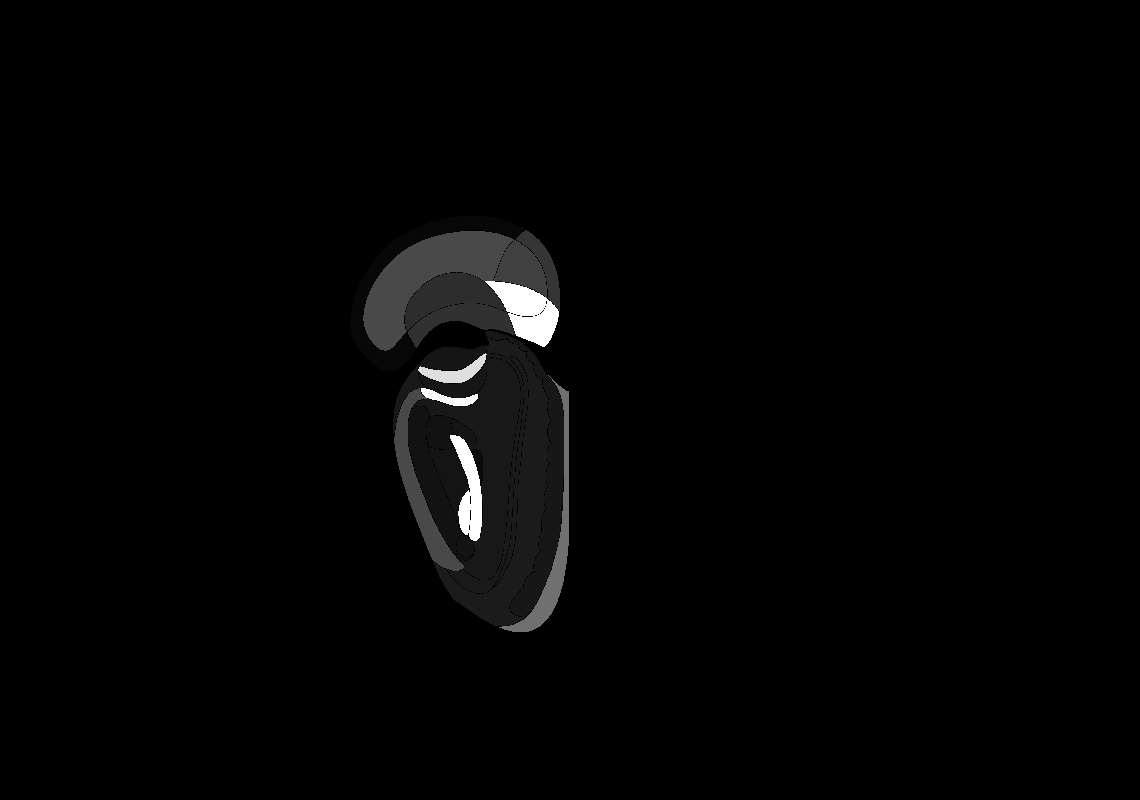

### 13_AP+3.1.tif

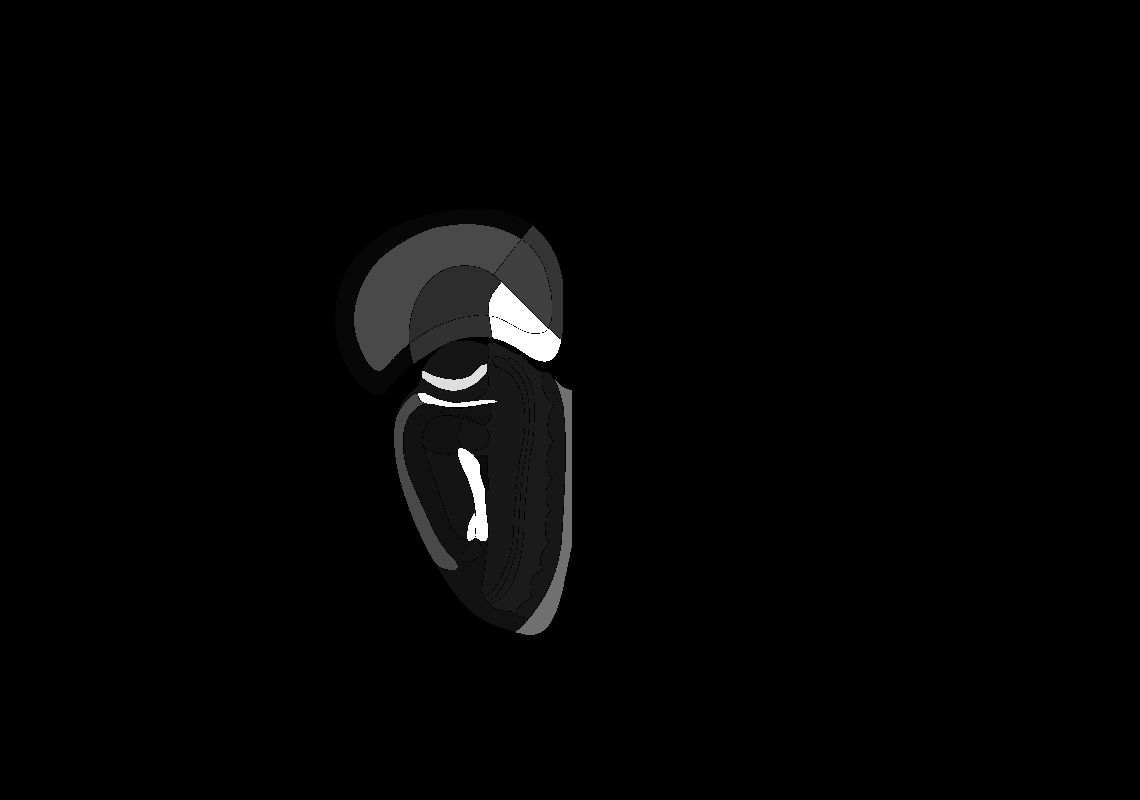

### 14_AP+3.0.tif

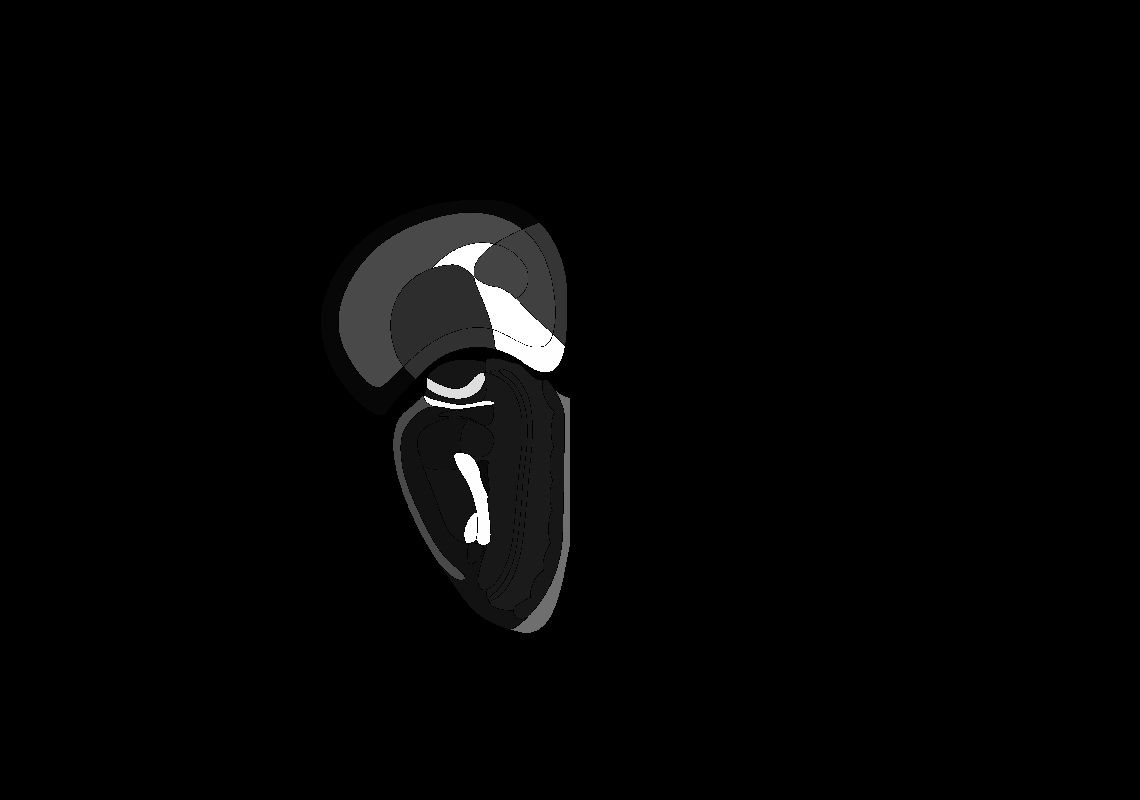

### 15_AP+2.9.tif

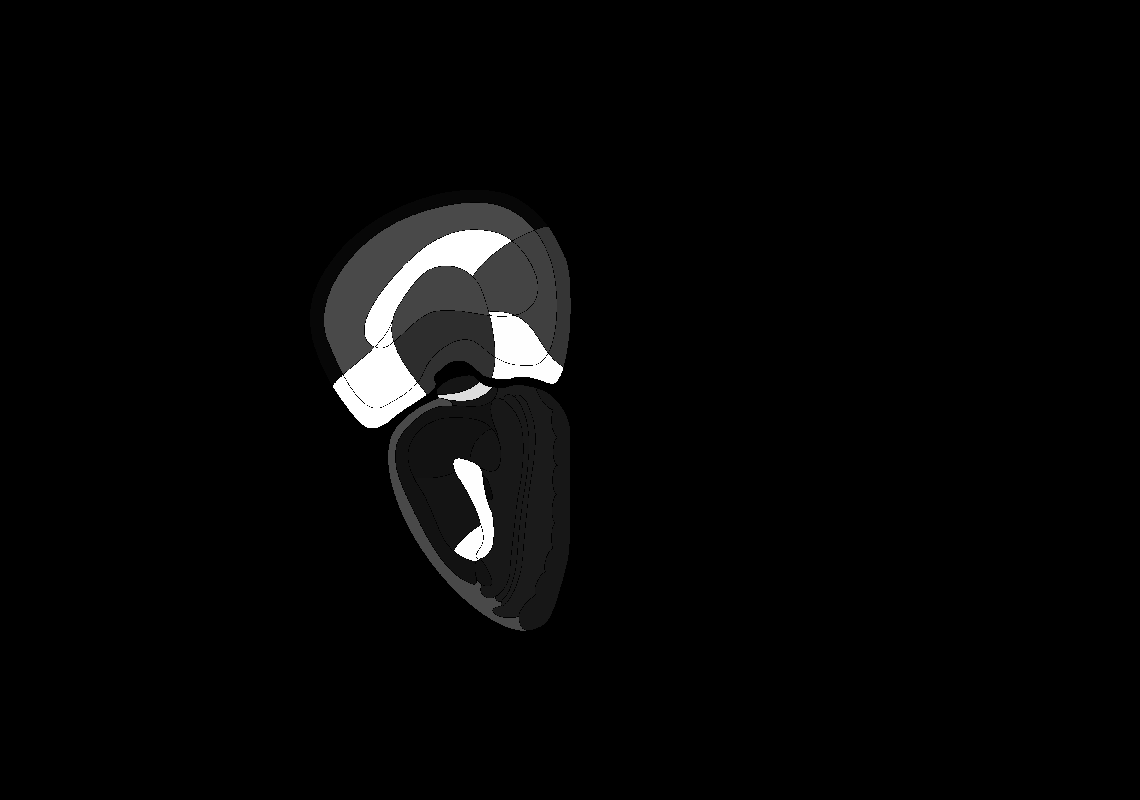

### 16_AP+2.8.tif

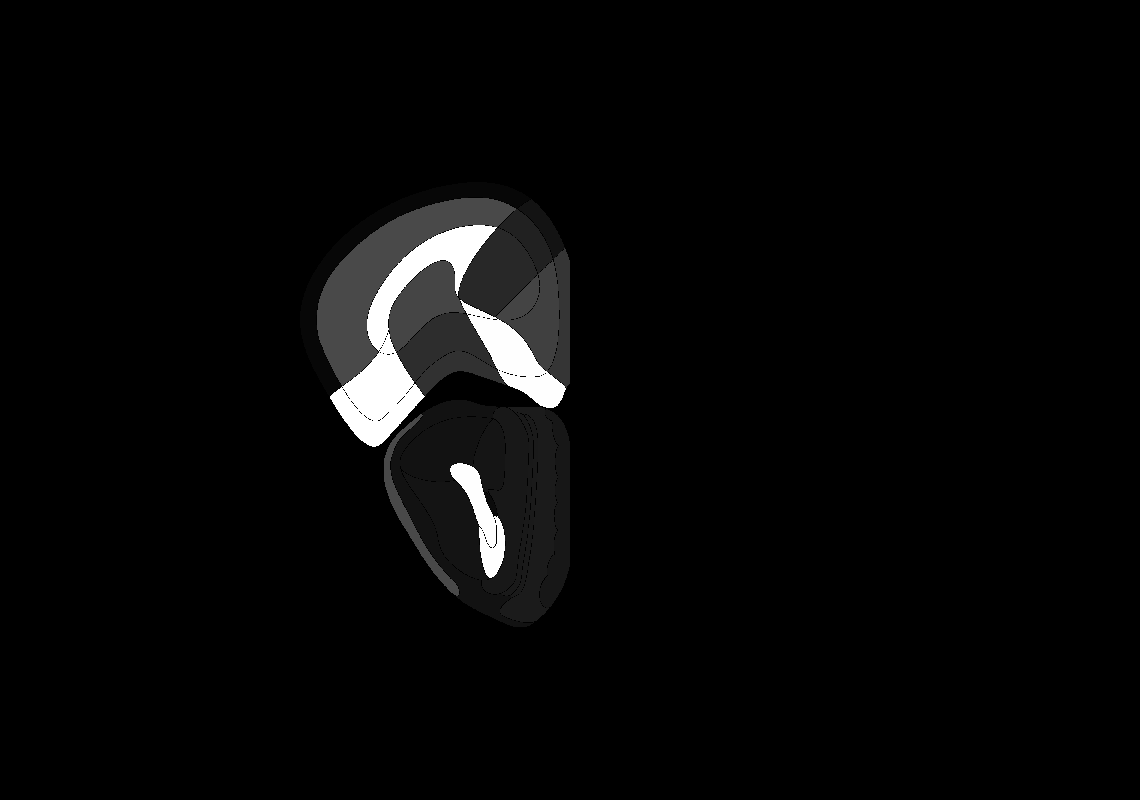

### 18_AP+2.6.tif

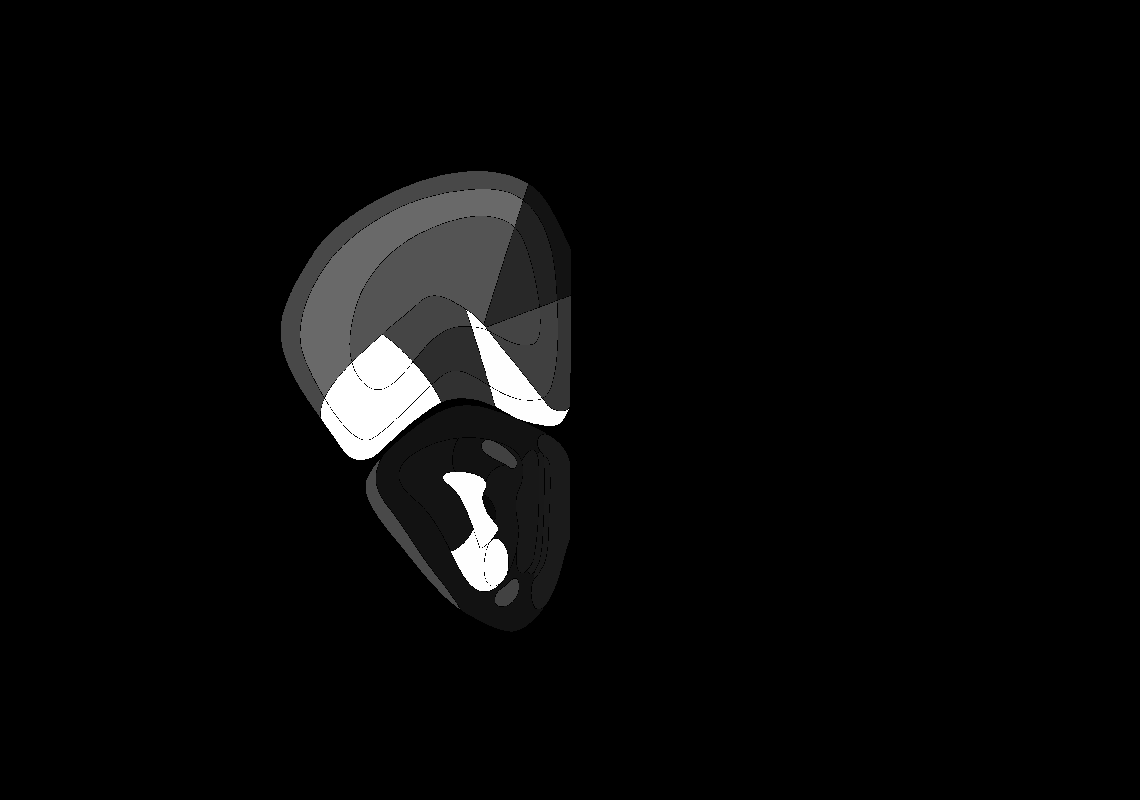

### 19_AP+2.5.tif

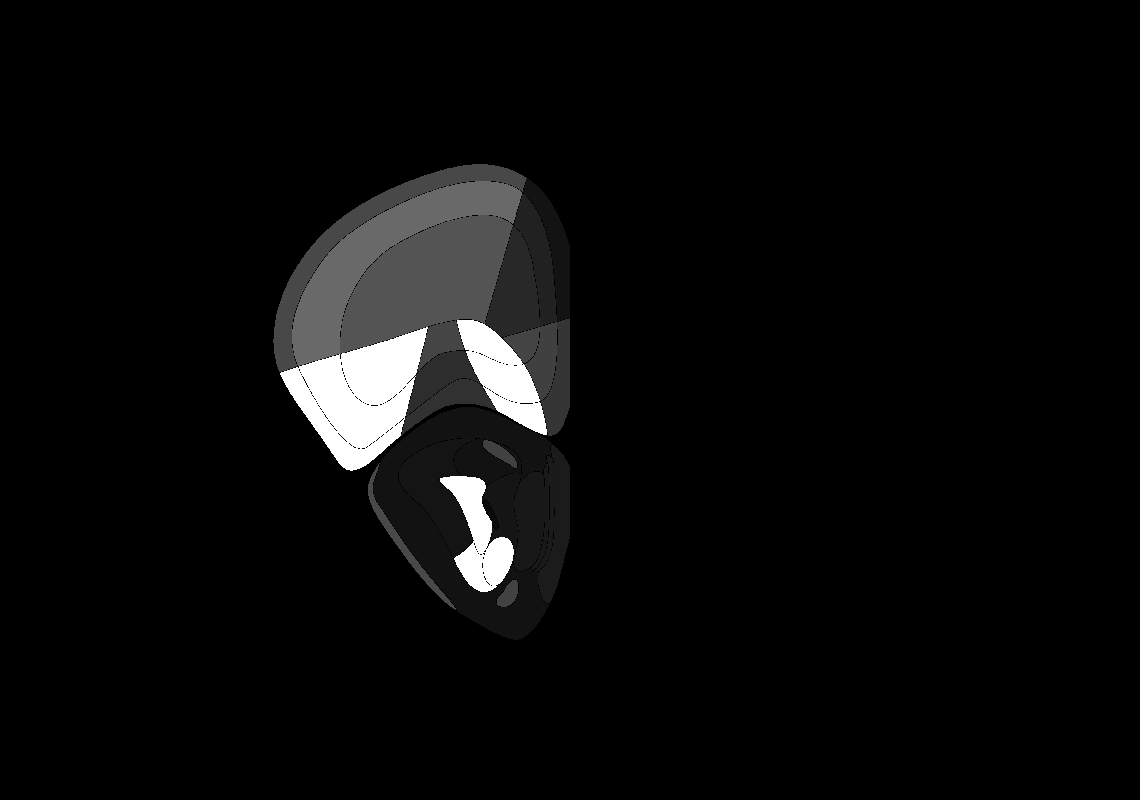

### 21_AP+2.3.tif

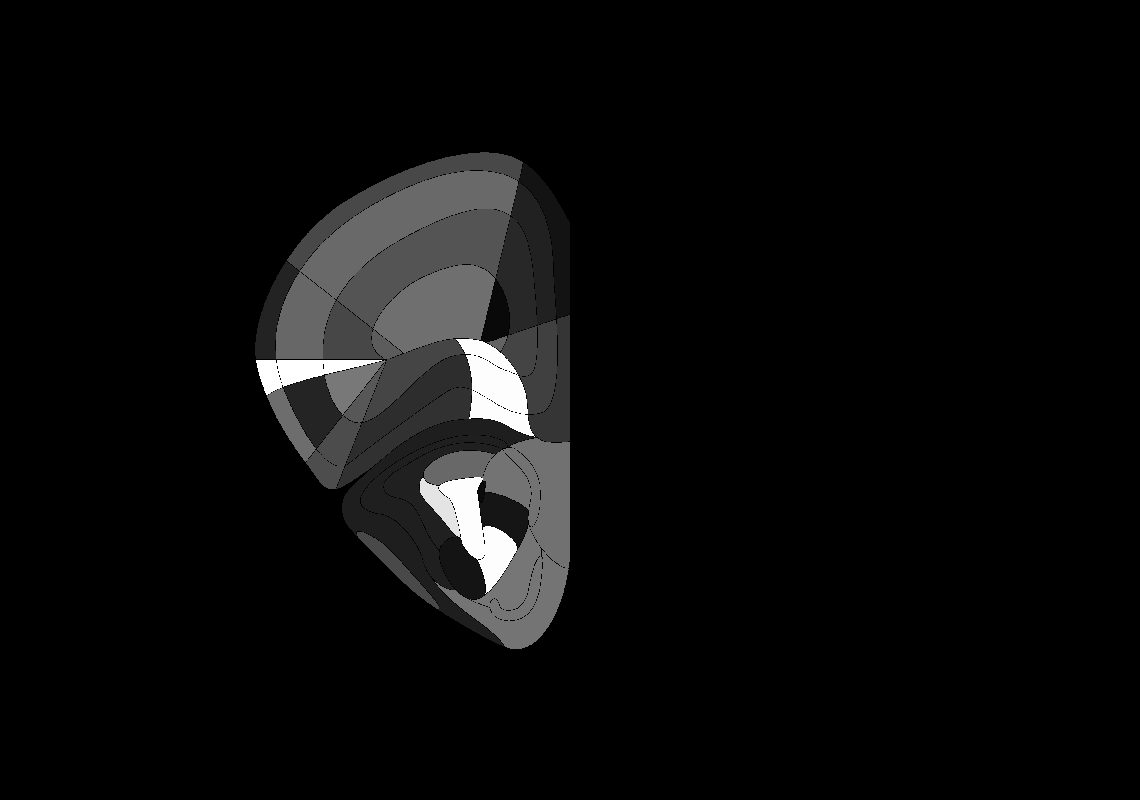

### 22_AP+2.2.tif

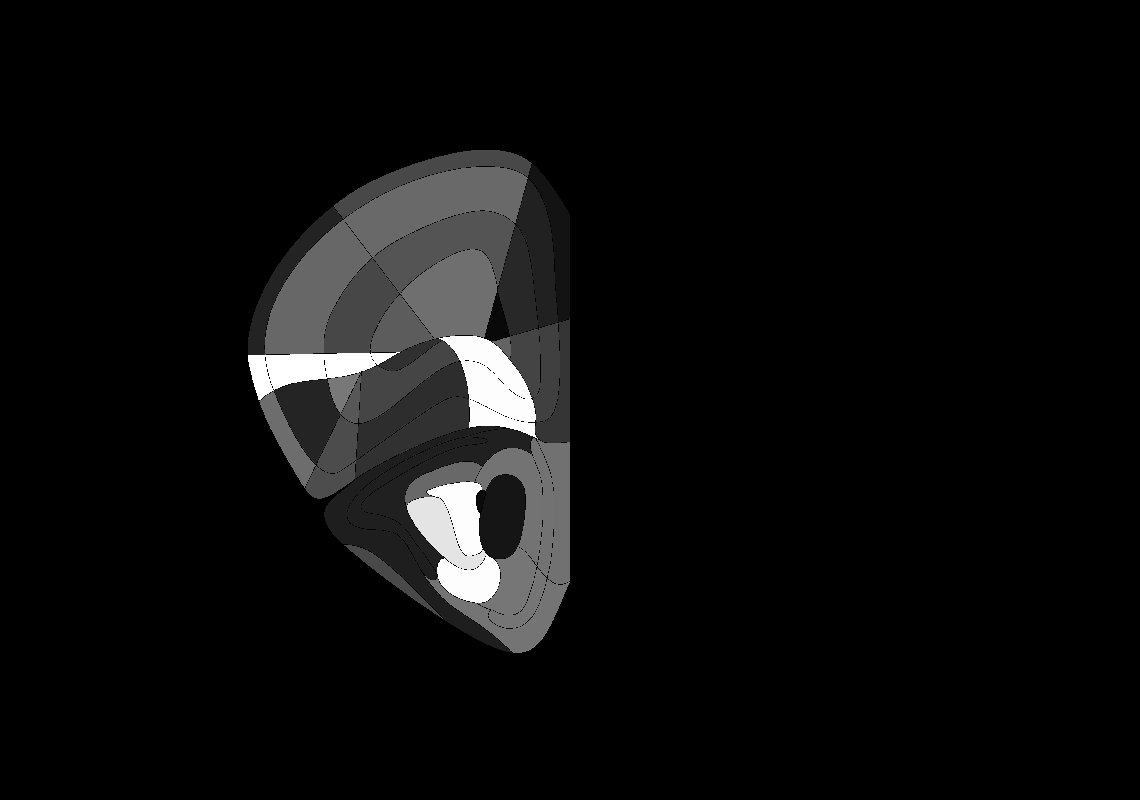

### 23_AP+2.1.tif

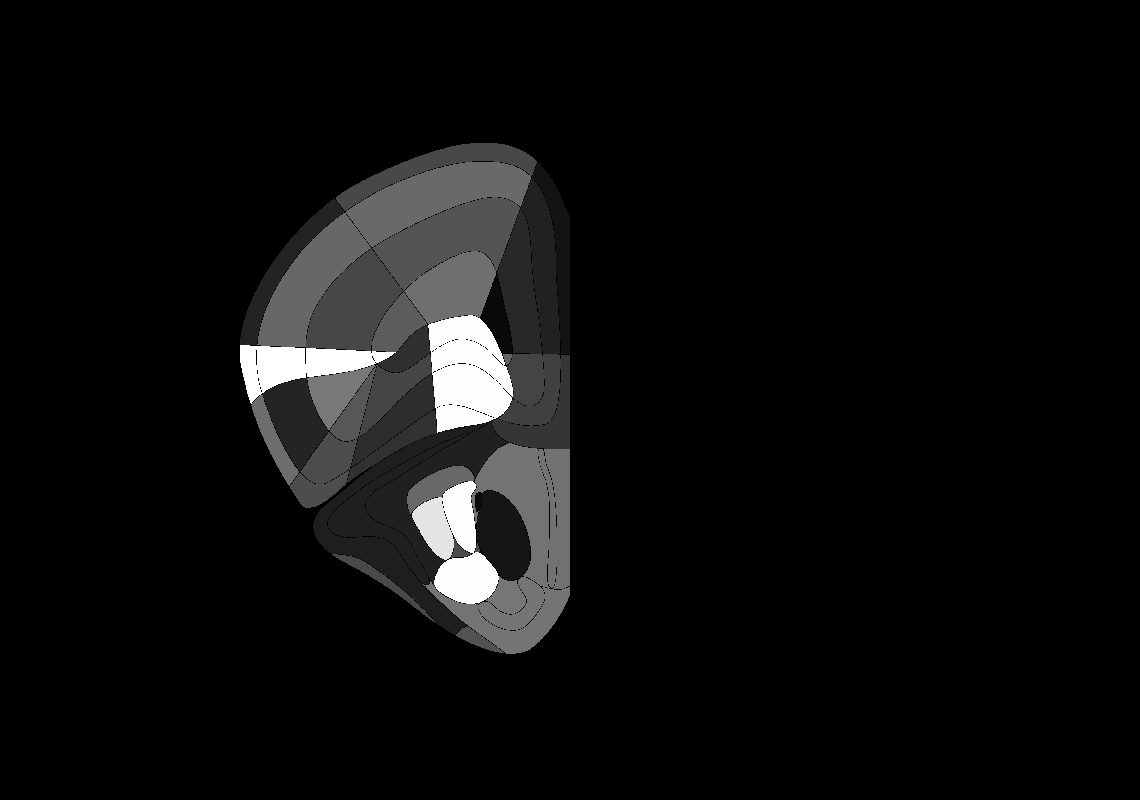

### 24_AP+2.0.tif

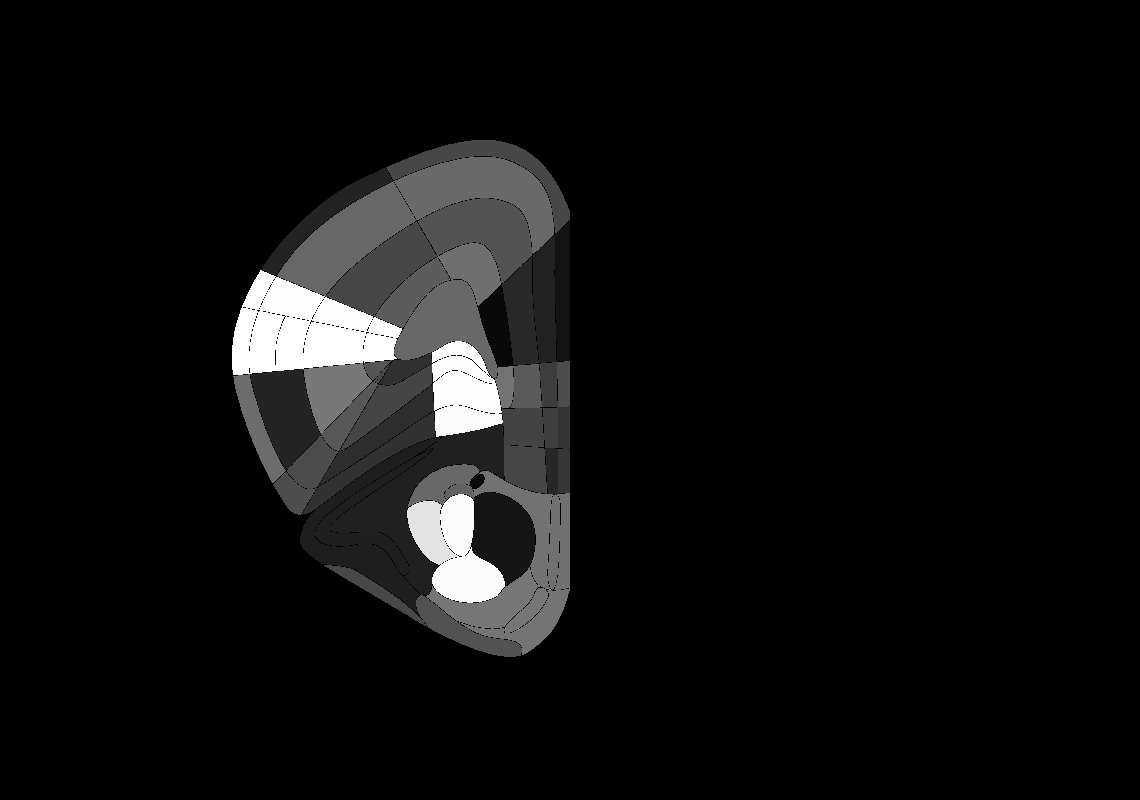

### 25_AP+1.9.tif

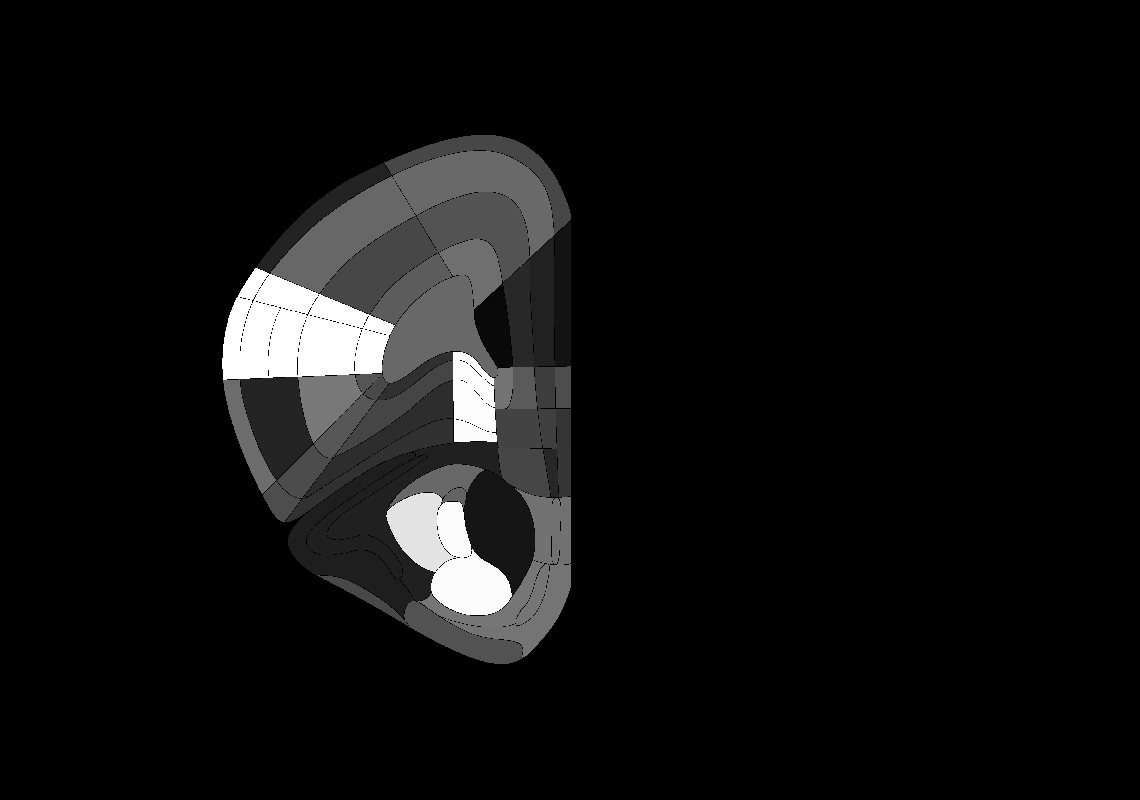

### 26_AP+1.8.tif

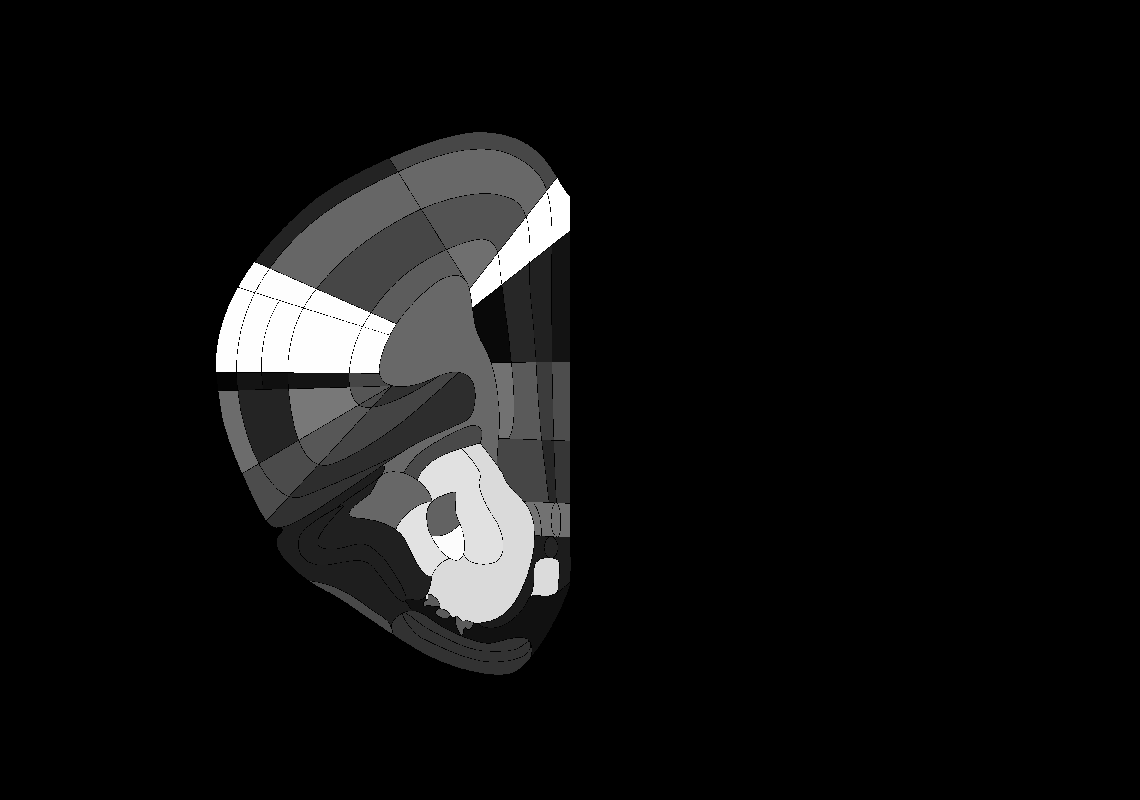

### 27_AP+1.7.tif

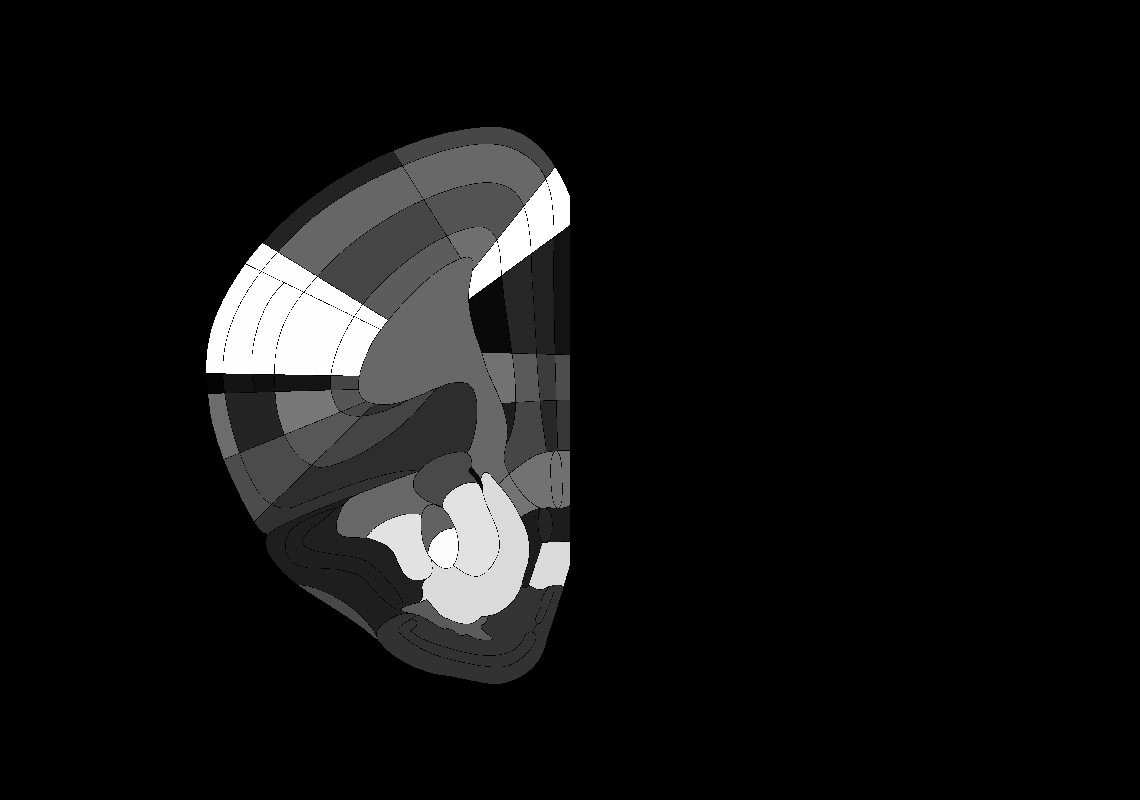

### 28_AP+1.6.tif

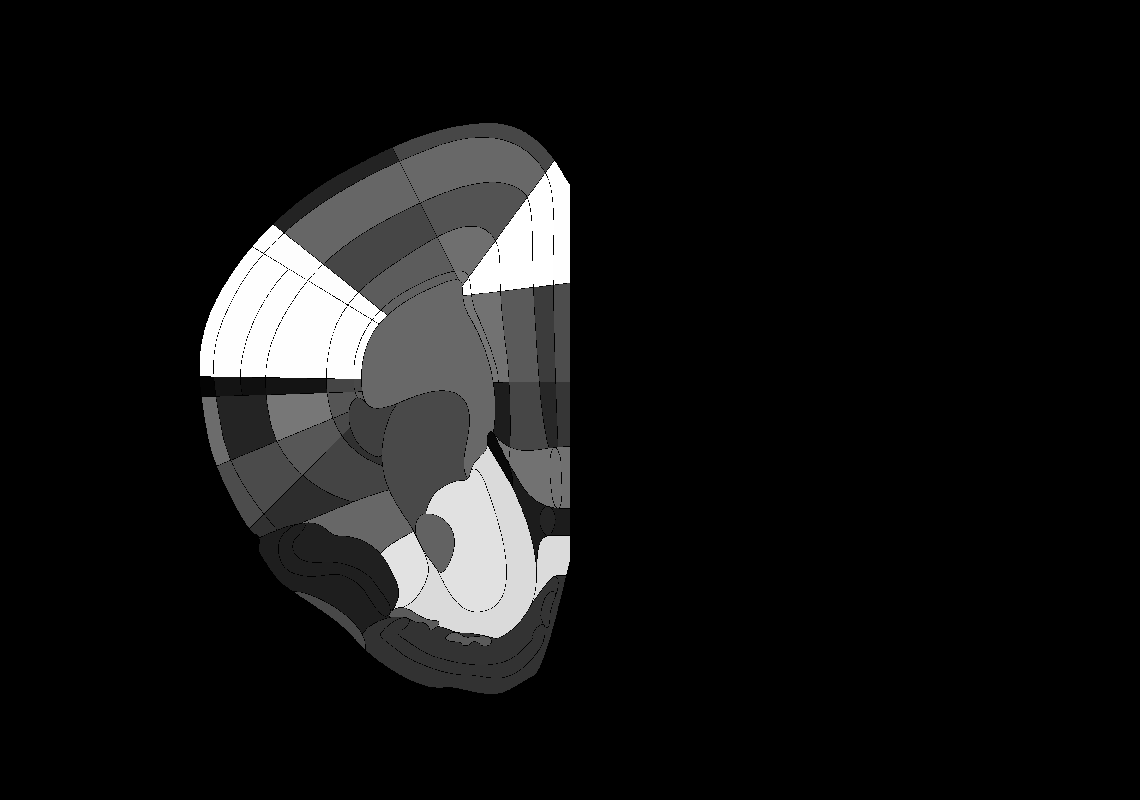

### 29_AP+1.5.tif

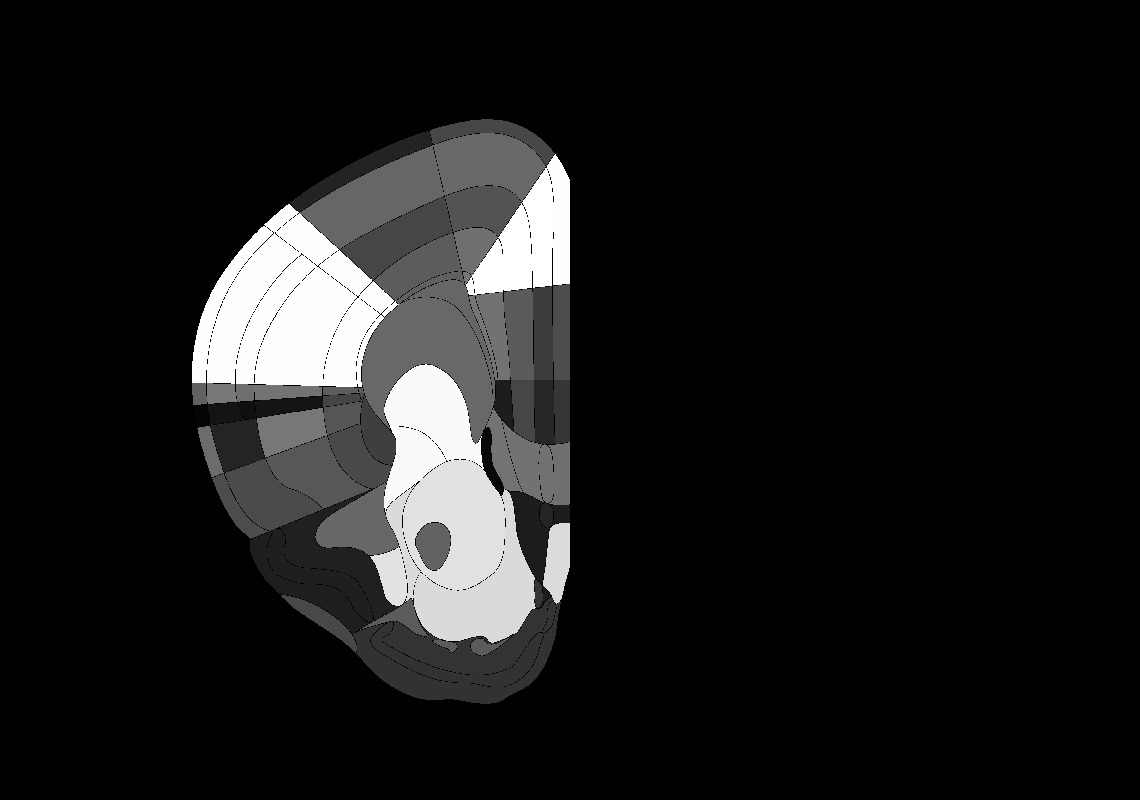

### 30_AP+1.4.tif

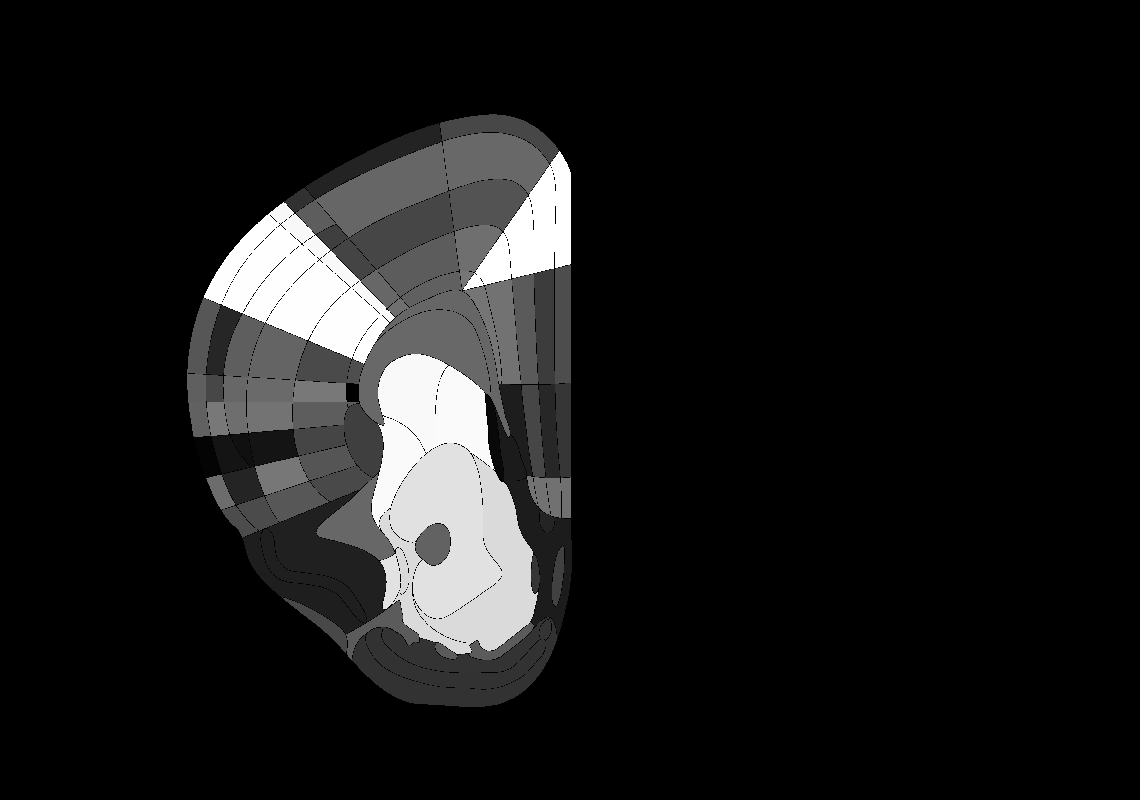

### 31_AP+1.3.tif

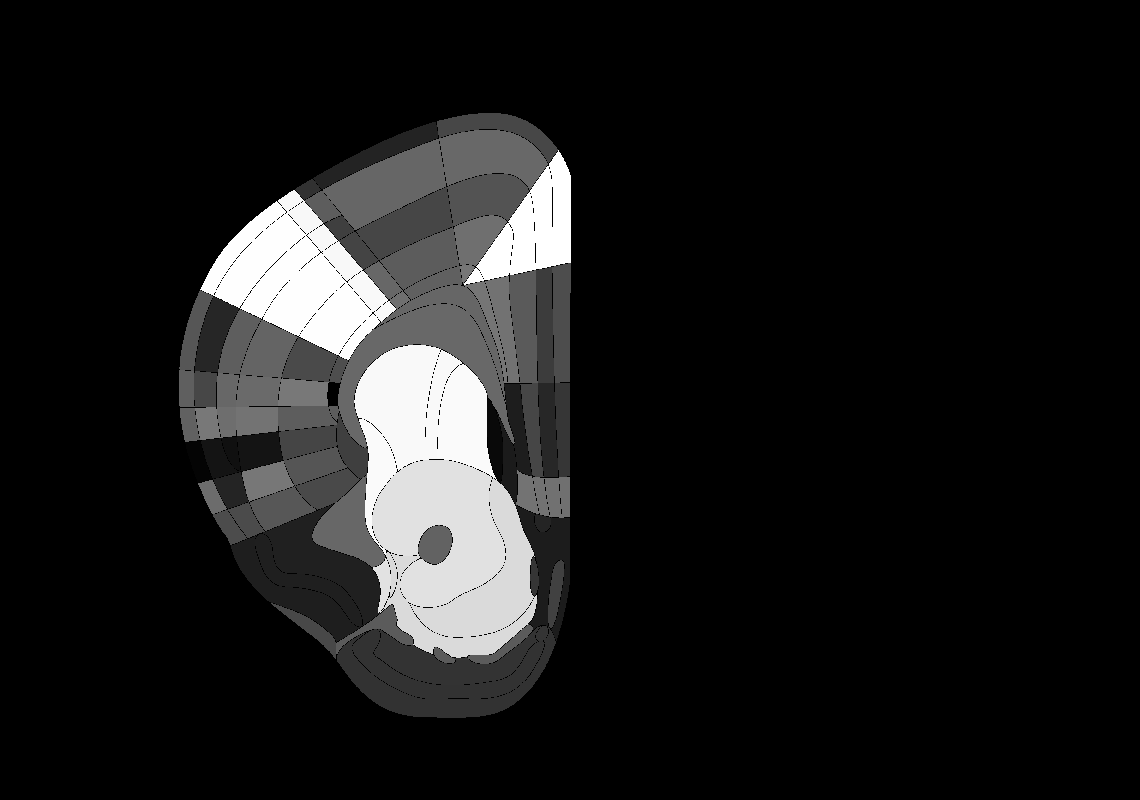

### 32_AP+1.2.tif

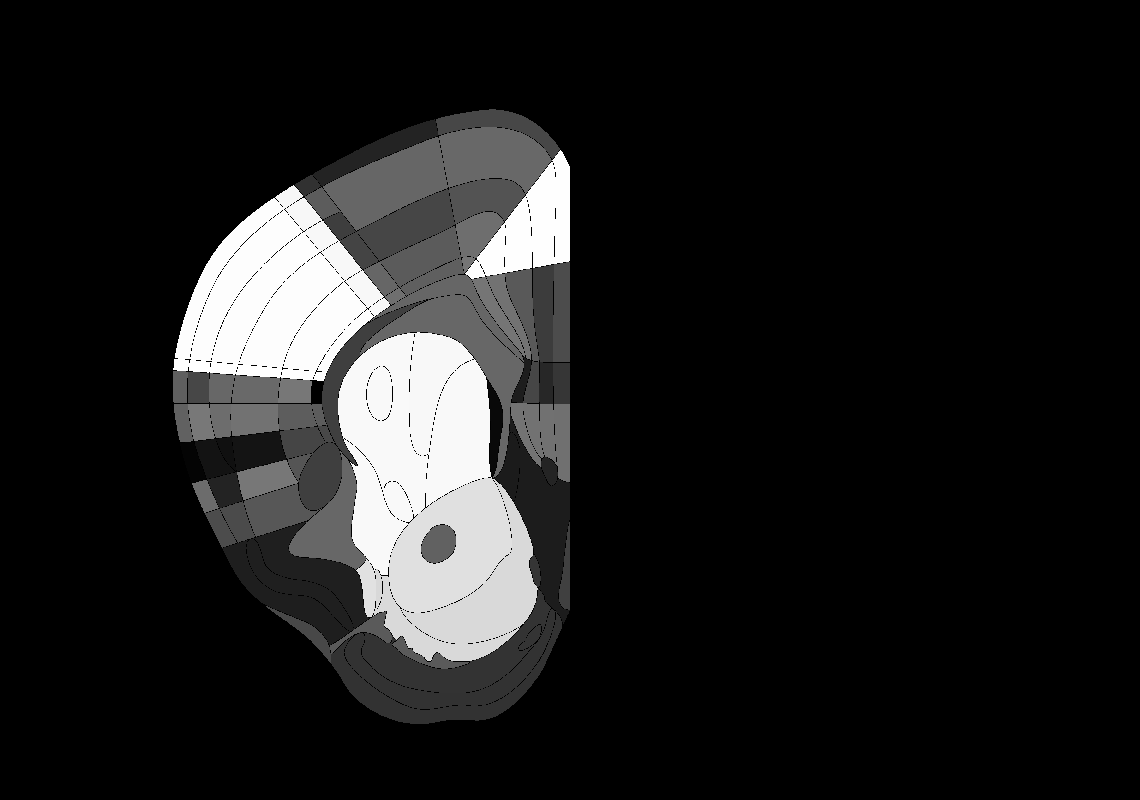

### 33_AP+1.1.tif

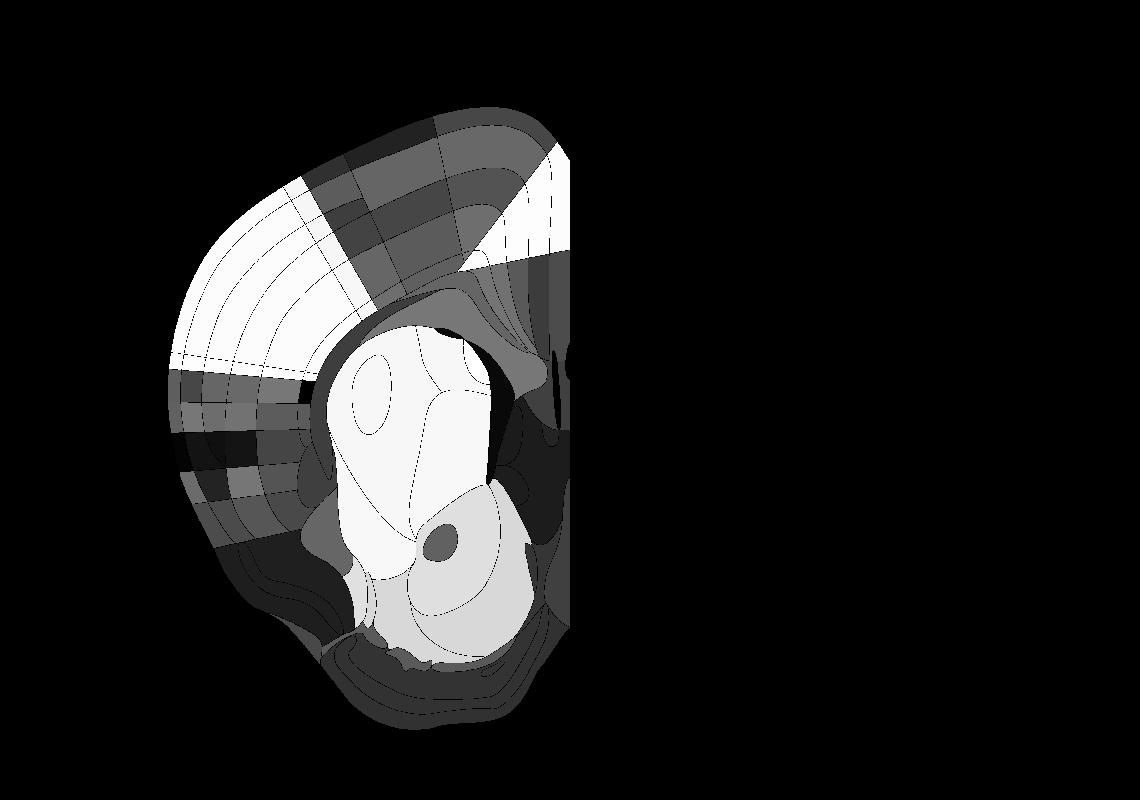

### 34_AP+1.0.tif

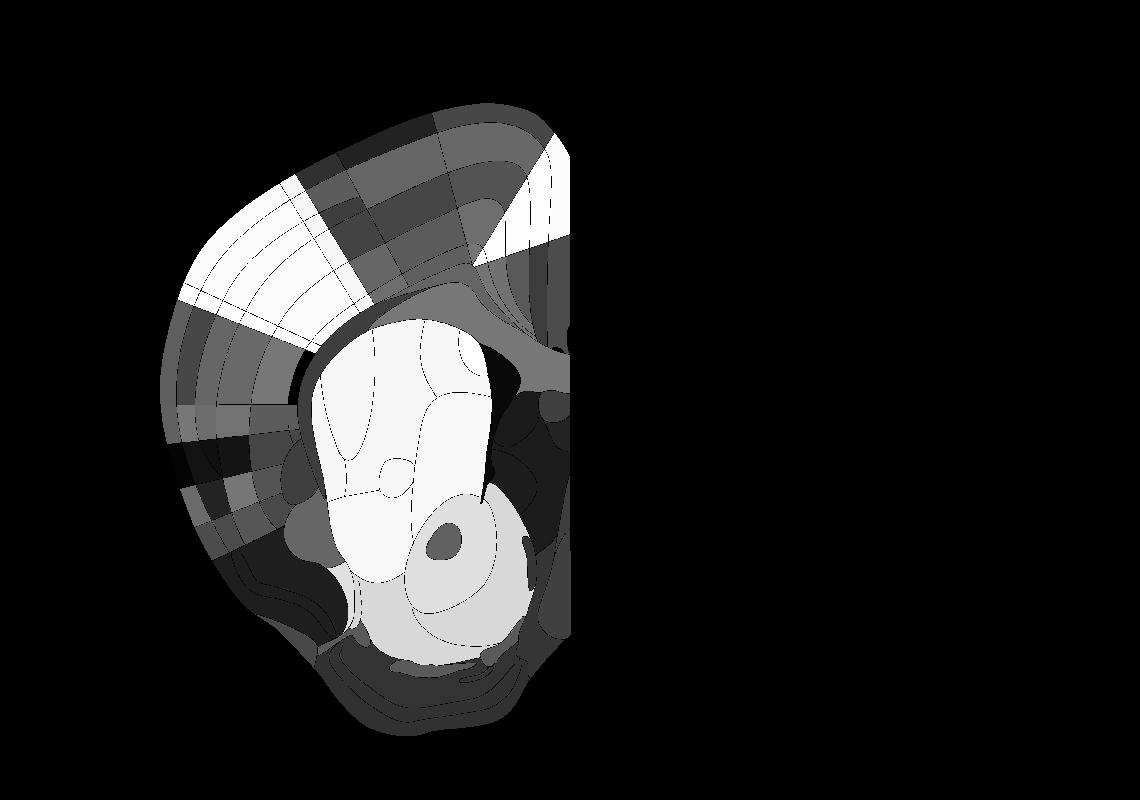

### 35_AP+0.9.tif

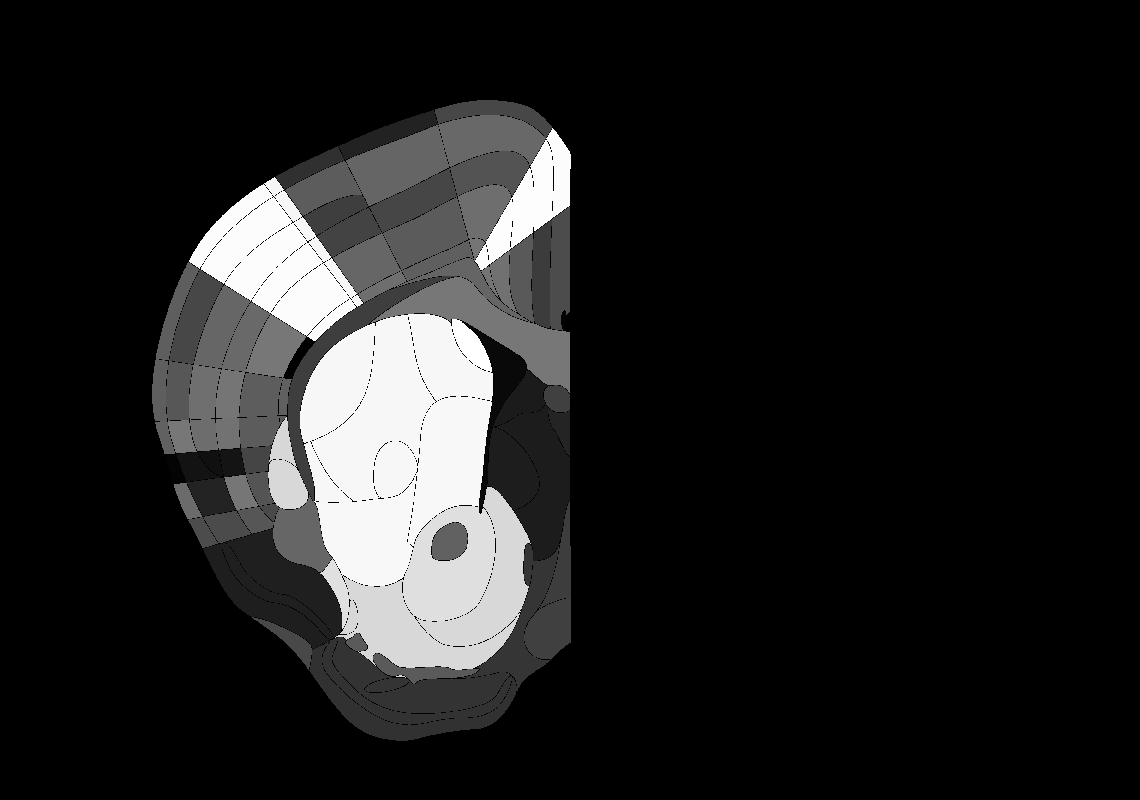

### 36_AP+0.8.tif

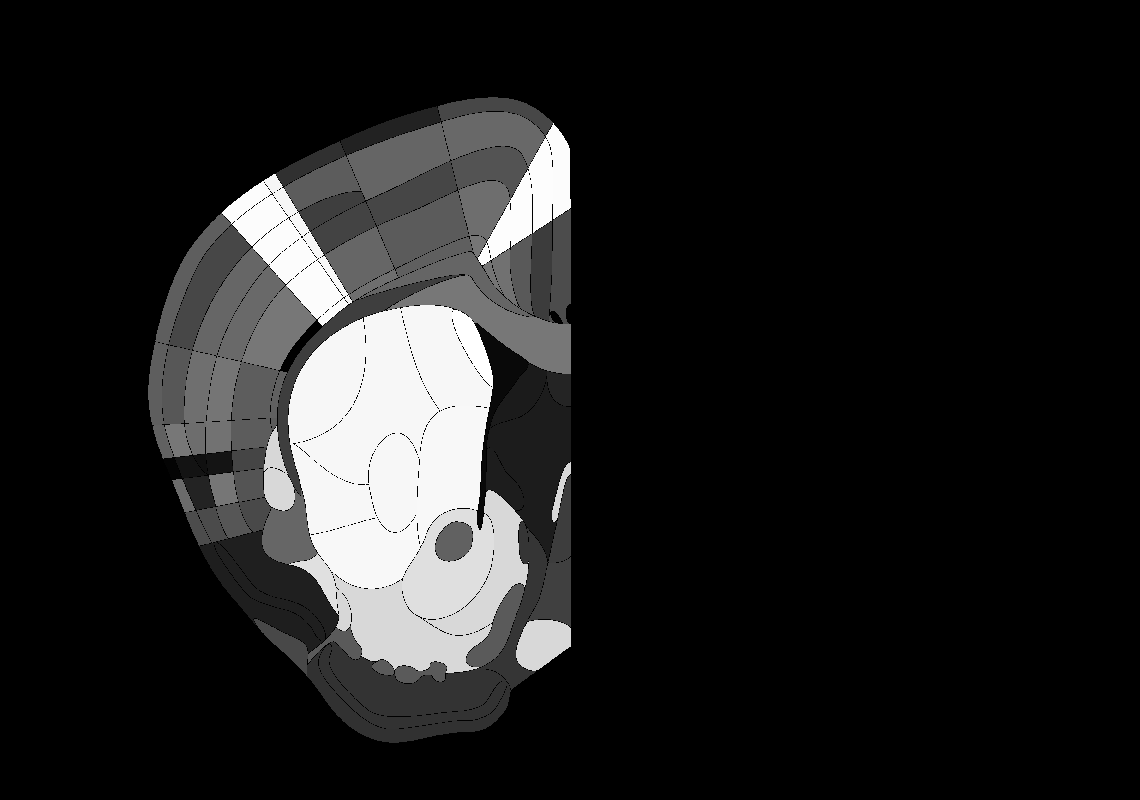

### 37_AP+0.7.tif

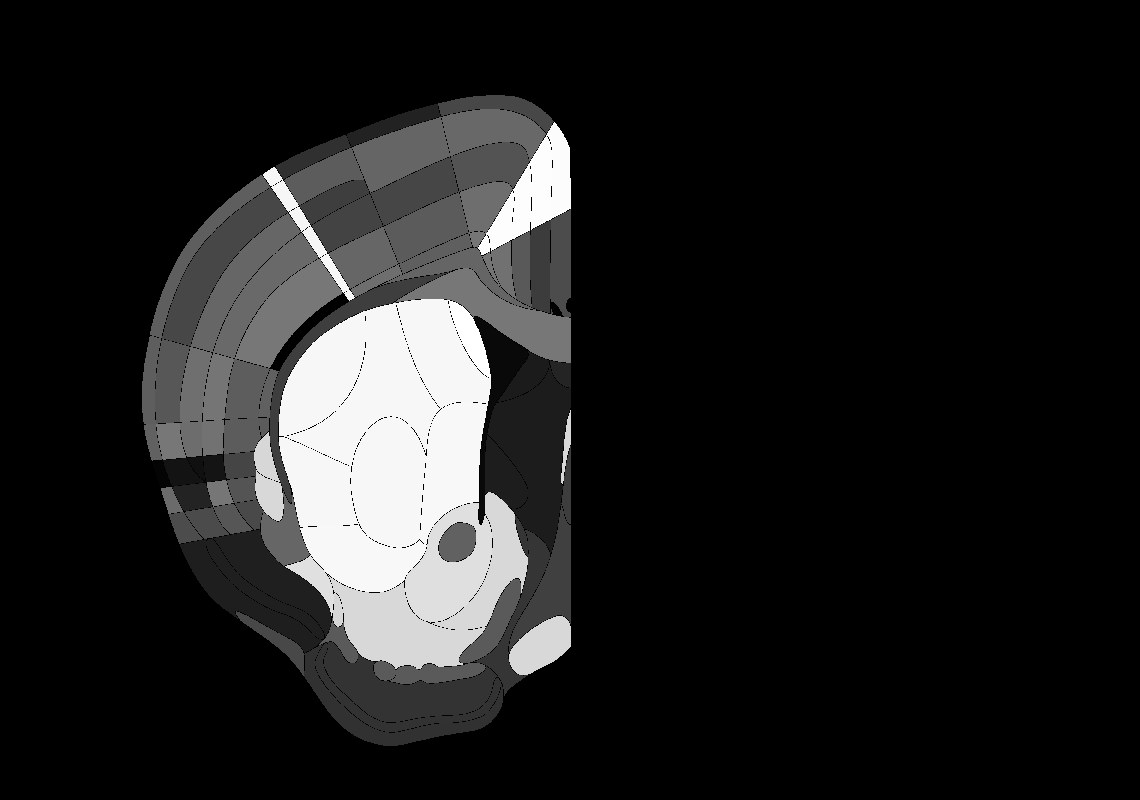

### 38_AP+0.6.tif

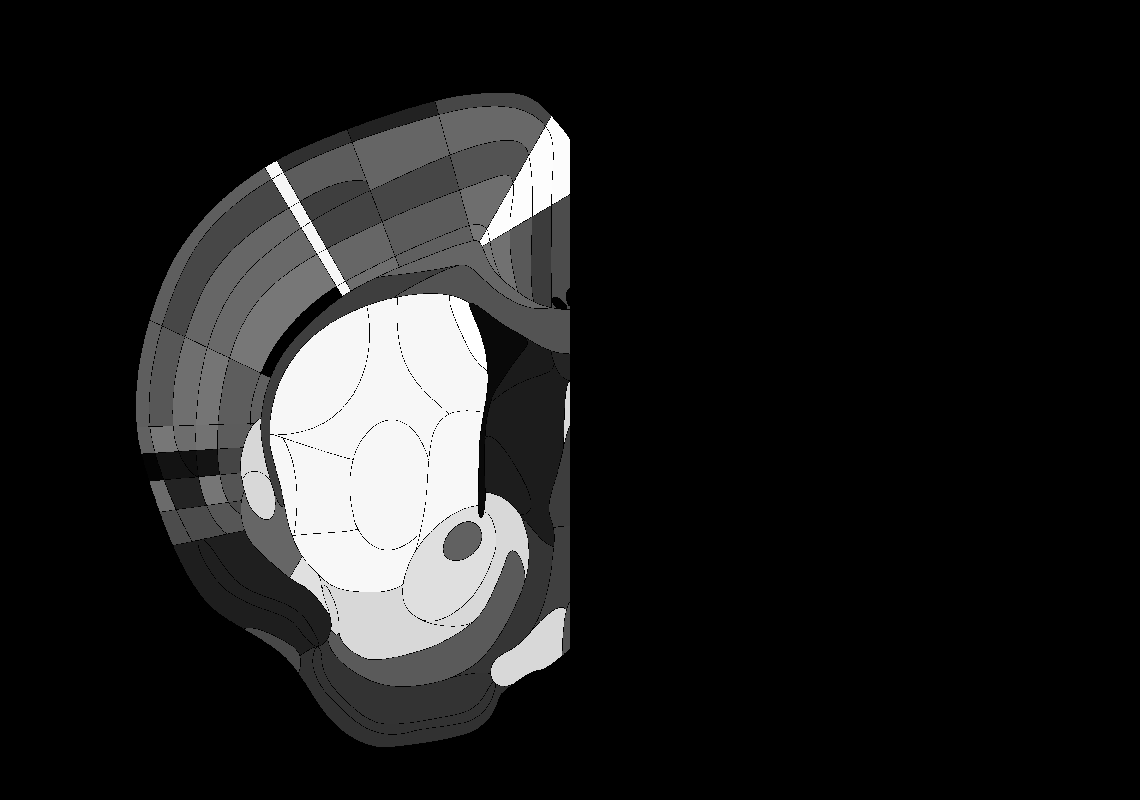

### 39_AP+0.5.tif

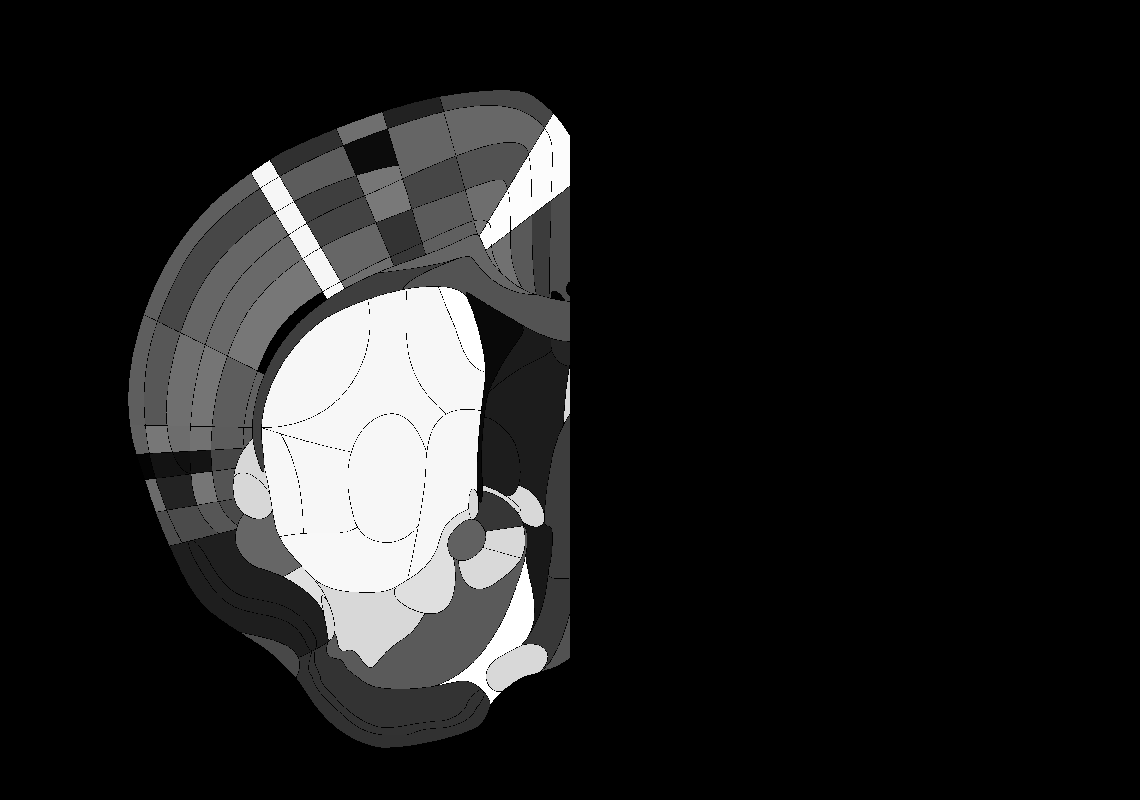

### 40_AP+0.4.tif

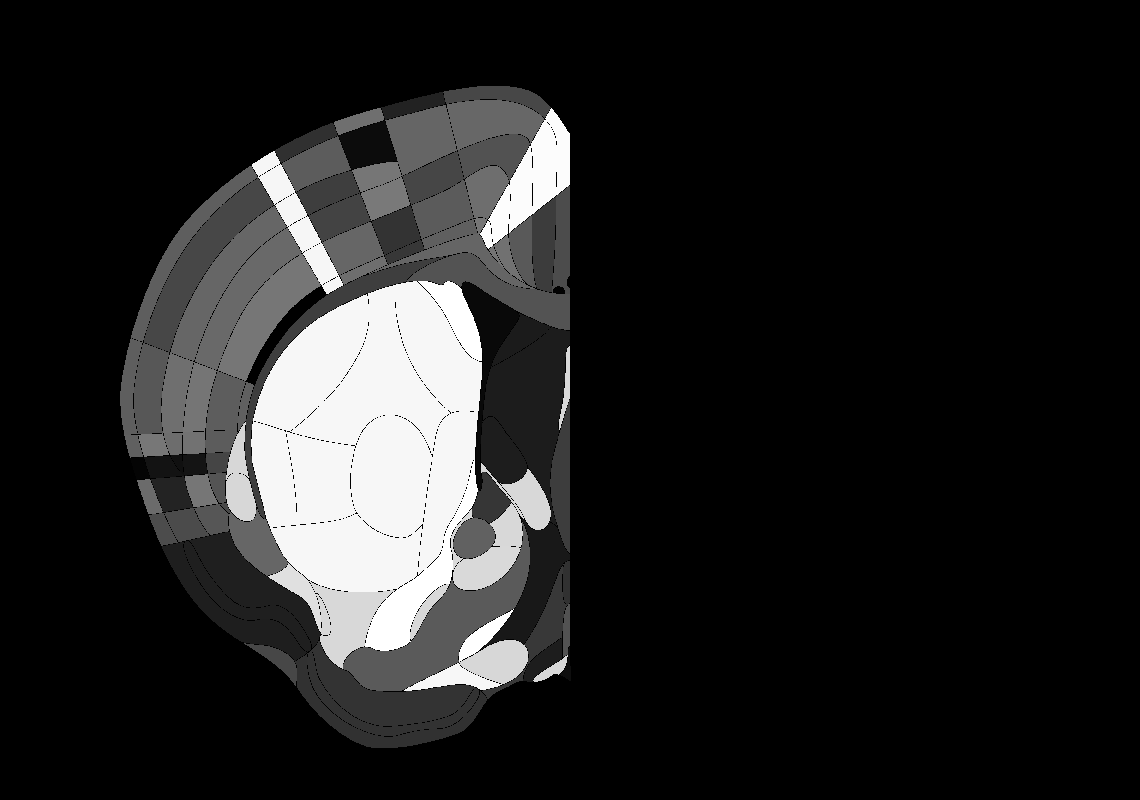

### 41_AP+0.3.tif

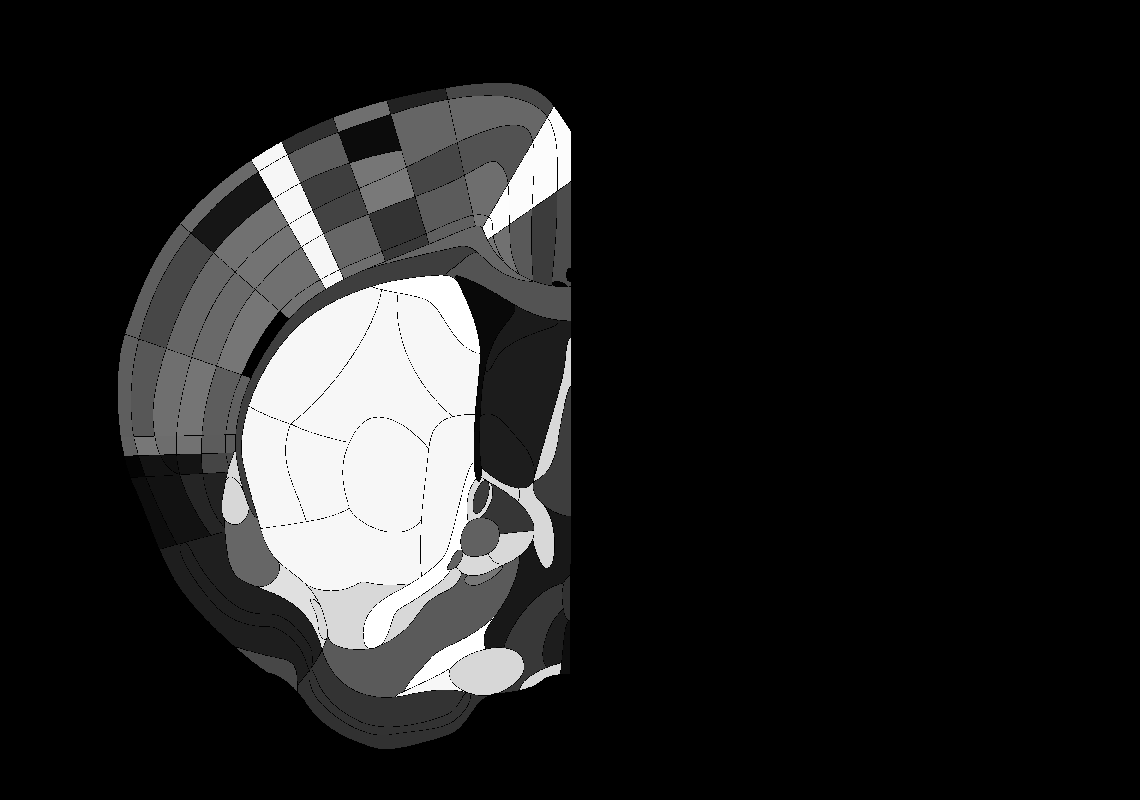

### 43_AP+0.1.tif

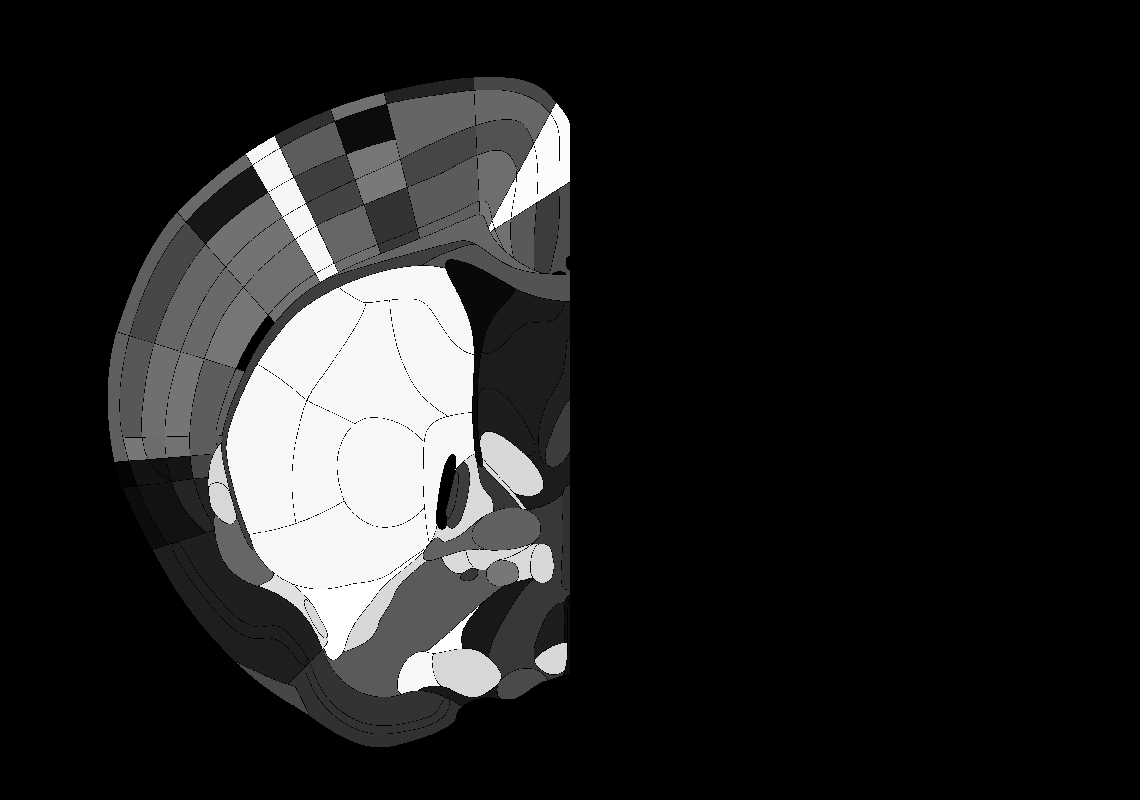

### 44_AP+0.0.tif

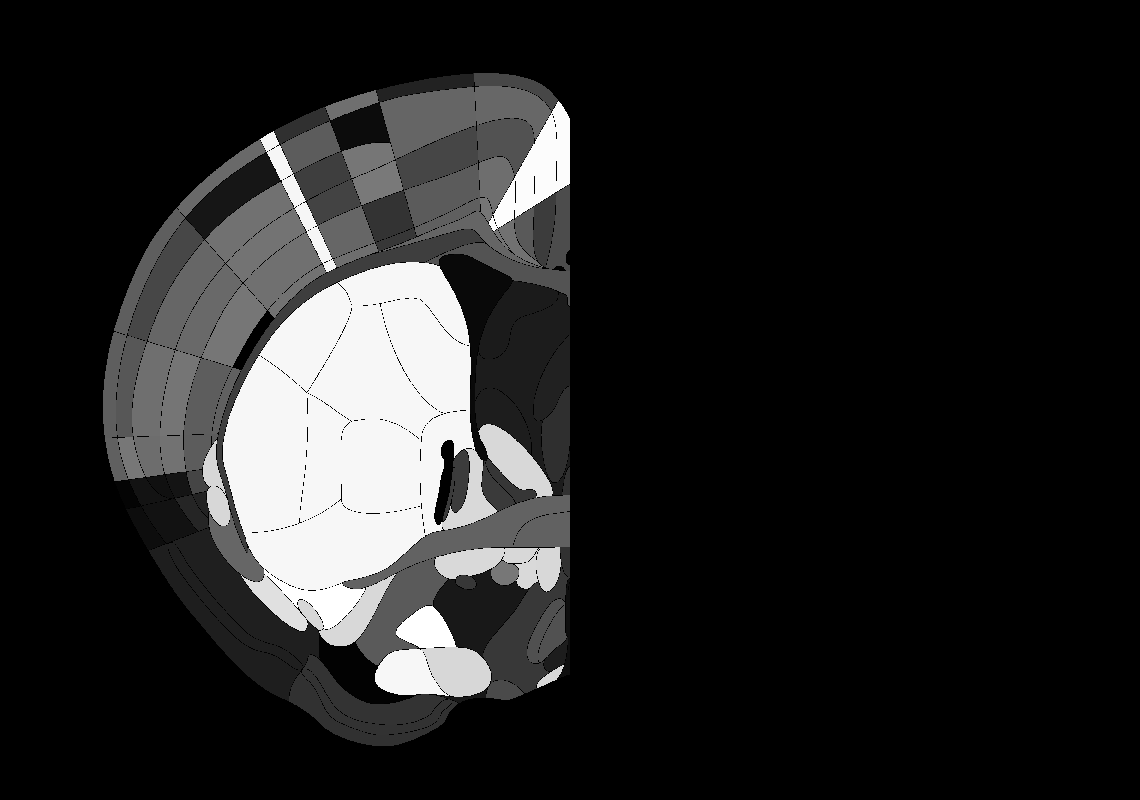

### 46_AP-0.2.tif

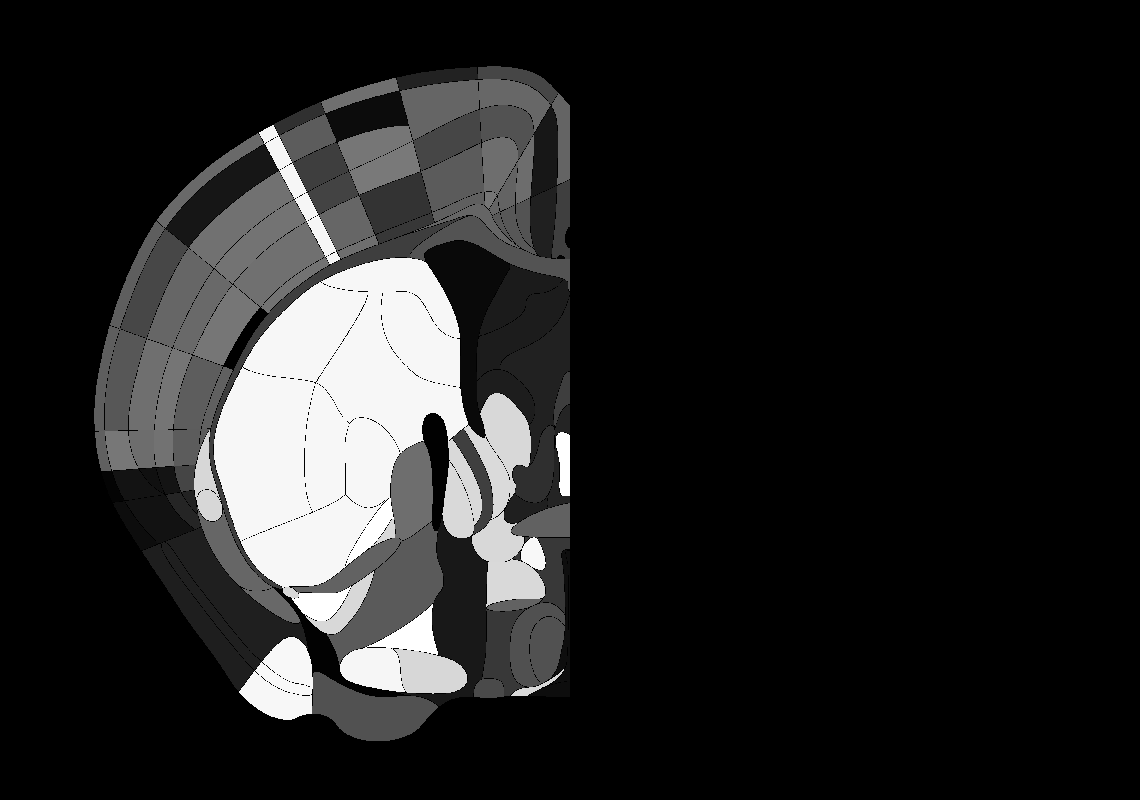

### 47_AP-0.3.tif

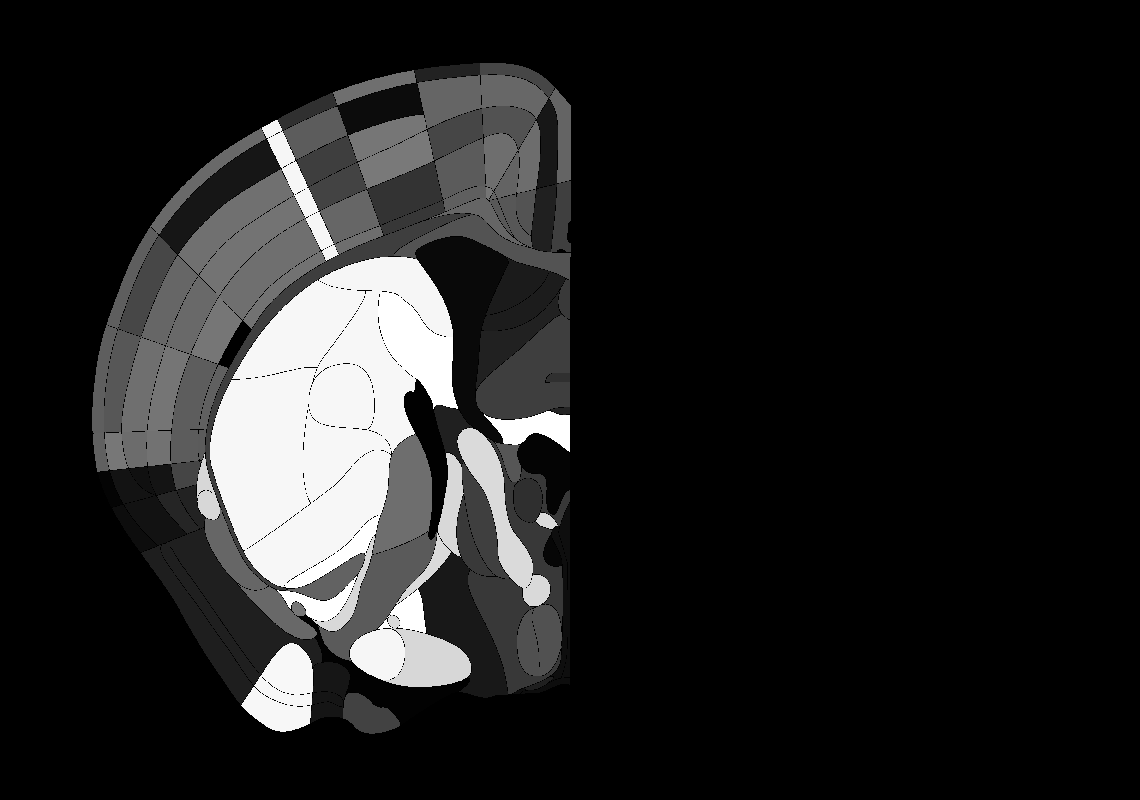

### 48_AP-0.4.tif

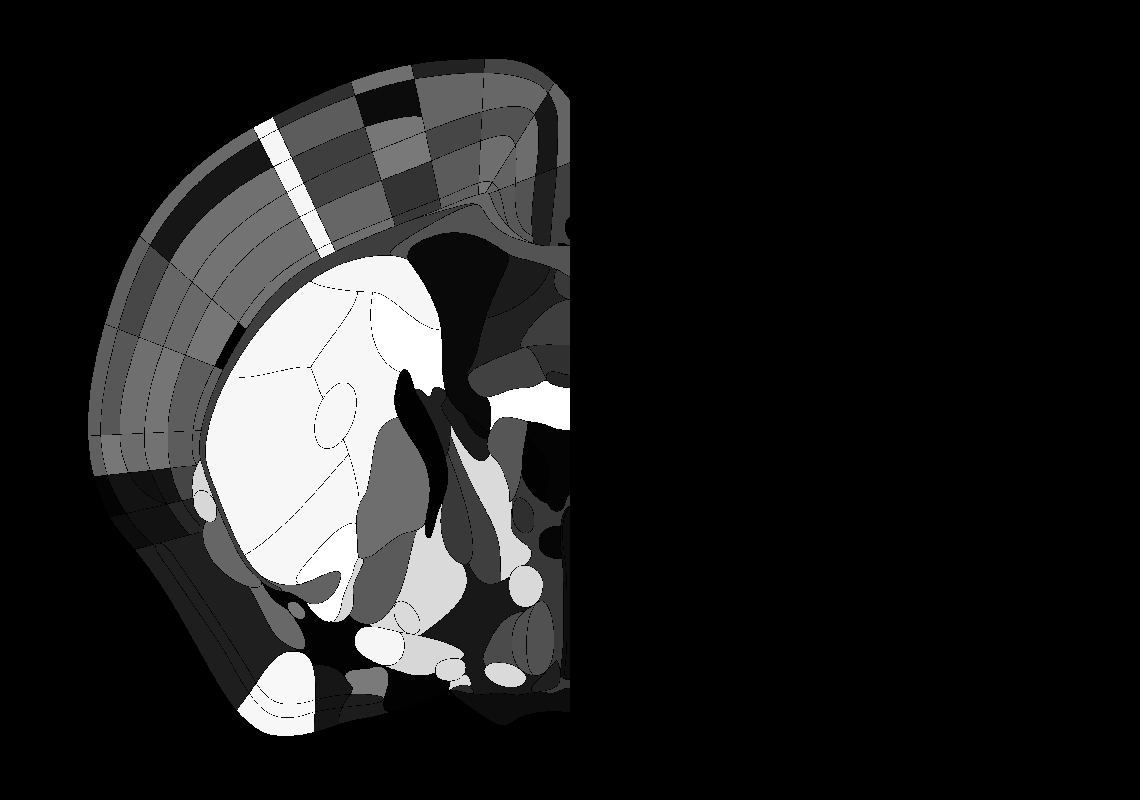

### 49_AP-0.5.tif

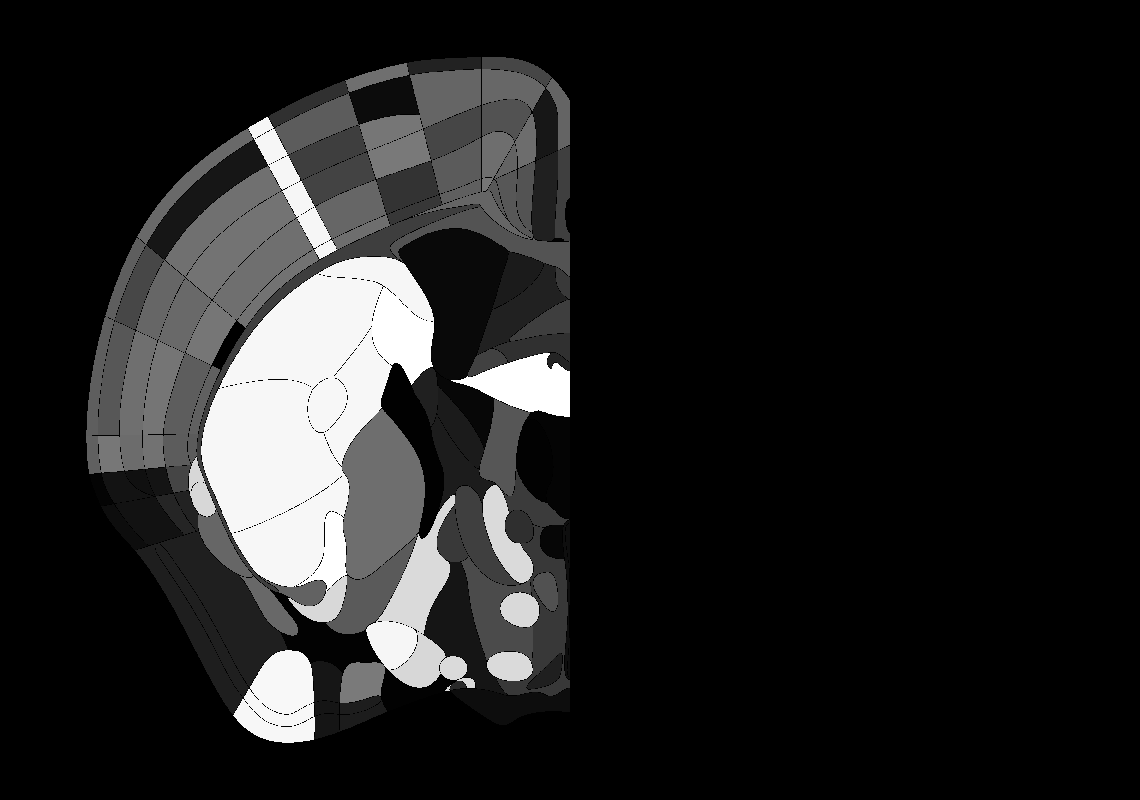

### 50_AP-0.6.tif

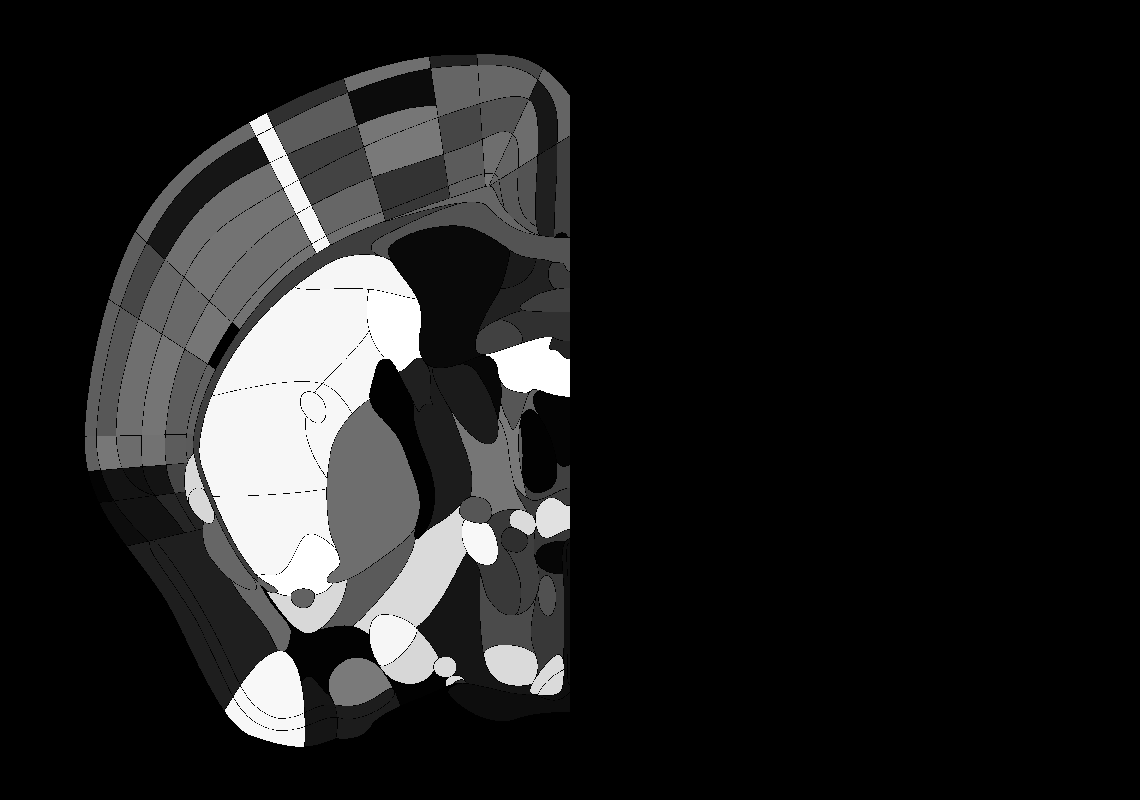

### 53_AP-0.9.tif

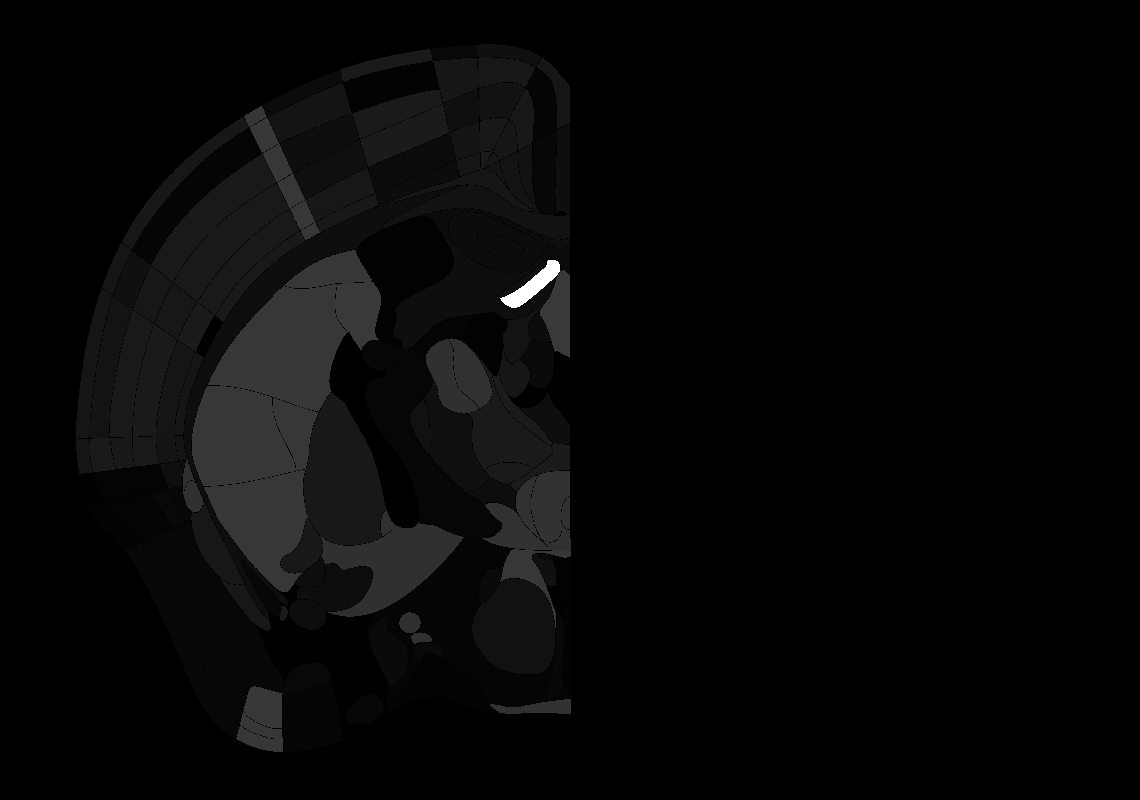

### 54_AP-1.0.tif

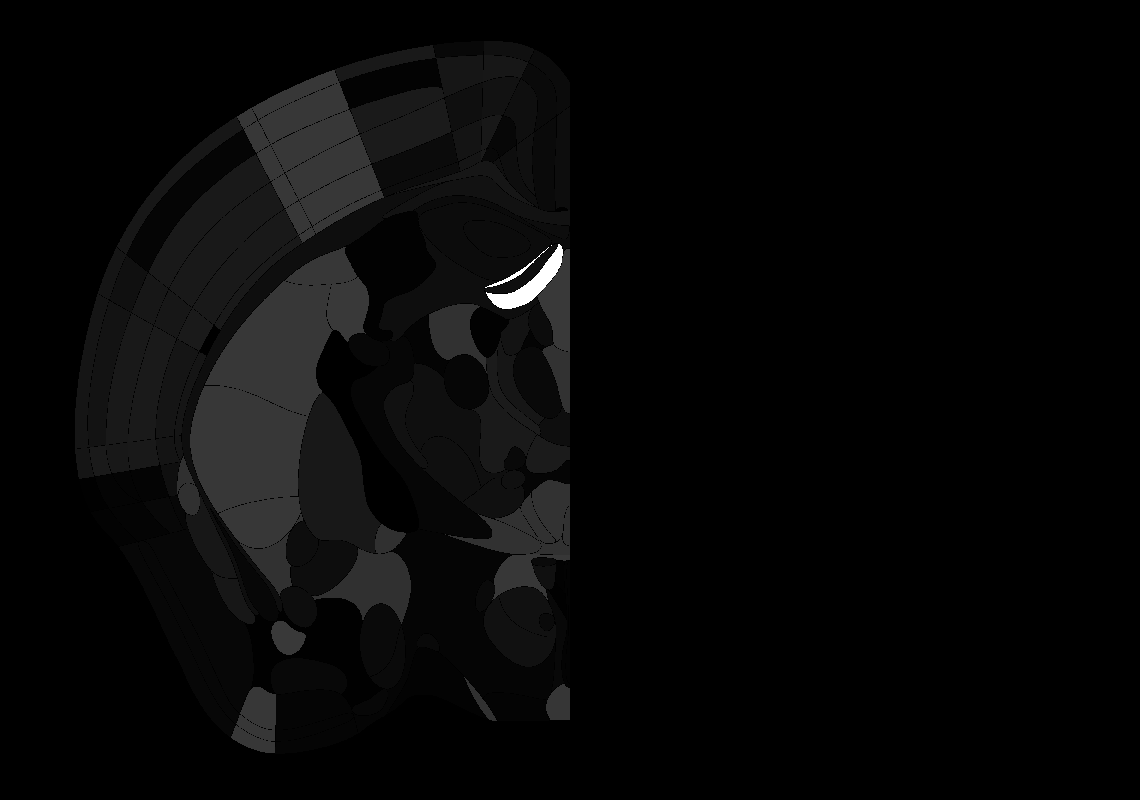

### 55_AP-1.1.tif

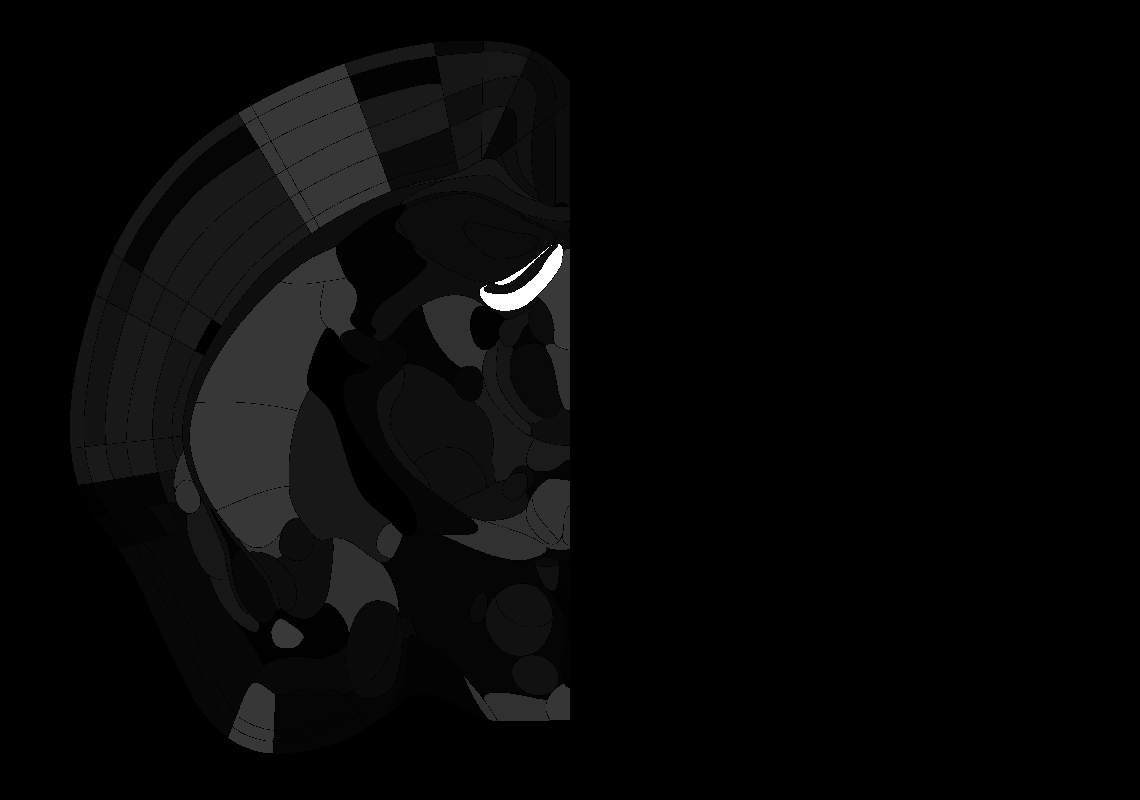

### 57_AP-1.3.tif

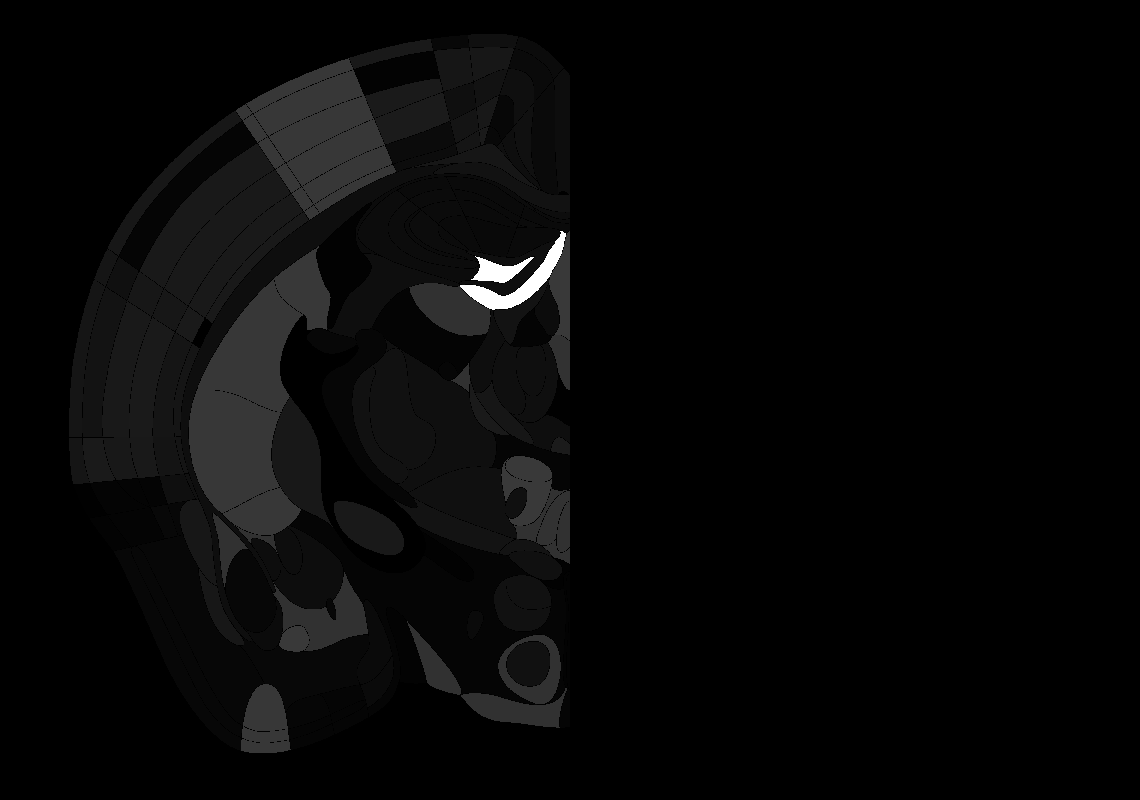

### 58_AP-1.4.tif

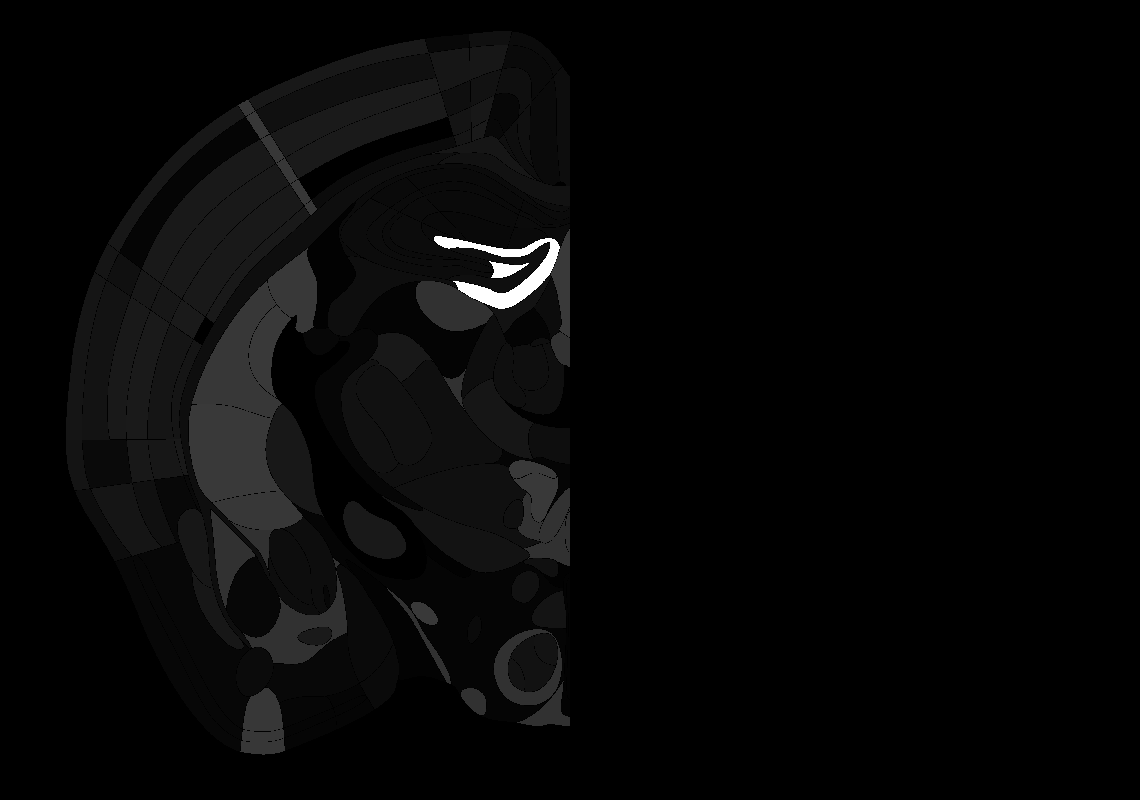

### 59_AP-1.5.tif

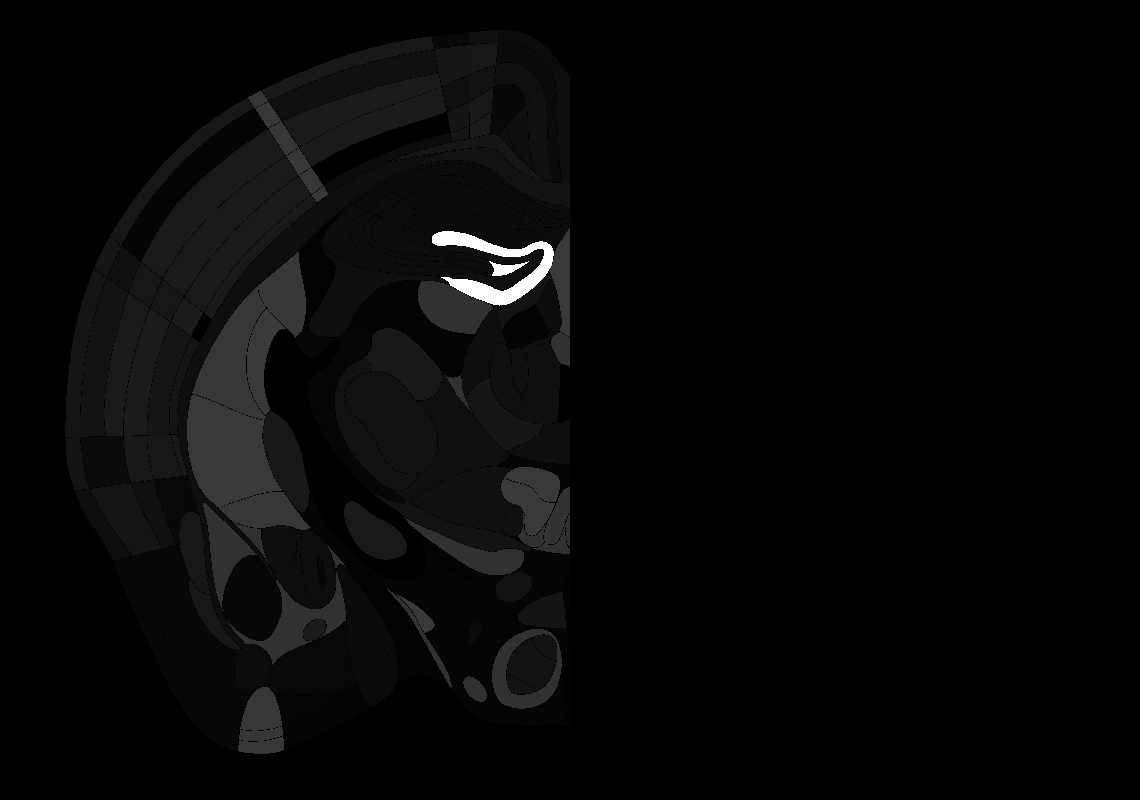

### 60_AP-1.6.tif

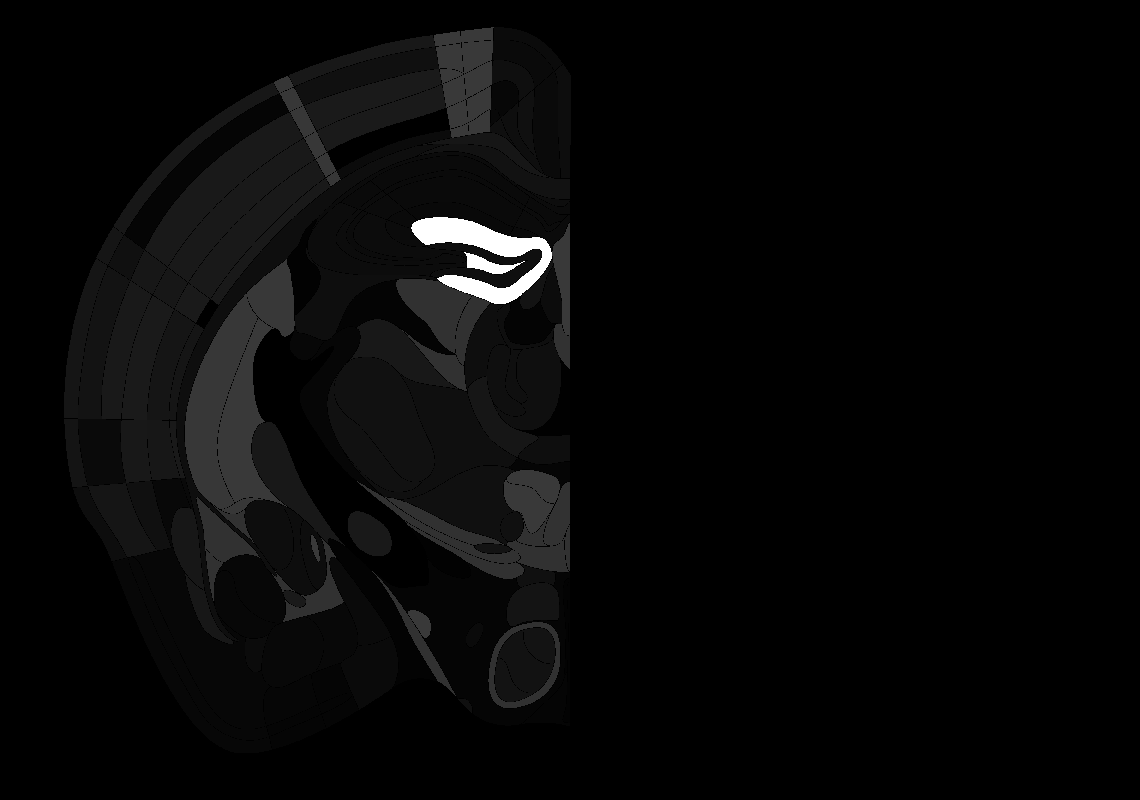

### 62_AP-1.8.tif

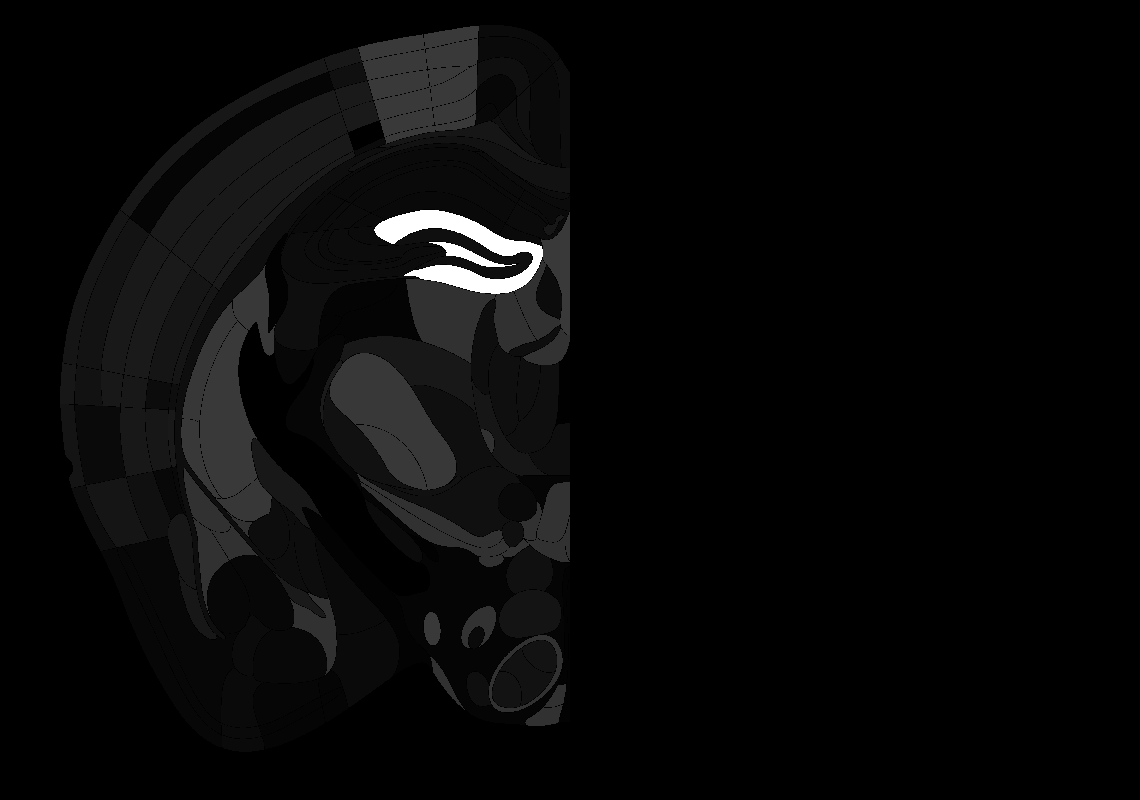

### 63_AP-1.9.tif

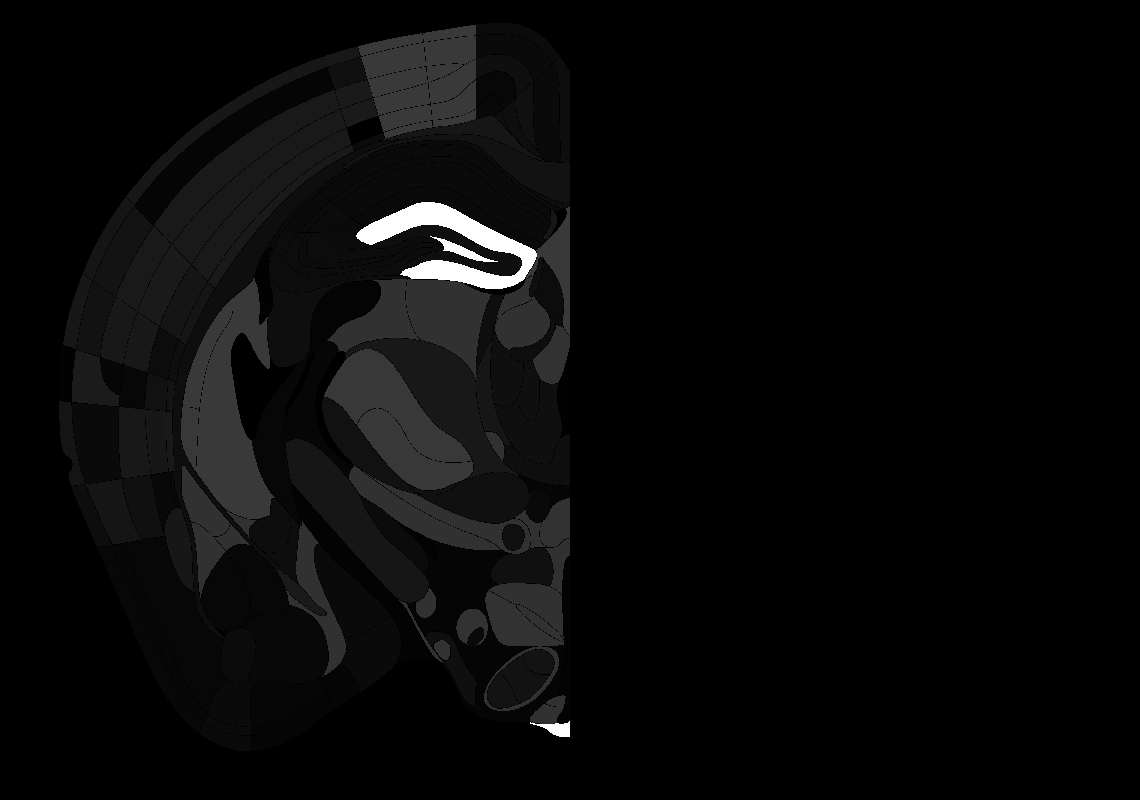

### 64_AP-2.0.tif

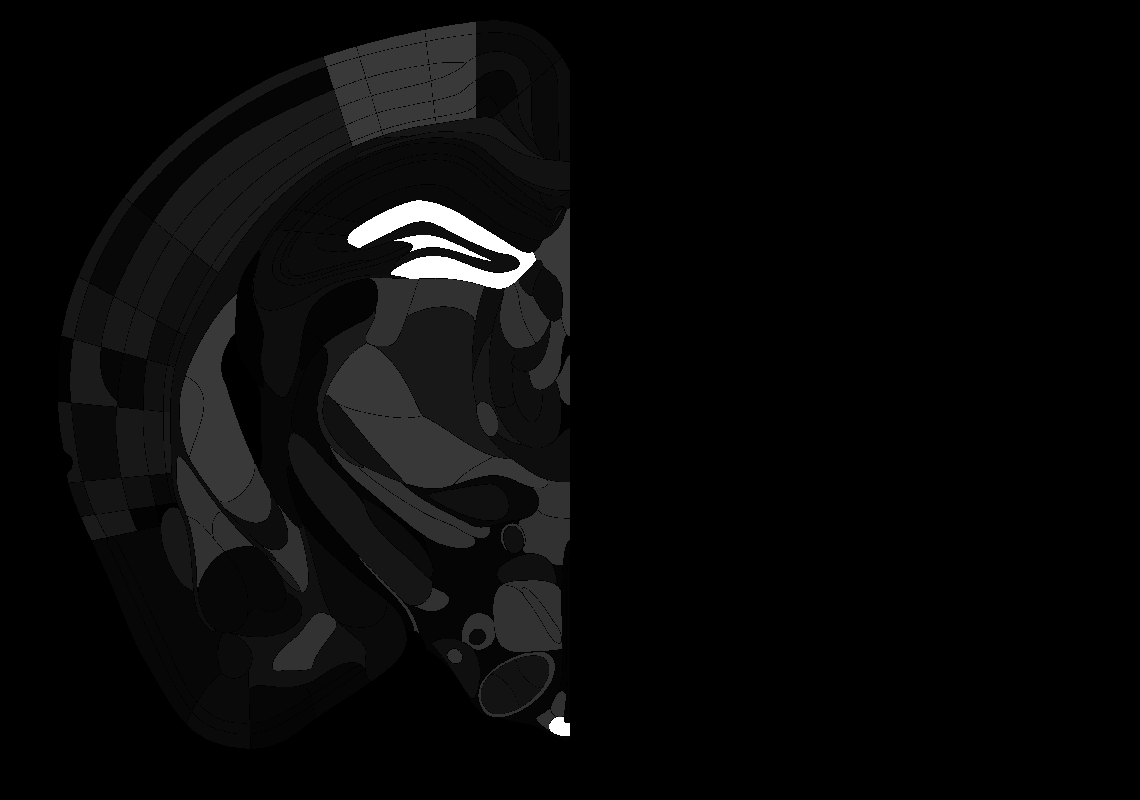

### 65_AP-2.1.tif

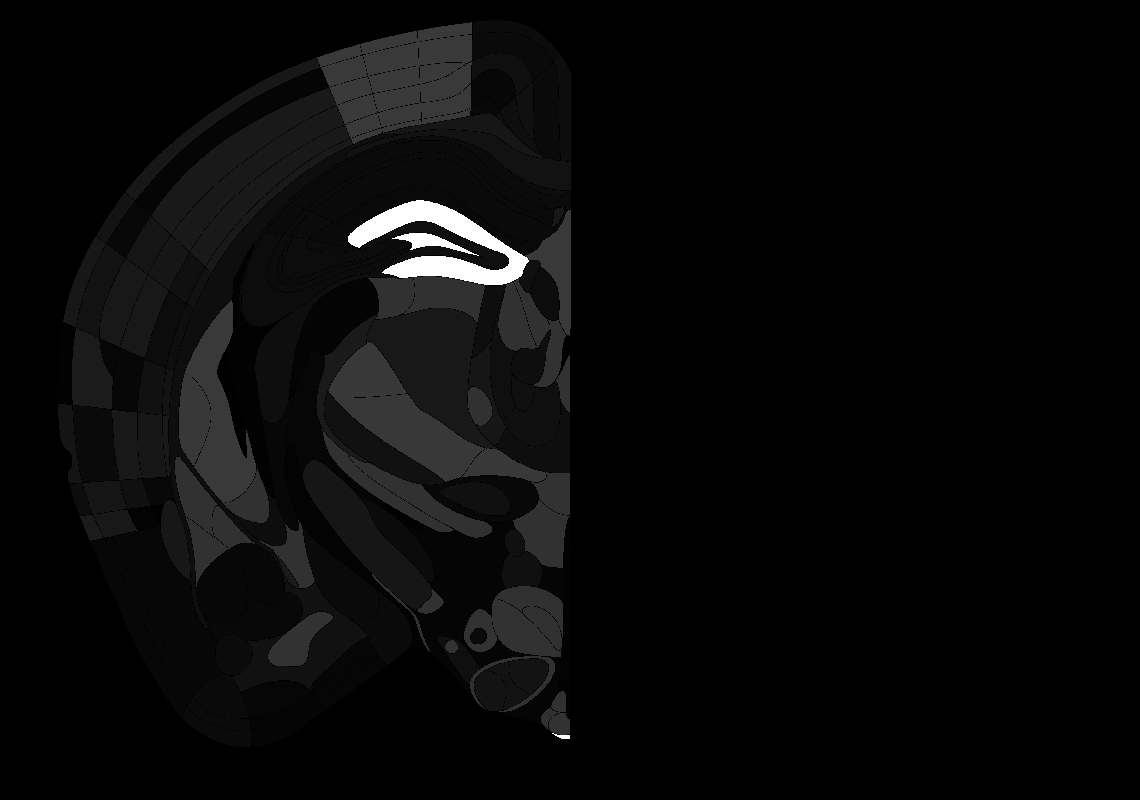

### 66_AP-2.2.tif

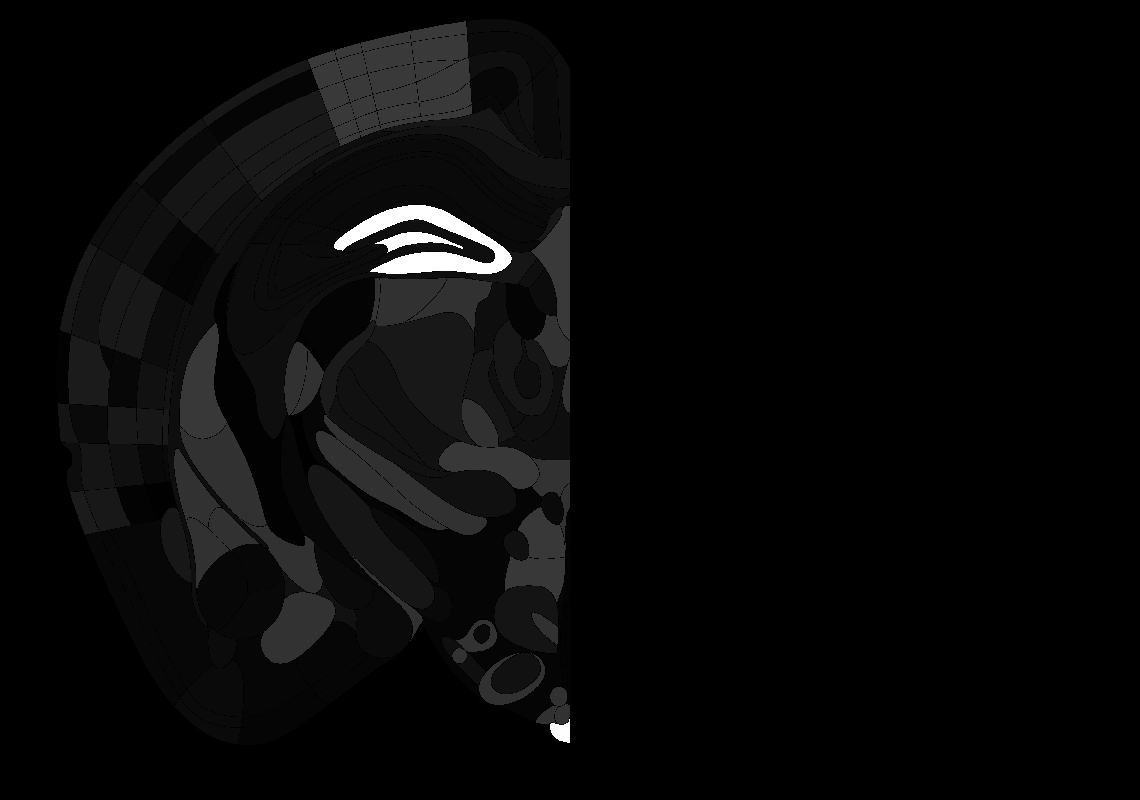

### 67_AP-2.3.tif

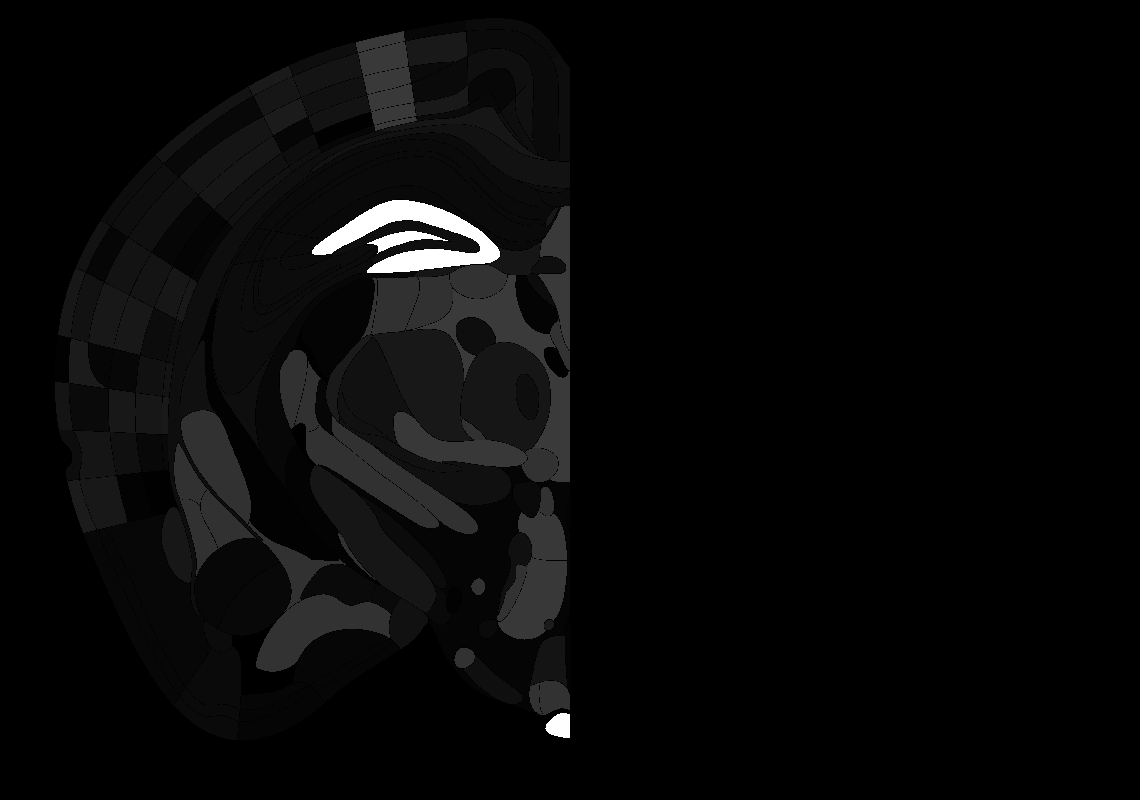

### 68_AP-2.4.tif

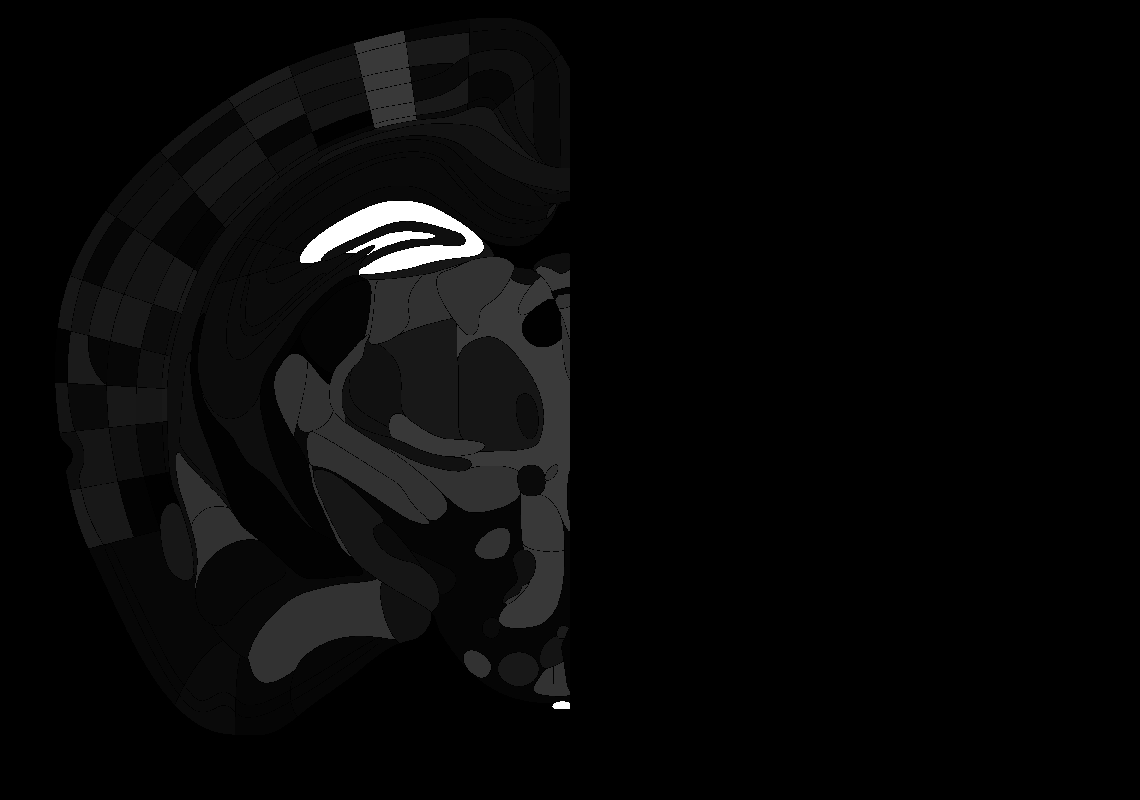

### 69_AP-2.5.tif

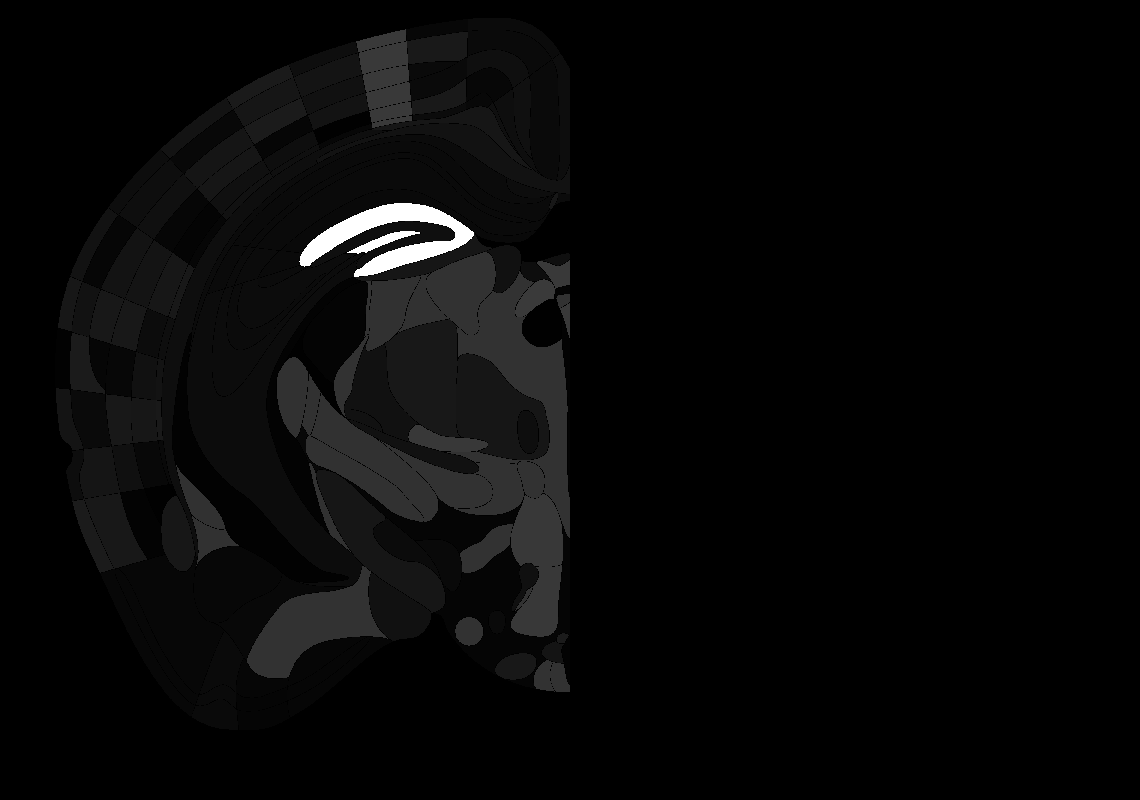

### 70_AP-2.6.tif

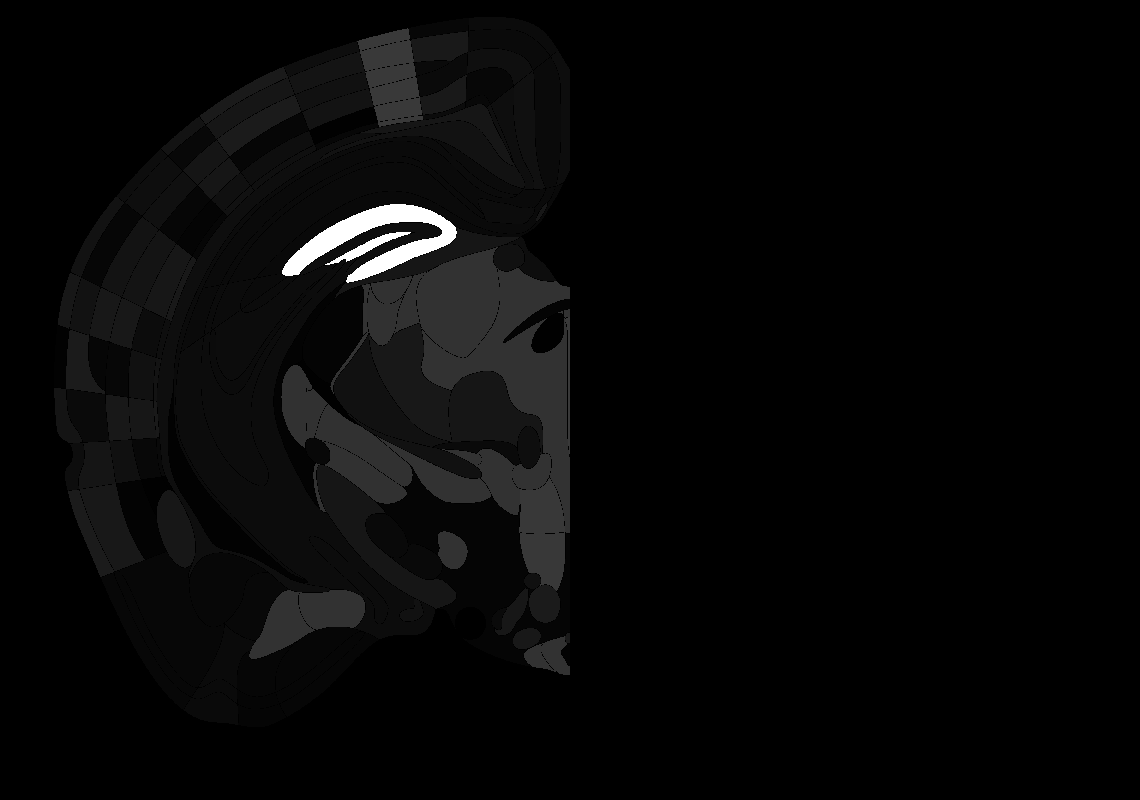

### 71_AP-2.7.tif

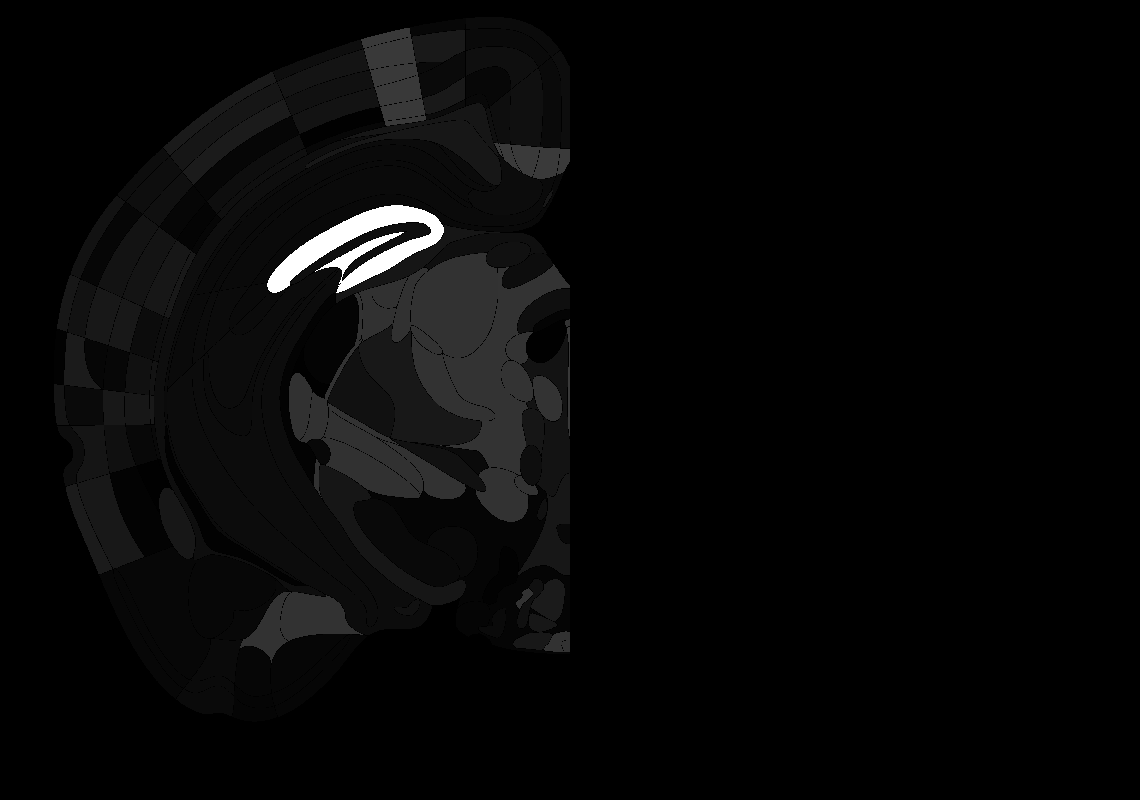

### 72_AP-2.8.tif

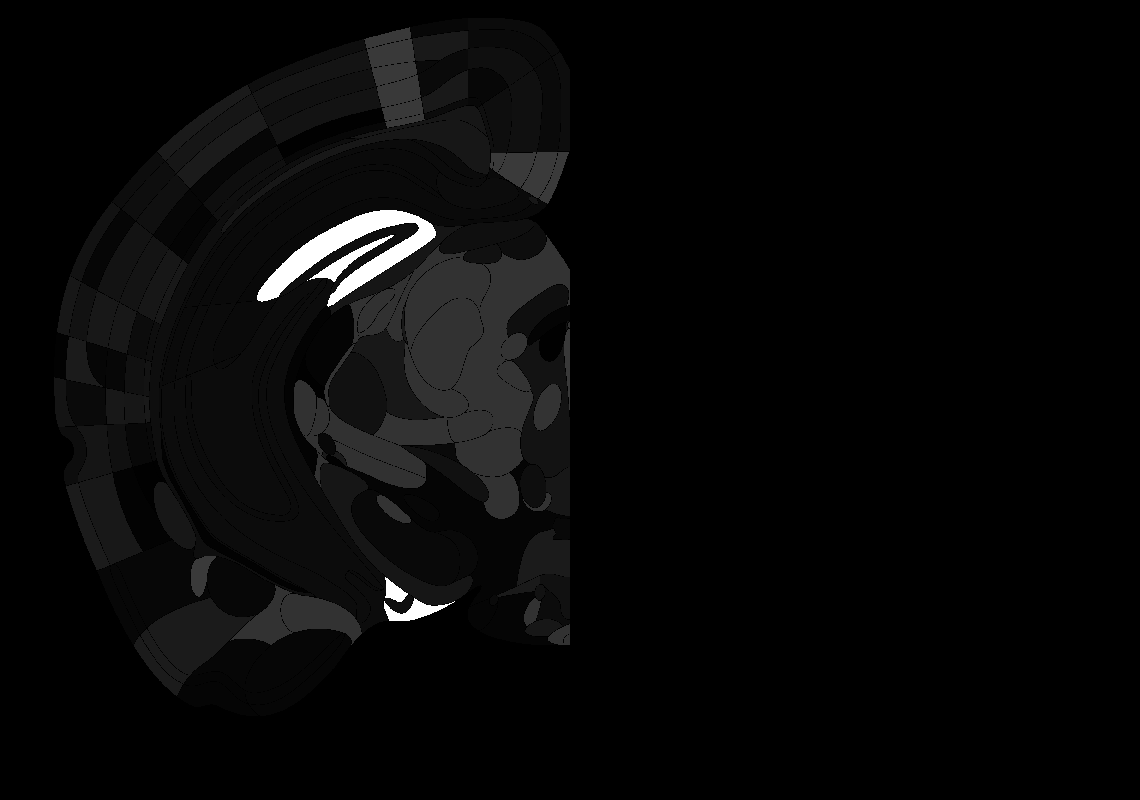

### 73_AP-2.9.tif

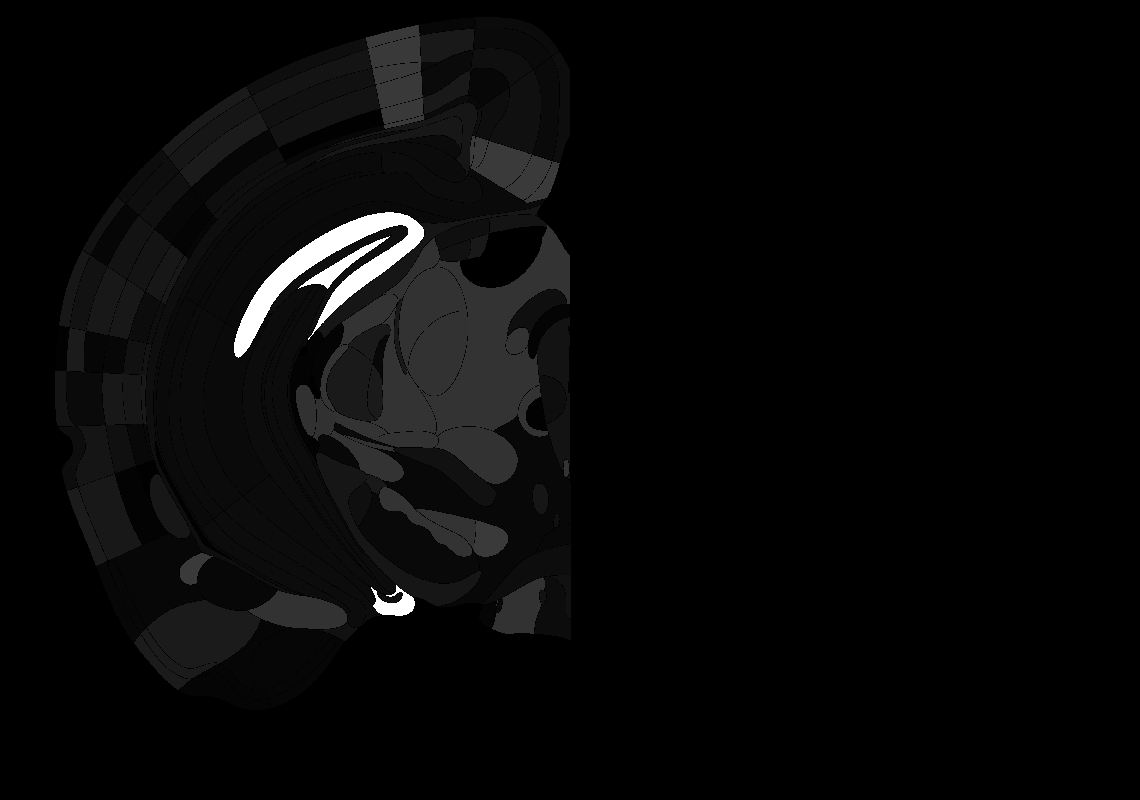

### 74_AP-3.0.tif

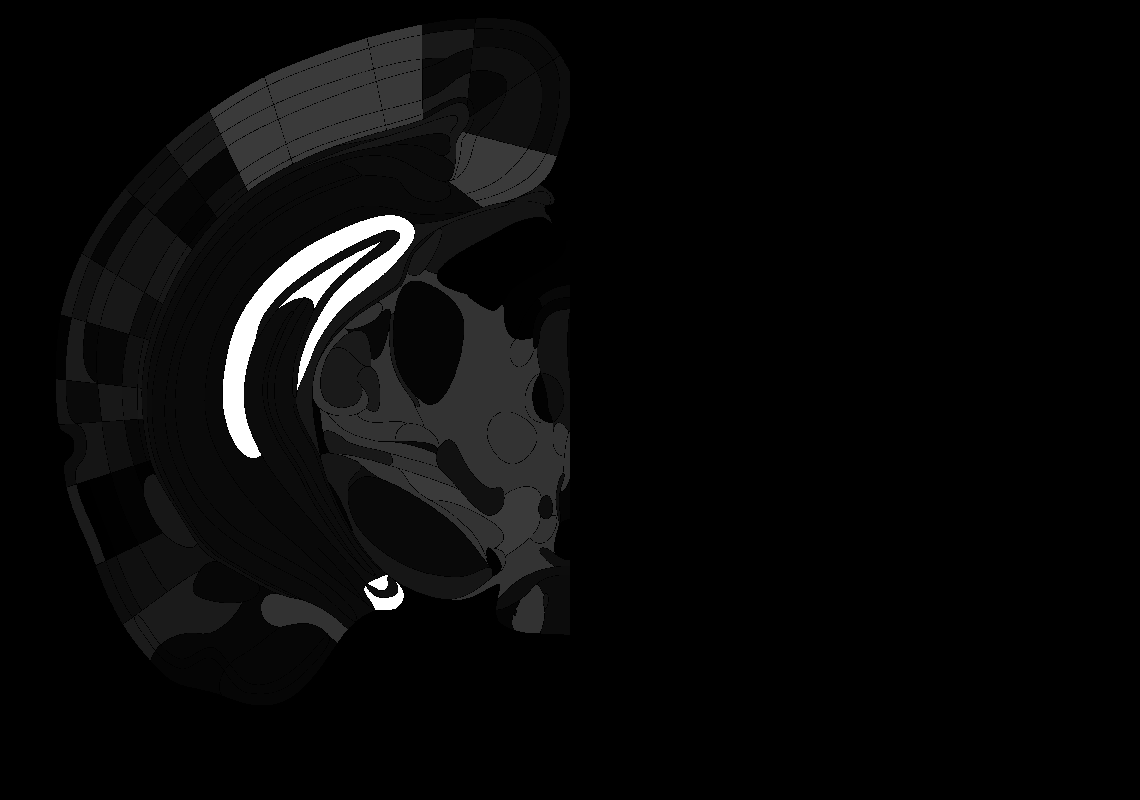

### 76_AP-3.2.tif

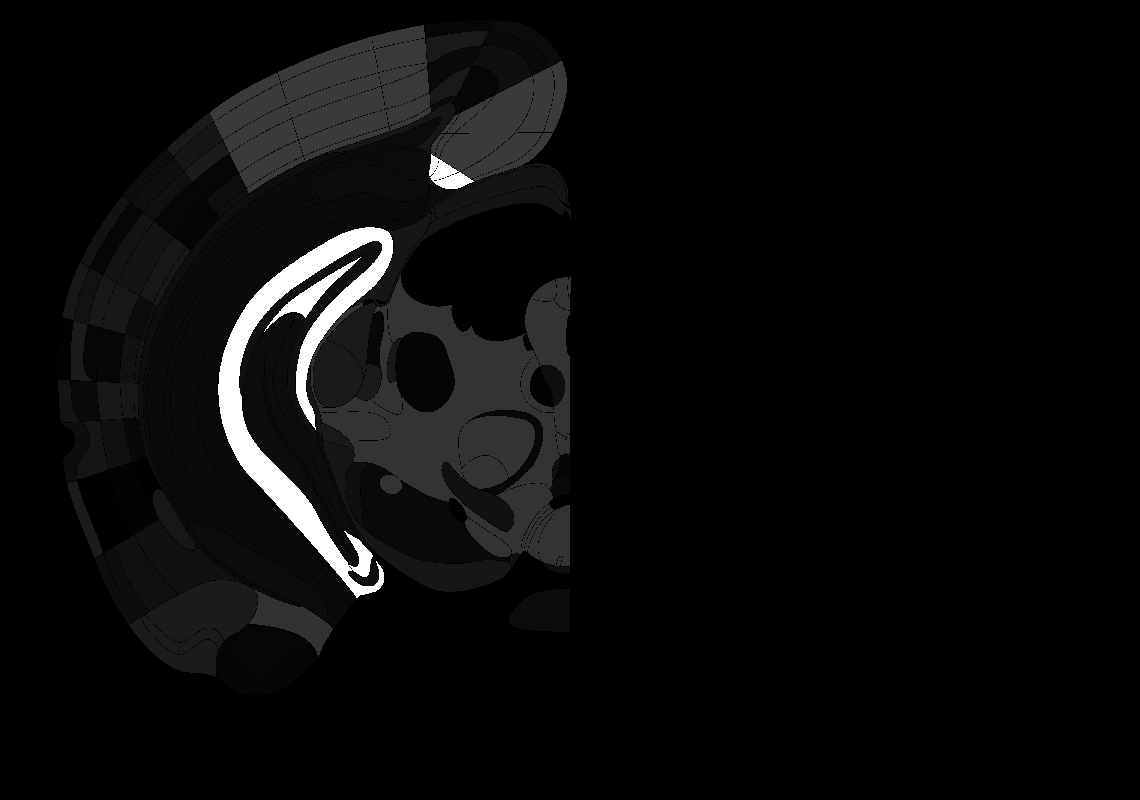

### 77_AP-3.3.tif

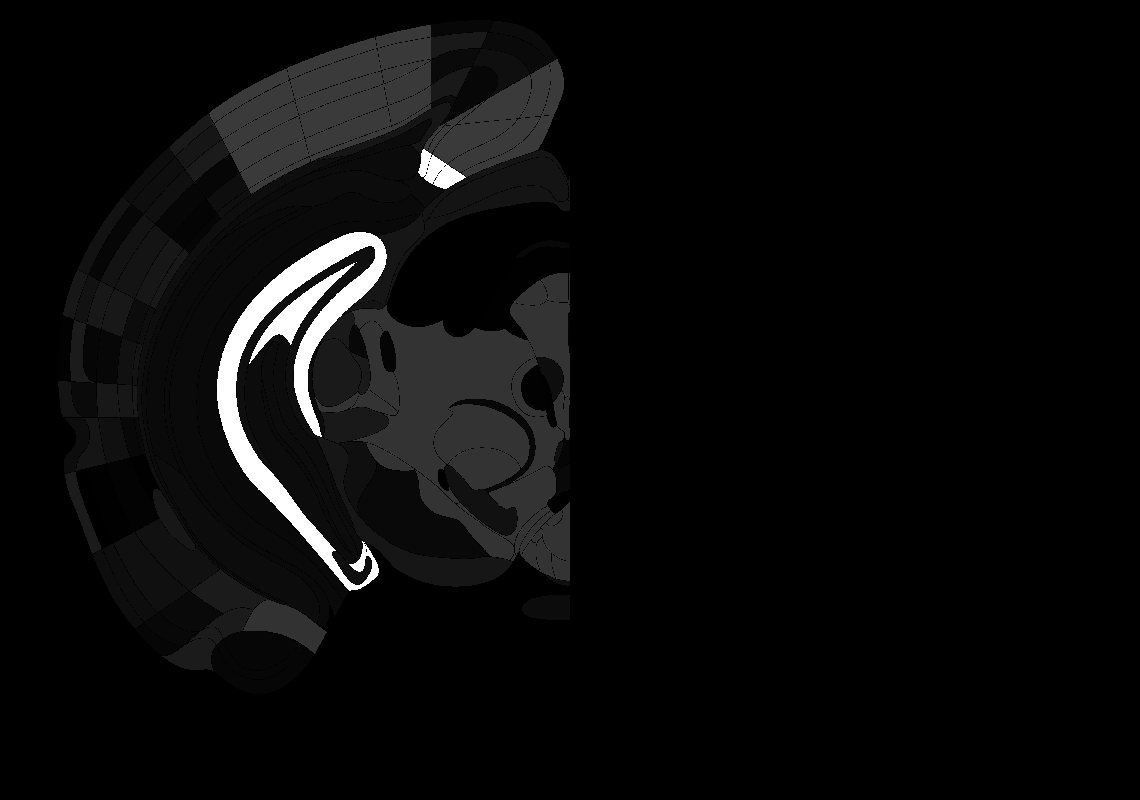

### 79_AP-3.5.tif

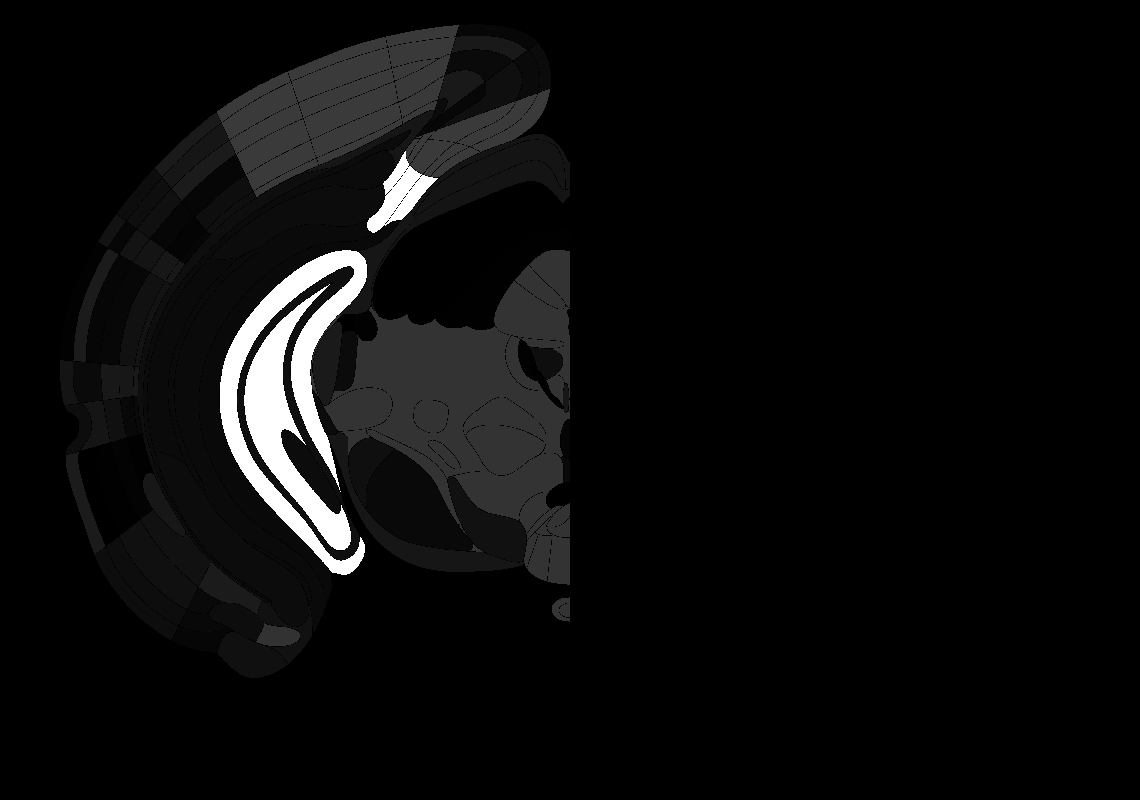

### 80_AP-3.6.tif

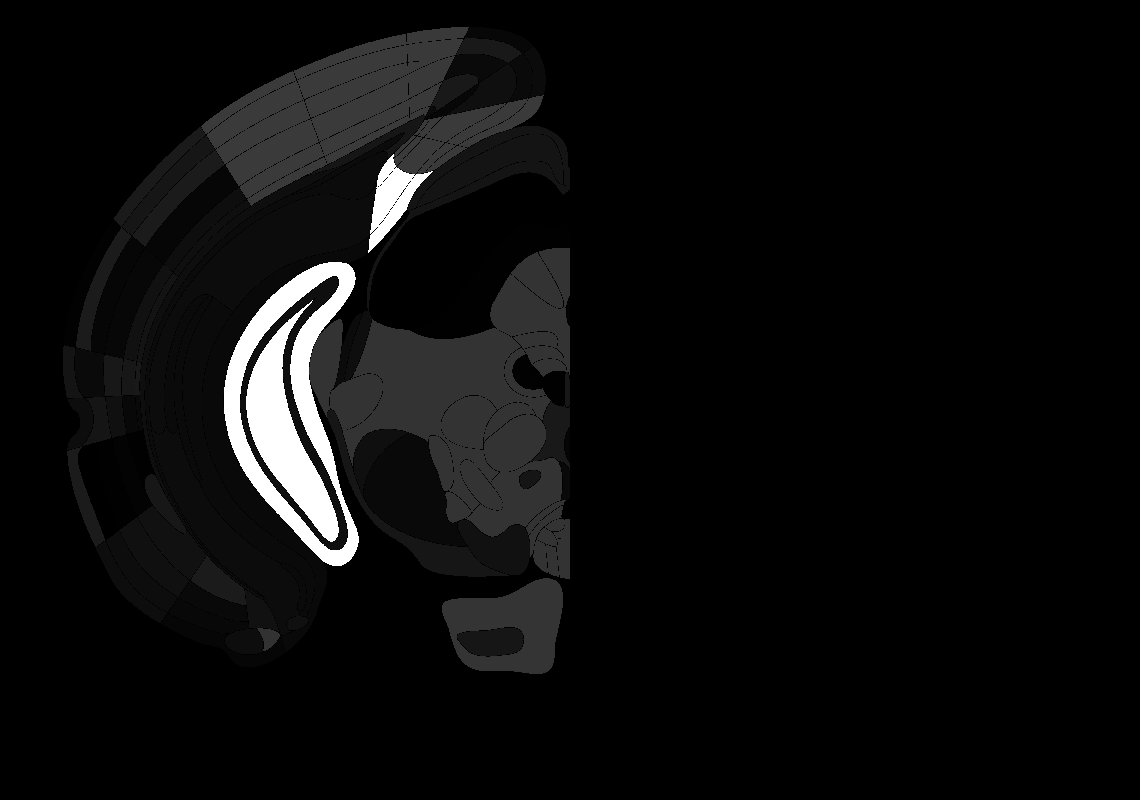

### 81_AP-3.7.tif

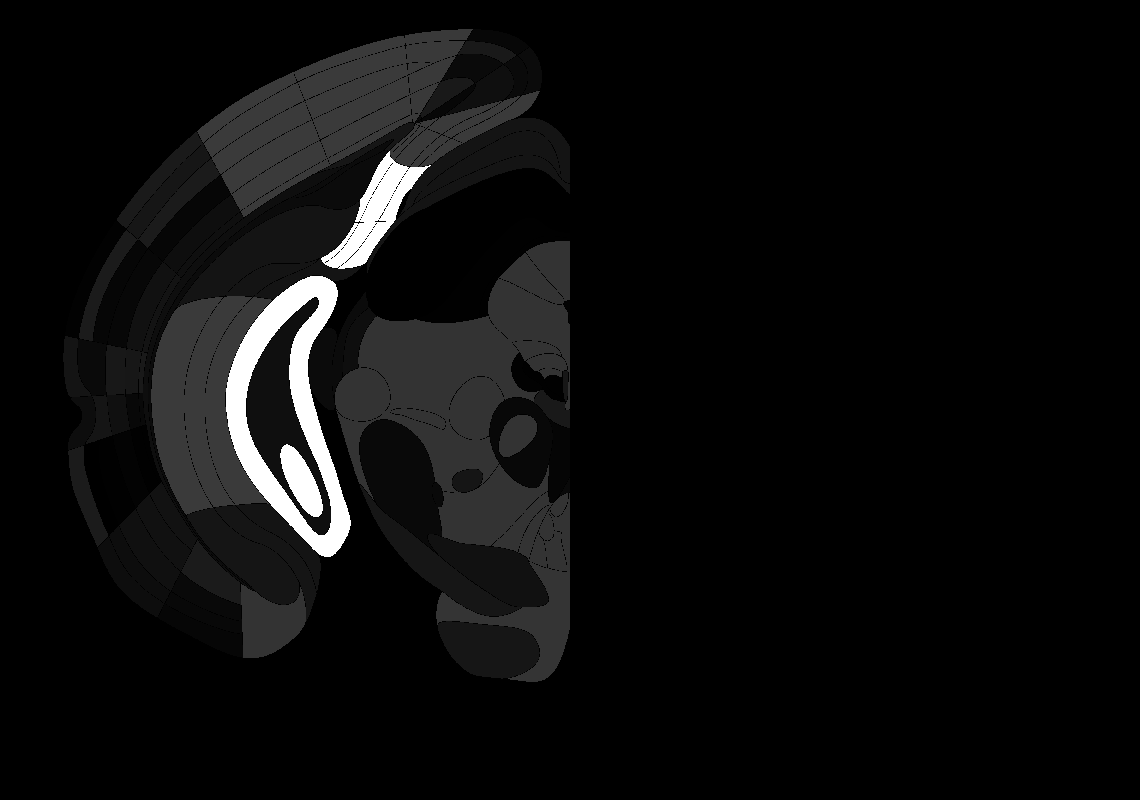

### 82_AP-3.8.tif

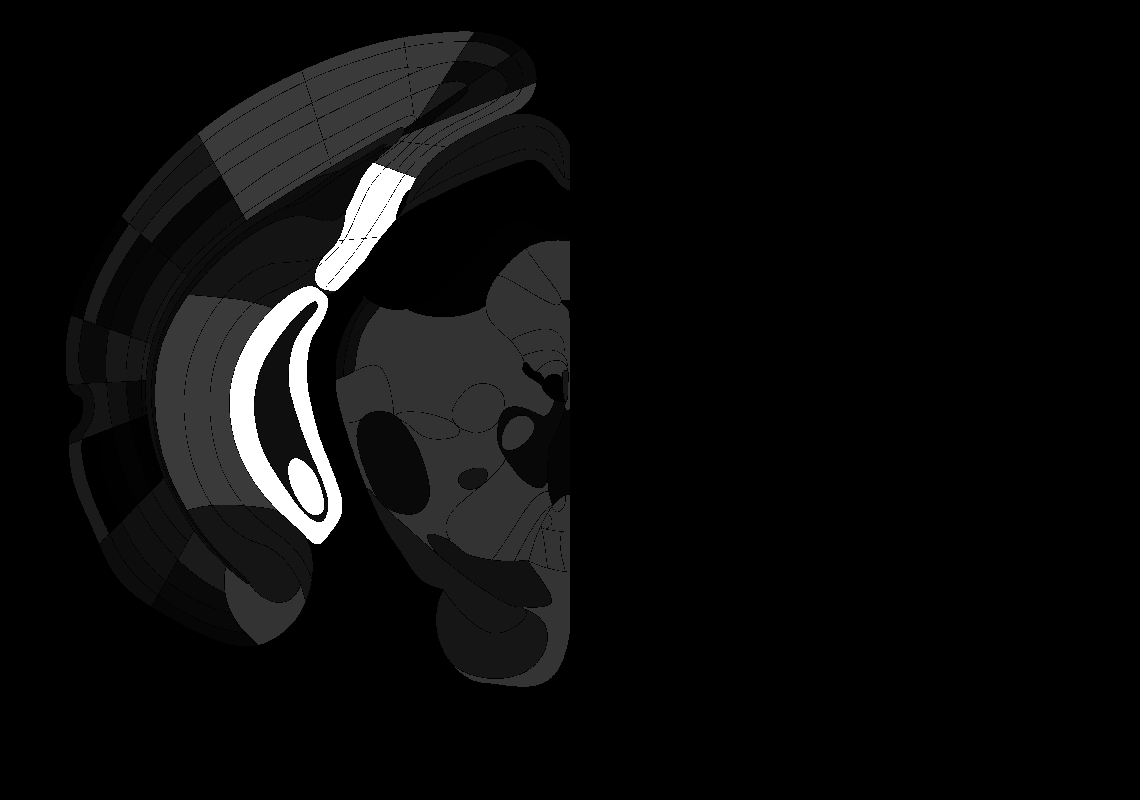

### 83_AP-3.9.tif

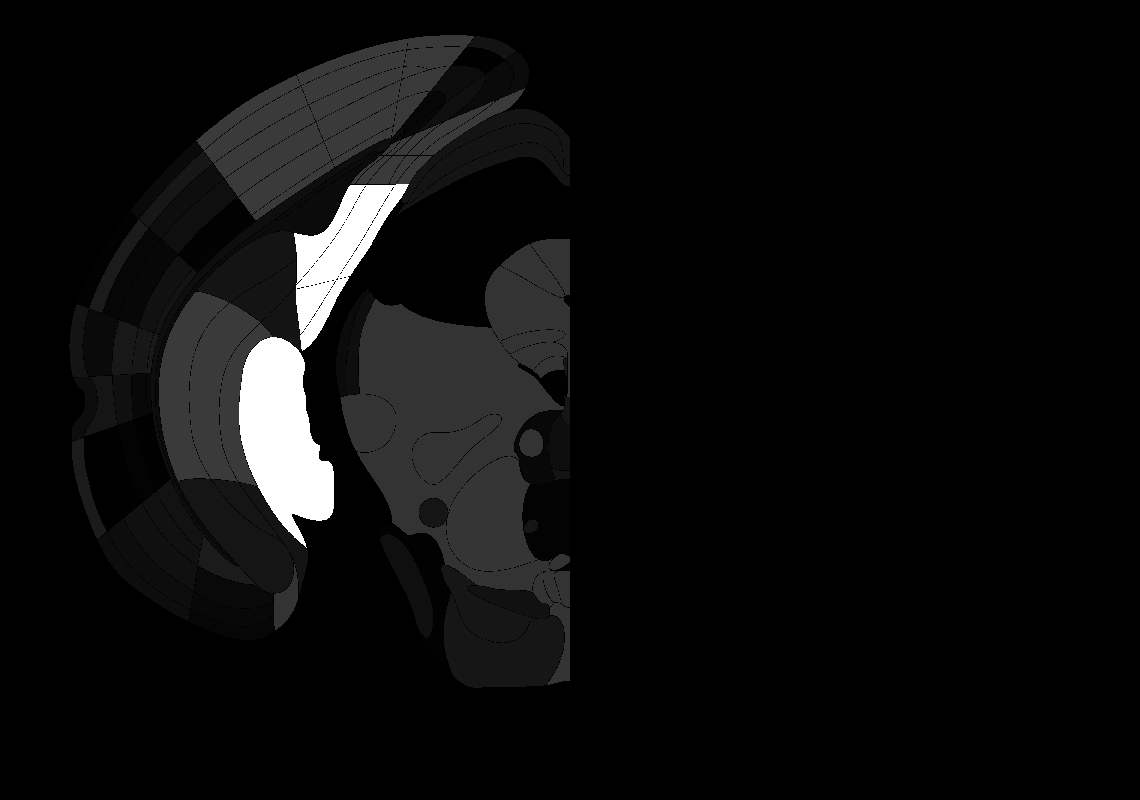

### 84_AP-4.0.tif

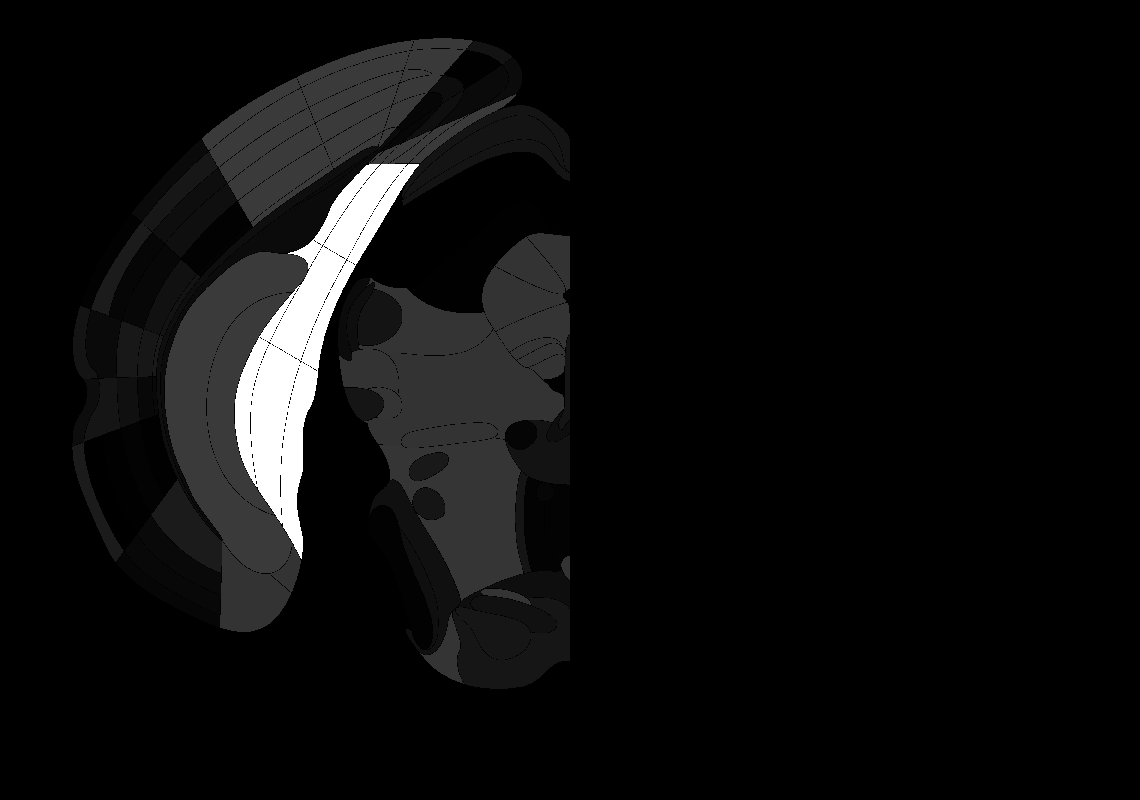

### 86_AP-4.2.tif

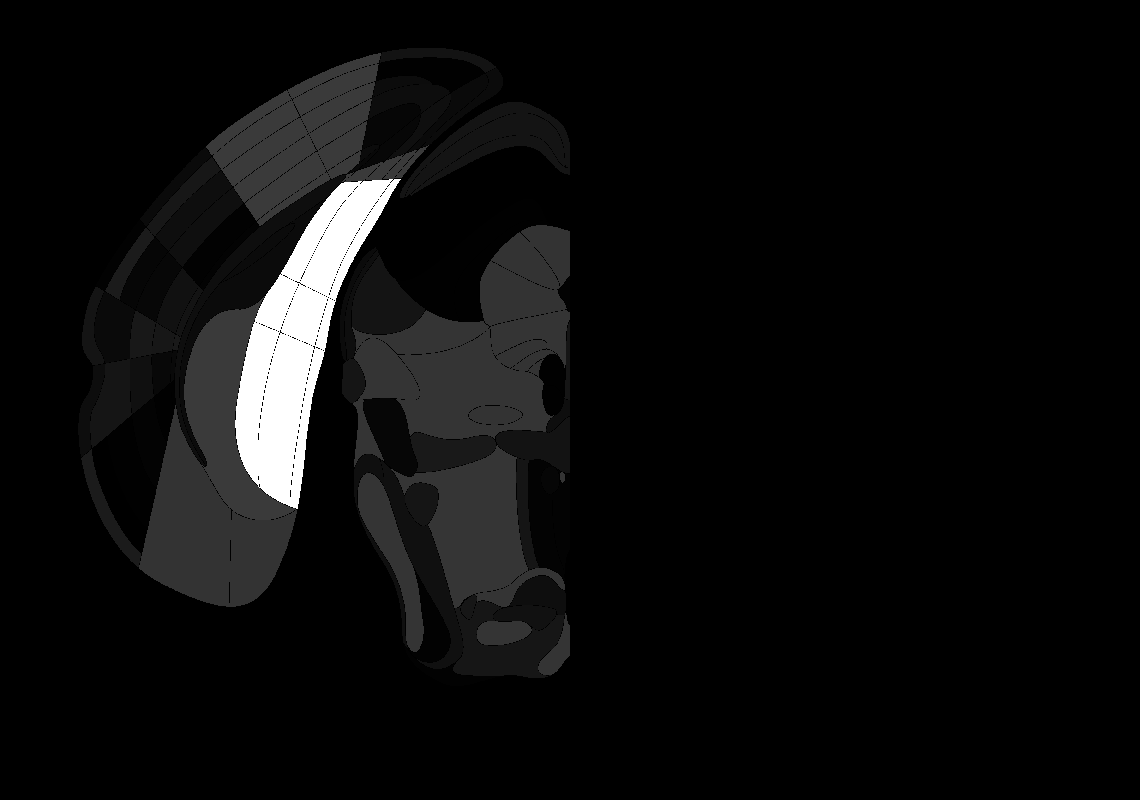

### 87_AP-4.3.tif

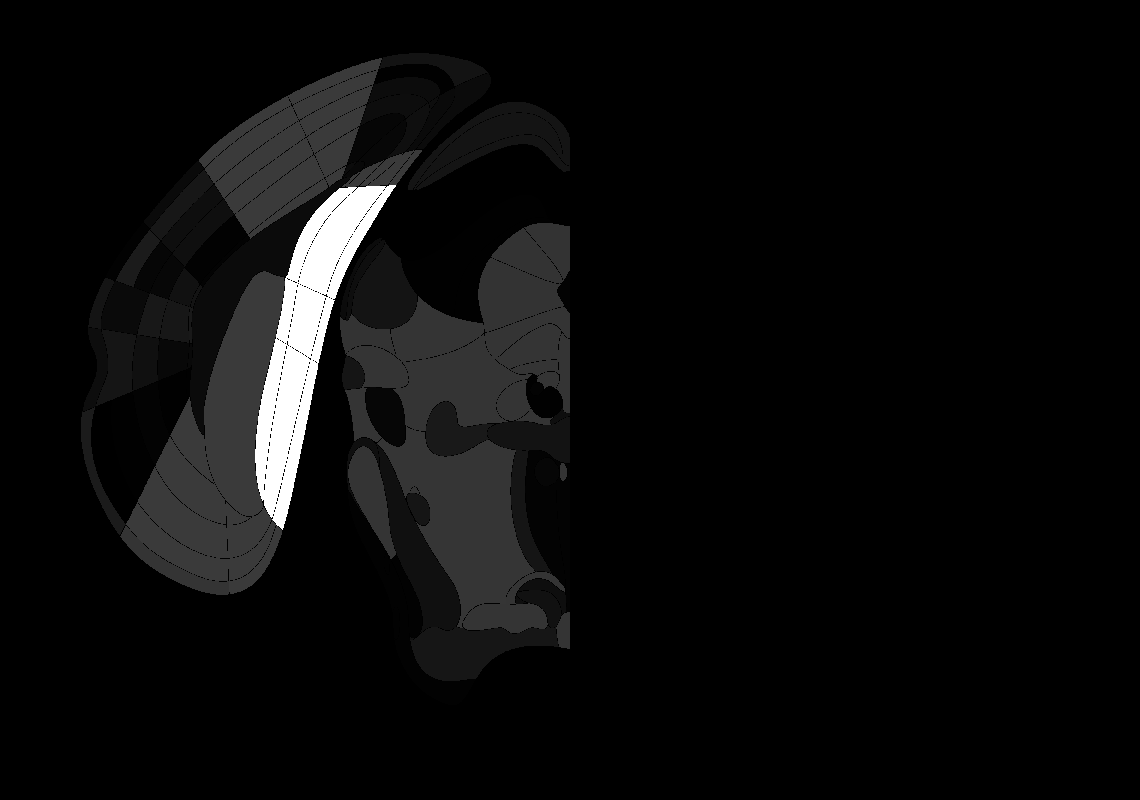

### 89_AP-4.5.tif

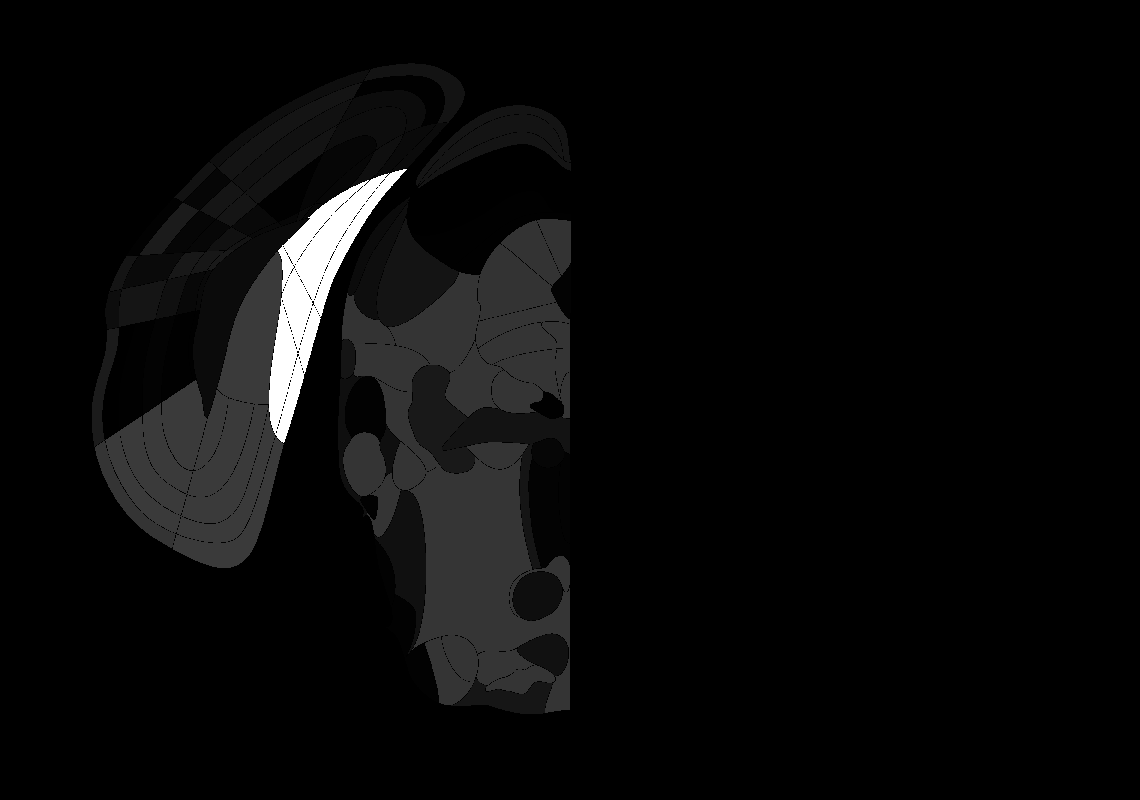

### 90_AP-4.6.tif

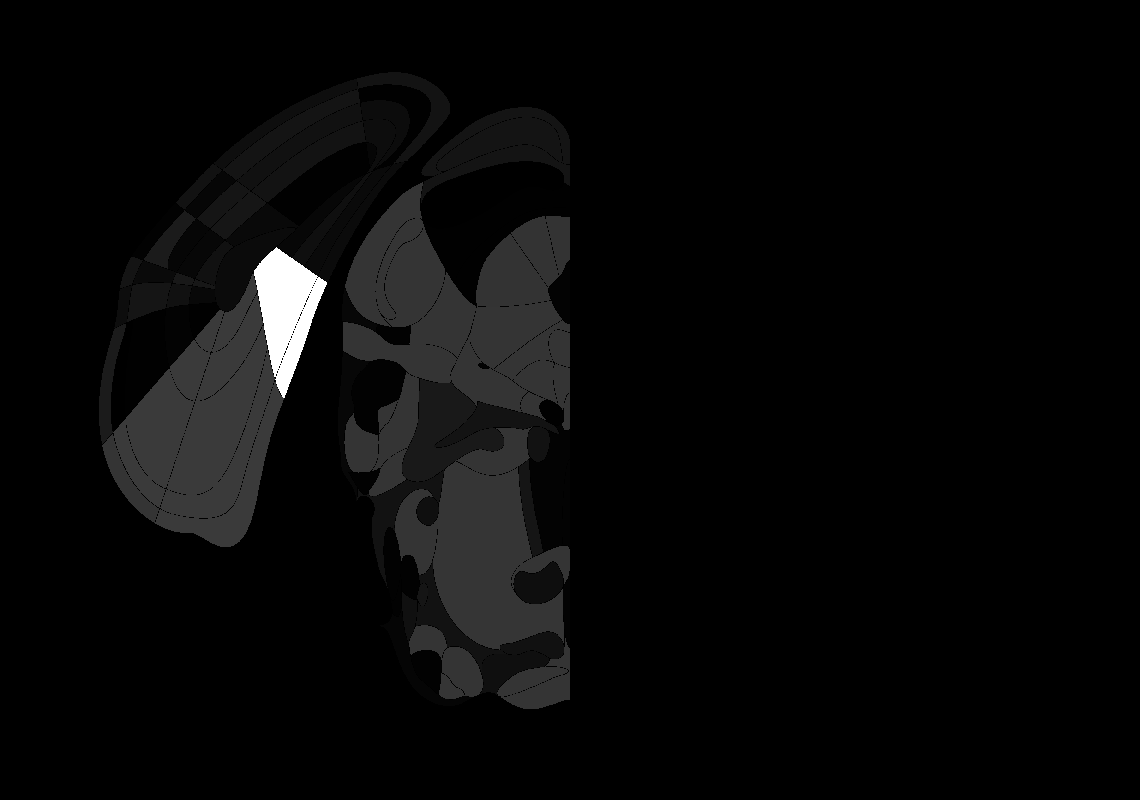

### 93_AP-4.9.tif

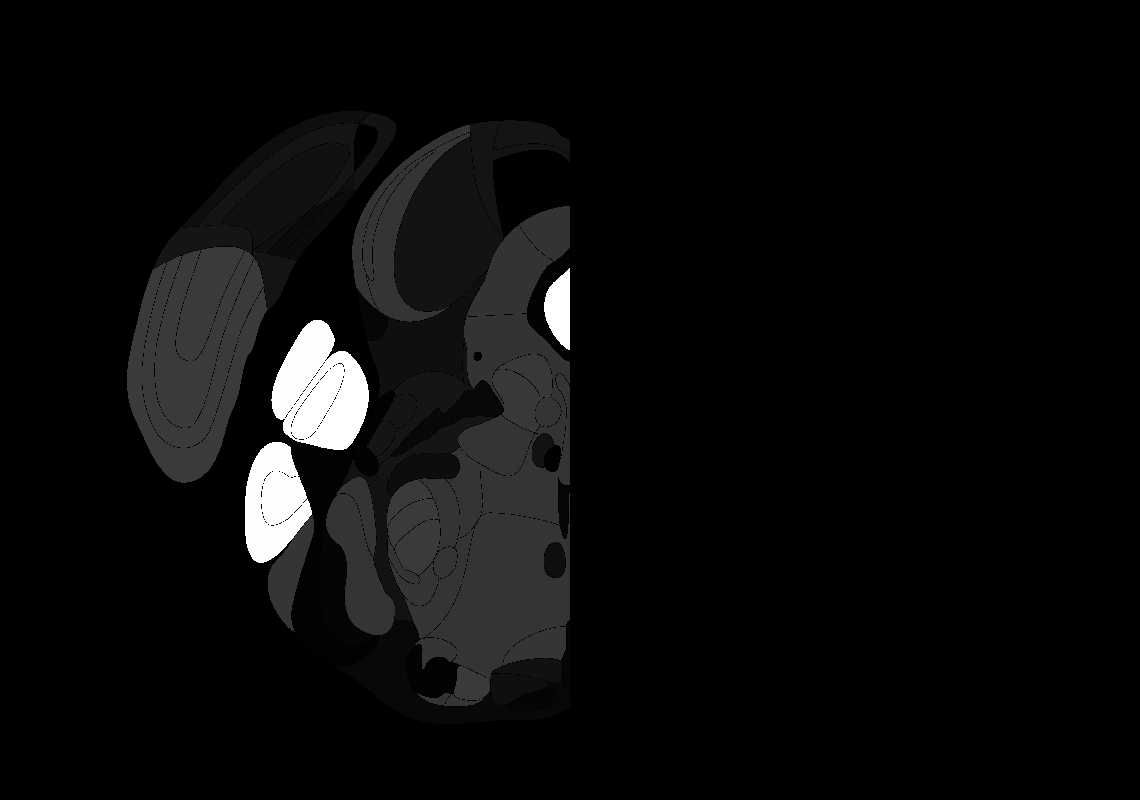

### 94_AP-5.0.tif

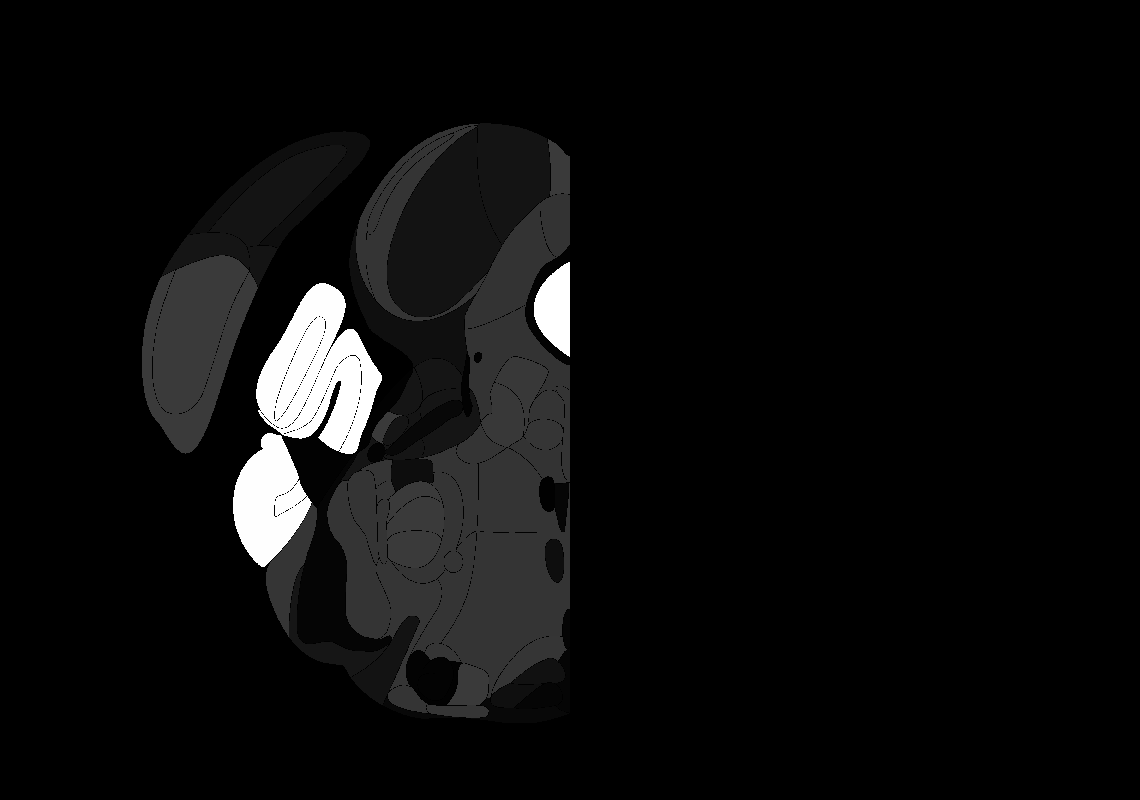

### 95_AP-5.1.tif

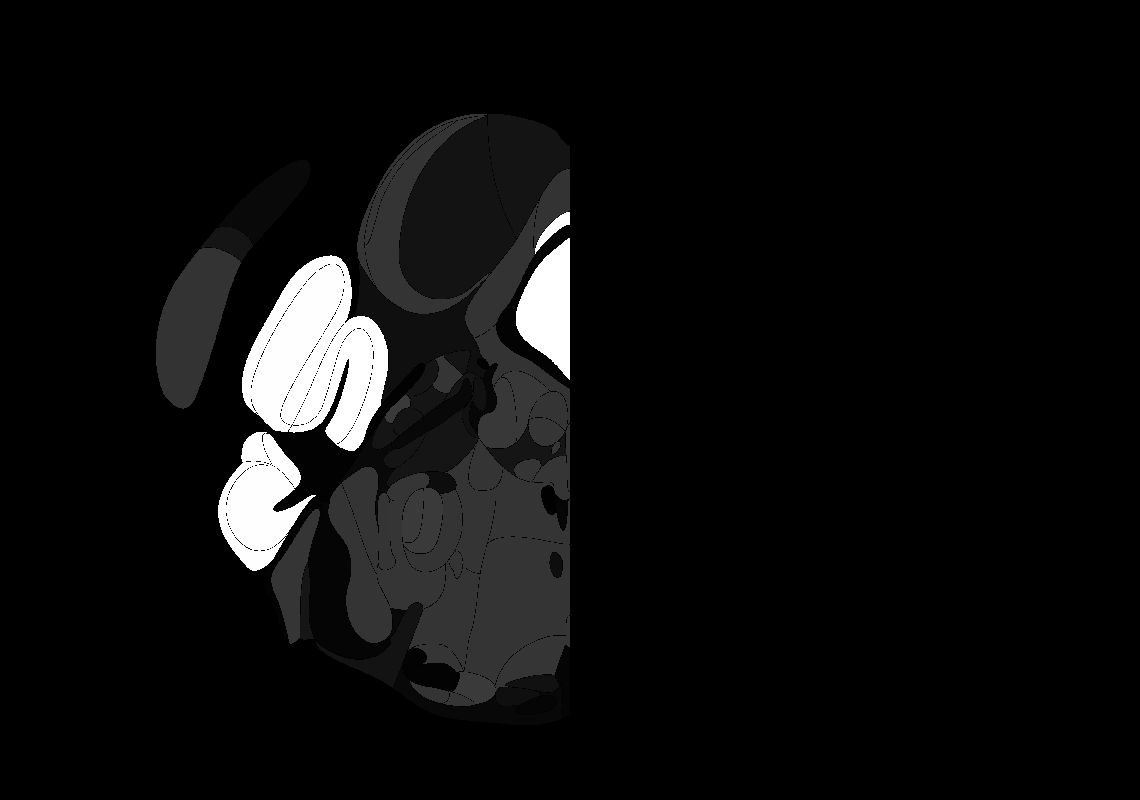

### 97_AP-5.3.tif

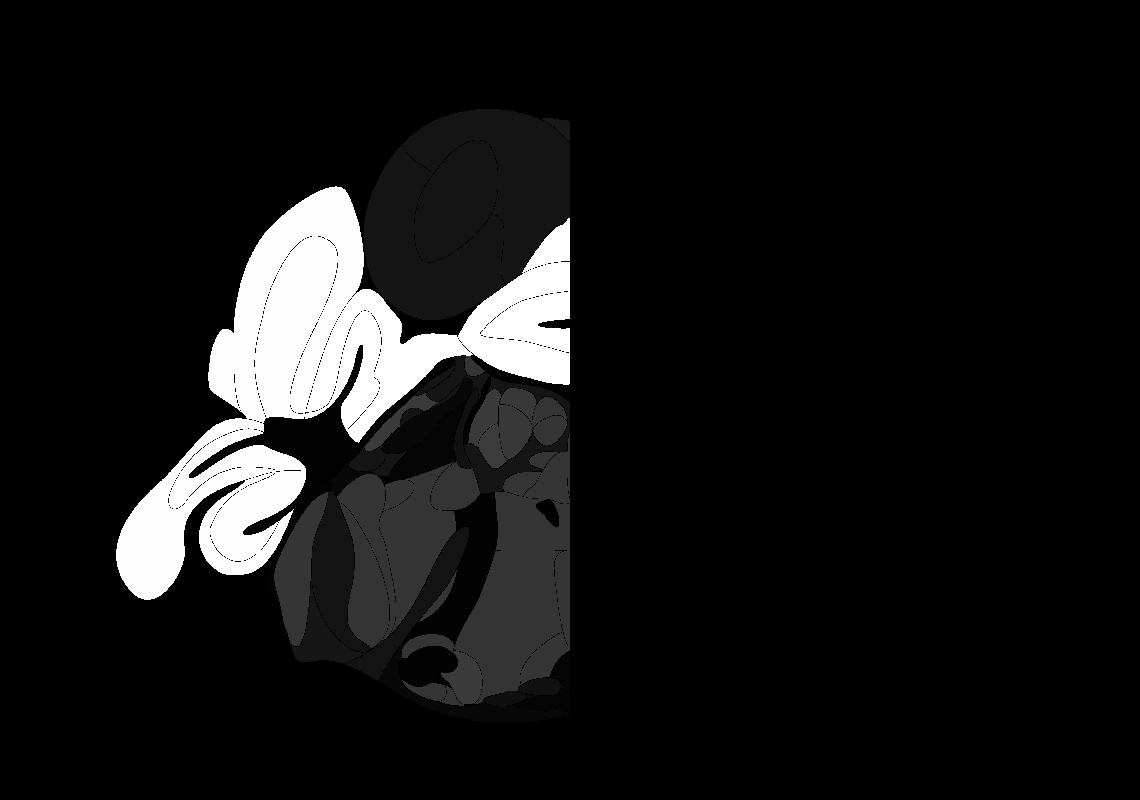

### 98_AP-5.4.tif

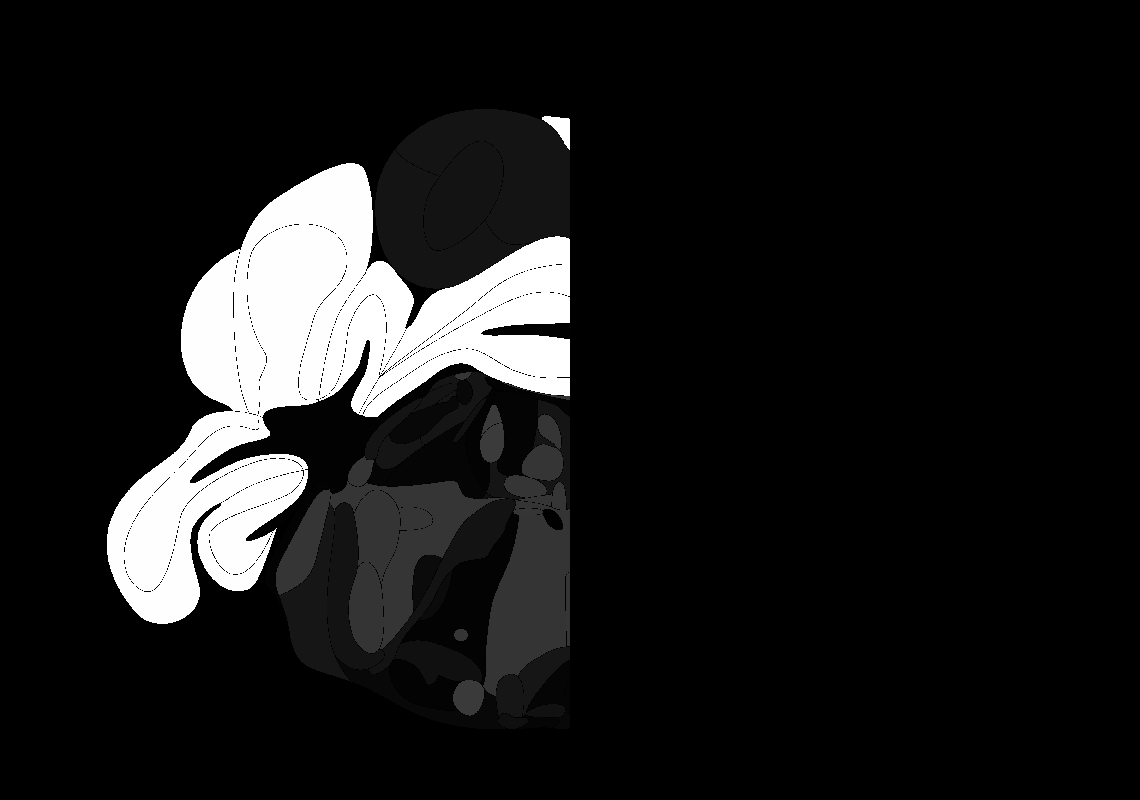

### 99_AP-5.5.tif

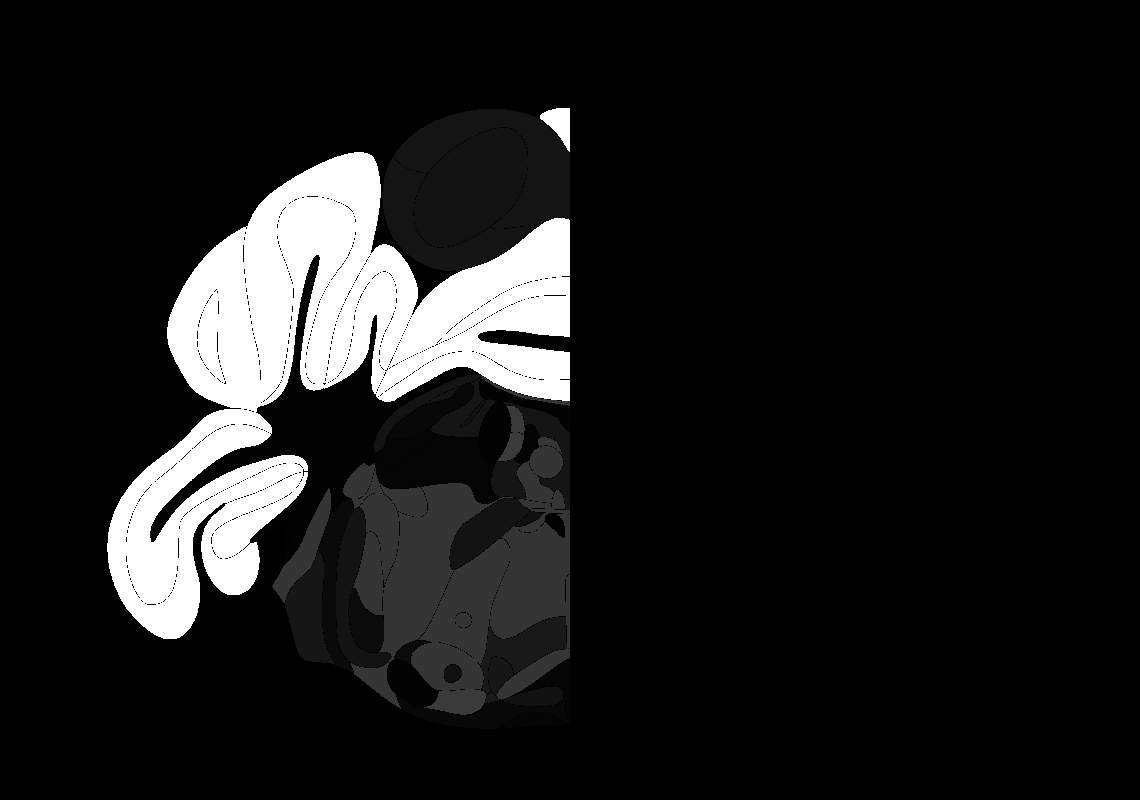

### 100_AP-5.6.tif

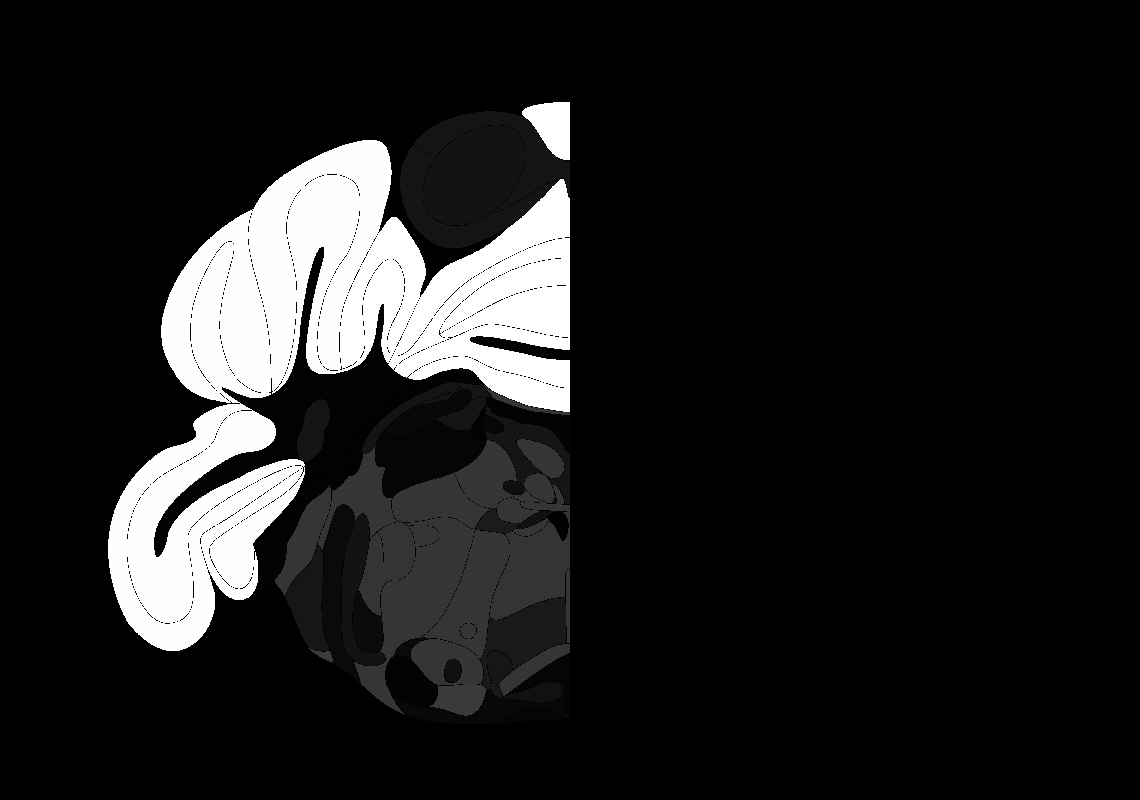

### 101_AP-5.7.tif

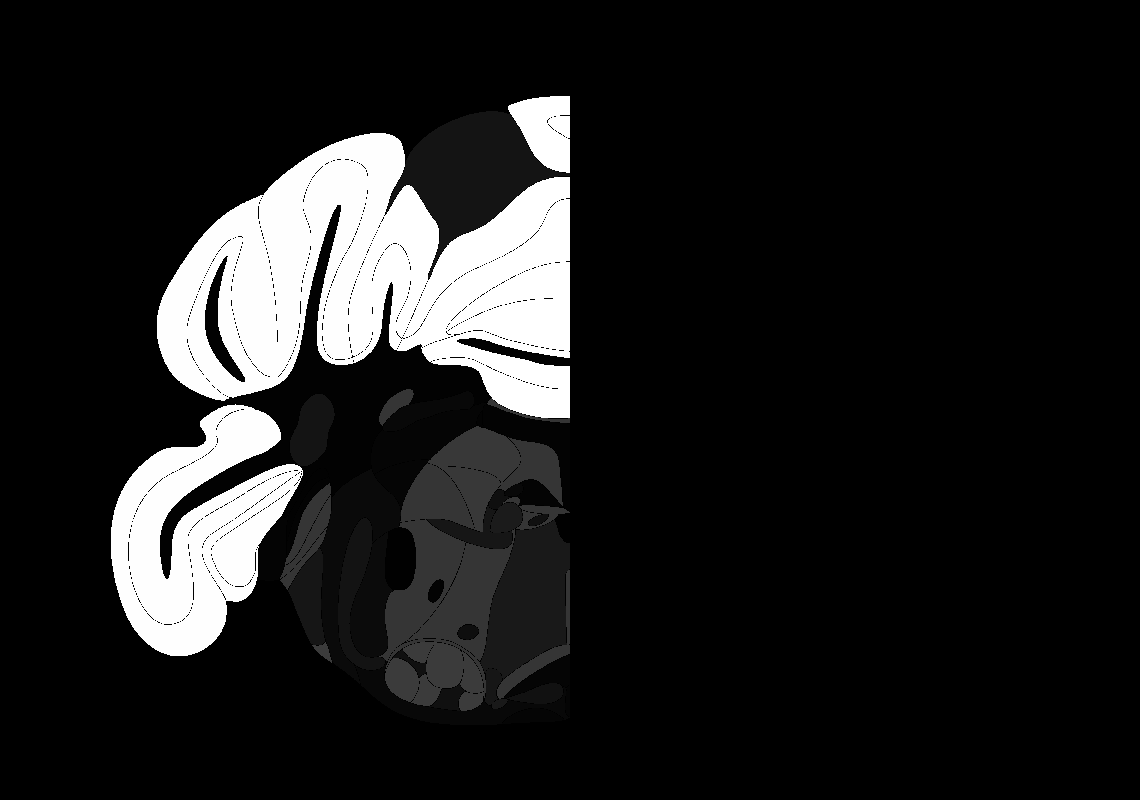

### 102_AP-5.8.tif

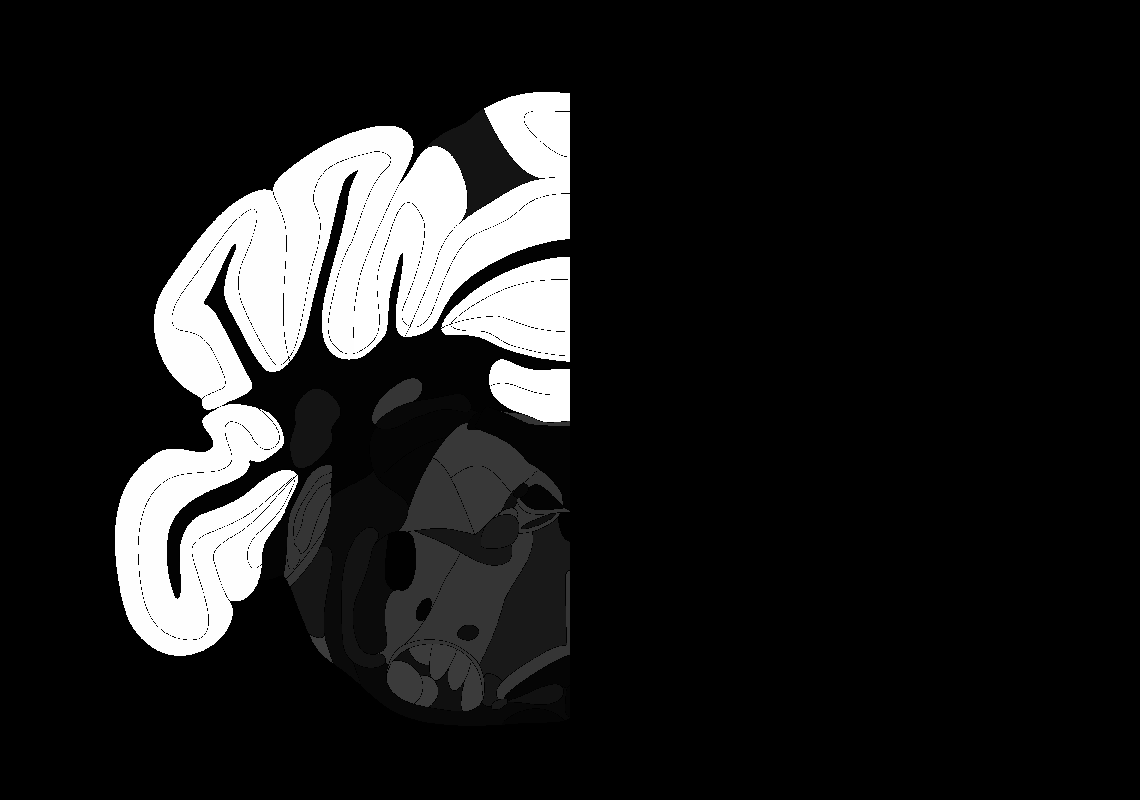

### 103_AP-5.9.tif

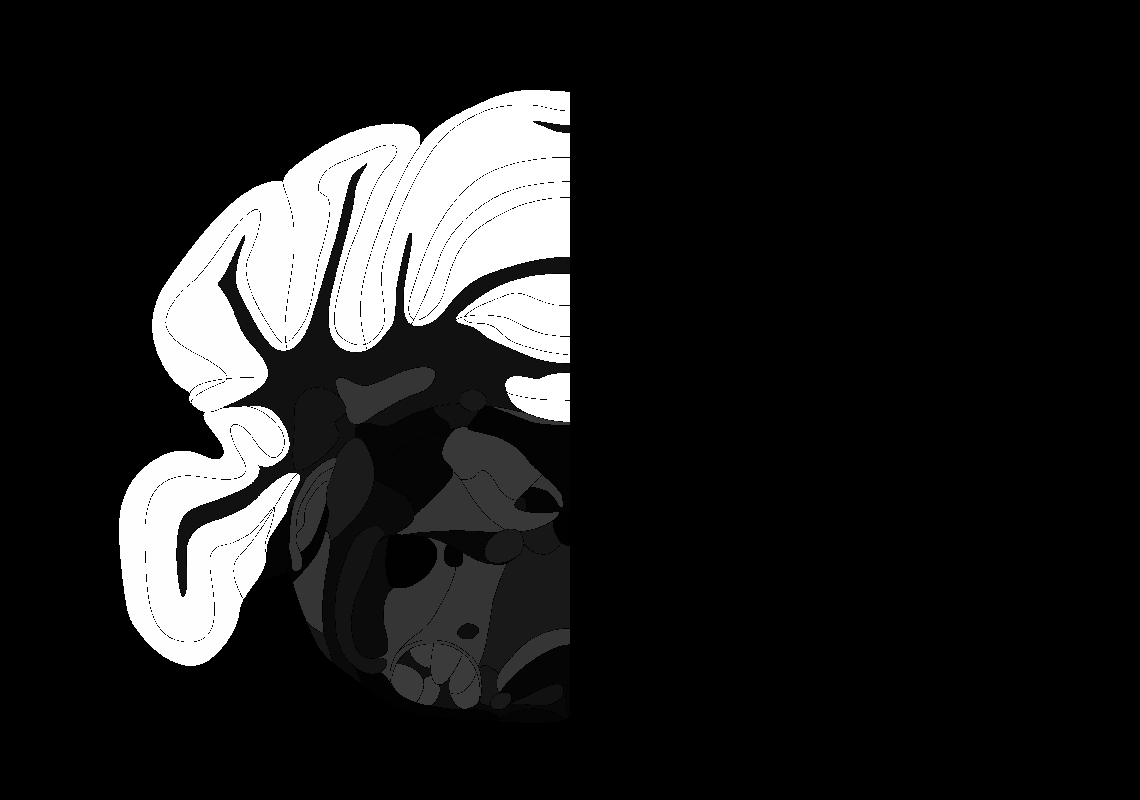

### 104_AP-6.0.tif

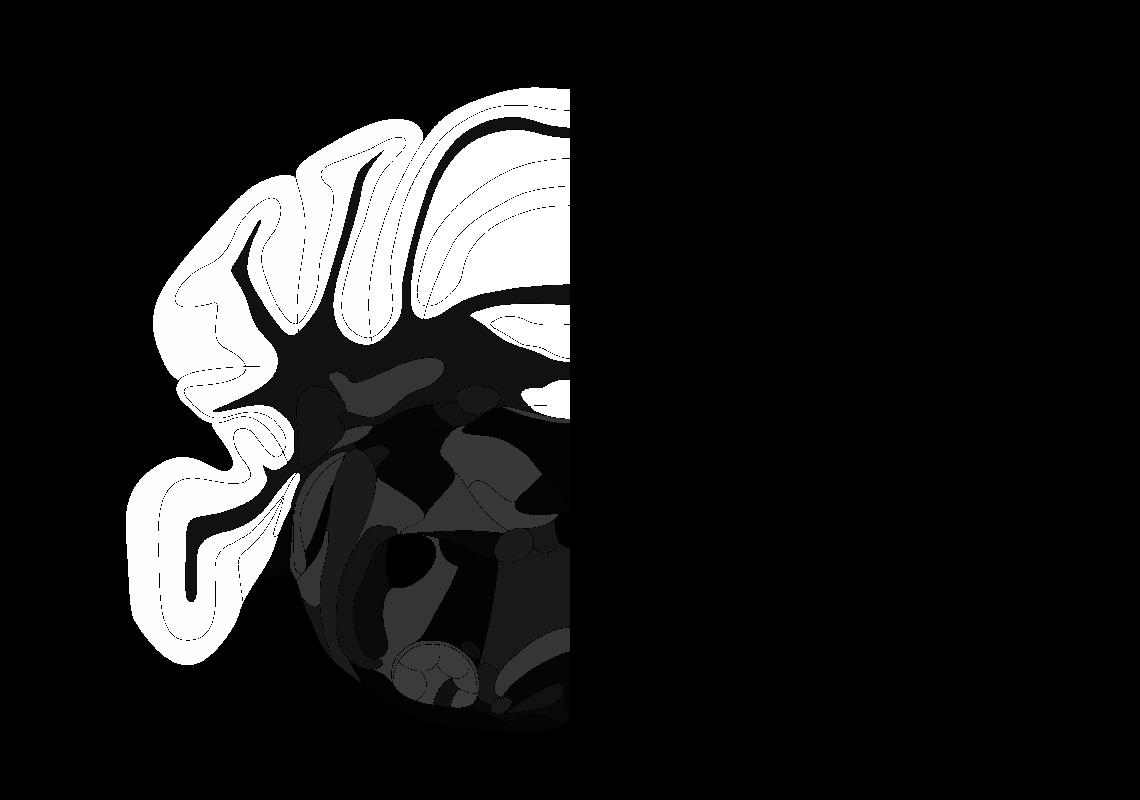

### 105_AP-6.1.tif

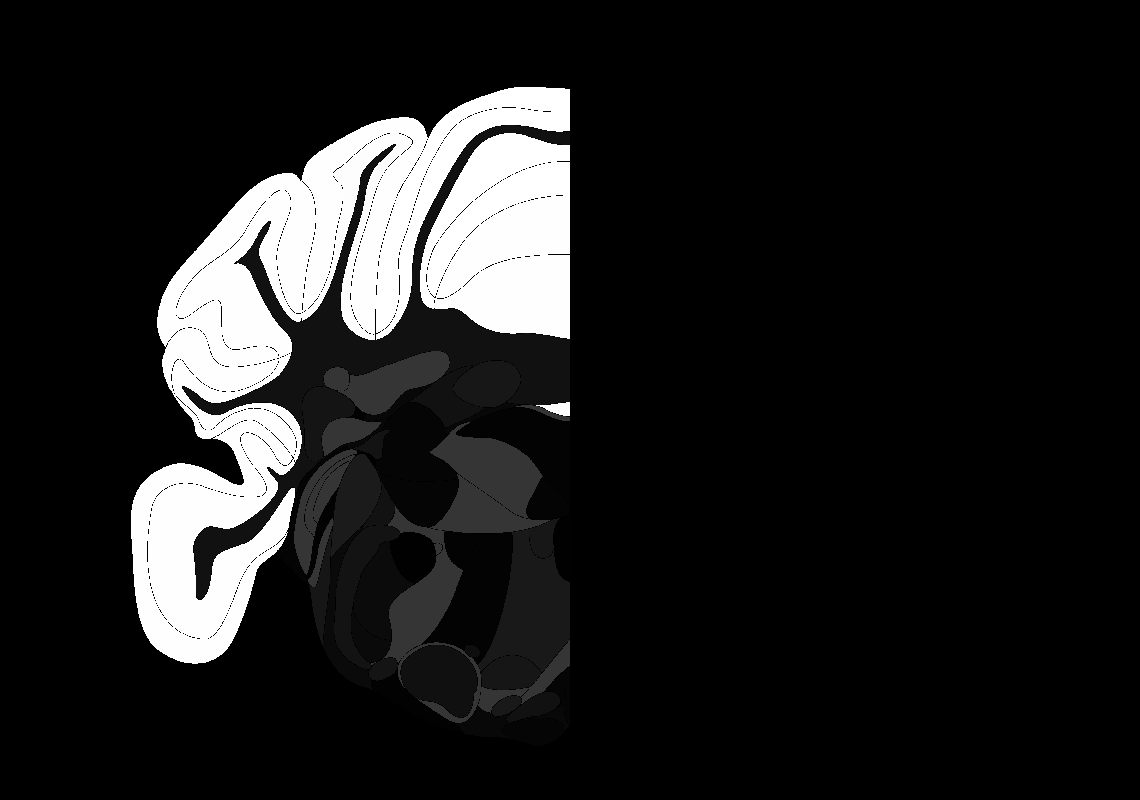

### 106_AP-6.2.tif

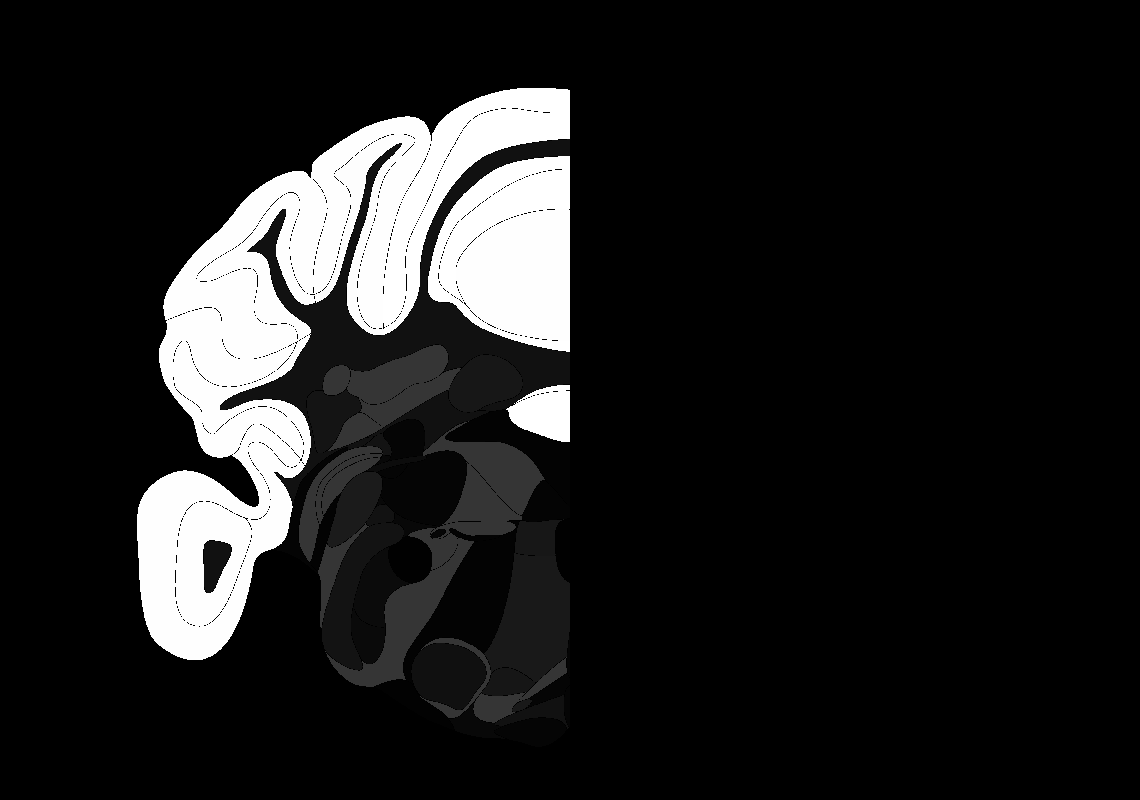

### 107_AP-6.3.tif

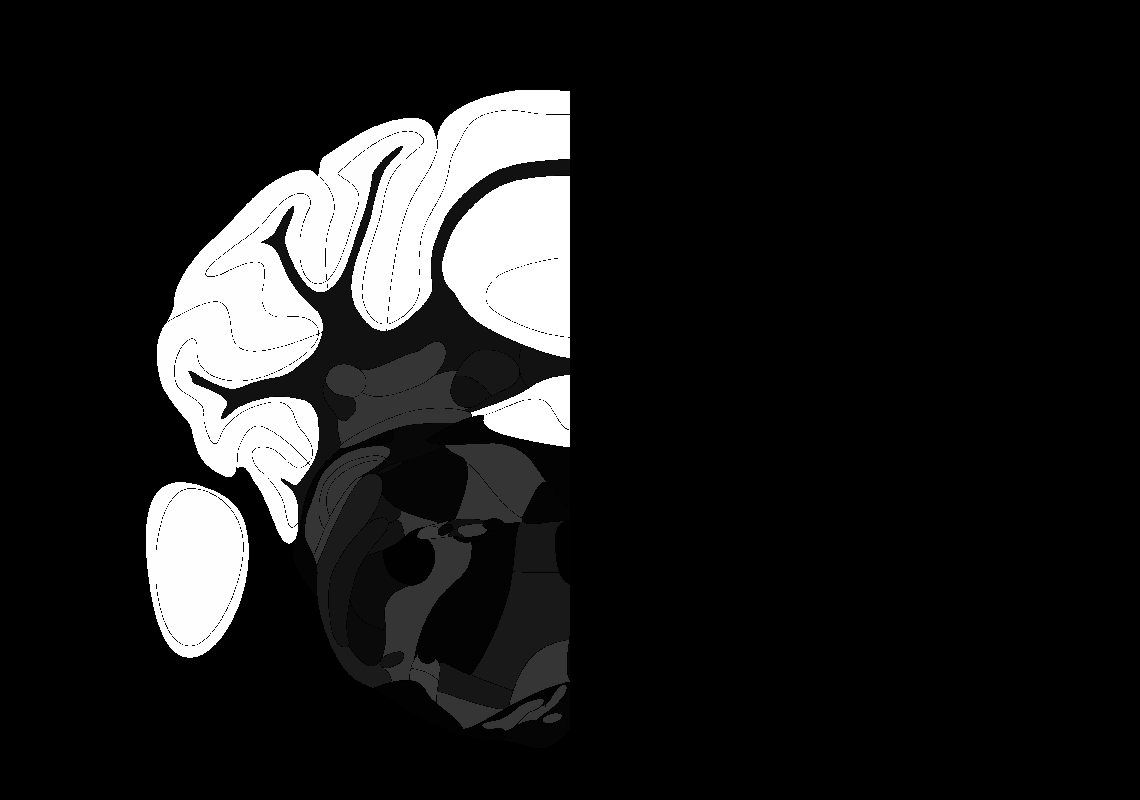

### 109_AP-6.5.tif

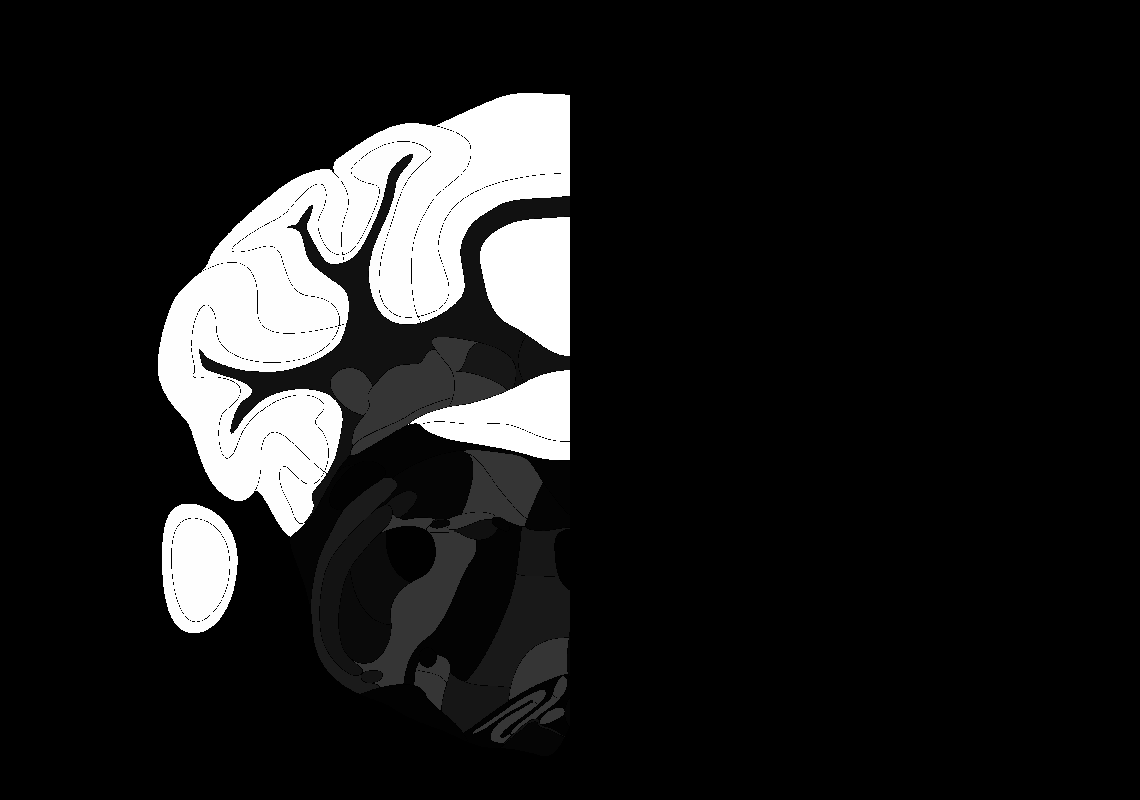

### 110_AP-6.6.tif

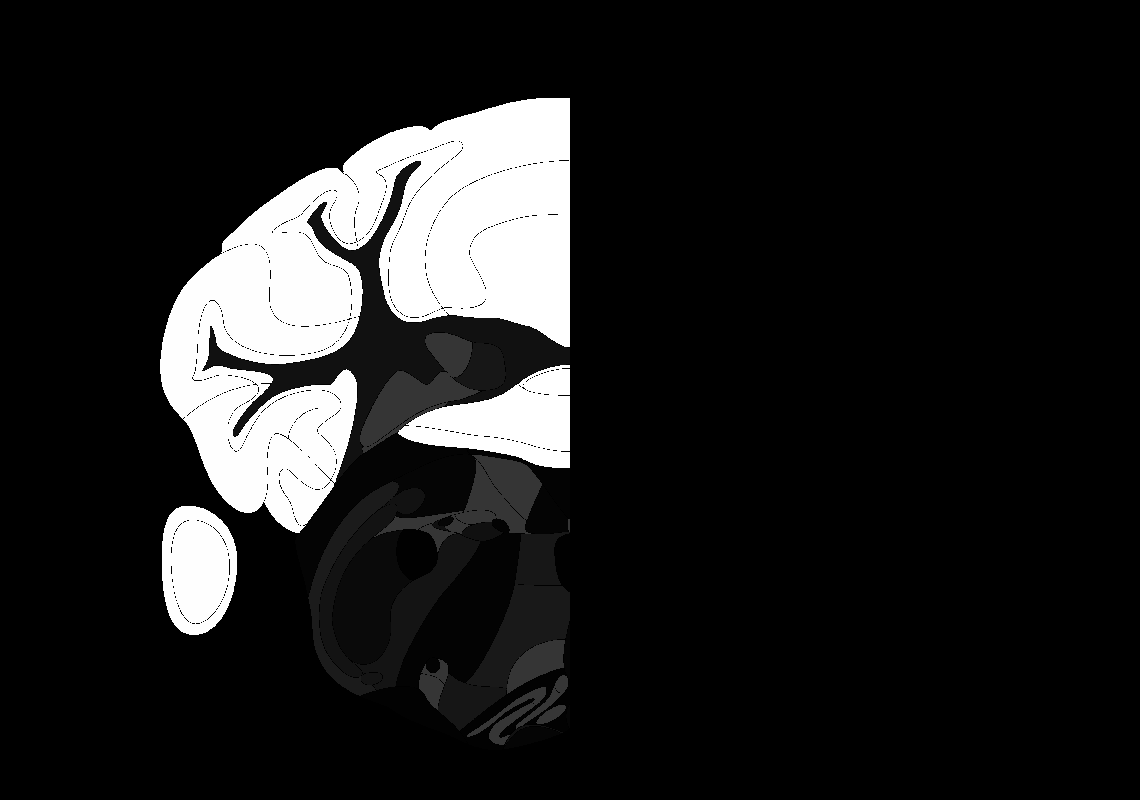

### 111_AP-6.7.tif

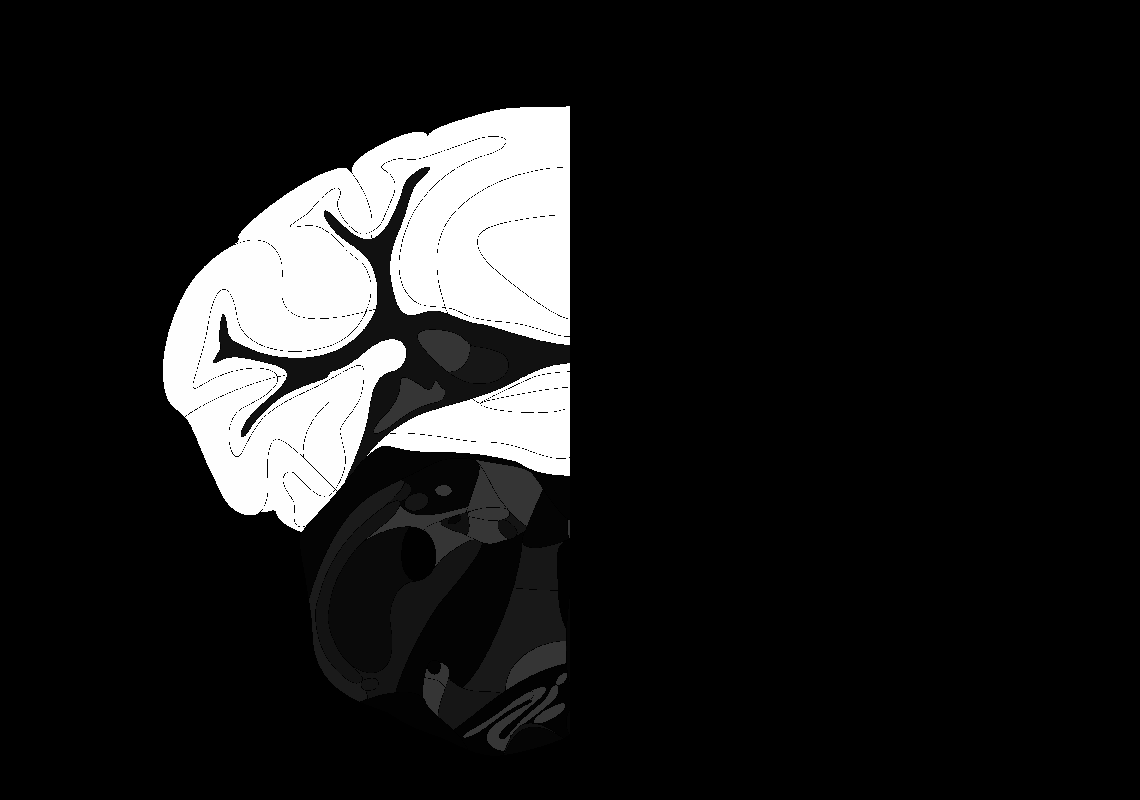

### 112_AP-6.8.tif

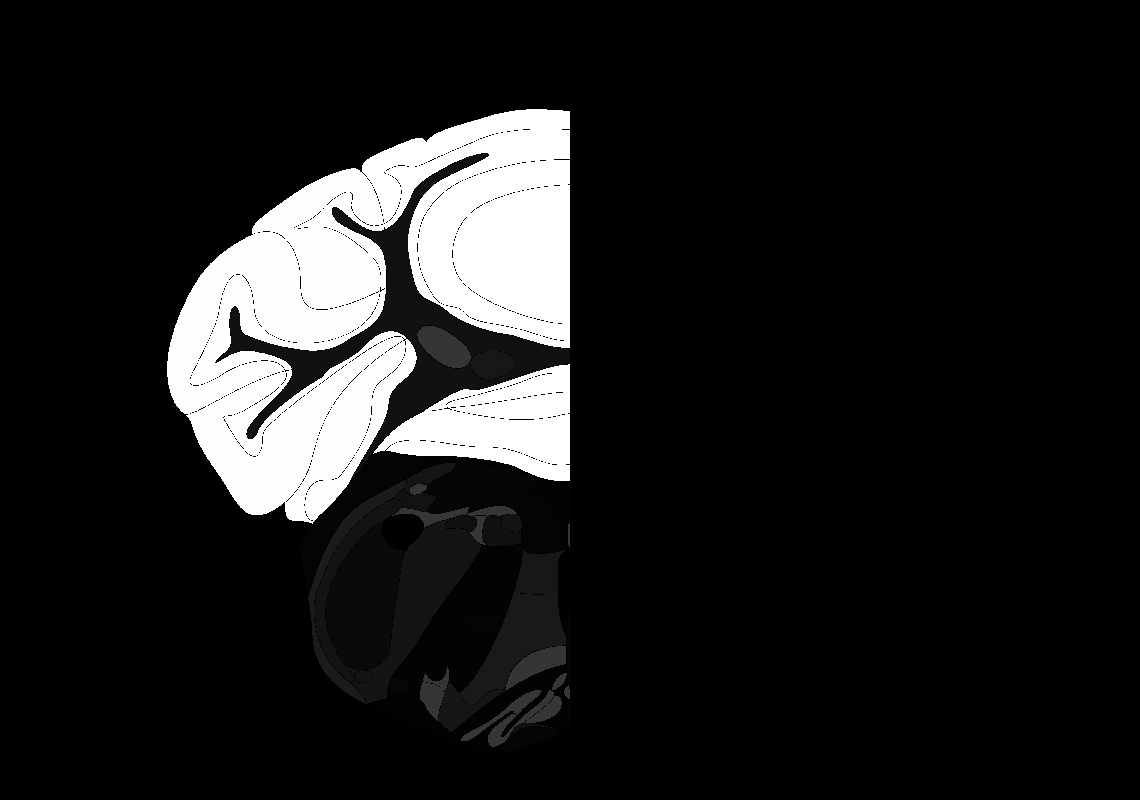

### 113_AP-6.9.tif

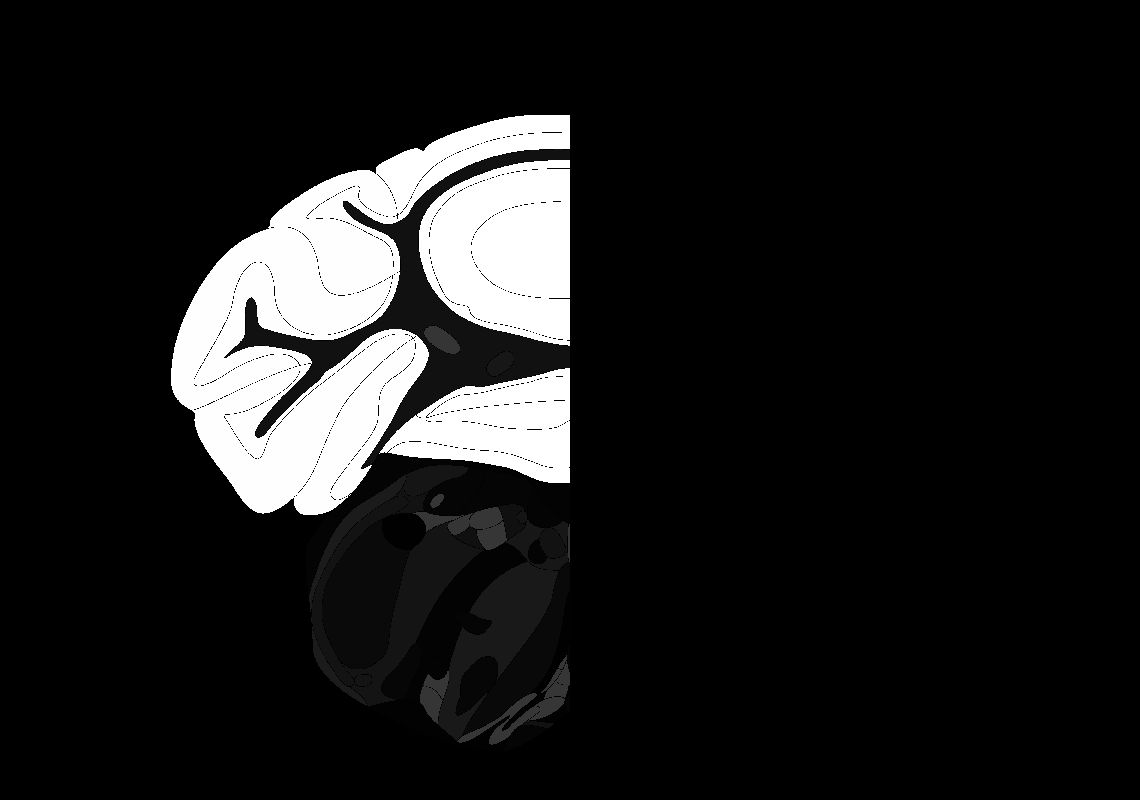

### 114_AP-7.0.tif

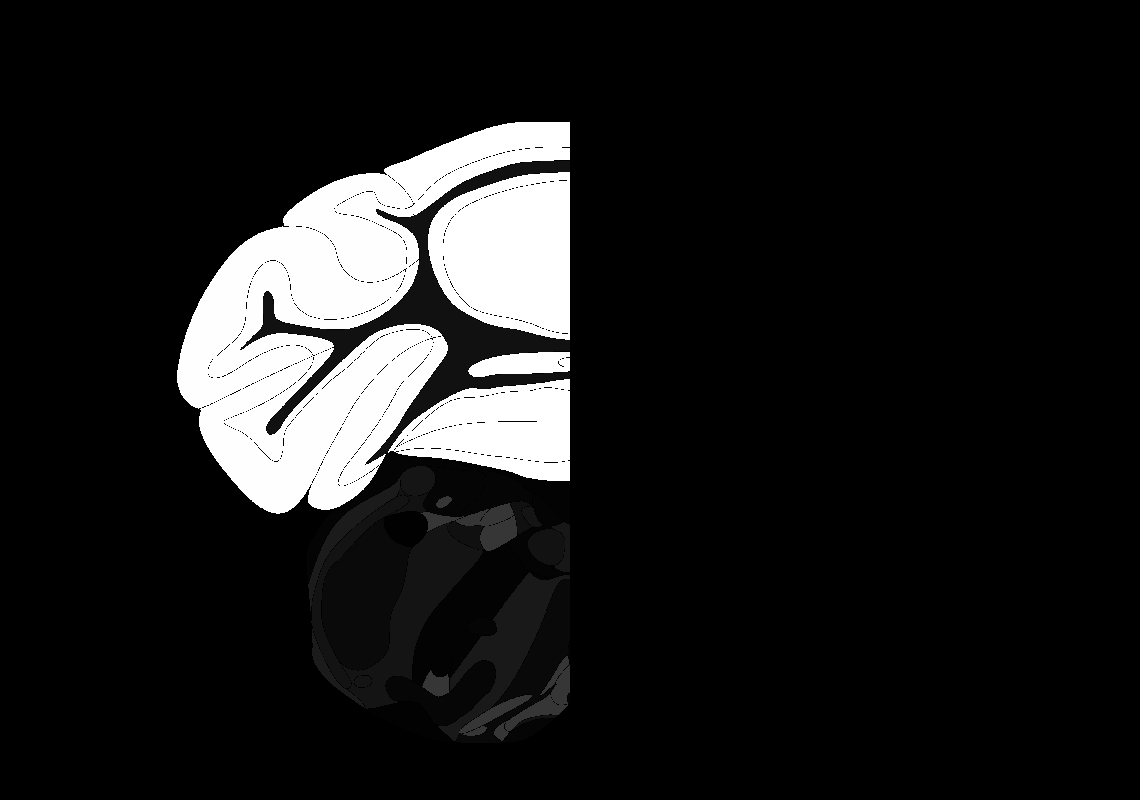

### 115_AP-7.1.tif

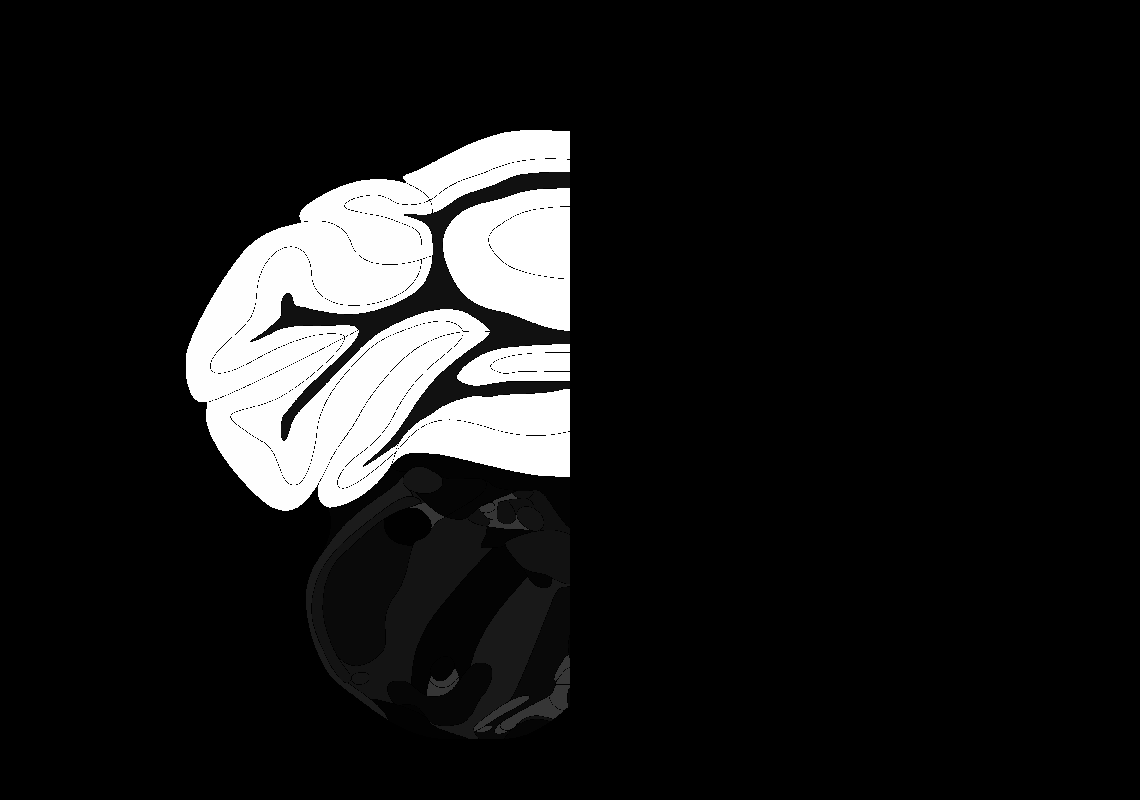

### 117_AP-7.3.tif

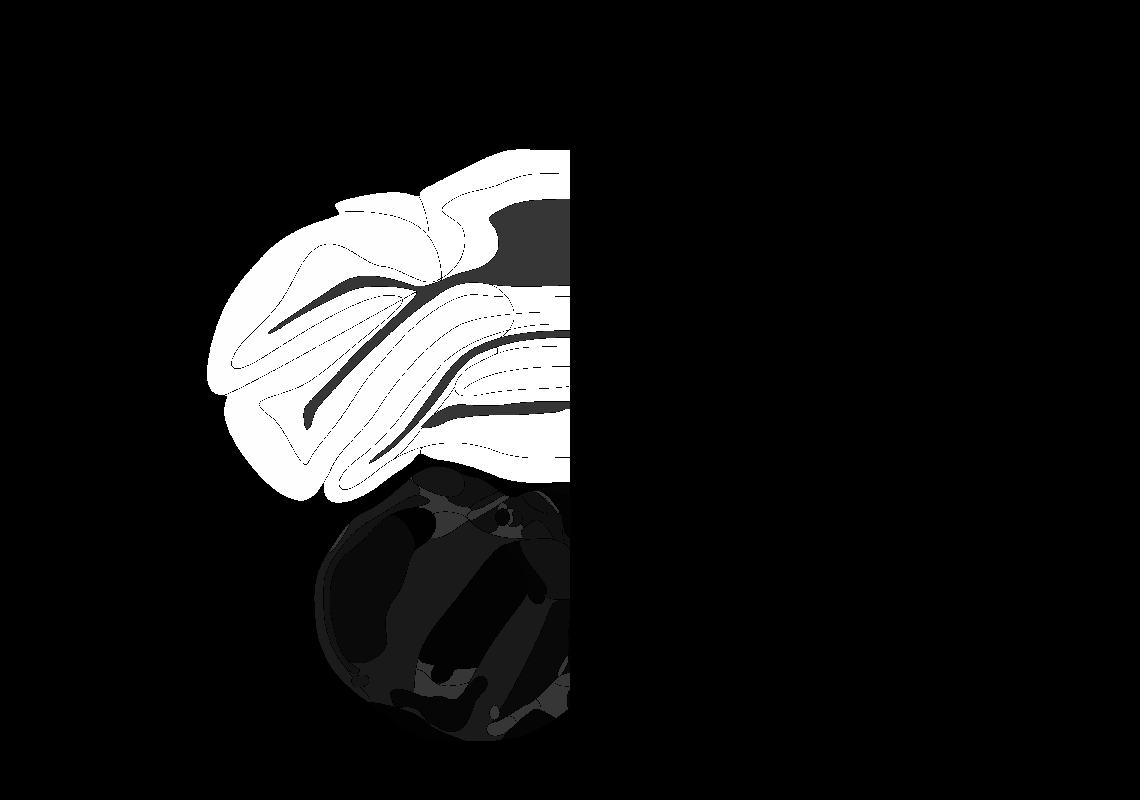

### 118_AP-7.4.tif

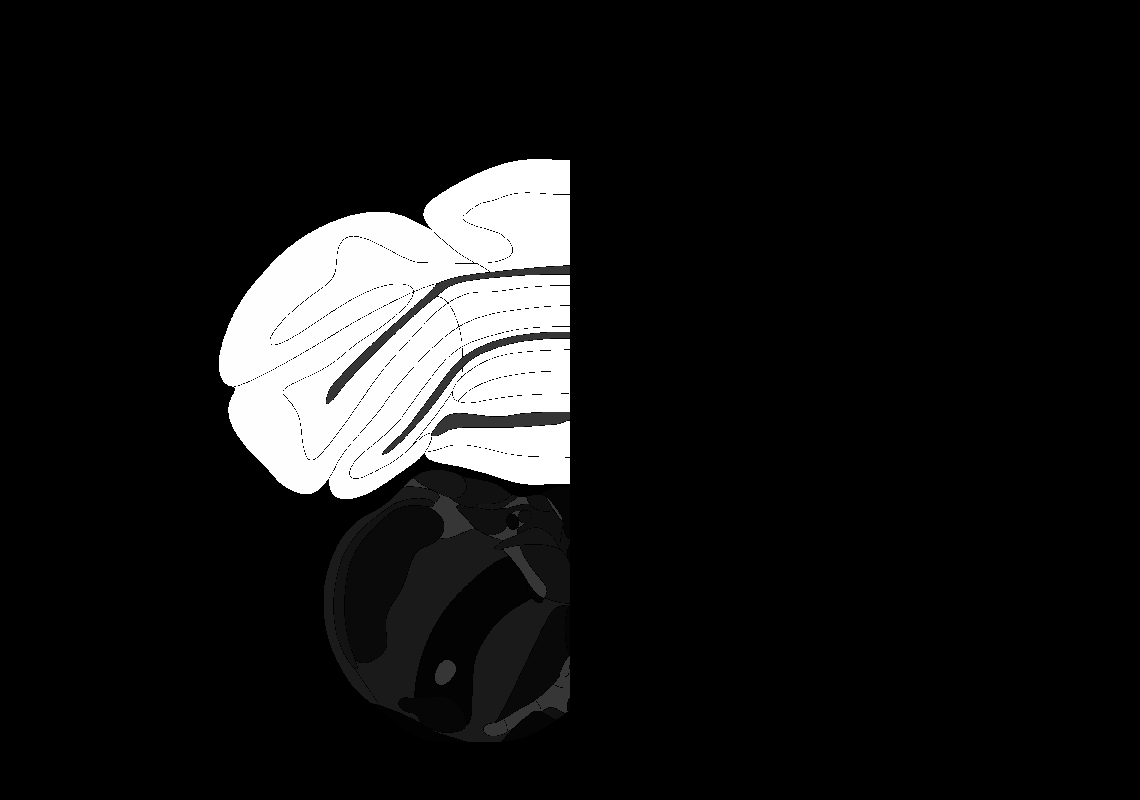

### 119_AP-7.5.tif

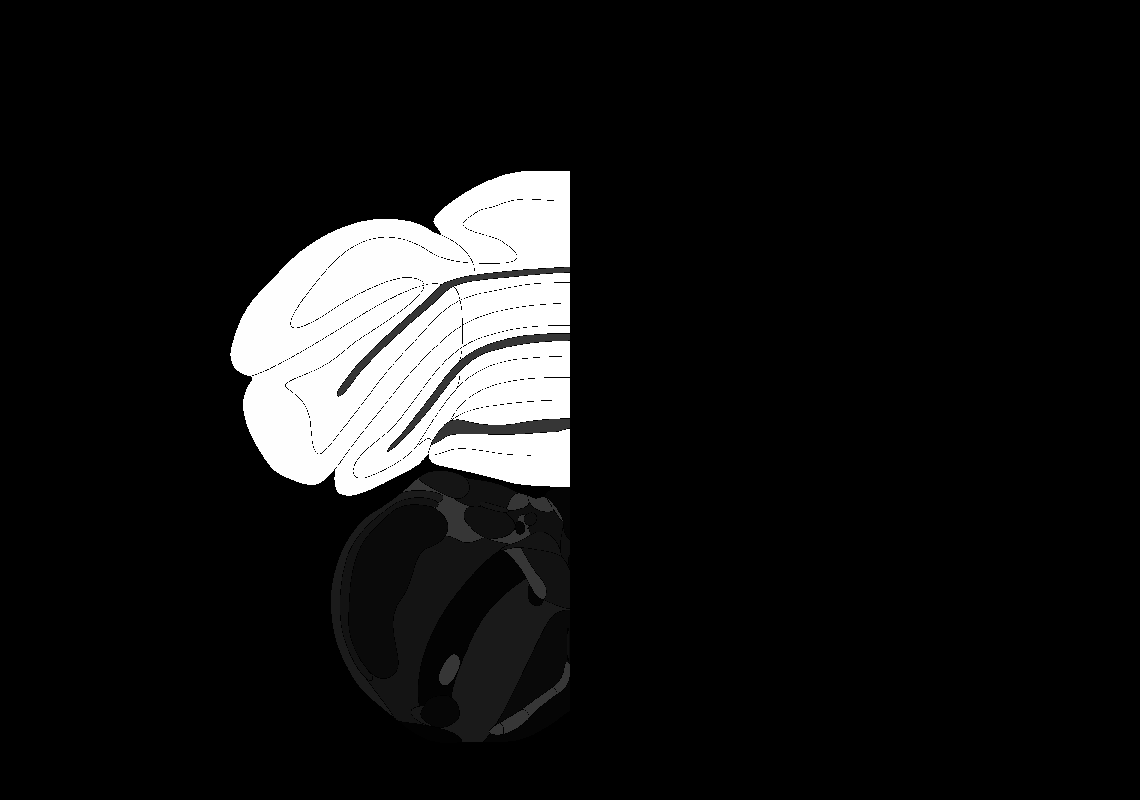

### 120_AP-7.6.tif

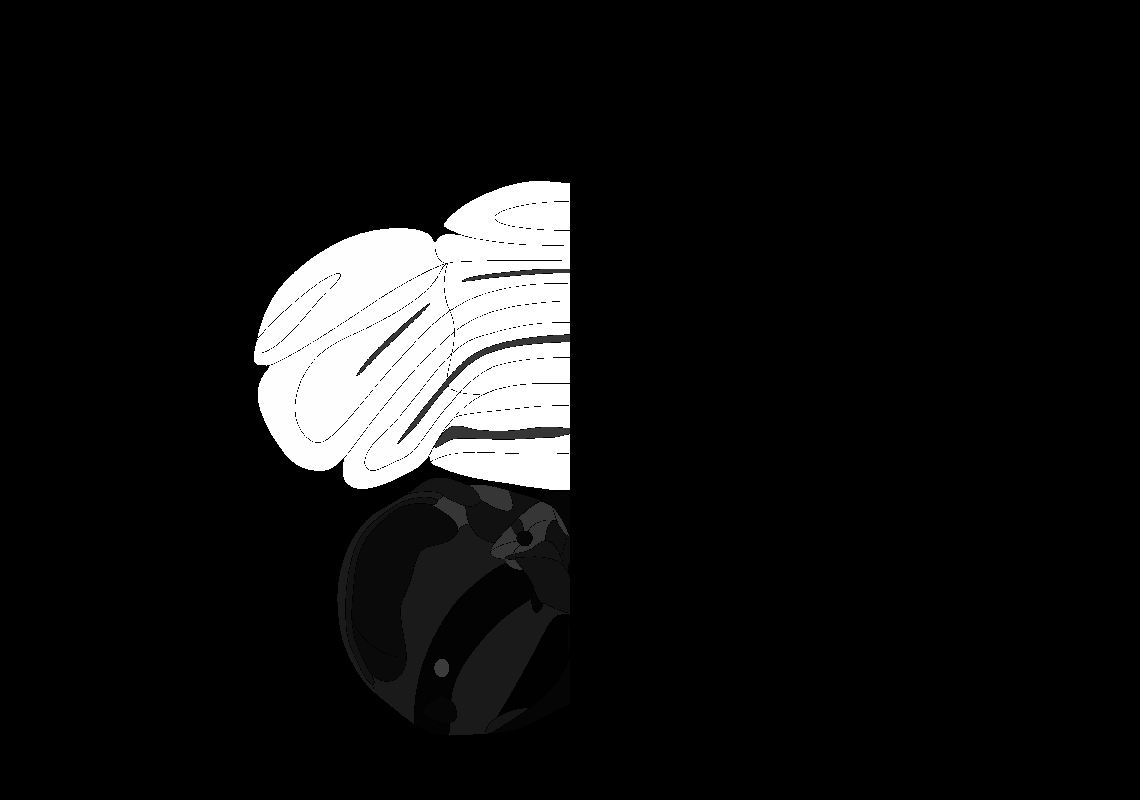

### 123_AP-7.9.tif

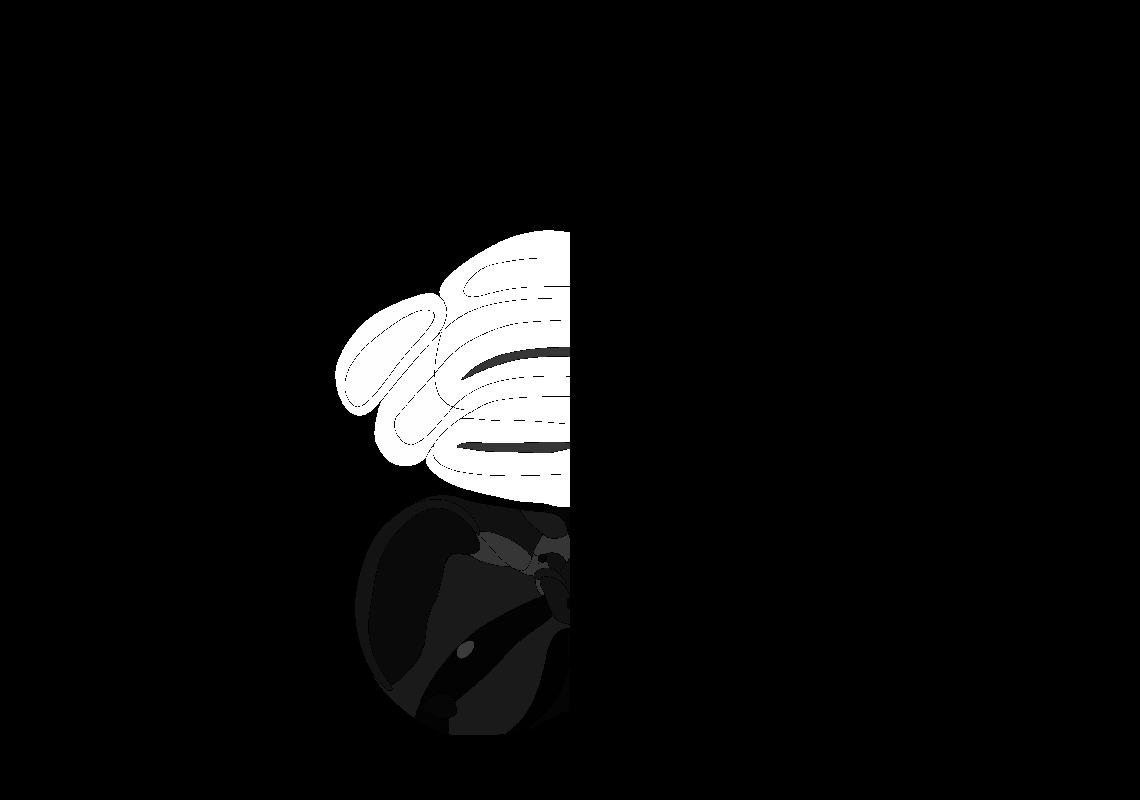
