## Supplementary figures and images for "Enhanced and Unified Anatomical Labeling for a Common Mouse Brain Atlas"

### AllenCCF_Z078.tif

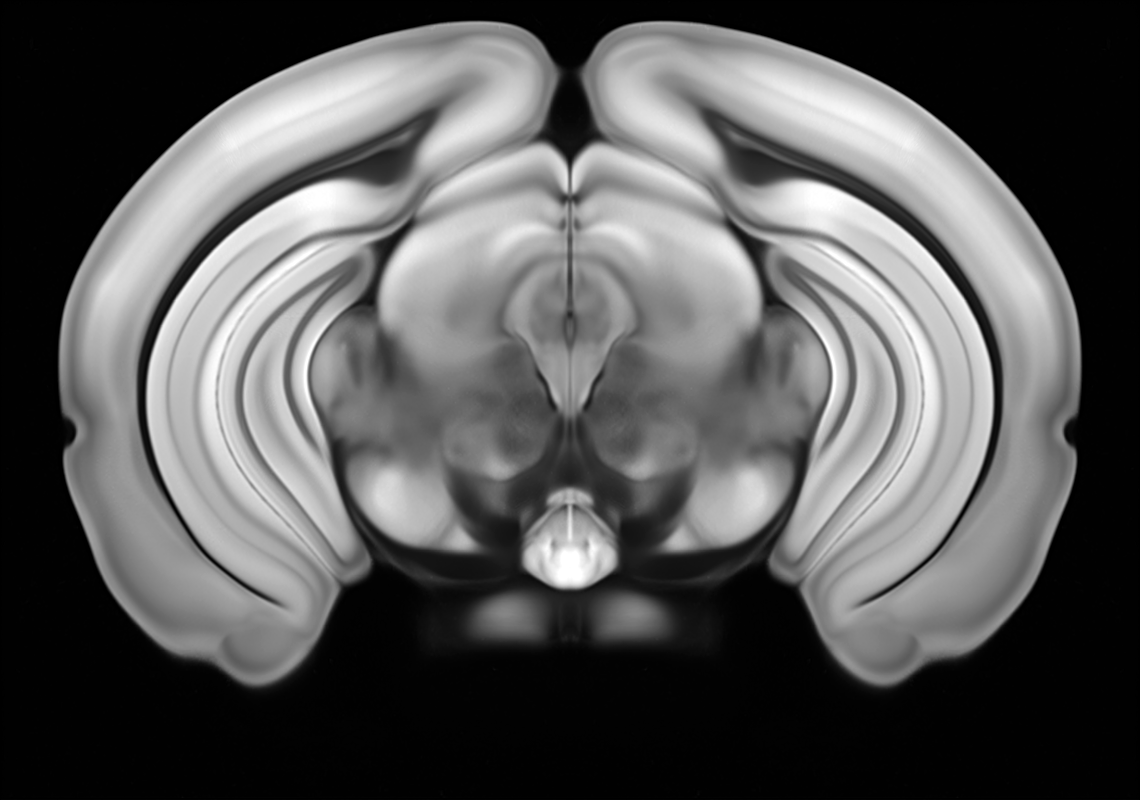

### AllenCCF_Z079.tif

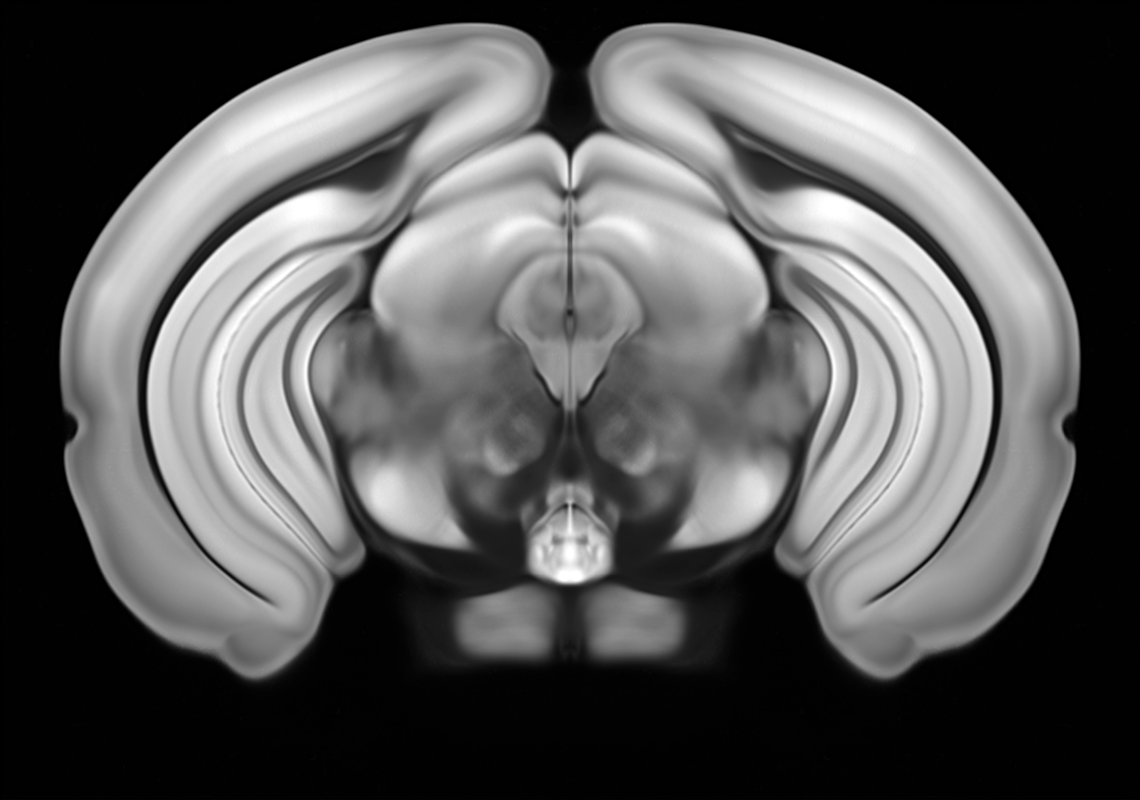

### AllenCCF_Z080.tif

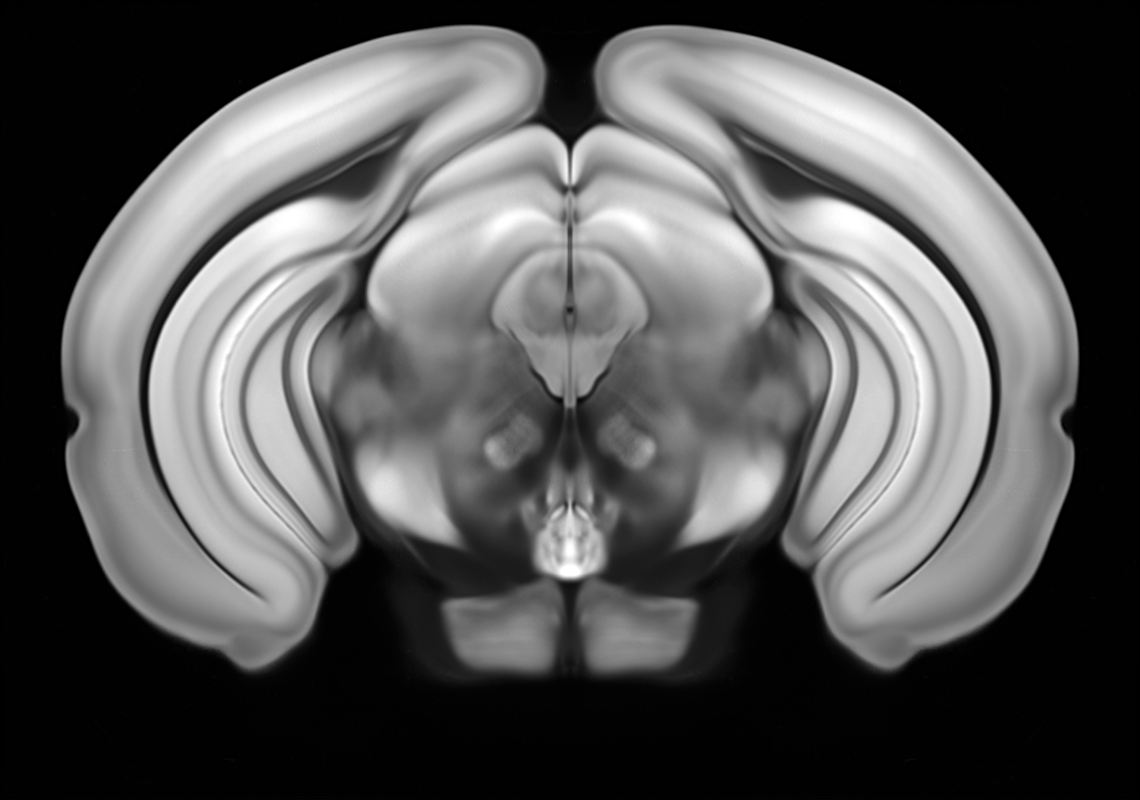

### AllenCCF_Z081.tif

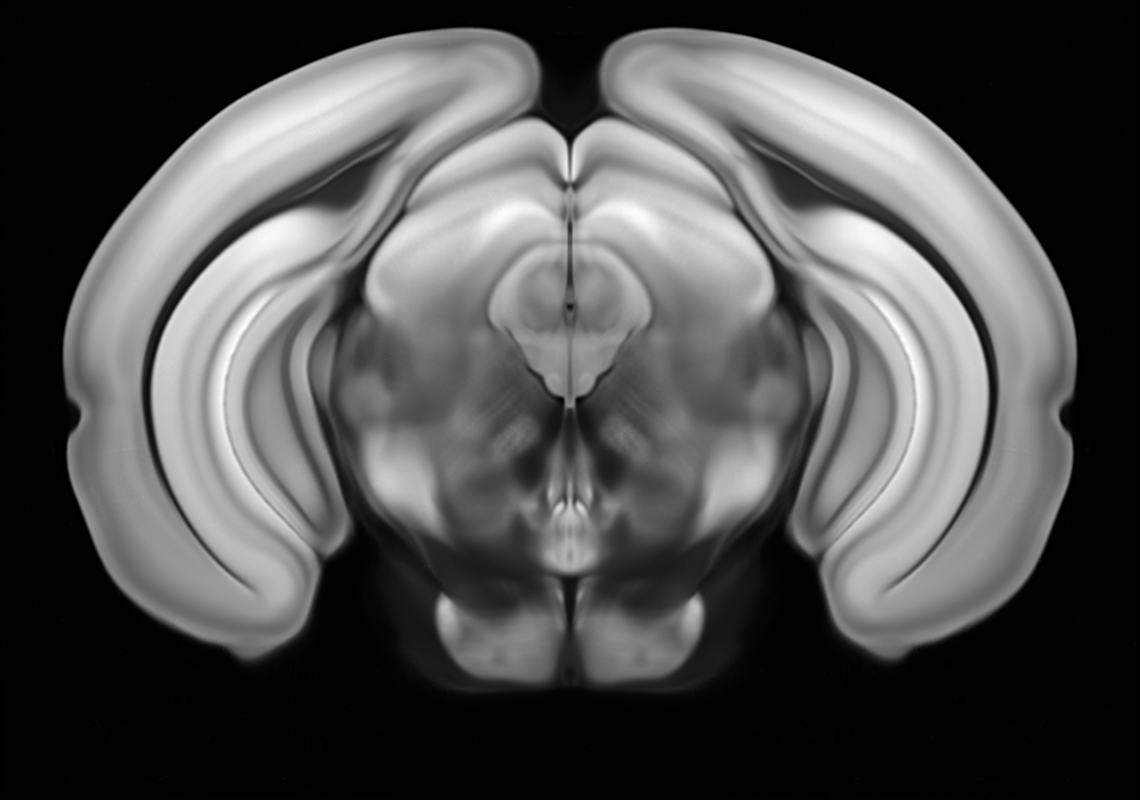

### AllenCCF_Z082.tif

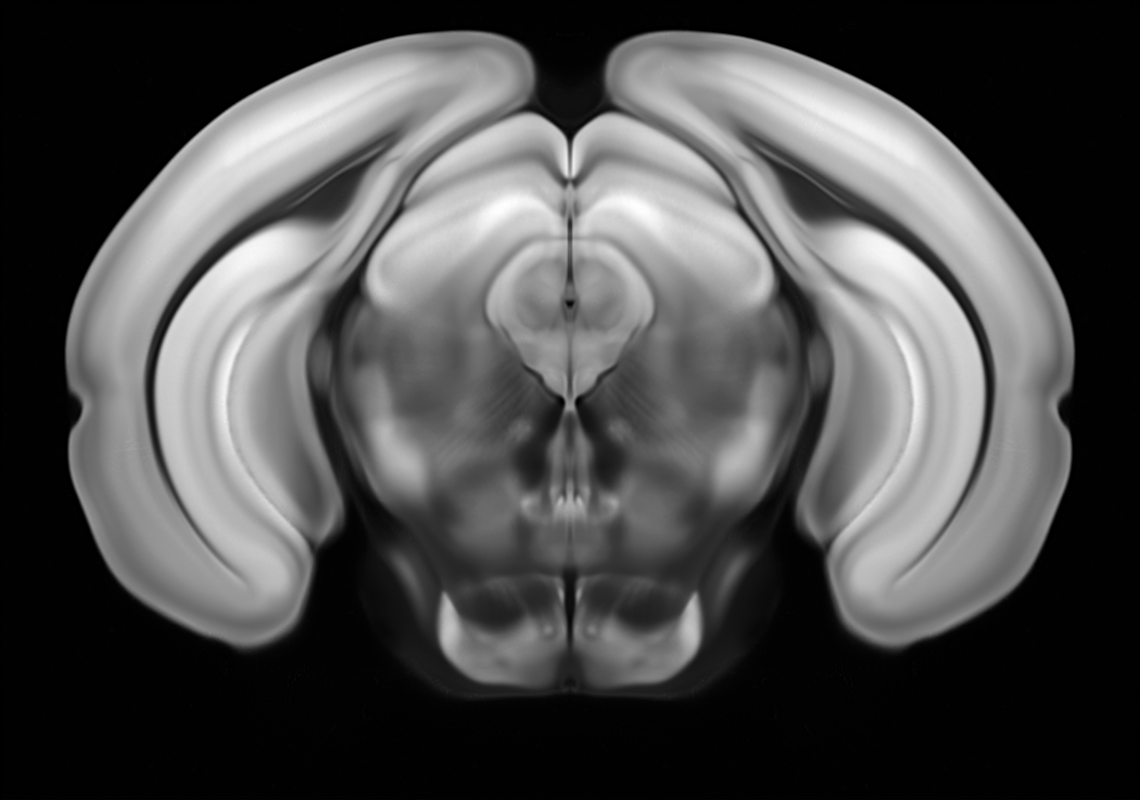

### AllenCCF_Z083.tif

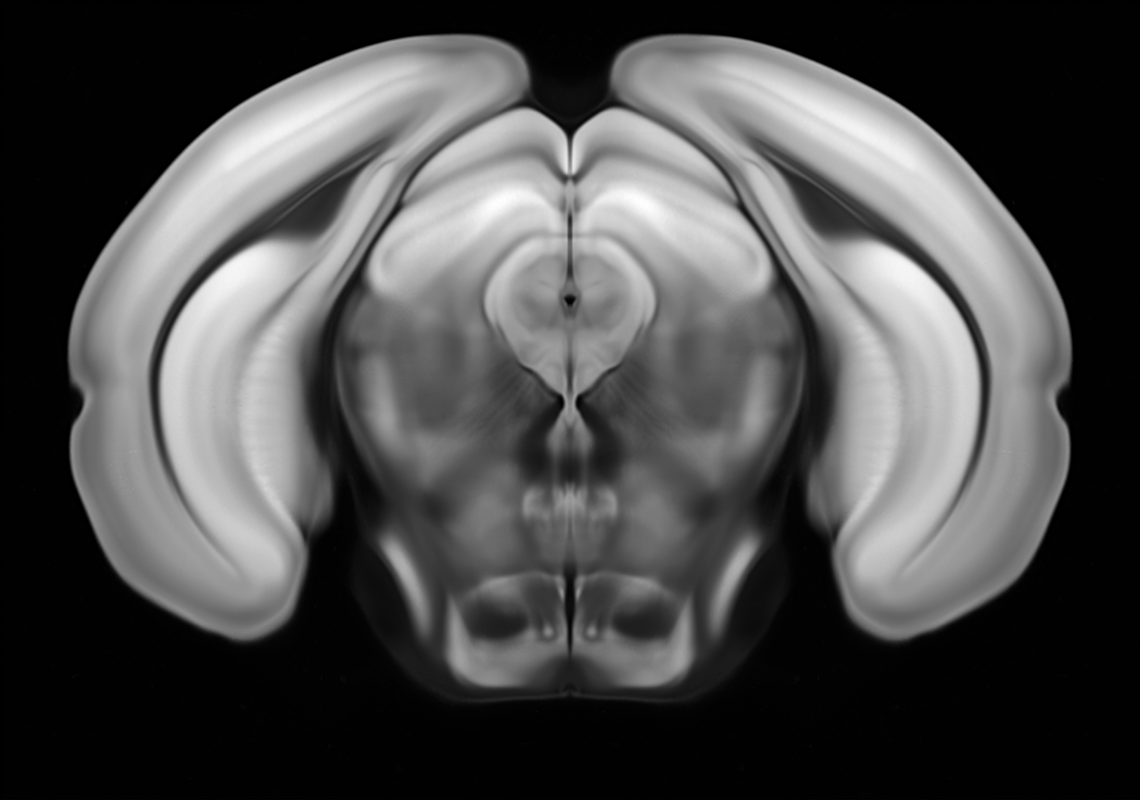

### AllenCCF_Z084.tif

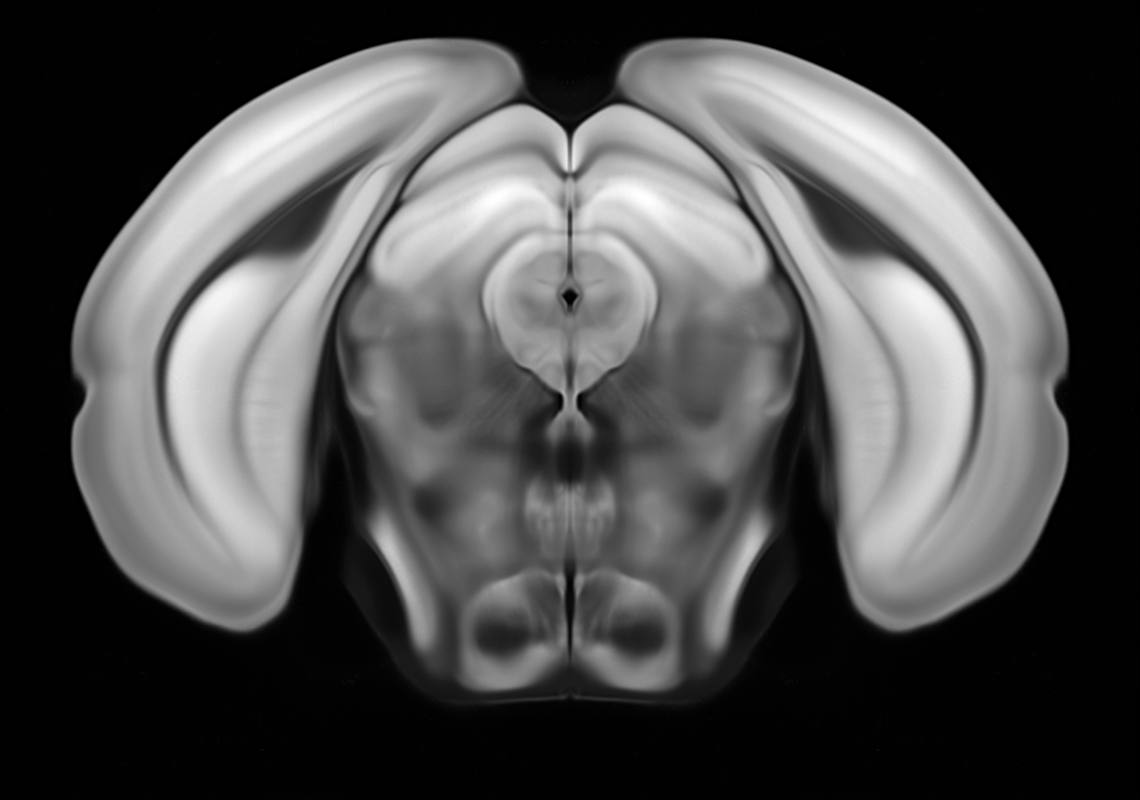

### AllenCCF_Z085.tif

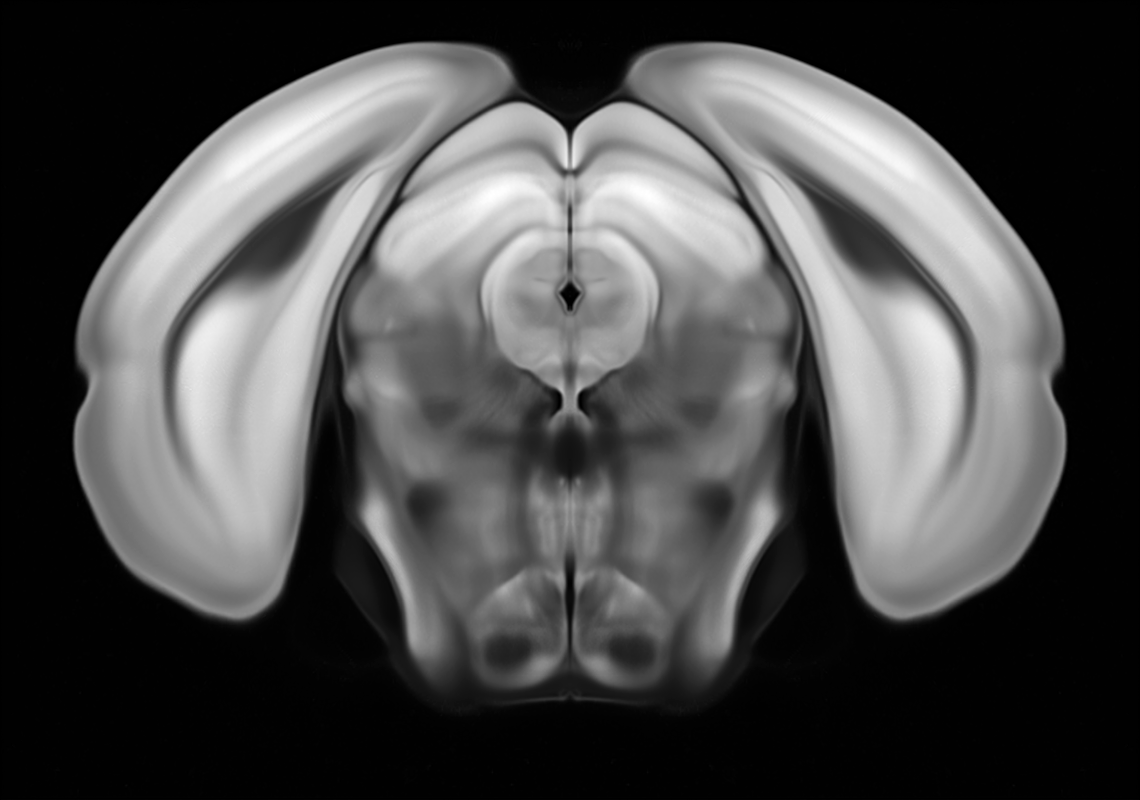

### AllenCCF_Z086.tif

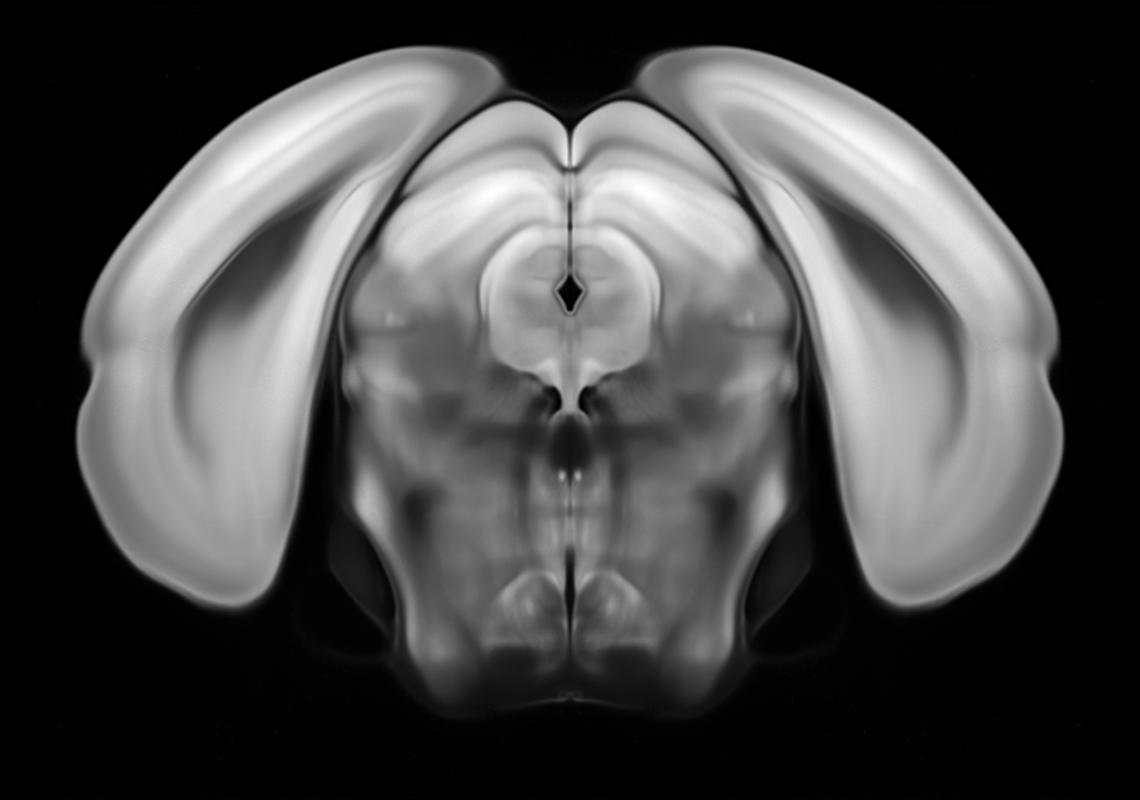

### AllenCCF_Z087.tif

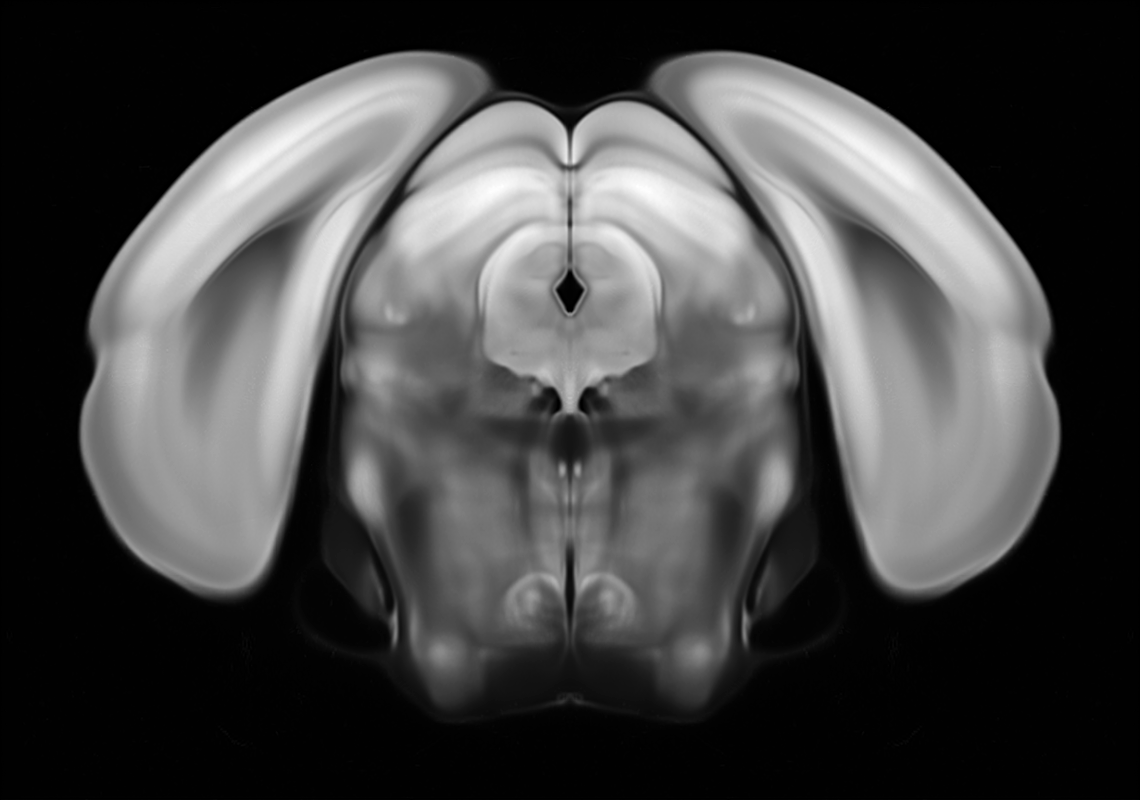

### AllenCCF_Z088.tif

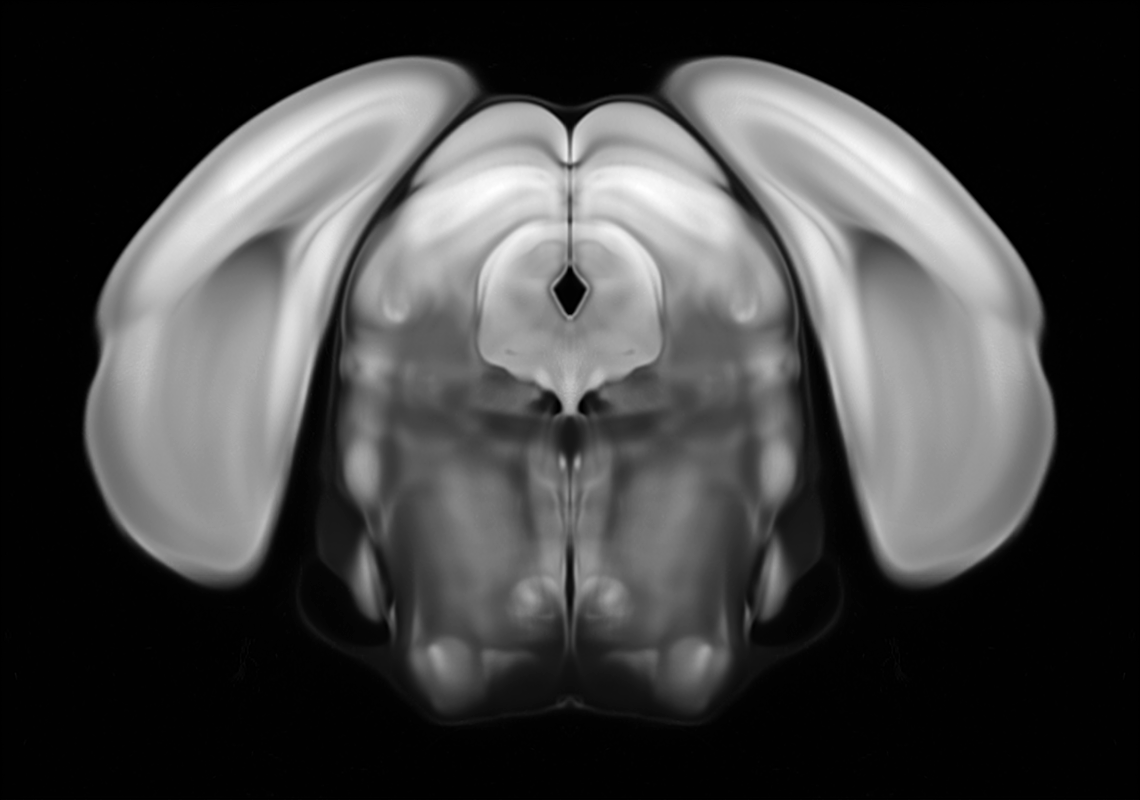

### AllenCCF_Z089.tif

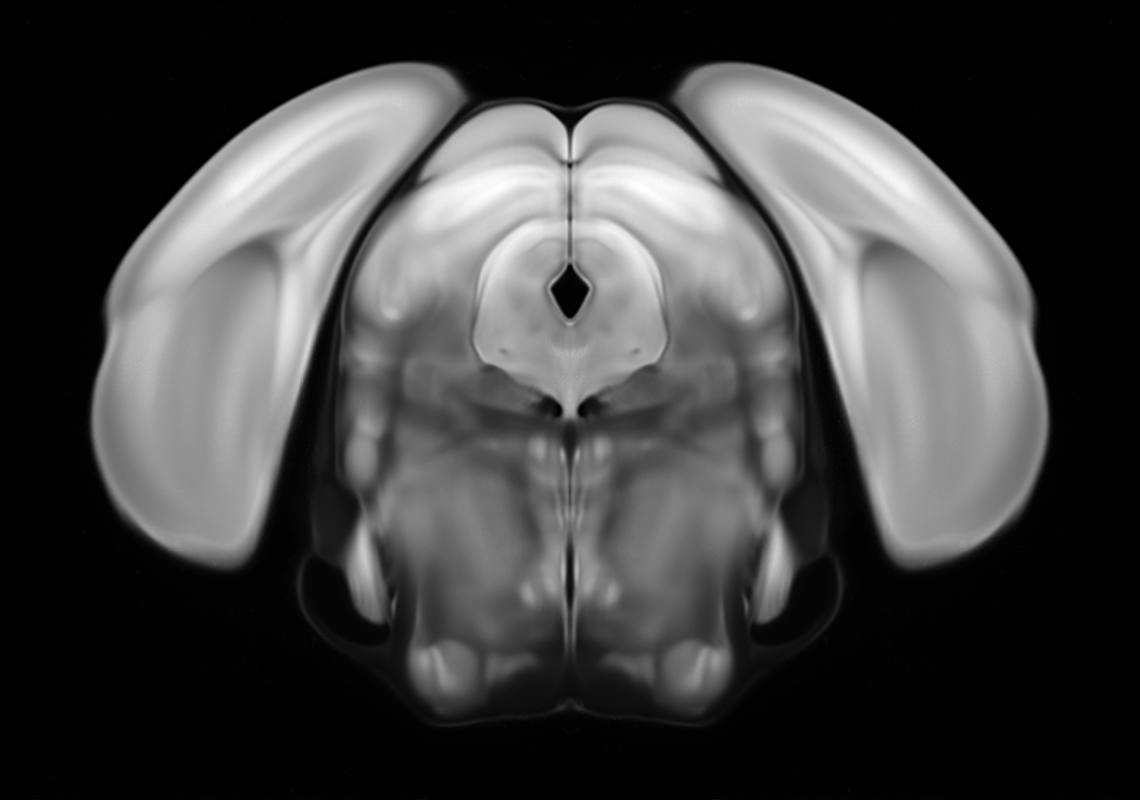

### AllenCCF_Z090.tif

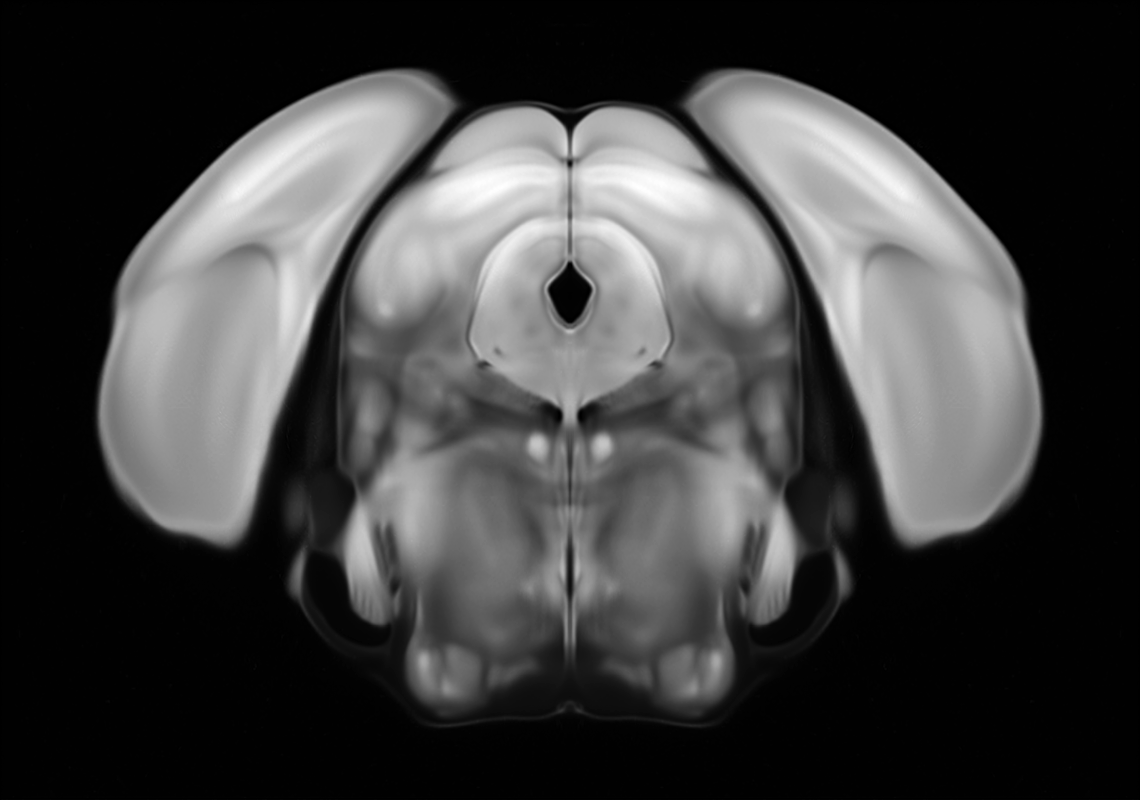

### AllenCCF_Z091.tif

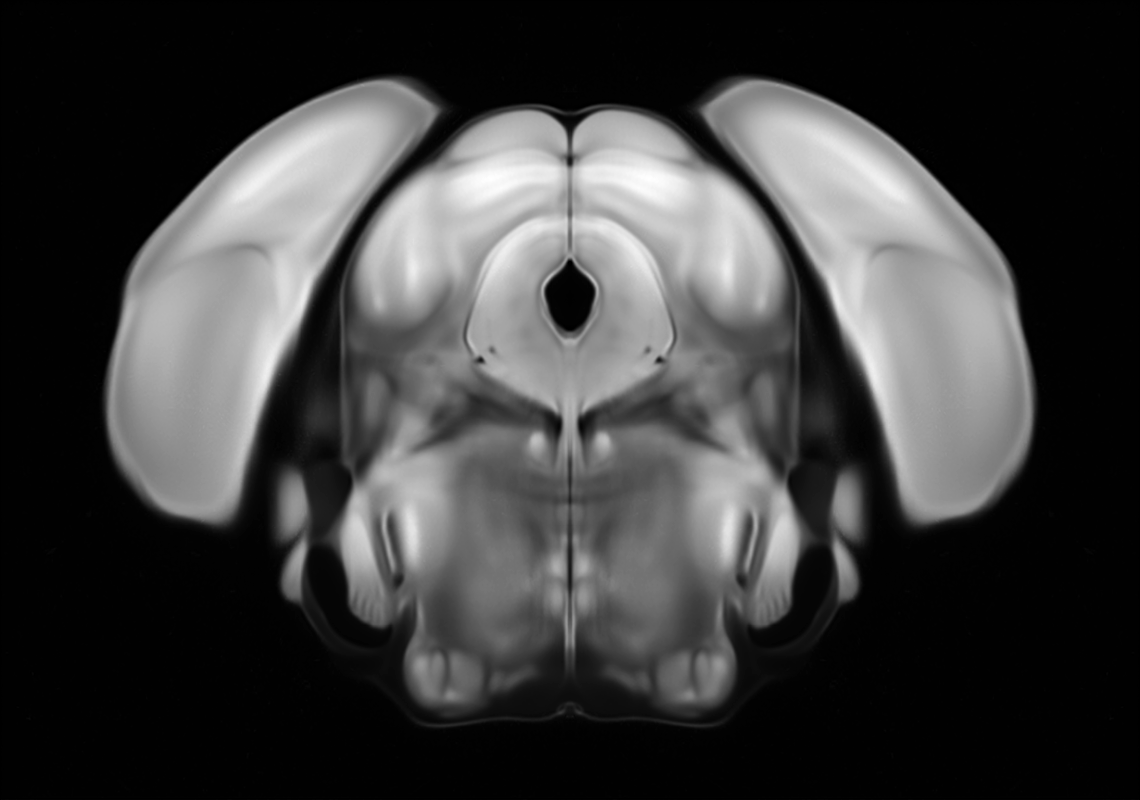

### AllenCCF_Z092.tif

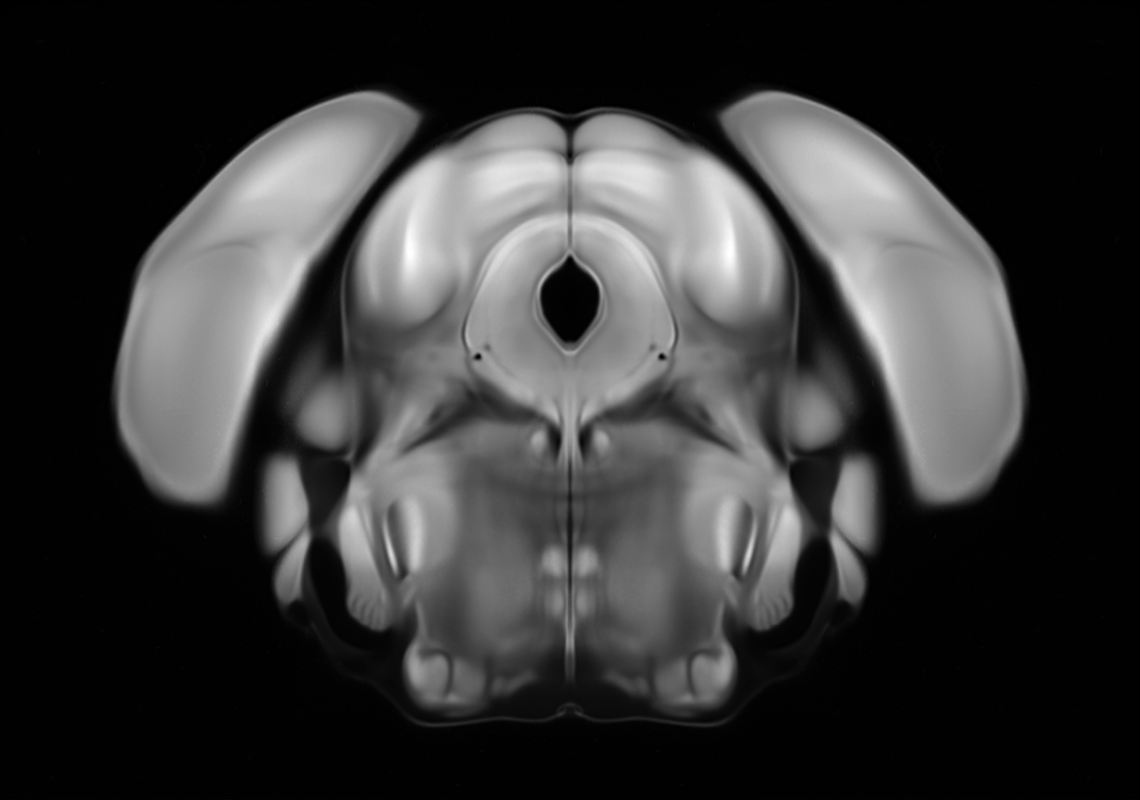

### AllenCCF_Z093.tif

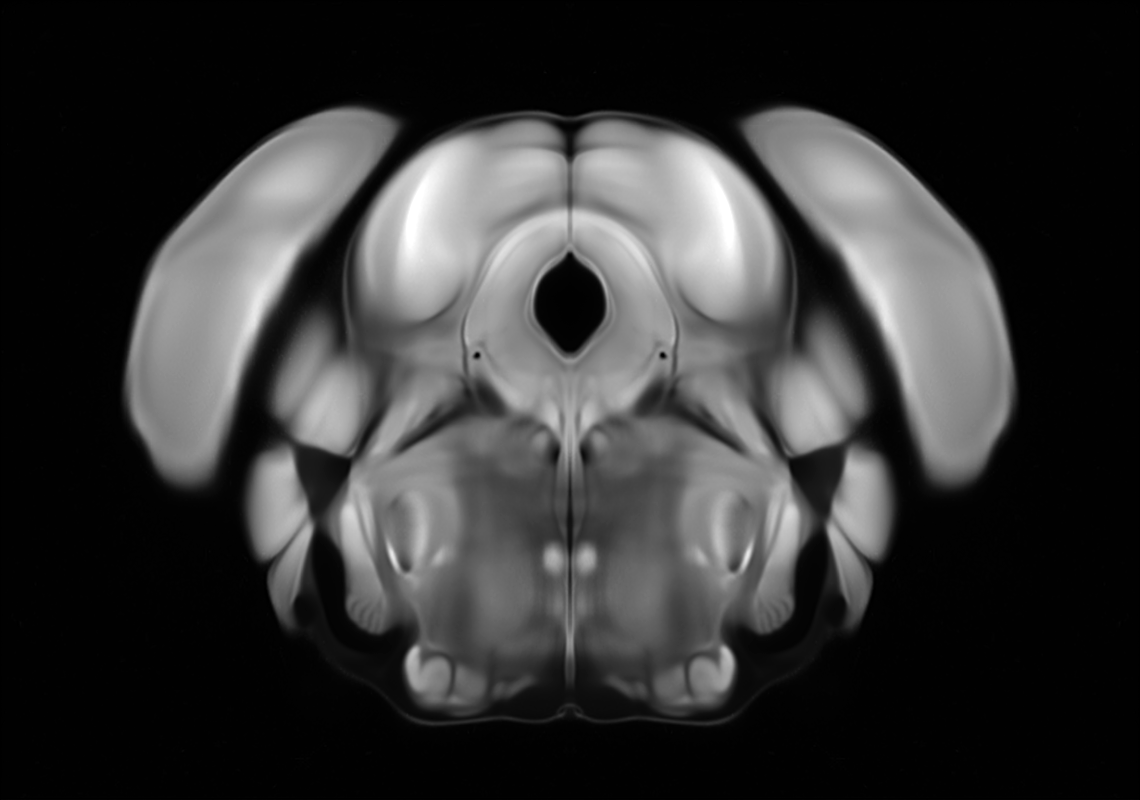

### AllenCCF_Z094.tif

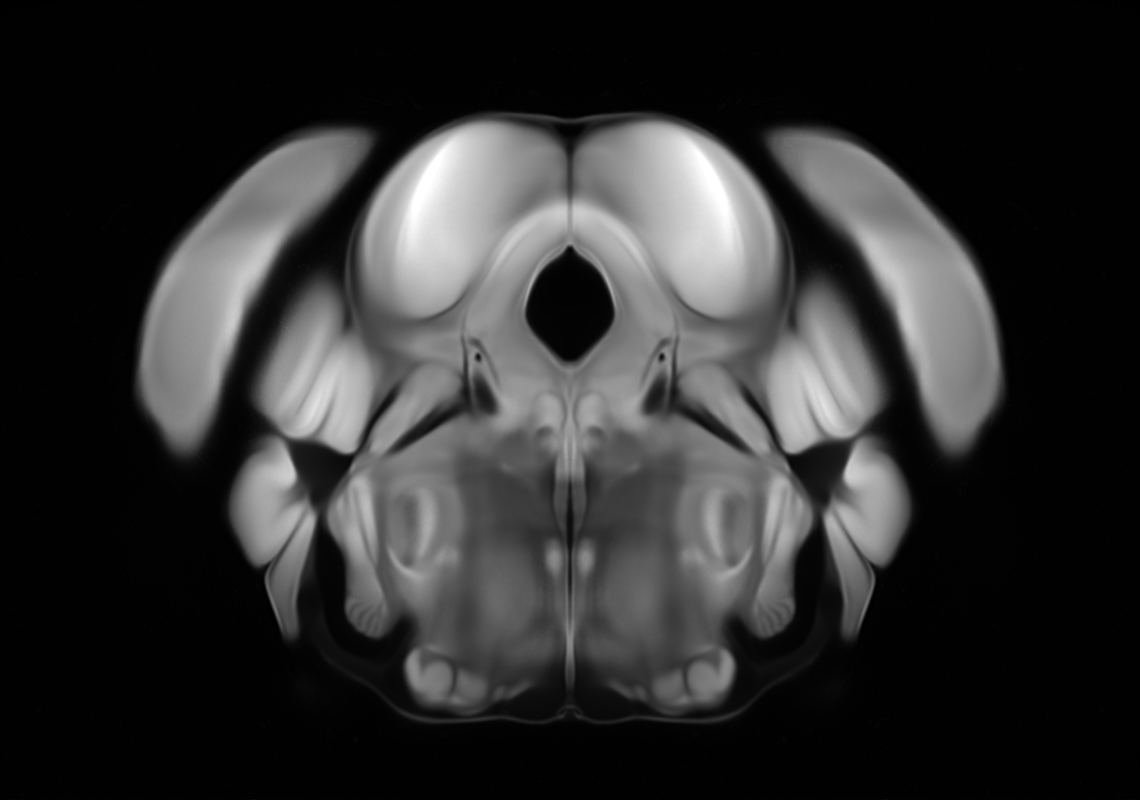

### AllenCCF_Z095.tif

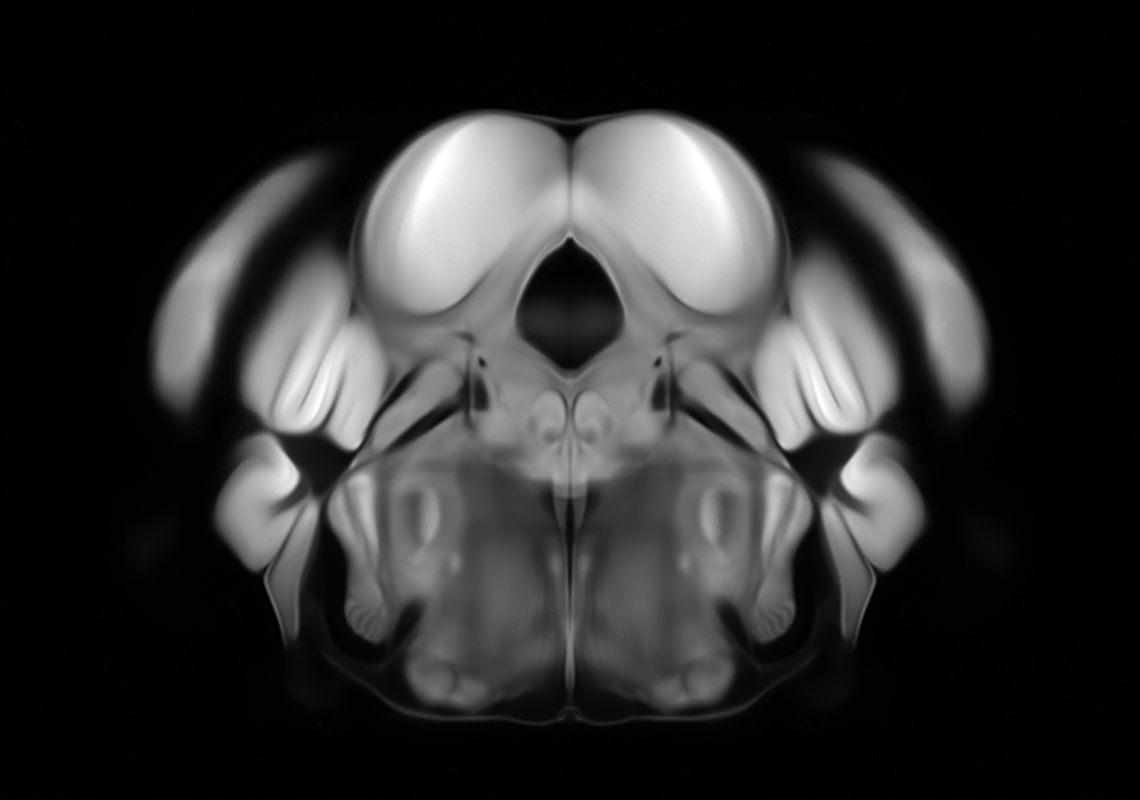

### AllenCCF_Z096.tif

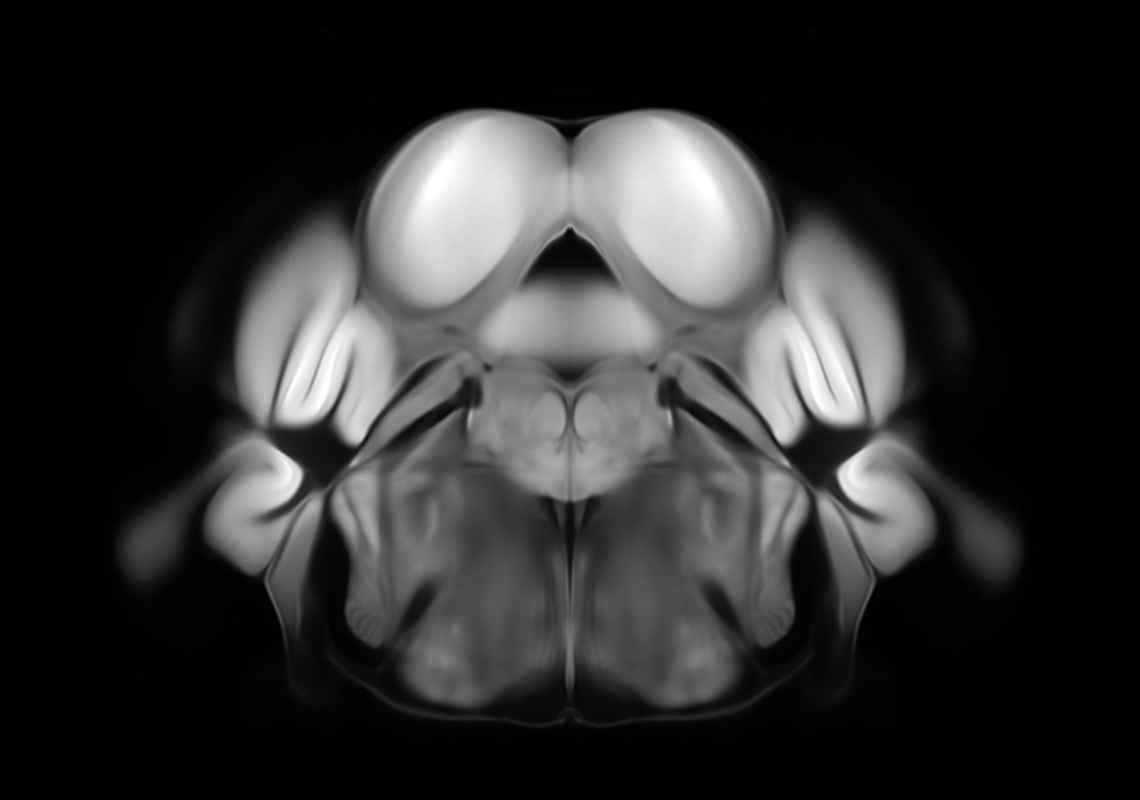

### AllenCCF_Z097.tif

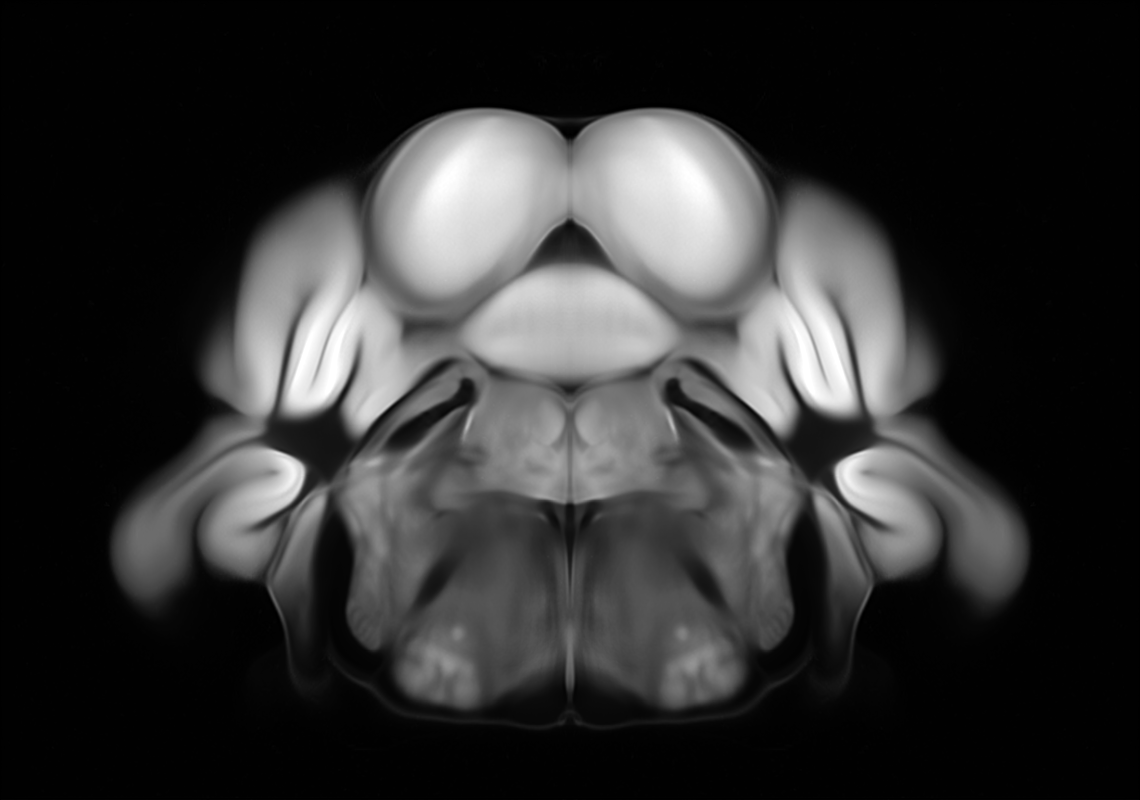

### AllenCCF_Z098.tif

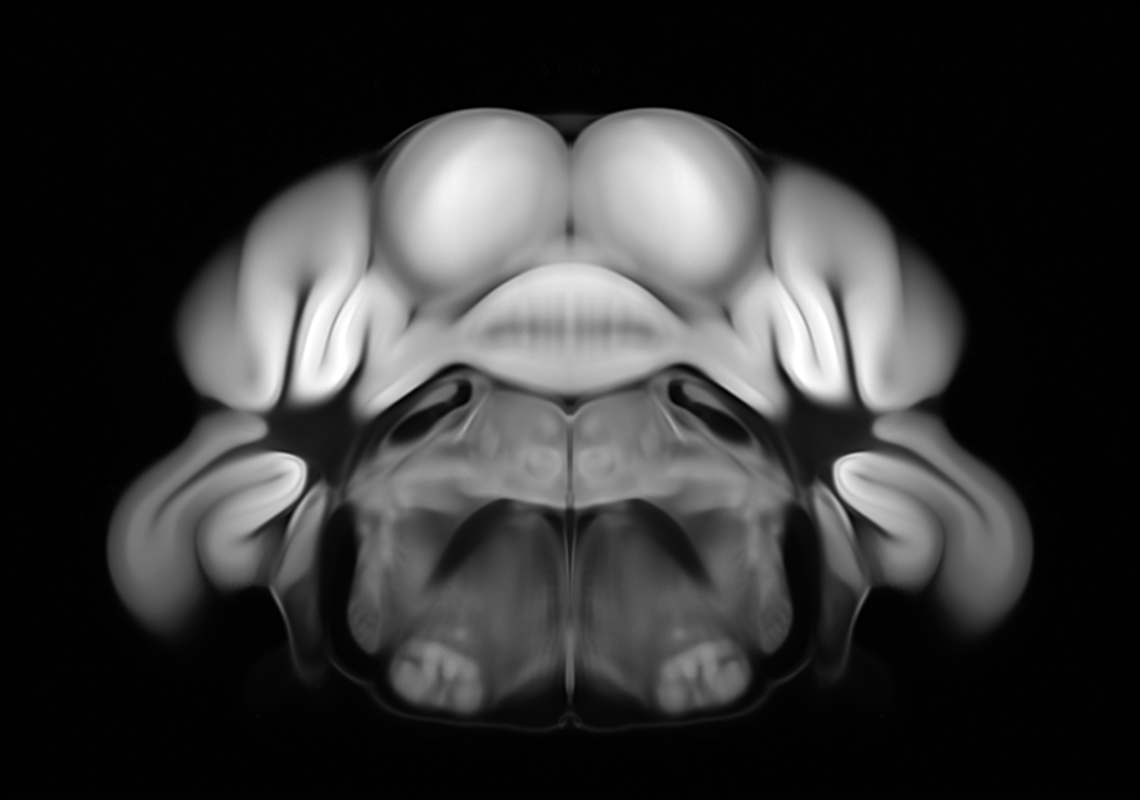

### AllenCCF_Z099.tif

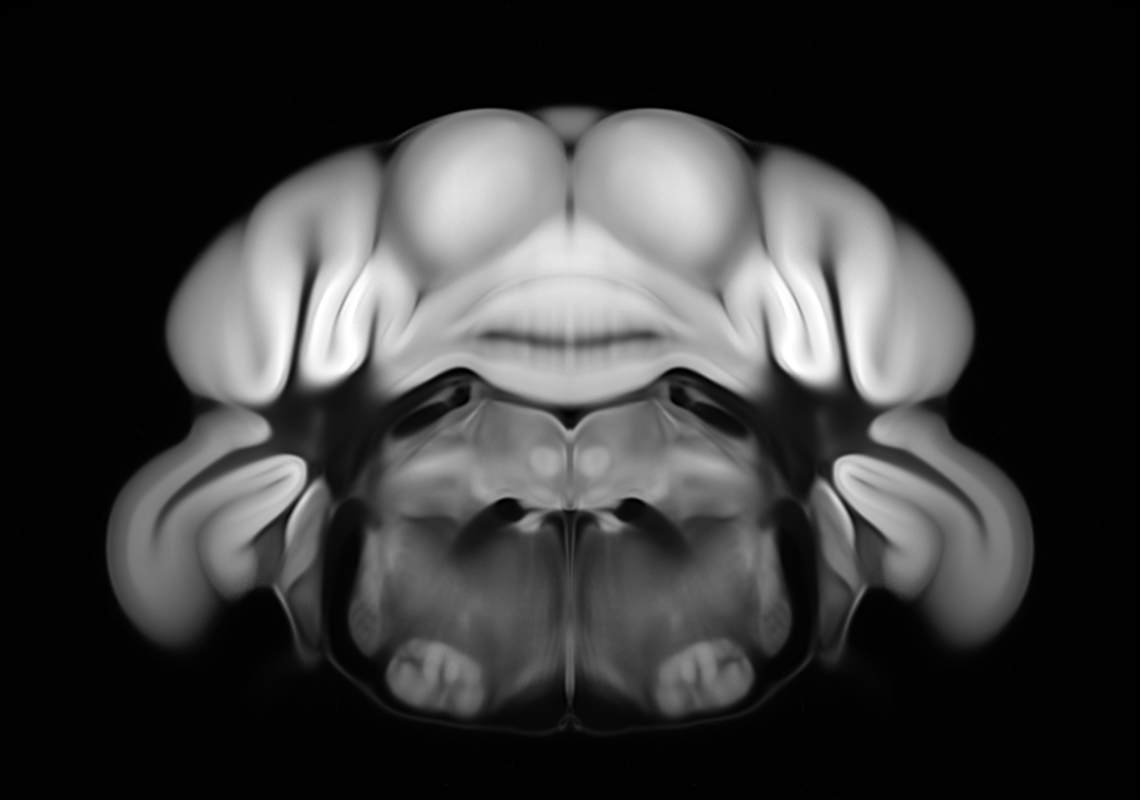

### AllenCCF_Z100.tif

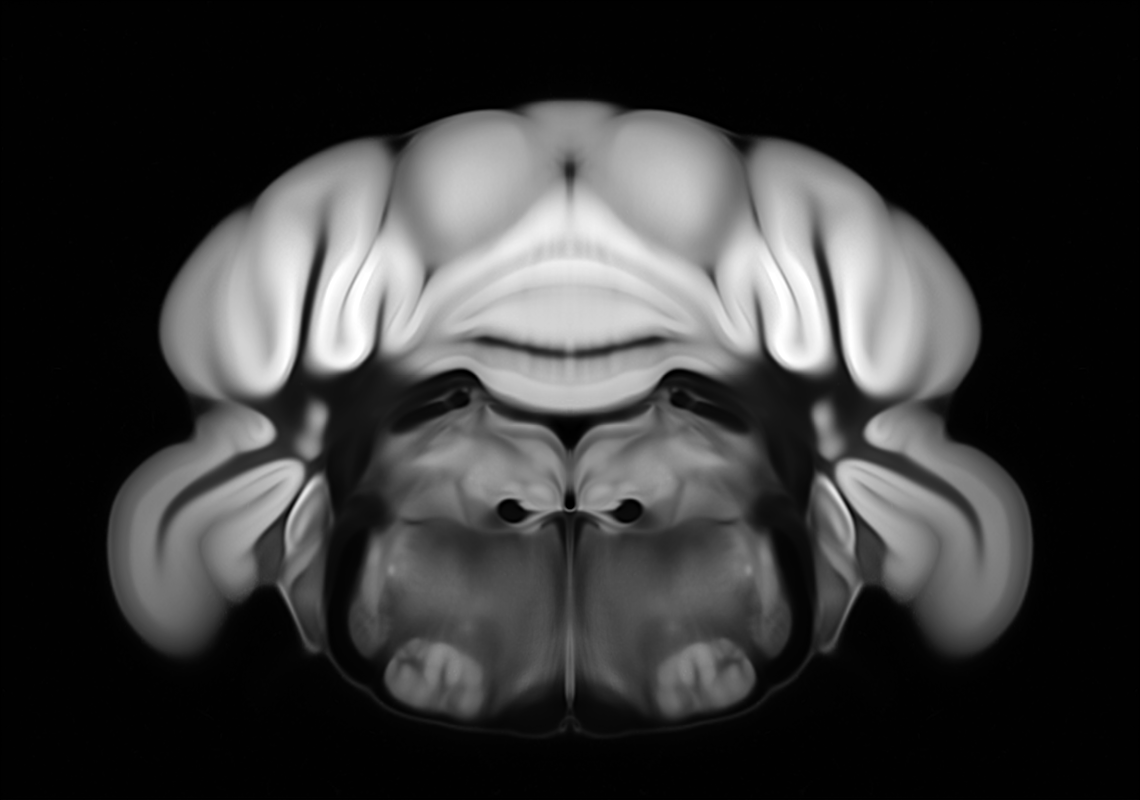

### AllenCCF_Z101.tif

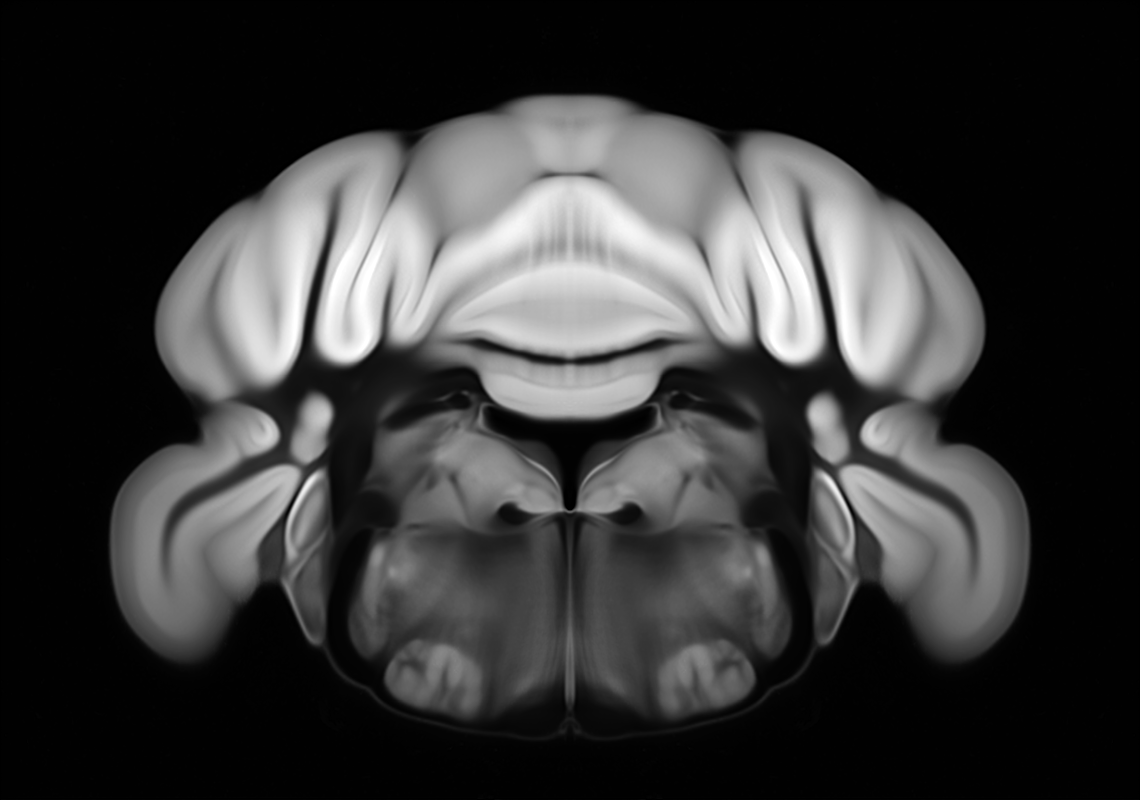

### AllenCCF_Z102.tif

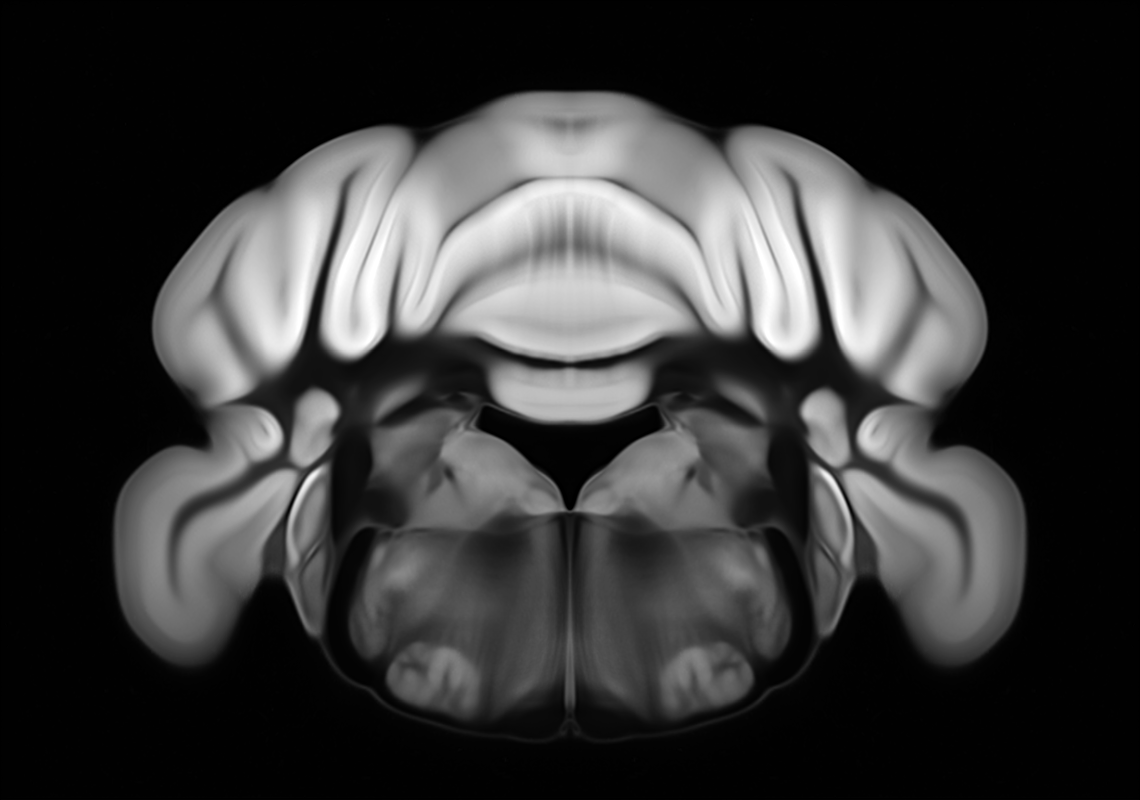

### AllenCCF_Z103.tif

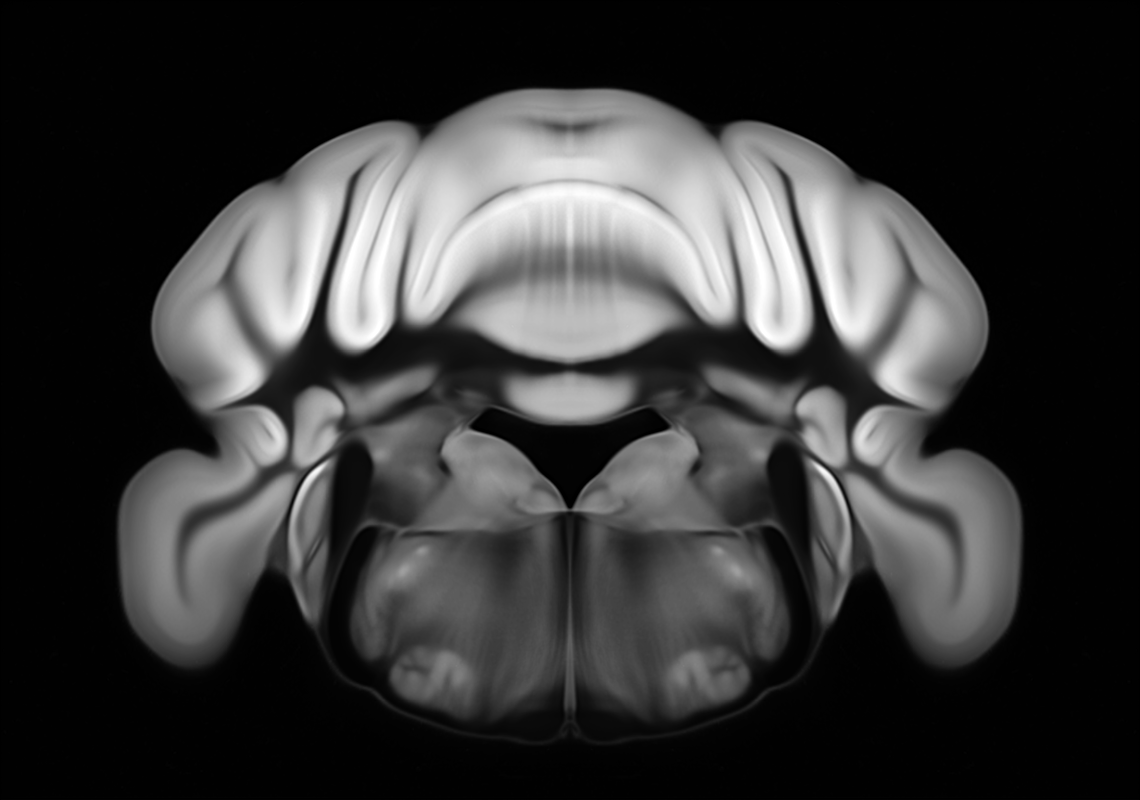

### AllenCCF_Z104.tif

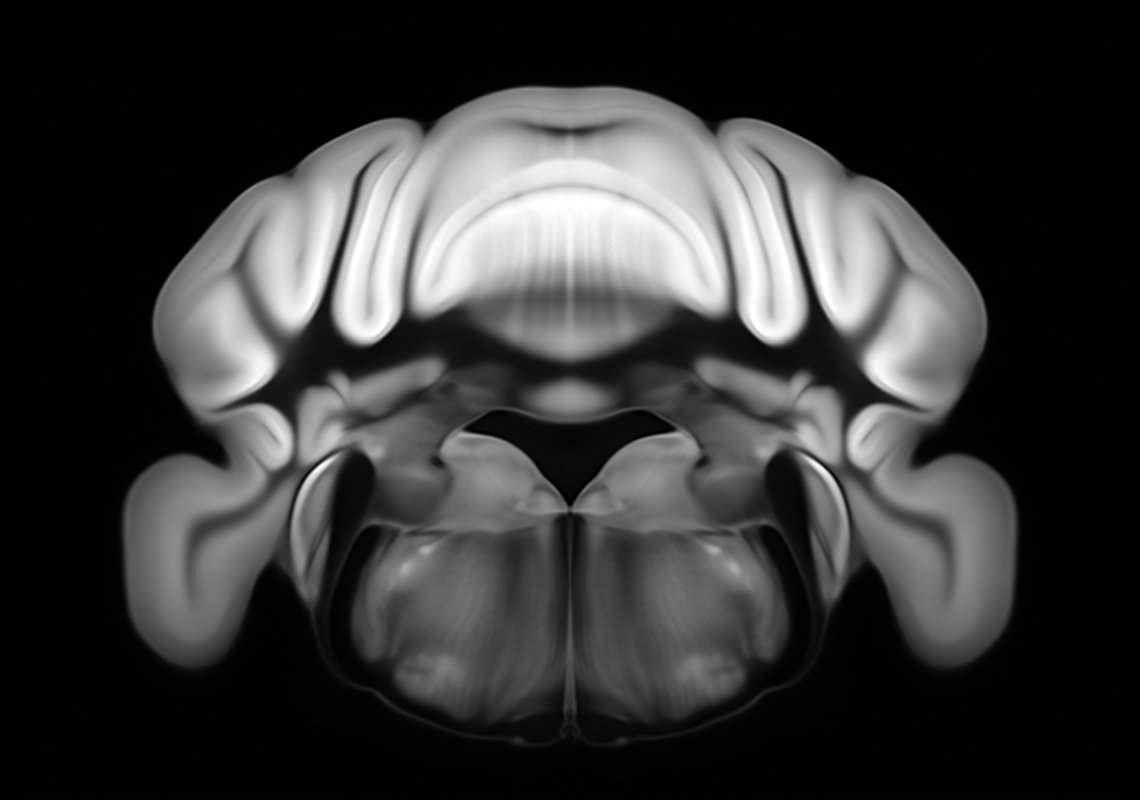

### AllenCCF_Z105.tif

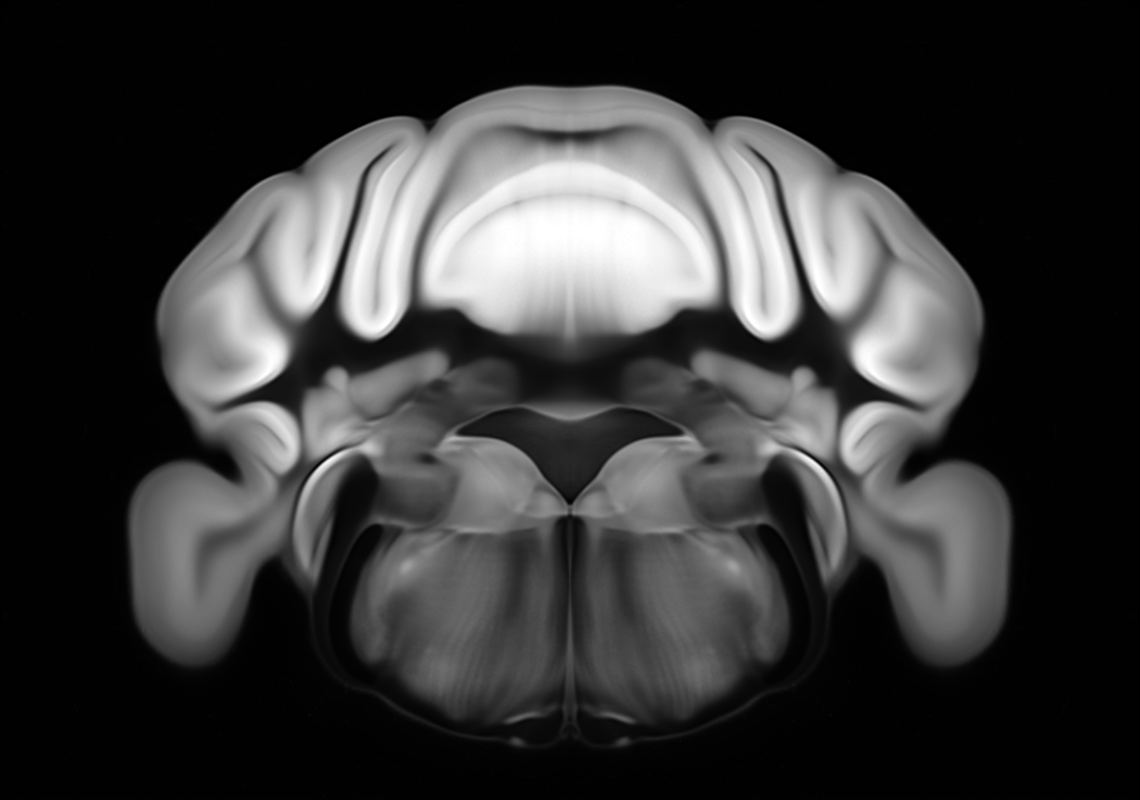

### AllenCCF_Z106.tif

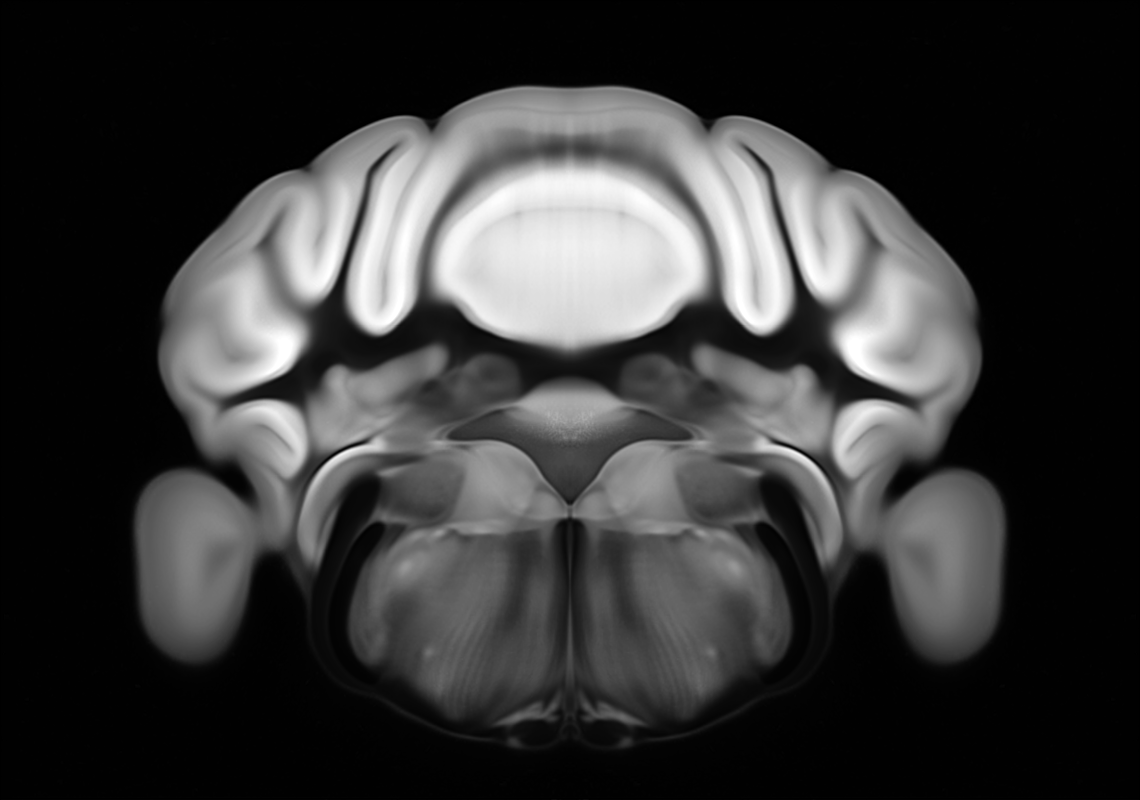

### AllenCCF_Z107.tif

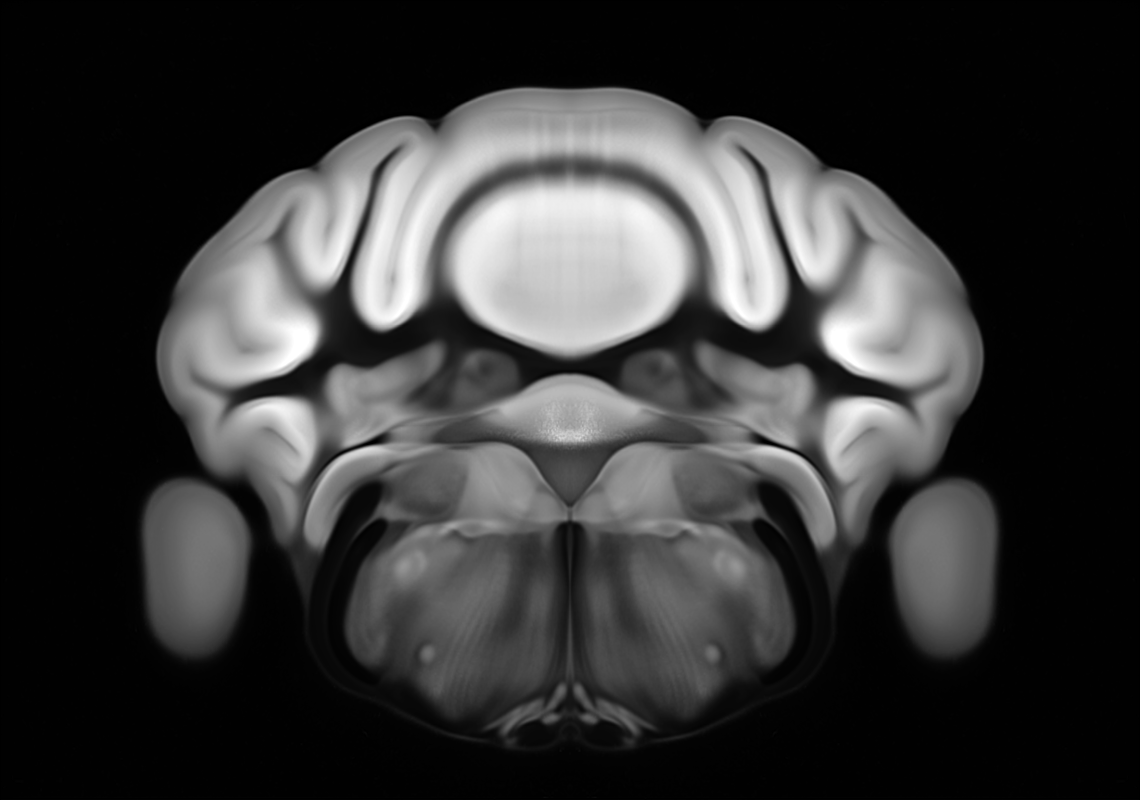

### AllenCCF_Z108.tif

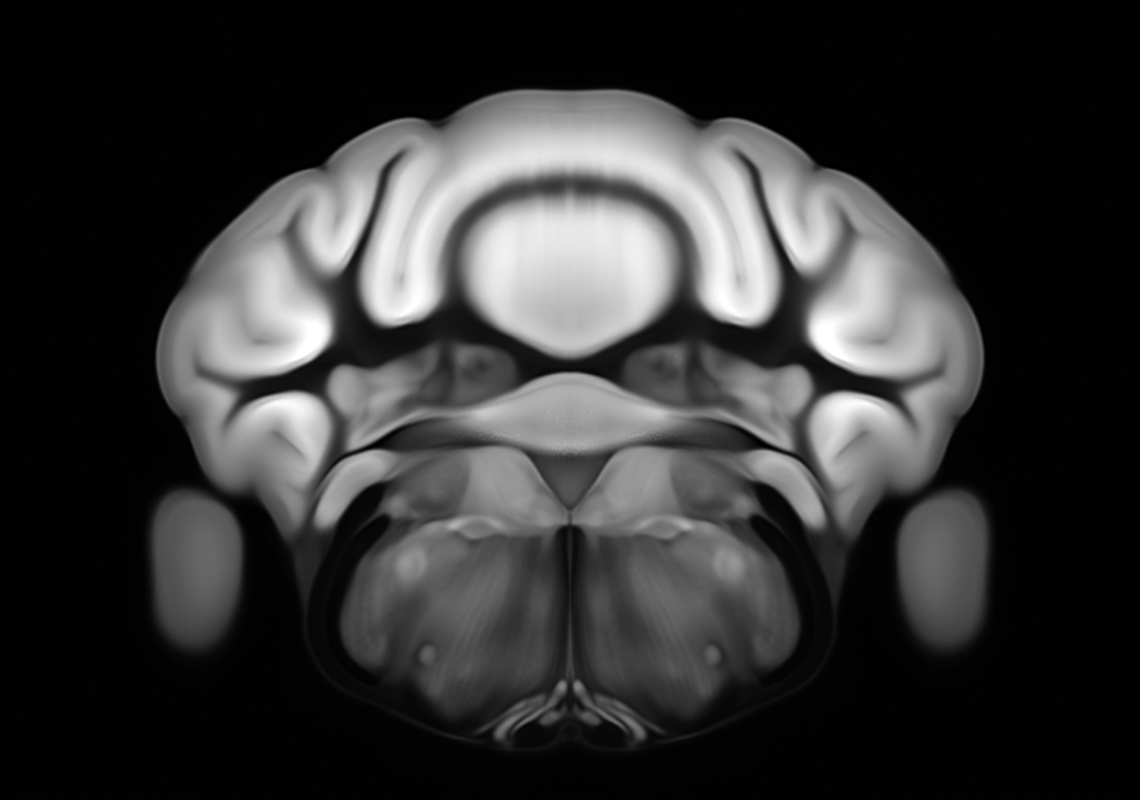

### AllenCCF_Z109.tif

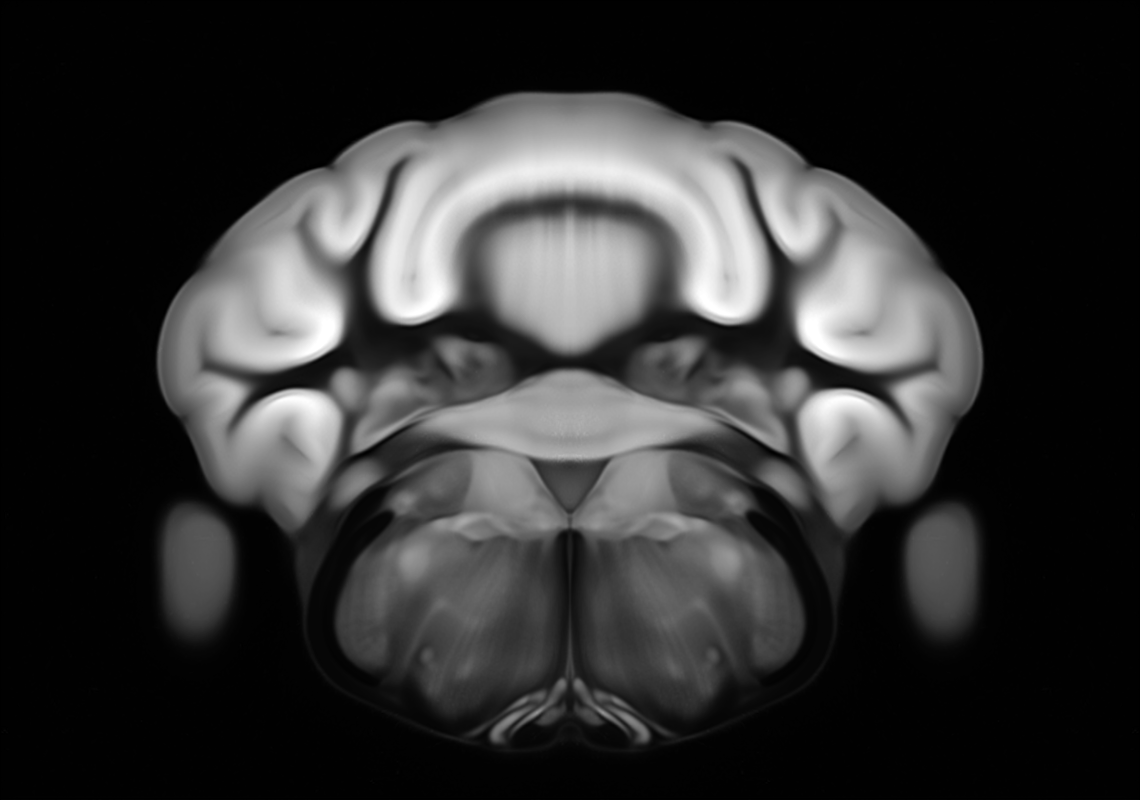

### AllenCCF_Z110.tif

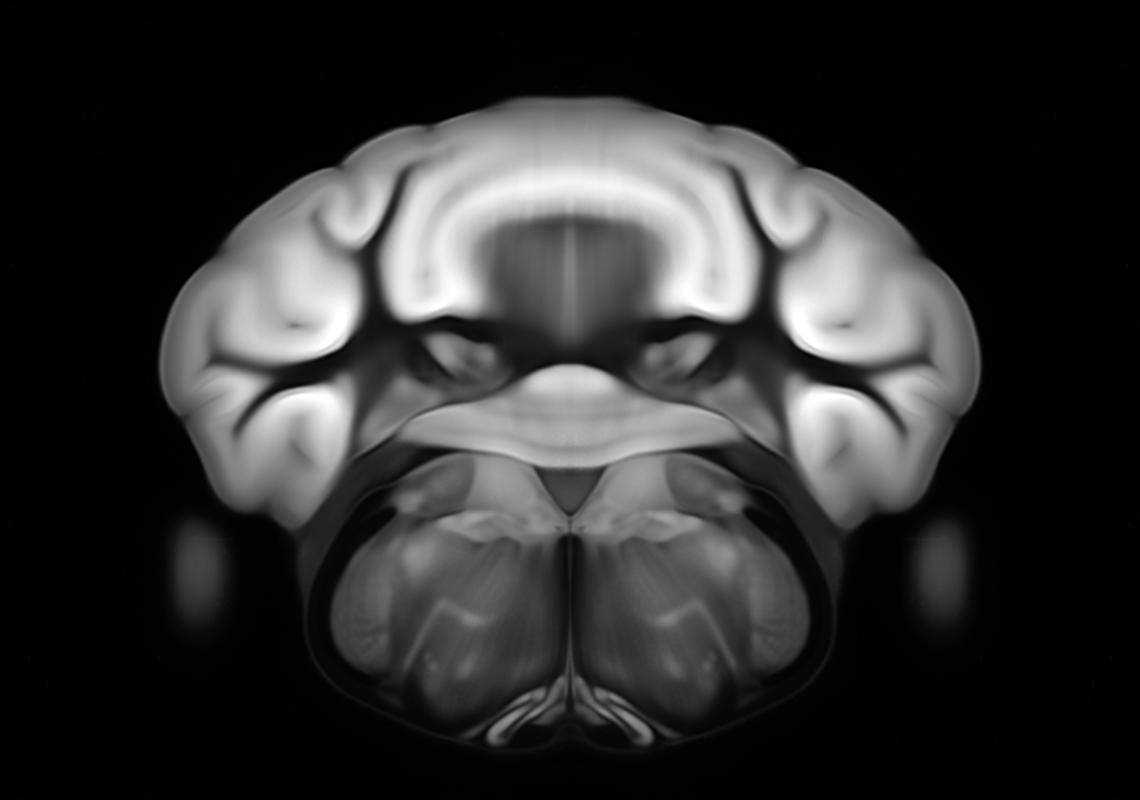

### AllenCCF_Z111.tif

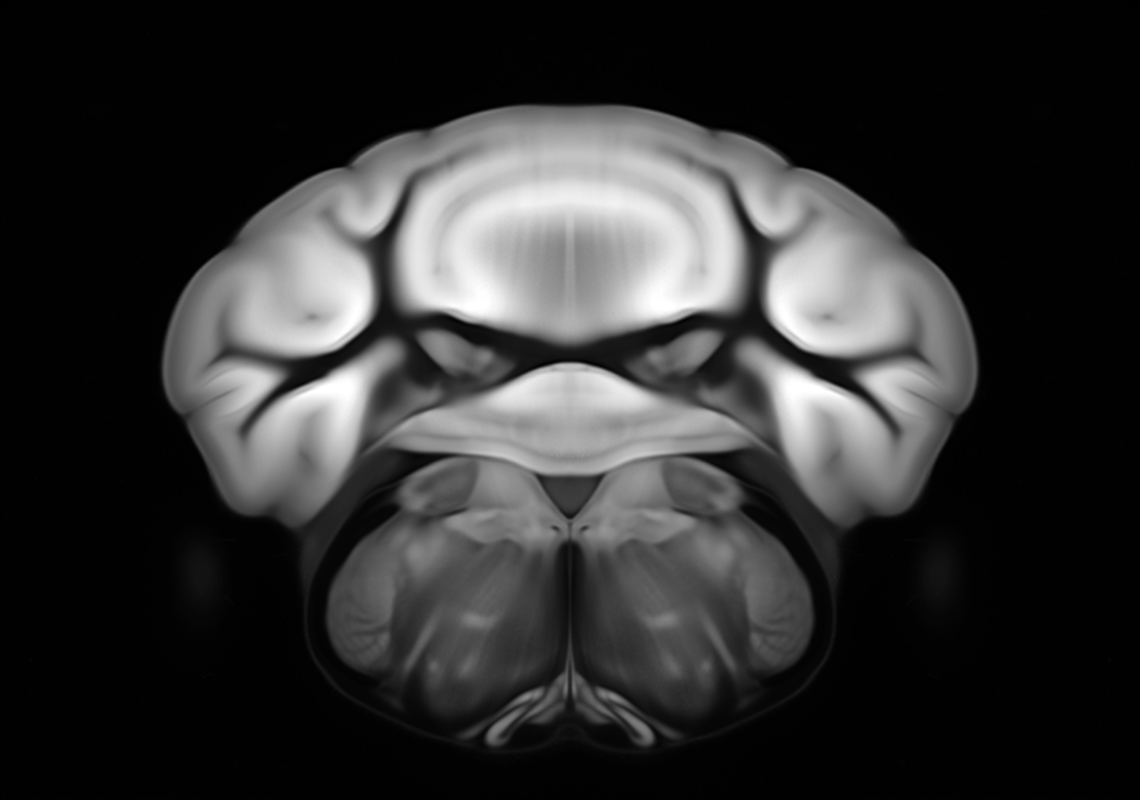

### AllenCCF_Z112.tif

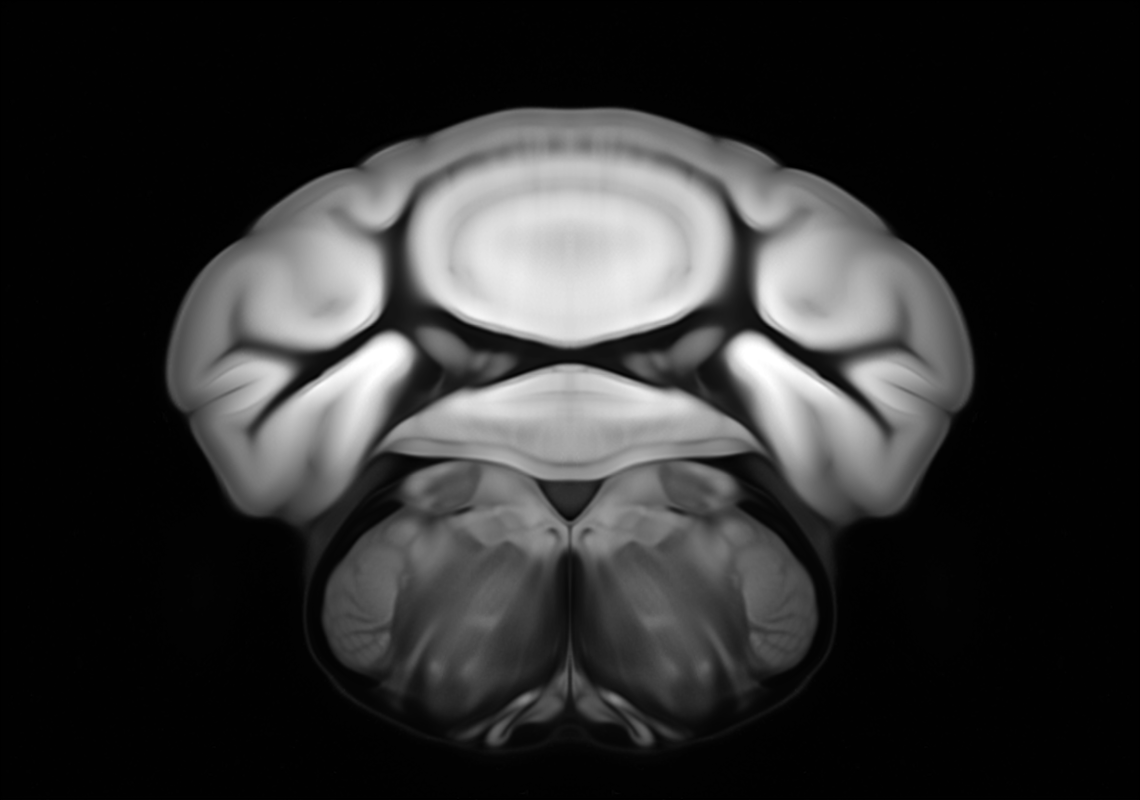

### AllenCCF_Z113.tif

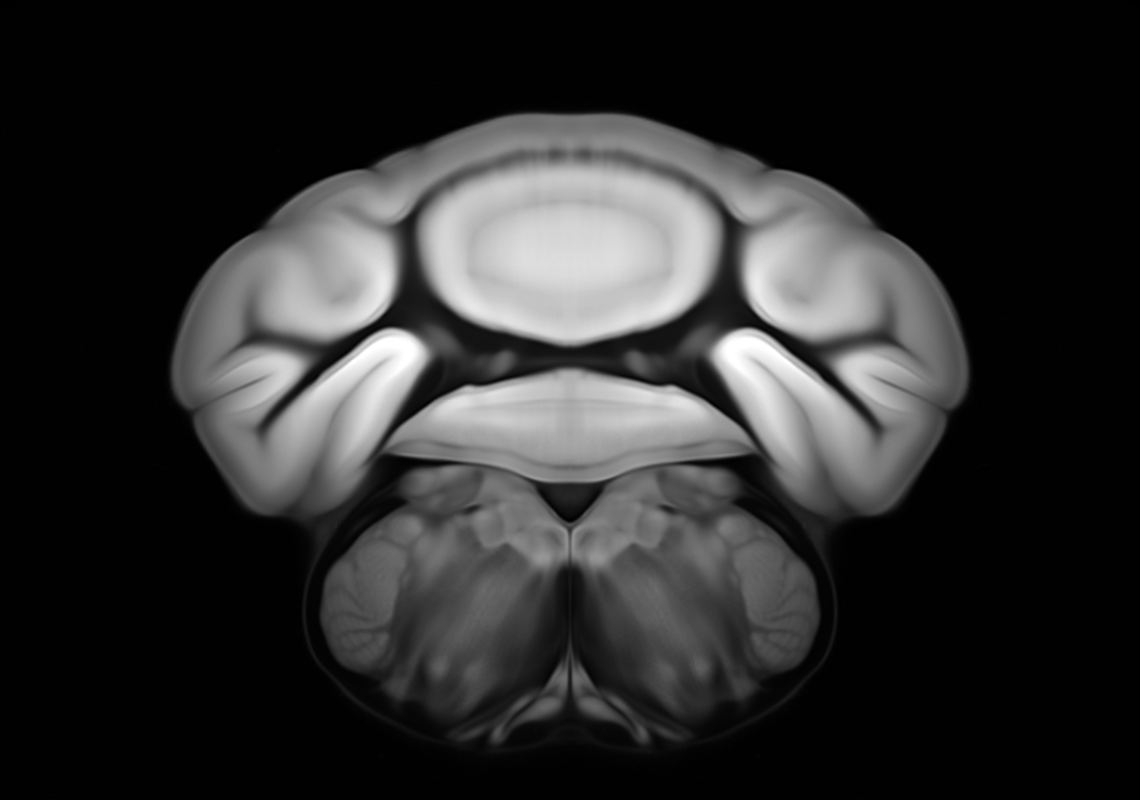

### AllenCCF_Z114.tif

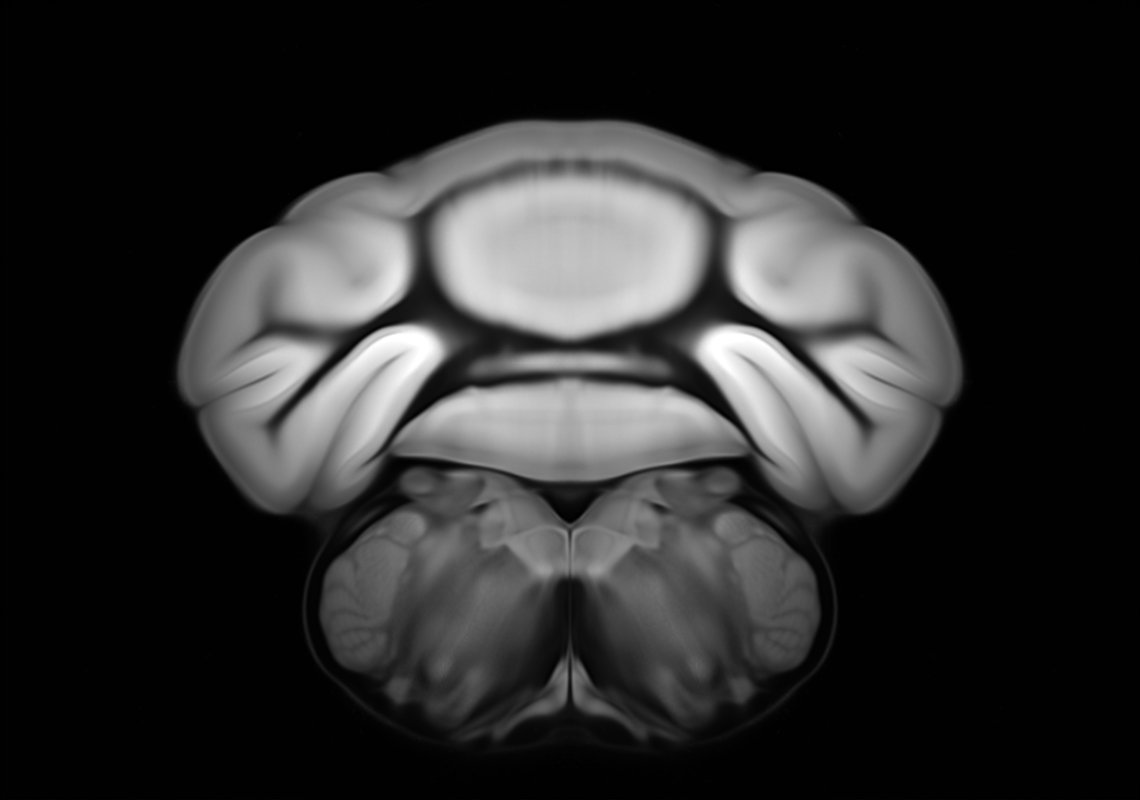

### AllenCCF_Z115.tif

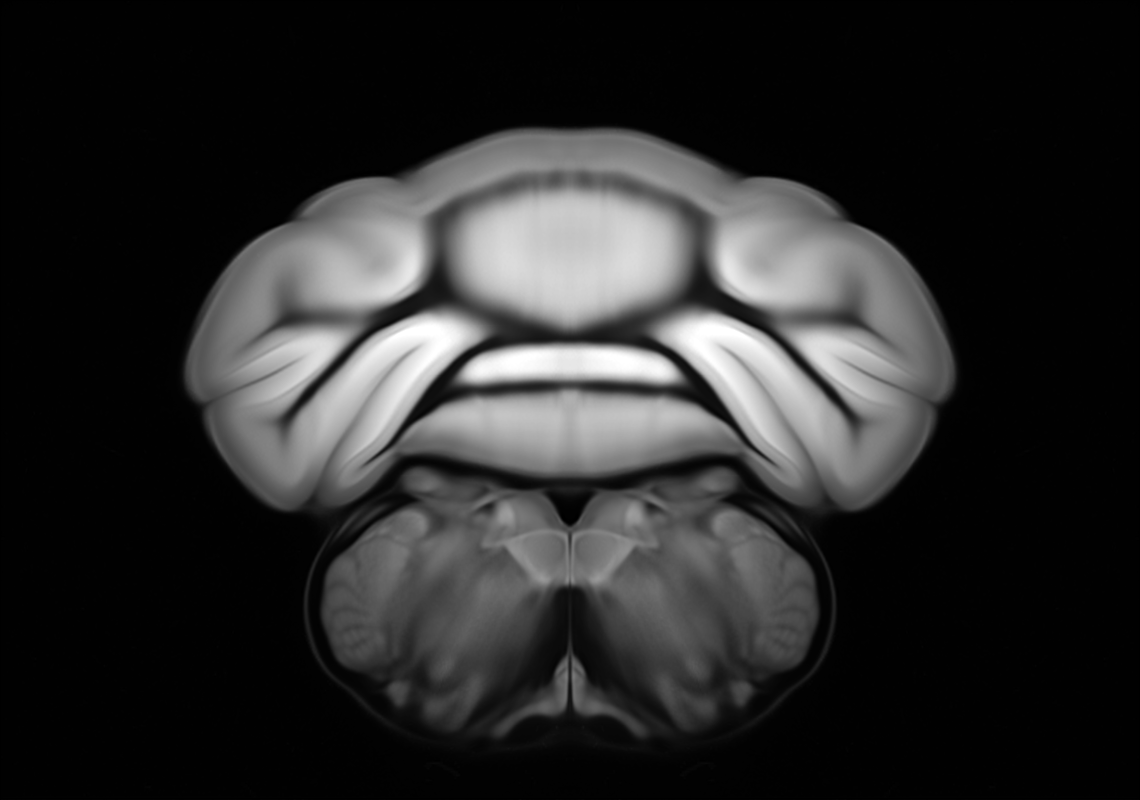

### AllenCCF_Z116.tif

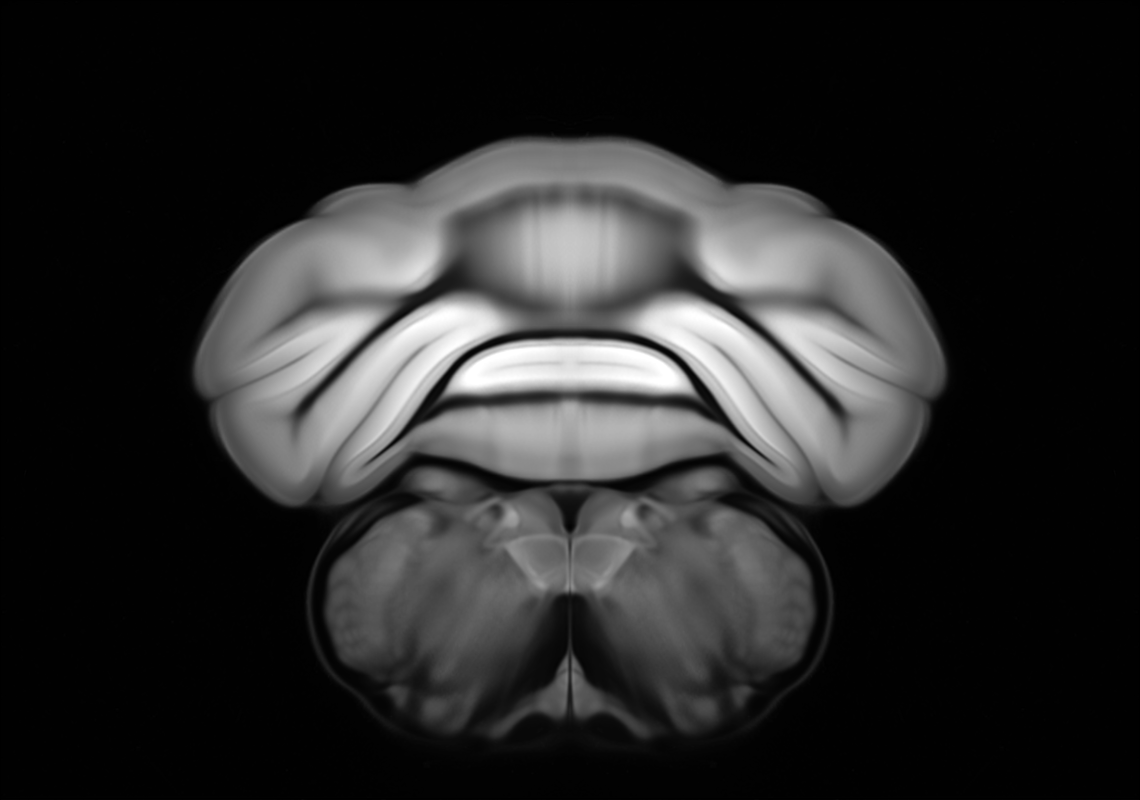

### AllenCCF_Z117.tif

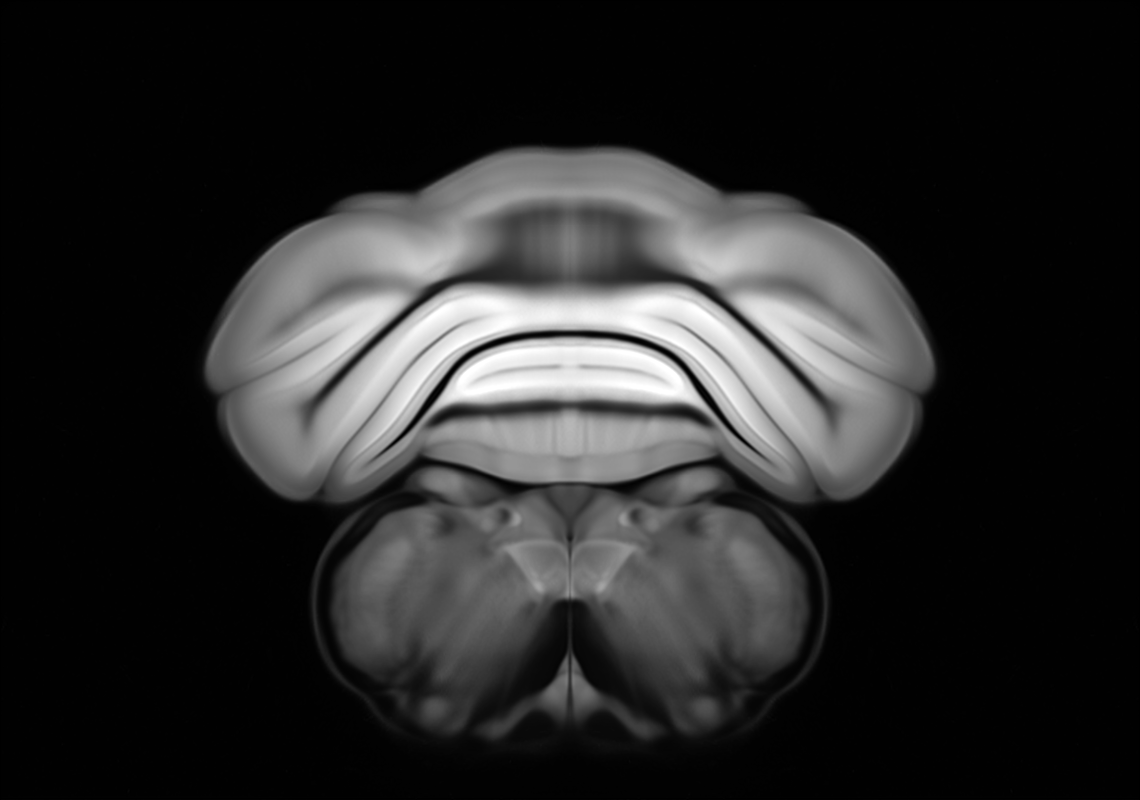

### AllenCCF_Z118.tif

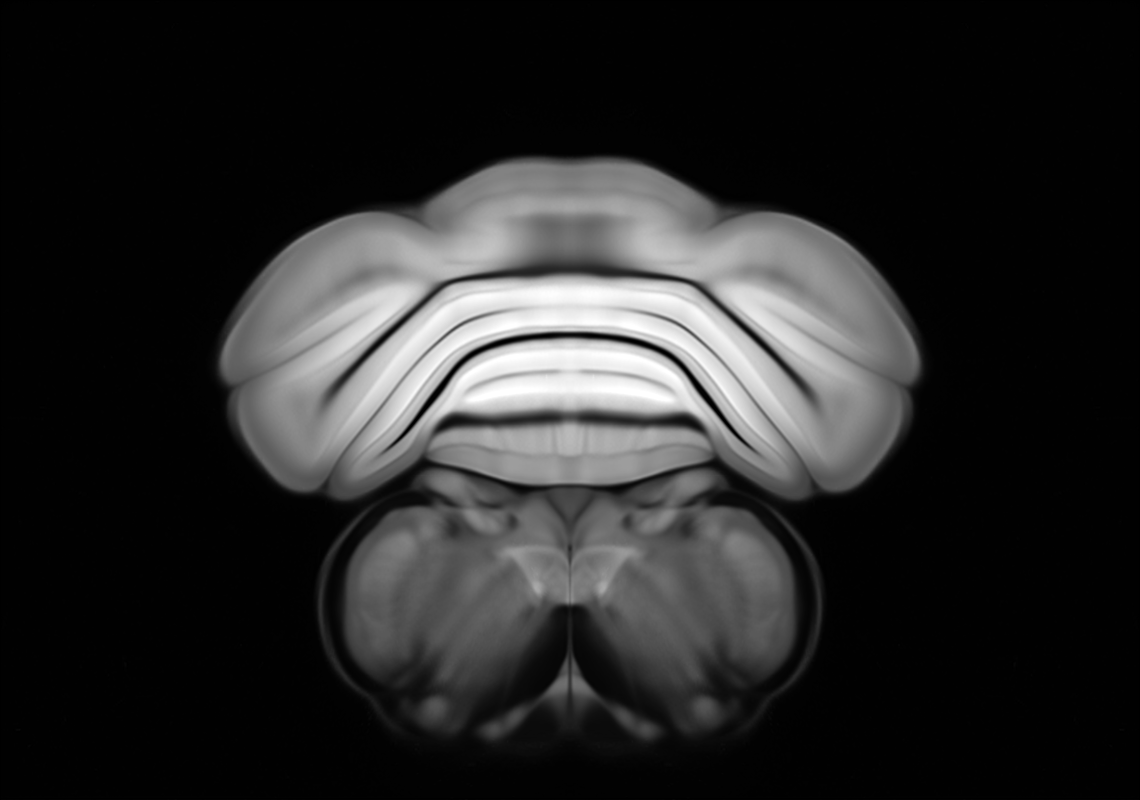

### AllenCCF_Z119.tif

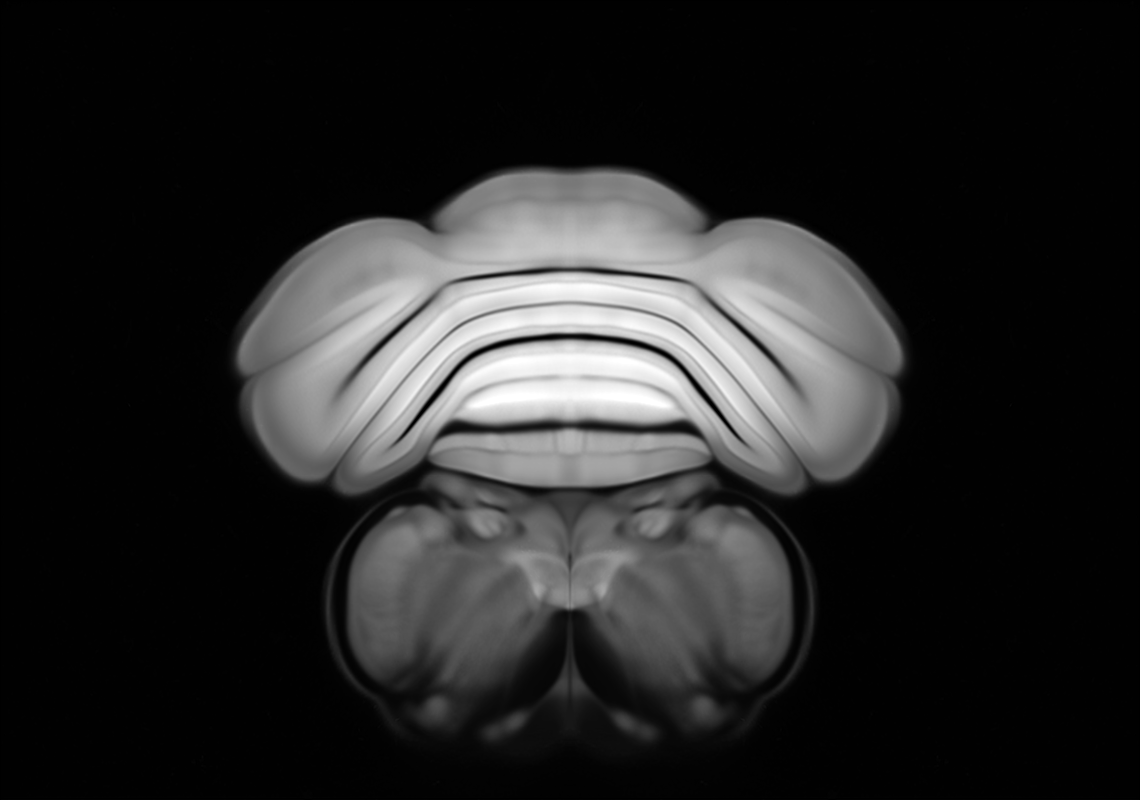

### AllenCCF_Z120.tif

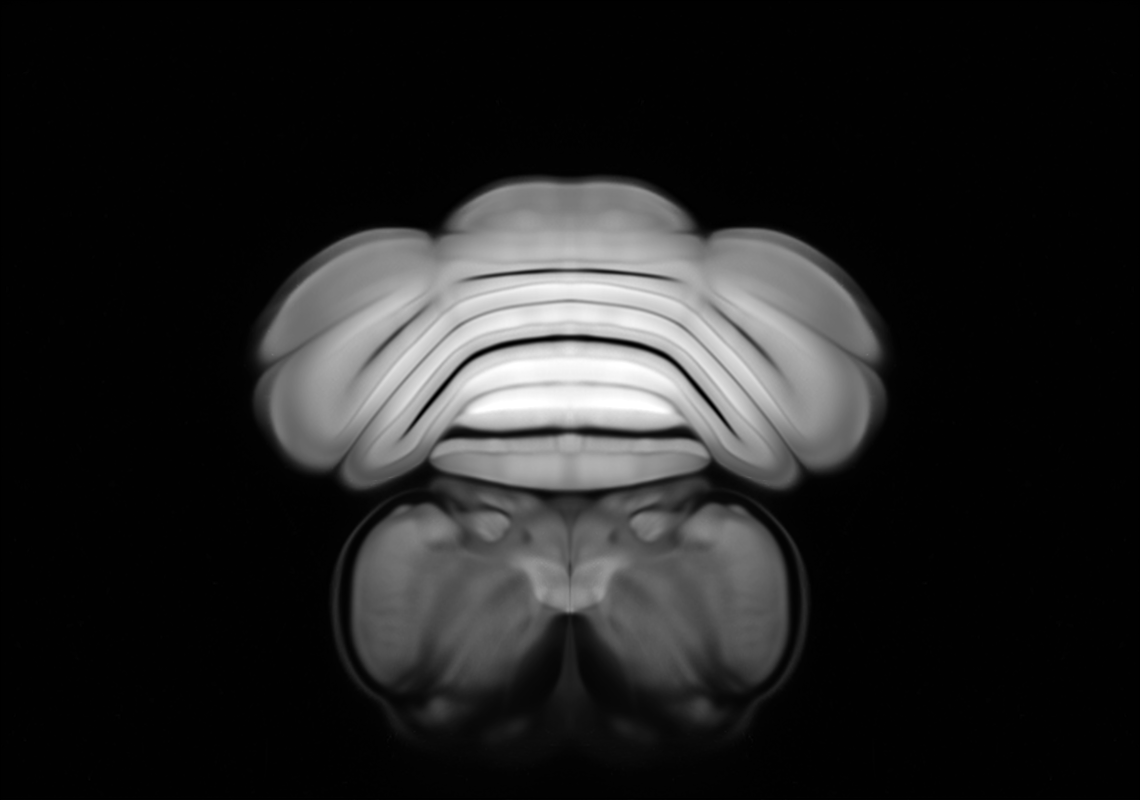

### AllenCCF_Z121.tif

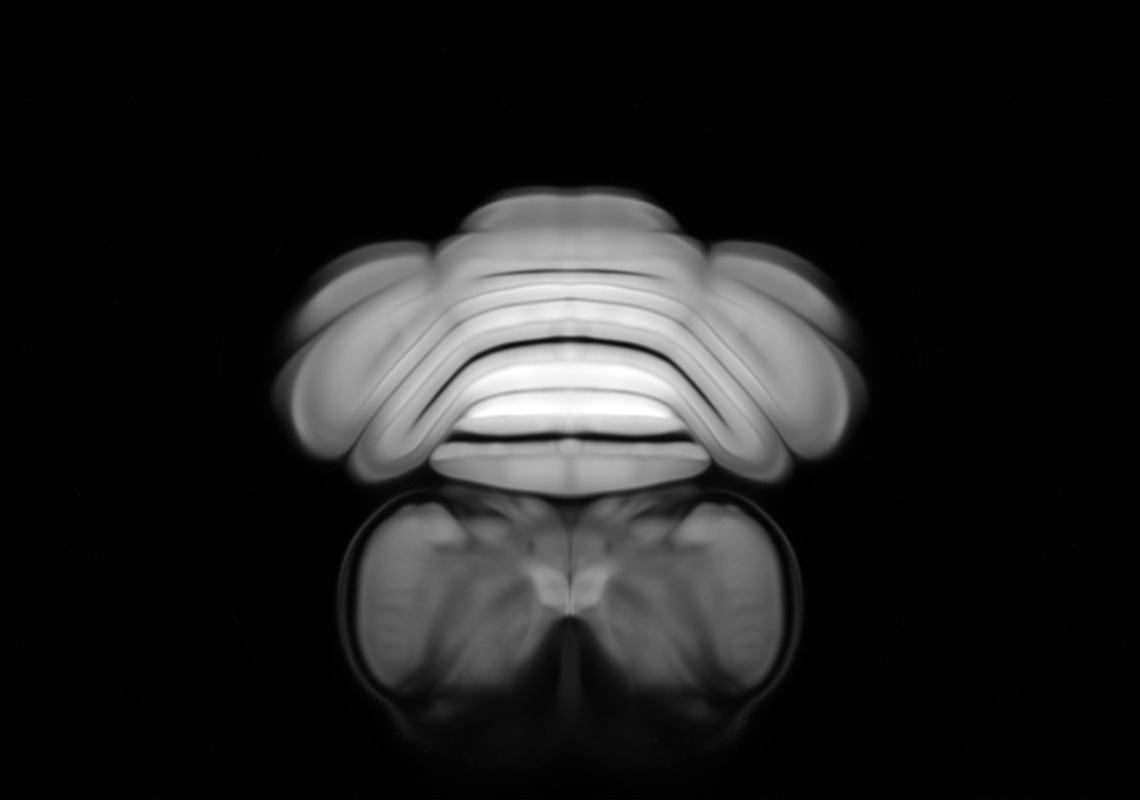

### AllenCCF_Z122.tif

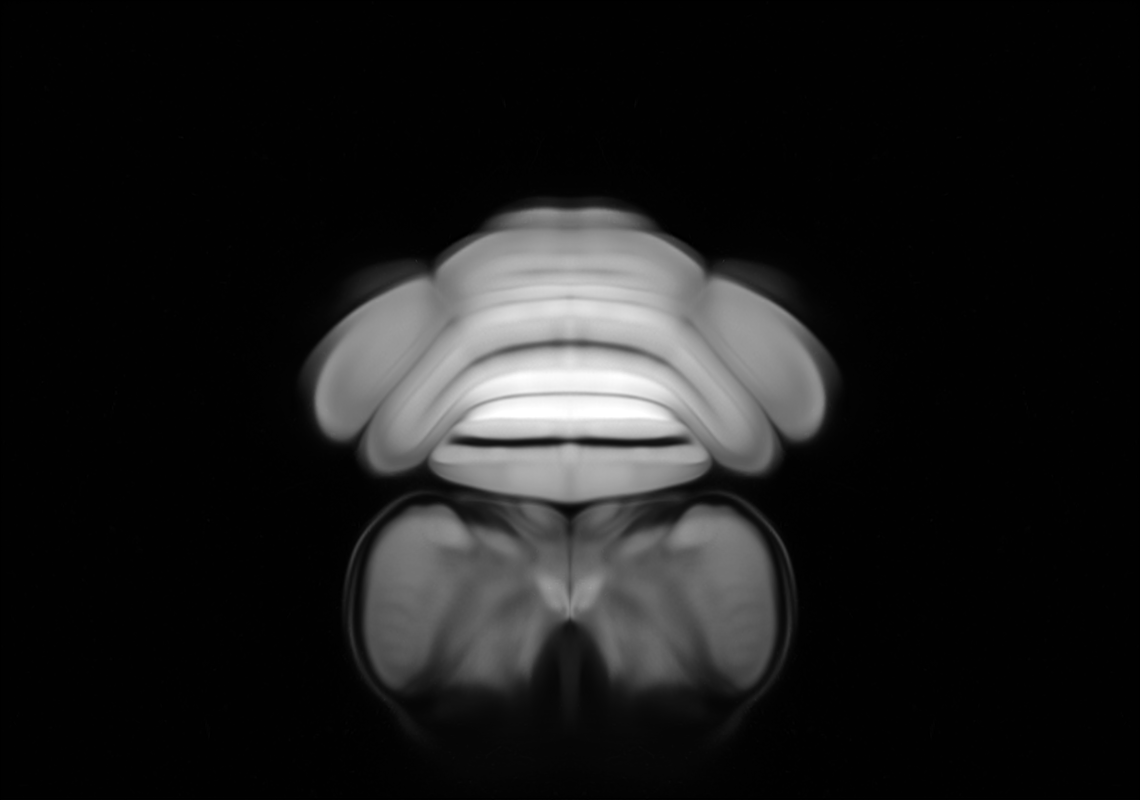

### AllenCCF_Z123.tif

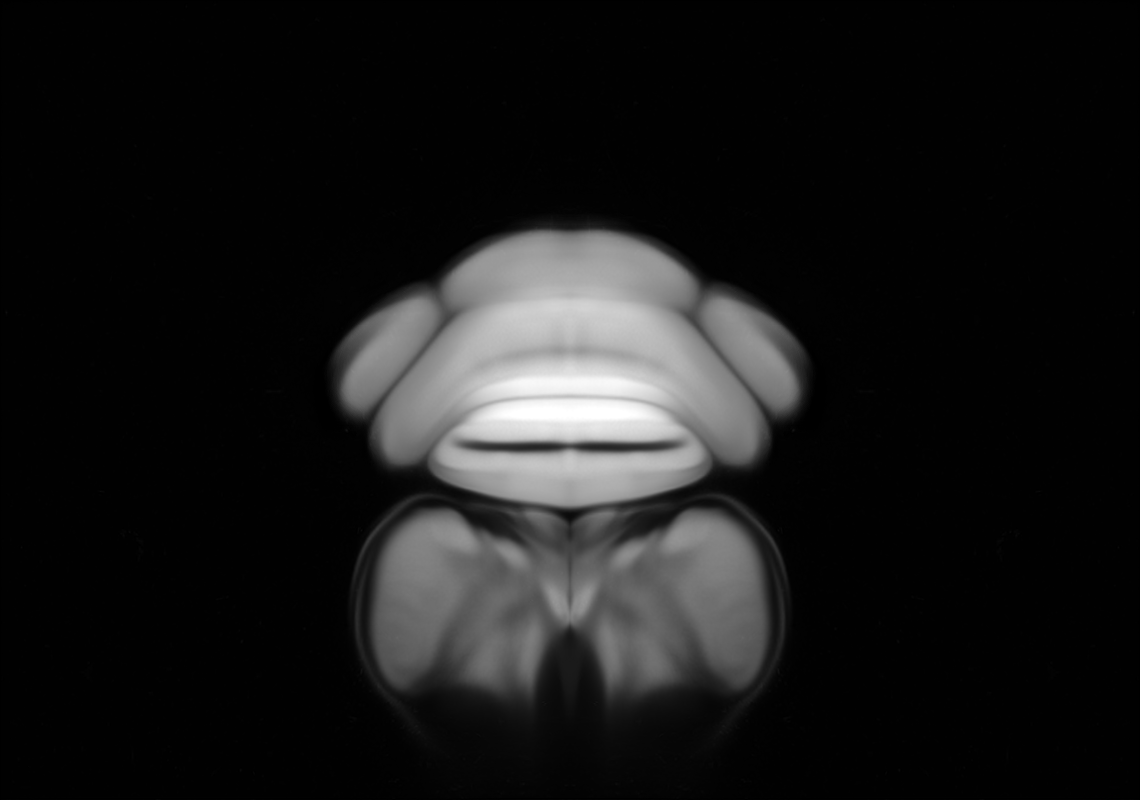
